# Linker histone dH1K27 dimethylation marks *Drosophila* heterochromatin independently of H3K9 methylation

**DOI:** 10.1101/2021.06.22.449135

**Authors:** Jordi Bernués, Andrea Izquierdo-Boulstridge, Oscar Reina, Lucía Castejón, Elena Fernández-Castañer, Núria Leal, Nancy Guerrero-Pepinosa, Carles Bonet-Costa, Olivera Vujatovic, Fernando Azorín

## Abstract

Post-translational modifications (PTMs) of histones are important epigenetic determinants and specific core histones PTMs correlate with functional chromatin states. However, despite linker histone H1s are heavily post-translationally modified, little is known about the genomic distribution of H1s PTMs and their association with epigenetic chromatin states. Here, we address this question in *Drosophila* that encodes a single somatic linker histone, dH1. We previously reported that dH1 is dimethylated at K27 (dH1K27me2). Here, we show that dH1K27me2 is a major PTM of *Drosophila* heterochromatin. At mitosis, dH1K27me2 accumulates at pericentromeric heterochromatin, while, in interphase cells, it is also detected at intercalary heterochromatin. ChIPseq experiments show that dH1K27me2 enriched regions cluster at both the assembled and unassembled heterochromatin regions of all four *Drosophila* chromosomes. More than 98% of the dH1K27me2 enriched regions map to heterochromatic repetitive DNA elements, including transposable elements, simple DNA repeats and satellite DNAs. We also show that dH1K27me2 is independent of H3K9 methylation, as it is equally detected in flies carrying a H3K9R mutation. Moreover, dH1K27me2 is not affected by depletion of Su(var)3-9, HP1a and Su(var)4-20. Altogether these results suggest that dH1K27me2 is a novel epigenetic mark of *Drosophila* heterochromatin that acts upstream of the major Su(var)3-9/HP1a pathway of heterochromatin formation.

## INTRODUCTION

The eukaryotic genome is structured in the form of chromatin, which is built up by the monotonous repetition of a fundamental unit, the nucleosome. Nucleosomes are formed by about 147 bp of DNA wrapped around a protein core particle formed by two copies of each of the core histones (H2A, H2B, H3 and H4). Onto this elemental unit a fifth linker histone H1 binds at the entry/exit point of the DNA, interacting with an additional 15-50 bp DNA fragment (reviewed in (1)). Core histones are subjected to a plethora of post-translational modifications (PTMs) that affect nucleosome structure and function in an epigenetic way (reviewed in(2)). In this regard, several functional chromatin states have been described on the basis of specific core histones PTMs and chromatin associated proteins (3,4). In particular, the silent gene-poor heterochromatic regions are enriched in H3K9me2/3 and H4K20me3, and in Heterochromatin Protein 1 (HP1). As a matter of fact, a main pathway for heterochromatin formation involves H3K9 methylation by the histone methyl-transferase (HMT) Su(var)3-9 and subsequent binding of HP1 (reviewed in (5)) that, in its turn, mediates recruitment of the Su(var)4-20 HMT that methylates H4K20 (6,7). However, it is appealing how so little is known about linker histones H1 PTMs and their potential correlation with functional epigenetic states (8).

Histone H1 is usually present in multiple isoforms in vertebrates (up to 11 in mouse and human) subdivided into somatic and germline variants (reviewed in (9)). In *Drosophila*, there is only a somatic variant (dH1) and a germline variant (BigH1) (10,11). All isoforms present a similar structure containing N- and C-terminal unstructured tails, and a central globular domain. In mice tissues and in human MCF7 and HeLa cells, many PTMs have been identified in the somatic H1 isoforms including phosphorylation of S and T, and acetylation, formylation, methylation and ubiquitilation of K (12). Additionally, di-methylation of K and methylation of R were also described in mice (12). Notably, in both species these PTMs were present all along the H1 sequence (not restricted to the N- and C-terminal tails) and all isoforms were shown to bear many PTMs (12). However, their genomic distribution and functional relevance remain largely unknown. Similarly, in *Drosophila*, there is a lot of information on core histones PTMs but little information on linker histones PTMs (8). In this regard, multiple PTMs were reported in the somatic *Drosophila* dH1 linker histone in Schneider S2 cells (13). Amongst them, dimethylation of lysine 27 (dH1K27me2) in the N-terminal tail was identified. Here, we further characterize this modification. We show that dH1K27me2 is a major PTM of *Drosophila* heterochromatin that is detected throughout development in both interphase cells and mitotic chromosomes. We also show that dH1K27me2 occurs in heterochromatin independently of H3K9 methylation, suggesting that it acts upstream of the Su(var)3-9/HP1a pathway for heterochromatin formation.

## MATERIALS AND METHODS

### Fly stocks and antibodies

dH1^RNAi^ has been described elsewhere (14). UAS-dH1::GFP and UAS-dH1K27A::GFP stocks were prepared by inserting the corresponding ORFs (WT or carrying a K27A mutation generated by PCR) into pEGFP (Clontech) to generate fusion proteins and then the inserts were cloned into pUAST (details can be provided on request). Constructs were injected into *Drosophila w^1118^* embryos and transgenic flies were obtained. H2AvD2::GFP flies were kindly provided by Dr Saint (15). yw; ΔHisC, 2xUAS-YFP; 12xHWT and yw; ΔHisC, 2xUAS-YFP; 12xK9R flies were kindly provided by Dr Duronio (16). w^1118^, NubGal4, UAS-Dic2 and UAS-Su(var)4-20^RNAi^ (BL-32892) fly stocks were obtained from Bloomington. UAS-HP1a^RNAi^(V31995) and UAS-Su(var)3-9^RNAi^ (V101494) were obtained from VDRC (Vienna).

Primary antibodies used were rabbit αdH127me2 (described in (13)), rat αHP1a (described in (17)) and rabbit αdH1 (generously provided by Dr Kadonaga and described in (14)). All other antibodies were commercially available: mouse αGFP (Roche, 11814 460 001), rabbit αGFP rabbit (life technologies, A11122), rabbit αH3K9me2 (Millipore, 07-441), and rabbit αH4K20me3 (Abcam, Ab9053). Secondary antibodies conjugated to Cyanine 2, 3 and 5 and to HRP were from Jackson.

### Cell culture and mitotic chromosomes preparations

*Drosophila* cell lines S2 and Kc167 were grown under standard conditions in Schneider’s medium supplemented with 10% fetal calf serum at 25°C. For metaphase chromosomes preparation 5 mL-flasks were grown to 3×10^6^ cells/mL. Then 240 μL of Colcemid (Roche, at 10 μg/mL) were added and incubated for 6 h at 25°C. To 1 mL of hypotonic buffer (IMAC: 50 mM glycerol, 5 mM KCl, 10 mM NaCl, 0,8 mM CaCl_2_, 10 mM sucrose) 300 μL of cells from above were added and incubated for 5 min. This cell suspension was impacted (200 μL) on slides using a Cytospin at 500 rpm for 10 min. Slides were allowed to dry at room temperature for 1 h and fixed in 4% *p*-formaldehyde (10 min), washed in PBS and then in PBS-T (PBS-0.2% Tween 20-0.1% BSA). Then αdH1K27me2 (1:200) or αdH1 (1:4000) in PBS-T was added and incubated 1h in a dark moist chamber at room temperature, washed with PBS-T and then treated with secondary α-rabbit-Cy3 antibody (1:400 in PBS-T) for 45 min. After washing with PBS-T and PBS, samples were mounted in MOWIOL-0.2ng/μL DAPI and visualized on a Leica SPE confocal microscope.

For neuroblast preparation, 3-5 brains were dissected in 0.7% NaCl and washed twice in the same solution. Brains were incubated at room temperature with 100 μL Colcemid (at 10 μg/mL with 0.7% NaCl) in a dark moist chamber for 2 h. After washing in PBS, they were treated with dissociation buffer (0.54 mL 0.8% Na-citrate, 0.06 mL 10Xdispase/collagenase solution) for 10 min at 37°C. Then, they were fixed in 4% *p*-formaldehyde for 30 min at room temperature. Brains were transferred to an ethanol-clean coverslip containing 16 μL PBS, covered with a slide and squashed (beat with a needle onto the coverslip until disaggregation). Slides were immersed in liquid nitrogen, coverslips were removed and chromosomes were immunostained with αdH1K27me2 (1:200) or αdH1 (1:4000) as above. Images were recorded with a Leica SPE confocal microscope.

### Polytene chromosome preparations

Salivary glands from *Drosophila w^1118^* strain or from the indicated crosses were dissected from 3^rd^ instar larvae and polytene chromosomes prepared as described in (18). Primary antibodies used were: mouse αGFP (1:50), rabbit αGFP(1:400), rabbit αdH1 (1:4000), rat αHP1a (1:200), rabbit αdH1K27me2 (1:100), rabbit αH4K20me3 (1:100), and rabbit αH3K9me2 (1:100). Secondary antibodies (Jackson) were all used at 1:200. Images were recorded with a Leica SPE confocal microscope. For comparisons identical settings were used.

### Chromatin Immunoprecipitation (ChIP)

To a 25 mL-flask containing about 10^8^ S2 cells, fresh formaldehyde was added until a final concentration of 1.8% and incubated for 10 min at room temperature. Reaction was quenched by adding glycine up to 0.125M and further incubating for 10 min at room temperature. Cells were harvested with a cell scraper and spun down. Cell pellets were resuspended in 5mL PBS and, subsequently, washed with 10 mL Lysis Buffer A (10mM Hepes pH 7.9, 10mM EDTA, 10mM EGTA and 0.25% Triton X100), spun and washed again with 10 mL Lysis Buffer B (10 mM Hepes pH 7.9, 100 mM NaCl, 1mM EDTA, 0.5mM EGTA and 0.01% Triton X100). A last washing step was done adding 4.5mL TE and 0.5mL SDS 10%. After spinning, pellets were resuspended in TE and PMSF was added up to 1mM in a final volume of 4 mL where 40 μL of 10% SDS was added. Sonication was performed with BioRuptor by 2 sessions of 10 min with 30s ON and 30s OFF, on ice. Lysates were pooled in one tube, which was supplemented to obtain 1% Triton X100, 0.1% Deoxycholate and 140 mM NaCl solution and incubated 10 min at 4°C. After spinning, supernatants containing chromatin were recovered to use them for the immunoprecipitation. To check DNA size, 100 μL of sonicated chromatin was de-crosslinked by adding SDS and NaHCO_3_ to a final concentration of 1% and 0.1M respectively, and incubating overnight at 65°C. Then, regular DNA extraction with phenol/chloroform was performed. Subsequently, DNA precipitation was done with 100% ethanol and 30 μL of 3M sodium acetate followed by a washing step with 70% ethanol. Pellet was resuspended in 10 μL H_2_O and incubated 30 min at 37°C with 0.5μL RNAse A for RNA clearance. DNA sample was finally loaded on a 1.5% agarose gel to check DNA fragment size. Average size of DNA in these samples was ∼250 bp.

For chromatin immunoprecipitation, 500 μL aliquotes of cross-linked chromatin were pre-cleared with 50 μL of Protein A Sepharose beads (PAS) resuspended in RIPA buffer at 50% (140 mM NaCl, 10 mM Tris-HCl pH8, 1mM EDTA, 1% Triton X100, 0.1% SDS and 0.1% Deoxycholate) and incubated for 1h at 4°C. After centrifugation, supernatants were collected and incubated with 1 μL of αdH1K27me2 overnight at 4°C in a rotary wheel. Subsequently, 40 μL of PAS suspension were added to the samples and incubated for 3h at 4°C. Washing steps with RIPA (x5), LiCl ChIP buffer (250 mM LiCl, 10 mM Tris-HCl pH 8, 1 mM EDTA, 0.5% NP- 40 and 0.5% Deoxycholate) (x1) and TE (x2) were performed (10 min each). Beads were incubated with 40 μL TE and 0.5 μL RNAse A (10 mg/mL) for RNA clearance (30 min, 37°C). Antibody-DNA conjugates were eluted by addition of 50 μL 0.2M NaHCO_3_ and 10 μL 10% SDS. After centrifugation, supernatants were collected. Beads were re-extracted twice with 100 μL of Elution Buffer (1% SDS and 0.1M NaHCO_3_) and supernatants pooled (300 μL). A sample input of 50 μL of cross-linked chromatin was resuspended in Elution Buffer to obtain a final volume 300 μL. Both IP samples and Inputs were incubated overnight at 65°C. Then 3 μL of Proteinase K (at 10 mg/mL) were added and samples were incubated 3h at 56°C. DNA was phenol/chloroform extracted and precipitated overnight at −20°C. Finally, DNA pellets were resuspended in 15 μL H_2_O. DNA concentrations were determined by Qbit fluorometric quantitation. For ChIPseq experiments samples were then sent to an external facility for high-throughput Illumina sequencing. For ChIP-qPCR experiments, real-time qPCR analysis from samples prepared as described above was performed in triplicate using the Light Cycler system (Roche) and a Light Cycler 480 apparatus as described in (19). Primers used in these experiments are described in **Supplementary Table I**.

### ChIPseq analysis

Analysis of ChIPseq data was performed essentially as described (20). Quality of original FASTQ sequences were assessed using FASTQC v0.11. FASTQ files were aligned against the dm3 genome assembly using Bowtie2 (v2.2.2) using options -n 1 and replicates were pooled into a single BAM file. Potential overamplification artifacts were detected and removed using sambamba v0.5.1. IP vs Input enrichment assessment was performed using the htSeqTools R package version 1.12.0. and TDF coverage tracks were generated using IGVTools 2. Peak calling between duplicate filtered IP and Input samples was performed using MACS 1.4.2, using options -g dm and the resulting peaks were annotated against the dm3 genome using ChIPPeakAnno 3.16.1. Asessment of the presence of repeat and repetitive elements among called peaks was determined using RepeatMasker v4.3.0, with options -no-is and -species drosophila. Identified peak sequences were scanned for known and denovo motifs using Homer version 2.

A cluster analysis of read similarity was performed with those reads that failed to align in both ChIP samples and Input, by means of the CD-Hit software (v 4.5.4) (21). First, reads were grouped into a single Fasta file and flagged with their corresponding tag (ChIP or Input). Then, reads were sorted decreasingly by length and a sequence clustering was performed using a greedy incremental algorithm. The longest read became the representative of the first cluster and the remaining reads were compared to the representatives of existing clusters. If the similarity with any representative was above the given thresholds, it was grouped into that cluster. Otherwise, a new cluster was defined with that read as the representative. The thresholds used were sequence identity and alignment coverage. The sequence identity threshold used was the number of identical nucleotides in the alignment divided by the full length of the shorter sequence which should be equal or bigger than 0.95. Secondly, the alignment should cover a 40% of the length of both sequences. Then, the proportion of reads between ChIP samples and Input for each cluster was tested with a hypergeometric test with a Benjamini-Yekuteli adjusted p-value below 0.01.

ChIPseq data is deposited at NCBI GEO (GSE167018).

### Western blot (WB) and peptide dot blot assays

WB and peptide dot blot assays were performed as described in (13). Custom designed *Drosophila* peptides for dot blot assays were: dH1K27unmod (CKKVVQKKASGSAGT) and dH1K27me2 (CKKVVQ[2Me-K]KASGSAGT) were obtained from Caslo (Denmark), and H3K9me3 (residues 1-21) was from Upstate-Millipore. For WB, total extracts were prepared in 100 μL SDS-PAGE protein loading buffer (PLB) from 30-50 salivary glands. Additionally, H1-enriched extracts were also prepared (from either ∼100 μL of staged embryos; from salivary glands, brains and imaginal disks from 30-50 larvae, and abdomens and heads from some 10-20 adult flies) by 5% perchloric acid extraction for 1h at 4°C with rotation. Extracts were spun for 5 min at 20,000xg at 4°C, supernatants were recovered and material containing linker histone H1 was precipitated by addition of trichloroacetic acid up to 20% and 10 min spinning as above. Pellets were washed once with cold acetone/1M HCl (9:1 v/v) and twice with pure acetone. Pellets were allowed to dry completely at room temperature and finally they were resuspended in PLB. Antibodies used were: αdH1K27me2 (1:2000), αdH1 (1:20000), and αGFP (1:2000). Secondary HRP-conjugated antibodies were always used at 1:10000 (Jackson) and developed with ECL (G&E biochemicals). For quantification, films were scanned with a GS300 laser microdensitometer (Bio-Rad) and quantitative analysis was carried out using FiJi software.

## RESULTS

### dH1K27me2 is detected throughout development and across cell types

To analyze dH1K27me2 content across fly development we performed WB and IF analyses using αdH1K27me2 antibodies. Specificity of the αdH1K27me2 antibodies used in these experiments was shown in transgenic flies carrying UAS-constructs expressing either a dH1::GFP fused protein or a mutated dH1K27A::GFP form, which carries a K27A mutation that prevents methylation. Both constructs were expressed in salivary glands using a *Nub-*GAL4 driver. WB analysis of salivary glands extracts showed that both constructs were expressed to similar levels as detected using αGFP (**Supplementary Figure S1A**, right panel) and IF studies showed similar patterns of localization in polytene chromosomes (**Supplementary Figure S1B**). However, while αdH1K27me2 antibodies detected dH1::GFP in WB, they failed to detect the mutated dH1K27A::GFP protein (**Supplementary Figure S1A**, left panel). In addition, previous *in vitro* dot-blot assays using dH1 peptides containing K27me2, K28me2 or unmethylated K27 showed high selectivity of αdH1K27me2 for the methylated dH1K27me2 peptide (13) (see also **Supplementary Figure S1C**, left). Furthermore, αdH1K27me2 showed no significant reactivity against an H3 peptide carrying K9me3 (**Supplementary Figure S1C**, right panel). During embryo development, dH1 expression is zygotic being first detected around 2h after egg laying concomitant with the activation of the zygotic genome (ZGA) (11). We observed that dH1K27me2 appears as early as dH1 expression is detected (**Figure 1A**, lanes 1-4). Later, dH1K27me2 was also detected in third instar larvae in brains, imaginal disks and salivary glands, as well as in head and abdomen of adult male and female flies (**Figure 1B**). Notably, relative to total dH1, dH1K27me2 levels appeared rather constant in all fly tissues tested (**Figures 1A** and **1B**, bottom panels).

**Figure 1.**
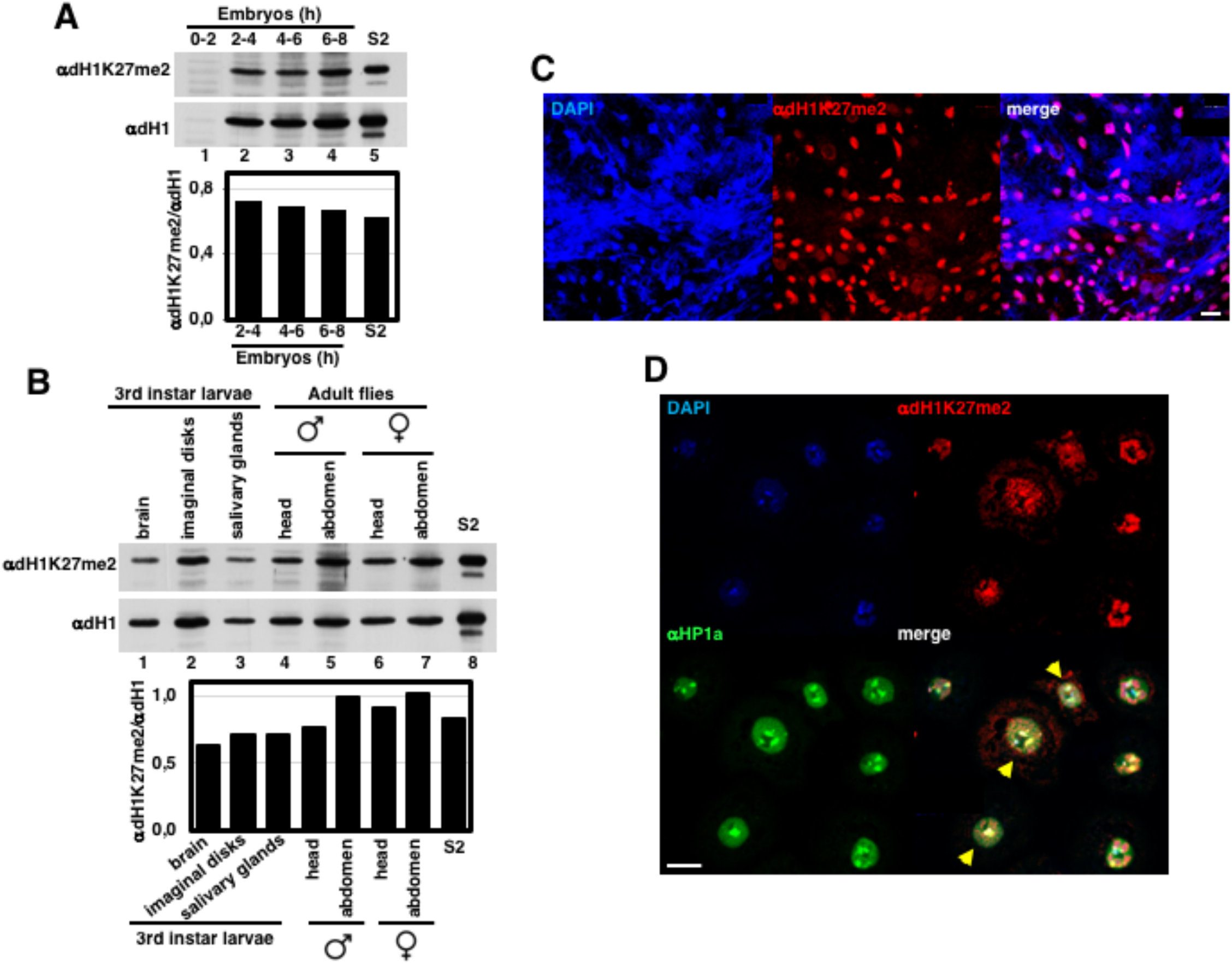
Developmental expression of dH1meK27me2. **A**) WB analysis with αdH1 and αdH1K27me2 of perchloric acid (PCA) extracts from *w^1118^* embryos staged after egg laying for the indicated time periods (lanes 1-4). Lane 5 shows a PCA extract from S2 cells as control. Quantification of the results is shown at the bottom as the dH1K27me2/dH1 ratio. **B**) WB analysis with αdH1 and αdH1meK27me2 of PCA extracts from the indicated *w^1118^* larval organs (lanes 1-3), and head and abdomen of adult male (lanes 4 and 5) and female (lanes 6 and 7) flies. Lane 8 shows a PCA extract from S2 cells as control. Quantification of the results is shown at the bottom. **C**) Immunostaining of *w^1118^* larval brain squashes with αdH1K27me2 is shown in red. DNA was stained with DAPI (in blue). Scale bar corresponds to 20μm. **D**) Immunostaining of interphase S2 cells with αdH1K27me2 (in red) and αHP1a (in green). DNA was stained with DAPI (in blue). Yellow arrows indicate nuclei in which αdH1K27me2 signal accumulates at αHP1a foci. Scale bar corresponds to 20μm.

In good agreement with these results, IF experiments performed in brain squashes of third instar larvae detected intense nuclear αdH1K27me2 immunostaining in interphase cells (**Figure 1C**). Similarly, in cultured S2 cells, a roughly uniform nuclear αdH1K27me2 immunostaining was observed in interphase (**Figure 1D**). Interestingly, αdH1K27me2 signal was generally detected at HP1a foci that mark pericentromeric heterochromatin (**Figure 1D**), where it highly concentrated in some cells (**Figure 1D**). Intense αdH1K27me2 immunostaining was also detected in the specialized interphase polytene chromosomes of salivary glands (**Figure 2A**). Notably, dH1K27me2 largely co-localized with HP1a at the chromocenter (**Figures 2A** and **2B**), which corresponds to the fused pericentromeric heterochromatin regions of all four *Drosophila* chromosomes. Strong αdH1K27me2 immunostaining was also observed all along the chromosome arms (**Figure 2A**). Overall dH1K27me2 distribution in the arms matched nicely with DAPI intense bands (**Figures 2A** and **2C)**, which have high dH1 content (22–24) and correspond to intercalary heterochromatic regions enriched in transposable elements and other repetitive DNA sequences (25,26). Noteworthy, while an intense HP1a immunostaining was observed on the heterochromatic chromosome 4, dH1K27me2 presence on chromosome 4 was low and mostly restricted to two weak bands that colocalized with DAPI bands (**Figure 2B**). Remarkably, dH1K27me2 was not detected at any other HP1a positive regions outside the chromocenter, namely euchromatic and telomeric regions (**Supplementary Figure S2**).

**Figure 2.**
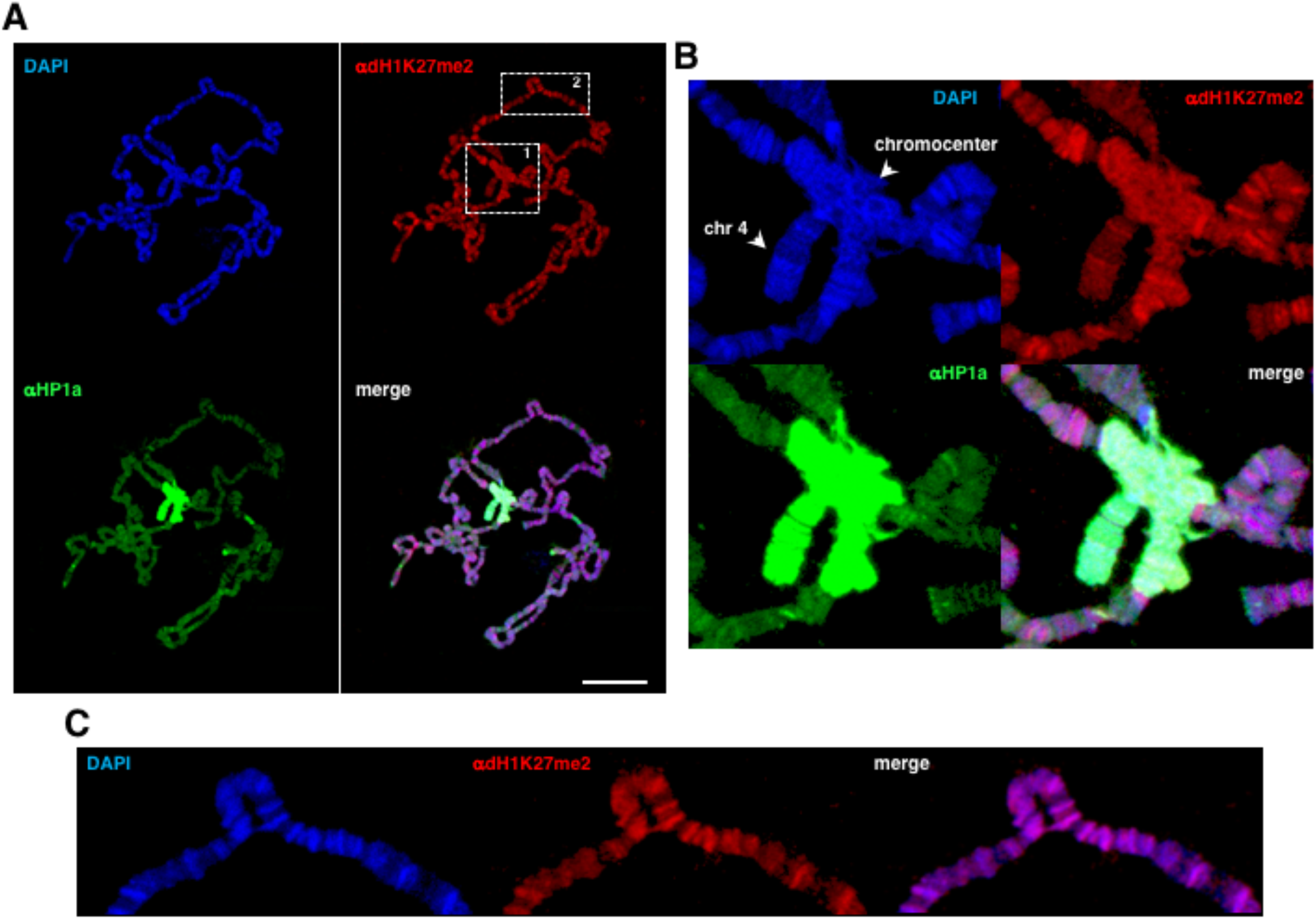
dH1K27me2 pattern in polytene chromosomes. **A**) Immunostaining of *w^1118^* polytene chromosomes with αdH1K27me2 (in red) and αHP1a (in green). DNA was stained with DAPI (in blue). Scale bar corresponds to 20μm. **B**) Enlarged image of region 1 in A. Arrows indicate the chromocenter and chromosome 4. **C**) Enlarged image of region 2 in A showing overlapping of αdH1K27me2 bands (in red) with DAPI bands (in blue).

Altogether these results suggest that dH1K27me2 is present across cell types throughout development, being distributed along chromatin.

### dH1K27me2 accumulates in heterochromatin

Previous results (13) showed that, in S2 cells, αdH1K27me2 immunostaining was largely restricted to pericentromeric heterochromatin in metaphase chromosomes. Here, we further confirmed this observation (**Figure 3B**, top panel) and extended it to metaphase chromosomes of larval neuroblasts (**Figure 3A**, left panel) and to a second cell line (Kc167) (**Figure 3C**). Note that immunostaining with αdH1 antibodies, which recognize total dH1, showed no such enrichment at pericentromeric heterochromatin (**Figures 3A**, right panel, and **3B**, bottom panel). To gain further insights on the genomic distribution of dH1K27me2, we performed ChIPseq analysis in S2 cells. This analysis identified 4,751 dH1K27me2 enriched regions (**Supplementary Table II**). Most of these regions clustered proximal to the pericentromeric heterochromatic regions of chromosomes 2 and 3, in the assembled heterochromatic regions of chromosomes 2 and 3 (chr2L&RHet and chr3L&RHet) and at the unassembled repetitive heterochromatic elements of the artificial chromosomes U and Uextra (**Figure 4A**). Interestingly, only 4 dH1K27me2 enriched regions were detected on the heterochromatic chromosome 4, in good agreement with IFs results reported above (**Figure 2B**). Moreover, although only 38% (N= 1805) of the dH1K27me2 enriched regions could be assigned to any particular chromatin state according to the *Drosophila* 9 chromatin states model (3), the vast majority of those (90%, N= 1603) overlapped with heterochromatin state 7 (**Figure 4B**). This enrichment was statistically significant as shown by permutation analysis (**Figure 4C**), while no significant overlap was observed with any of the other chromatin states even after removing all reads overlapping chromatin state 7. The high proportion of non-assigned (NA) dH1K27me2 enriched regions reflects their high abundance at the unassembled artificial heterochromatic chromosomes U and Uextra, which are not included in the *Drosophila* 9 chromatin states model (3). dH1K27me2 enriched regions usually appeared in clusters and extended for several kilobases (up to ∼70kb) that, in some cases, could be interrupted by regions of low dH1K27me2 content (**Figure 4D**, top panel). Silenced heterochromatic gene clusters, such as the *stellate* locus in pericentromeric heterochromatin of chromosome X, were also found enriched in dH1K27me2 (**Figure 4D**, bottom panel).

**Figure 3.**
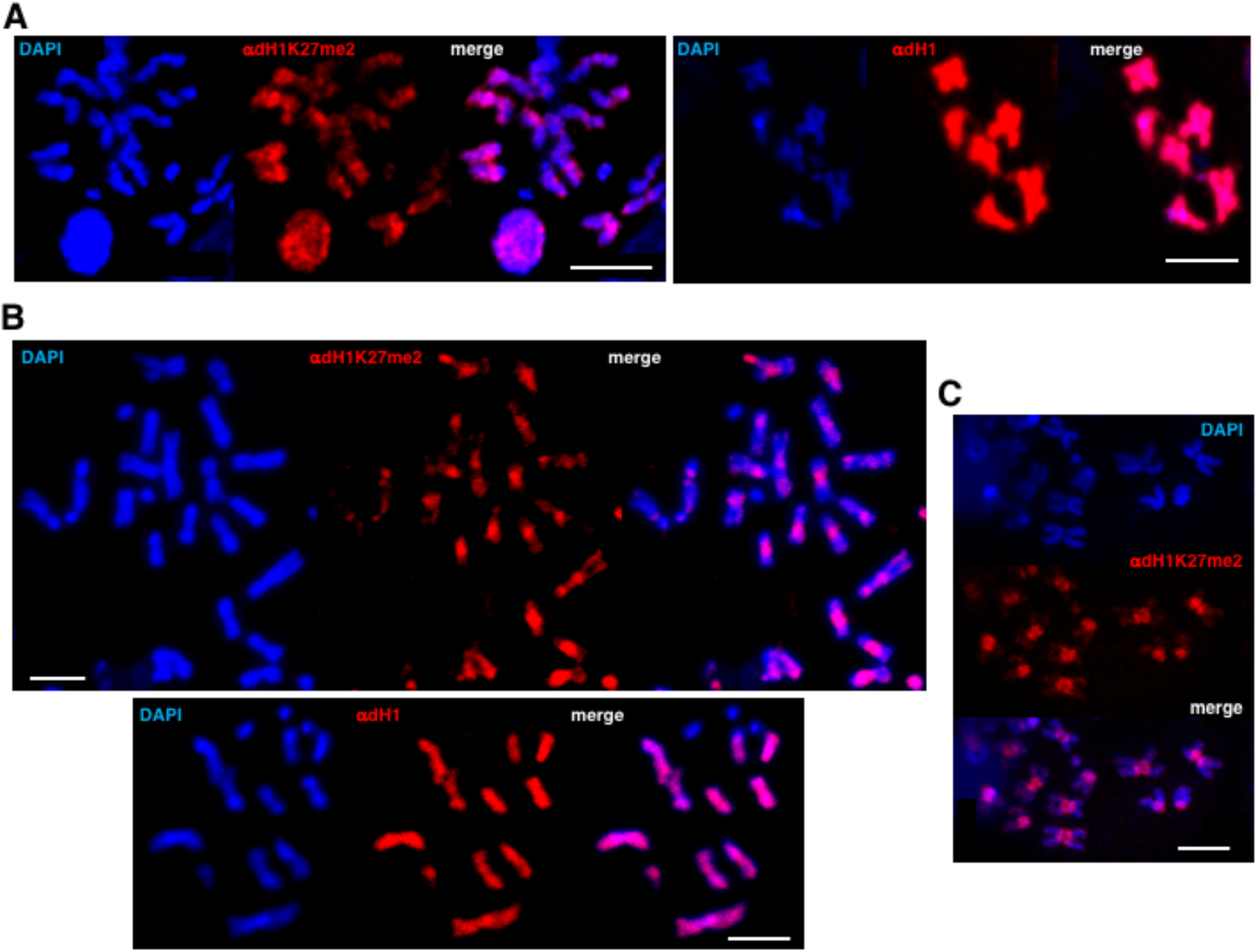
Pericentromeric accumulation of dH1K27me2 in metaphase chromosomes. **A**) Metaphase chromosome spreads from *w^1118^* larval brain squashes immunostained with dH1K27me2 (in red) (left) and αdH1 (in red) (right). DNA was stained with DAPI (in blue). Scale bar corresponds to 20μm. **B**) Metaphase chromosome spreads from S2 cells immunostained with αdH1K27me2 (in red) (top) and αdH1 (in red) (bottom). DNA was stained with DAPI (in blue). Scale bars correspond to 20μm. **C**) Metaphase chromosome spreads from Kc167 cells immunostained with αdH1K27me2 (in red). DNA was stained with DAPI (in blue). Scale bar corresponds to 20μm.

**Figure 4.**
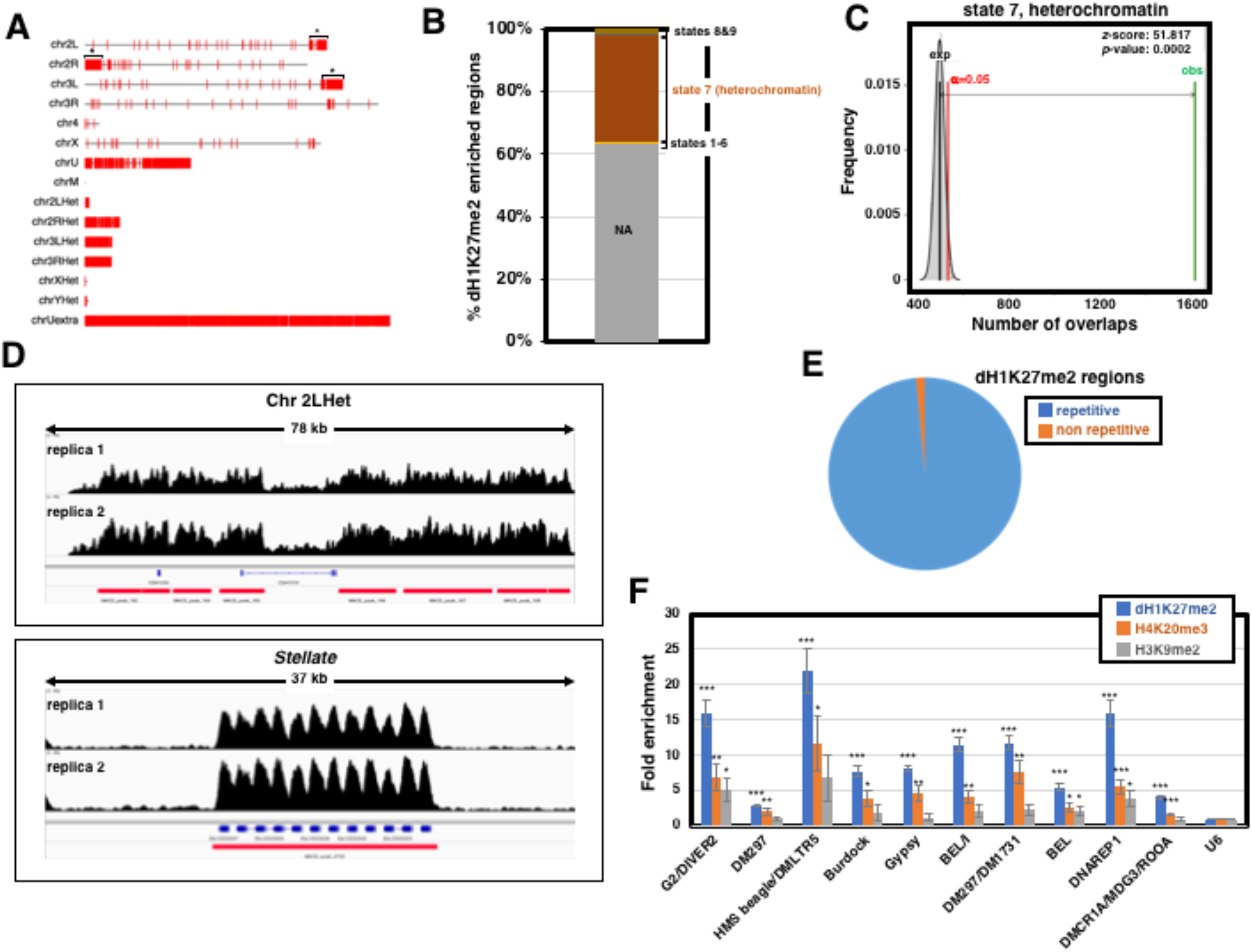
αdH1K27me2 ChIPseq analysis in S2 cells. **A**) Chromosomal distribution of the identified dH1K27me2 enriched regions. chr2L and chr2R, and chr3L and chr3R correspond to chromosome 2 and 3 left and right arms respect to the position of the centromere, respectively. Asterisks indicate dH1K27me2 enriched regions clustering at pericentromeric regions. chr4 and chrX are oriented with the centromere to the right. chr2LHet, chr2RHet, chr3LHet, chr3RHet, chrXHet and chrYHet correspond to partially assembled pericentromeric heterochromatin regions of the indicated chromosomes. chrU and chrUextra correspond to unassembled highly repetitive heterochromatic chromosome regions. chrM corresponds to the mitochondrial chromosome. **B**) The proportion of dH1K27me2 enriched regions assigned to each of the nine chromatin epigenetic states according to (3). NA: dH1K27me2 enriched regions that could not be assigned to any chromatin state. **C**) Permutation analysis showing statistical significance of the association of dH1K27me2 enriched regions with chromatin state 7 (heterochromatin) according to (3). The frequency of the number of overlaps is presented based on 5000 random permutations of the experimentally identified regions. The average expected number of overlaps (black) is compared with the observed number of overlaps (green). The α=0.05 confidence interval is indicated (red). z-score and permutation test p-value of the difference are also indicated. **D**) ChIPseq coverage profiles of dH1K27me2 across the indicated genomic regions of Chr2LHet (top) and the *Stellate locus* in ChrX (bottom) are presented for the two ChIPseq replicas. Genomic organization of the regions are indicated. **E**) Pie graph showing the proportion of identified dH1K27me2 enriched regions as a function of their content in repeated sequences. **F**) ChIPqPCR analysis with αdH1K27me2 (blue bars), αH4K20me3 (orange bars) and αH3K9me2 (gray bars) at the indicated genomic region is presented as the fold enrichment respect to *U6* as a control region. Results are the average of 2-4 independent experiments. Error bars are s.e.m. p-values respect to *U6* are indicated (*<0.1;**<0.05;***<0.001; two-tailed Student’s t-test).

A large majority of the dH1K27me2 enriched regions (4,683 regions, >98%) mapped to repetitive DNA sequences (**Figure 4E**). These included many transposable elements (TEs), both LTR and non-LTR retrotransposon and DNA transposons, but also long repeat sequences, satellite DNAs and a long list of simple repeated DNA sequences (**Supplementary Tables III** and **IV**). In good agreement with these results, ChIP-qPCR experiments detected significant dH1K27me2 enrichment in ten selected TEs with respect to a negative control region (**Figure 4F**). Enrichments ranged from 3-fold to 20-fold depending on the element analyzed. These regions were also enriched in H3K9me2 and H4K20me3, reflecting their heterochromatic character (**Figure 4F**). We also observed that an unusually high proportion of the reads (21-45%, depending on the replicate) failed to align to the reference genome. A cluster analysis of sequence similarity performed with the unaligned reads showed that they were highly enriched in uncatalogued simple repeat sequences (**Supplementary Table V**).

Altogether these results strongly suggest that dH1K27me2 largely accumulates at heterochromatic regions. Interestingly, DNA sequence motif analysis of the dH1K27me2 regions showed statistically significant enrichment in several motifs that, contrary to our expectations, were complex, did not represent any known satellite DNA sequences or simple repeat, and scored as binding sites for several transcription factors (TFs) (**Supplementary Table VI**).

### dH1K27me2 is independent of H3K9 methylation

Next, we analyzed the extent to which dH1K27me2 at heterochromatin is linked to known epigenetic hallmarks of heterochromatin. A main pathway of heterochromatin formation involves H3K9 methylation by Su(var)3-9 and subsequent binding of HP1a (5). Here, to analyze the relationship between dH1K27me2 and H3K9 methylation, we performed IF experiments in polytene chromosomes from flies carrying a H3K9R mutation that prevents H3K9 methylation. In these flies, deletion of the complete *HisC* locus, which contains ∼100 tandemly repeated copies of the 5kb histone repeat that carries one copy of each histone gene, was complemented by a transgenic construct containing 12 copies of the histone monomer either wild type (12xWT) or carrying a H3K9R mutation (12xH3K9R) (16). We observed that in H3K9R mutant flies the levels of H3K9me2 at the chromocenter were markedly reduced in polytene chromosomes (**Figure 5A**). In agreement, HP1a localization at the chromocenter was heavily reduced (**Figures 5A** and **5B**) (16). Remarkably, the pattern of dH1K27me2 remained largely unaffected since no signs of reduction at the chromocenter or redistribution could be appreciated (**Figure 5B**). Along the same lines, depletion of Su(var)3-9 did not affect the pattern of dH1K27me2. In these experiments, salivary glands from knockdown *Su(var)3*-*9^RNAi^* flies were squashed and immunostained together with salivary glands from control flies expressing an H2AvD2::GFP fusion protein. In this way, variability due to immunostaining conditions was minimized, while control and knockdown polytene chromosomes are easily distinguished on the basis of their GFP fluorescence. We observed that Su(var)3-9 depletion, which strongly reduced H3K9me2 (**Figure 6A**), did not affect the overall pattern of dH1K27me2, nor its accumulation at the chromocenter (**Figure 6B**). Similar experiments showed that HP1a depletion had no effect either on the pattern of dH1K27me2 (**Figure 7**). Another important hallmark of heterochromatin is H4K20 methylation by Su(var)4-20, which is recruited to heterochromatin by HP1a and H3K9 methylation (6,7). We observed that the pattern of dH1K27me2 distribution in polytene chromosomes was not affected by depletion of Su(var)4-20 (Figure 8B), while H4K20me3 was strongly reduced (Figure 8A).

**Figure 5.**
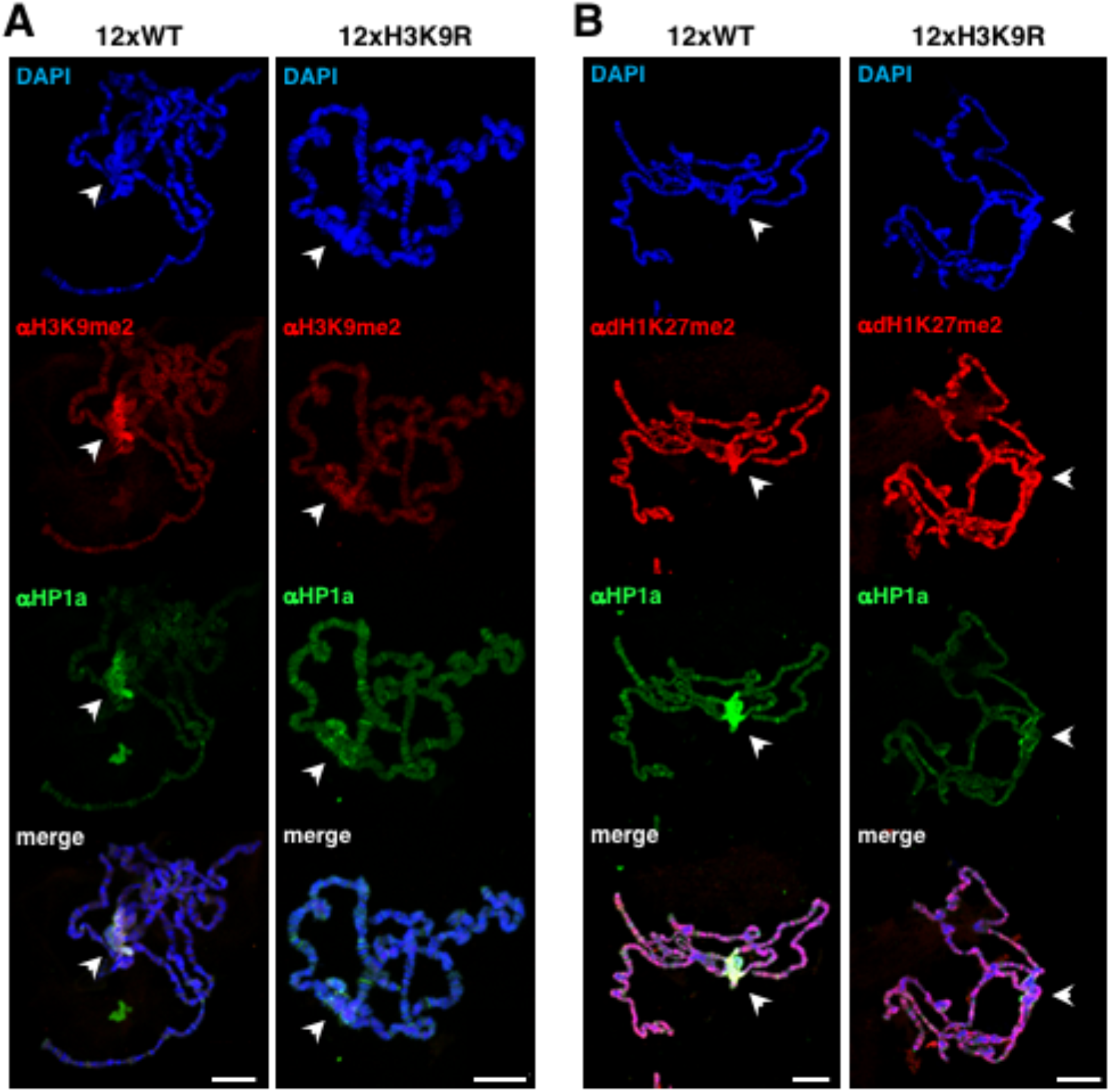
dH1K27me2 is independent of histone H3K9 methylation. **A**) Immunostaining with αH3K9me2 (in red) and αHP1a (in green) of polytene chromosomes from flies carrying 12 copies of a WT (left) or a mutated H3K9R histone repeat (right) in front of a deficiency of the *HisC* locus. DNA was stained with DAPI (in blue). Arrowheads indicate the chromocenter. Scale bars correspond to 20μm. **B**) As in A but stained with αdH1K27me2 (in red) and αHP1a (in green).

**Figure 6.**
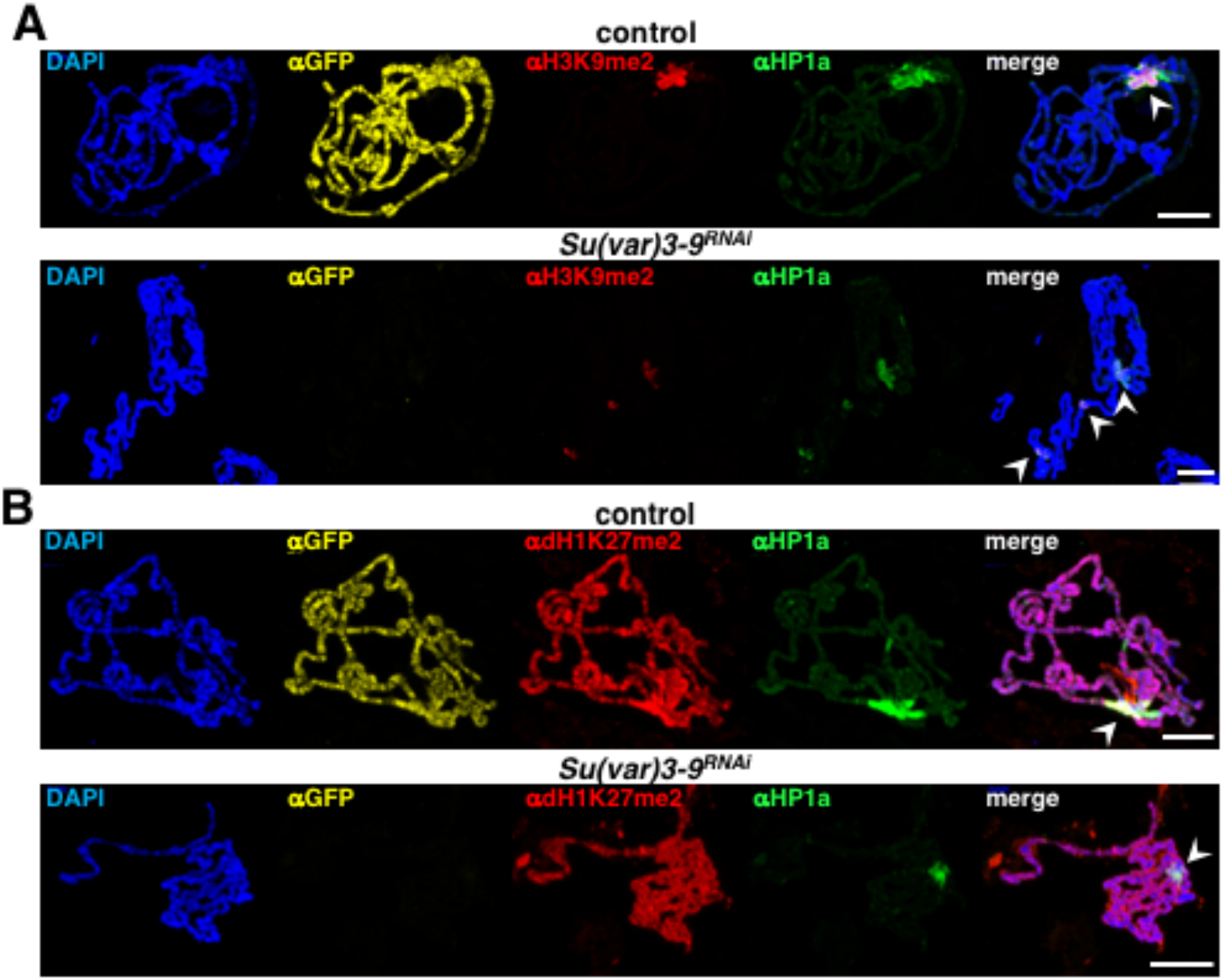
dH1K27me2 is not affected by Su(var)3-9 depletion. **A**) Polytene chromosomes from control flies expressing a H2AvD2::GFP fusion protein and from Su(var)3-9 depleted flies (*Su(var)3*-*9^RNAi^*) were mixed and co-immunostained with αGFP (in yellow) (to distinguish control and *Su(var)3*-*9^RNAi^* chromosomes), αH4K20me3 (in red) and αHP1a (in green). DNA was stained with DAPI (in blue). Arrowheads indicate the chromocenter. Scale bars correspond to 20μm. **B**) As in A but chromosomes were immunostained with αGFP (in yellow), αdH1K27me2 (in red) and αHP1a (in green).

**Figure 7.**
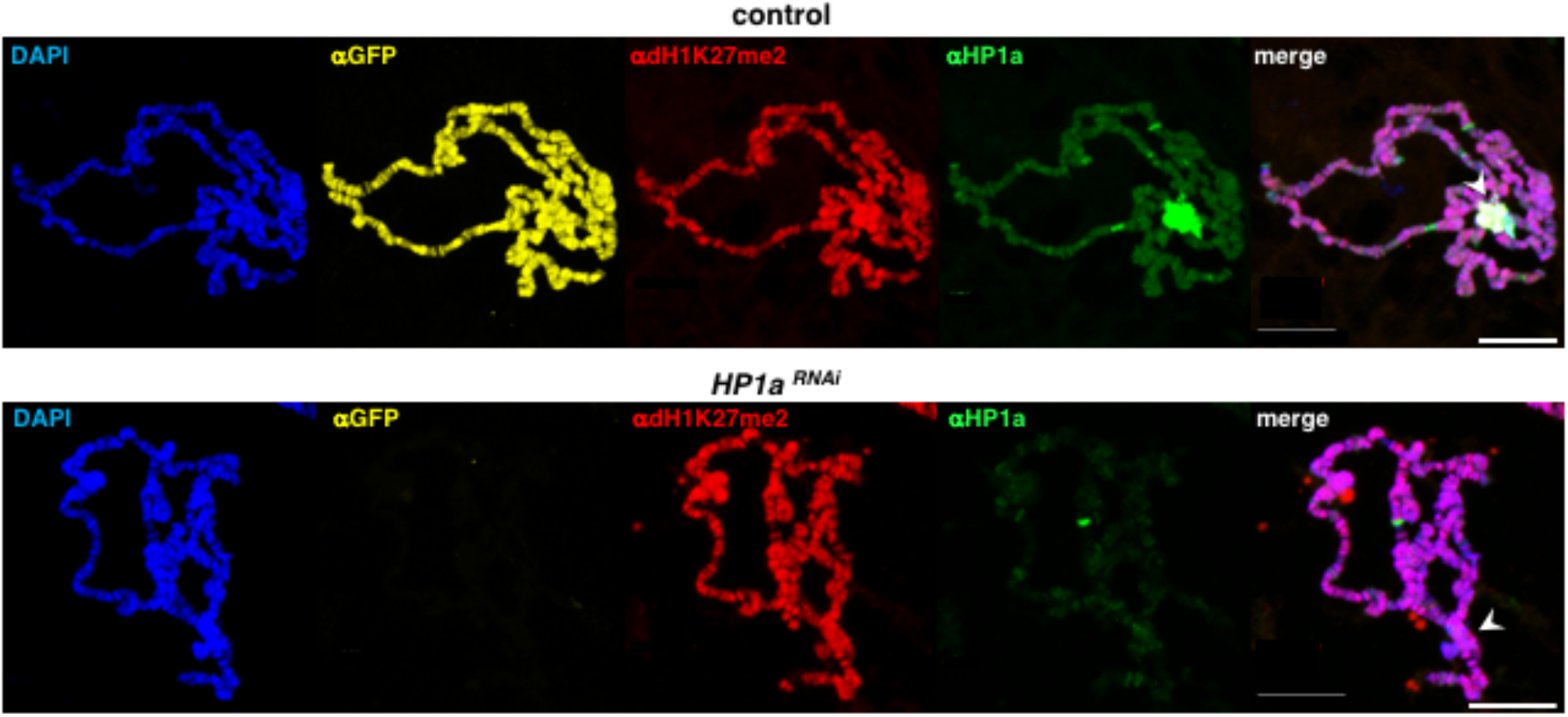
dH1K27me2 is not affected by HP1a depletion. Polytene chromosomes from control flies expressing a H2AvD2::GFP fusion protein and from HP1a depleted flies (*HP1a^RNAi^*) were mixed and co-immunostained with αGFP (in yellow) (to distinguish control and *HP1a^RNAi^* chromosomes), αdH1K27me2 (in red) and αHP1a (in green). DNA was stained with DAPI (in blue). Arrowheads indicate the chromocenter. Scale bars correspond to 20μm

**Figure 8.**
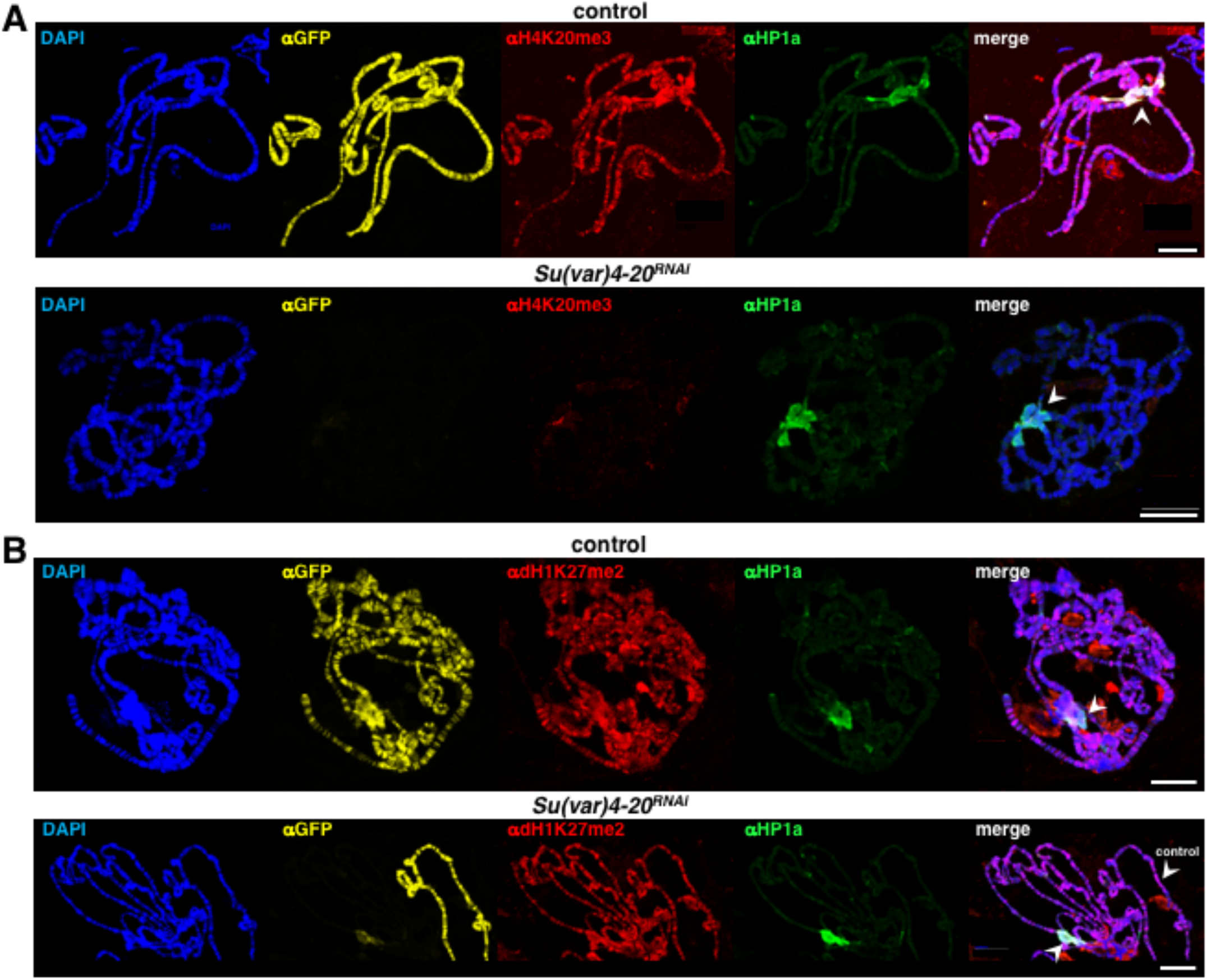
dH1K27me2 is not affected by Su(var)4-20 depletion. **A**) Polytene chromosomes from control flies expressing a H2AvD2::GFP fusion protein and from Su(var)4-20 depleted flies (*Su(var)4*-*20^RNAi^*) were mixed and co-immunostained with αGFP (in yellow) (to distinguish control and *Su(var)4*-*20^RNAi^* chromosomes), αH4K20me3 (in red) and αHP1a (in green). DNA was stained with DAPI (in blue). Arrowheads indicate the chromocenter. Scale bars correspond to 20μm. **B**) As in A but chromosomes were immunostained with αGFP (in yellow), αdH1K27me2 (in red) and αHP1a (in green). Note that in the lower *Su(var)4*-*20^RNAi^* panels an arm from a control chromosome is also visible in the same field allowing for a better comparison.

Altogether, these results suggest that dH1K27me2 at heterochromatin does not depend on the Su(var)3-9/H3K9 methylation/HP1a pathway of heterochromatin formation, nor on Su(var)4-20 or H4K20 methylation.

## DISCUSSION

The contribution of linker histones H1 to the epigenetic regulation of chromatin functions is currently under active study since increasing evidence is pointing out to far more than its known structural role (1). In this regard, like core histones, linker histones H1 are subjected to multiple PTMs, which are anticipated to have important contributions to the regulation of their functions. However, to date, data on their genomic distribution and functional contribution are scarce. Here we have provided strong evidence for dH1K27me2 as a new epigenetic modification that marks heterochromatin in *Drosophila*. dH1K27me2 is detected as early as histone H1 is expressed at cellularization and heterochromatinization begins (27), remaining constant thereafter through development. Our results suggest that dH1K27me2 associates with heterochromatin throughout the cell cycle. In mitosis, dH1K27me2 strongly accumulates at pericentromeric heterochromatin. However, in interphase, dH1K27me2 shows a broader genomic distribution. In the interphase polytene chromosomes, dH1K27me2 is detected in pericentromeric heterochromatin at the chromocenter, but, also, in intercalary heterochromatin at DAPI-intense bands. Similarly, in interphase cells, dH1K27me2 is not constrained to pericentromeric heterochromatin since significant αdH1K27me2 reactivity is detected at regions that do not show detectable αHP1a immunostaining. ChIPseq analysis suggests that these regions correspond to intercalary heterochromatin since the vast majority (>98%) of dH1K27me2 enriched sites detected in asynchronous S2 cell cultures, in which the proportion of mitotic cells is <5%, maps to repeated DNA elements, both TE and simple repeat sequences. These observations suggest that dH1K27me2 genomic distribution during cell cycle progression is dynamic, being released from intercalary heterochromatin in mitosis to accumulate at pericentromeric heterochromatin.

Interestingly, dH1K27me2 was not found enriched in chromosome 4, despite it is highly heterochromatic and enriched in both H3K9me2 and HP1a. Of note, the mechanisms behind heterochromatin formation in chromosome 4 are different since H3K9 methylation and HP1a binding depend on SETDB1 instead of Su(var)3-9 as in pericentromeric heterochromatin (28–30). Along the same lines, dH1K27me2 was not detected at telomeres, which are also heterochromatic and enriched in HP1a. In this regard, our ChIPseq analysis detected dH1K27me2 enrichment at the telomeric HET-A and TART retrotransposons. However, this enrichment likely reflects the presence of dH1K27me2 at pericentromeric heterochromatin of chromosomes Y and III, where HET-A and TART are also present (26,31,32). Finally, HP1a is detected at several euchromatic regions where, paradoxically, it is required for gene expression (33). At these regions, dH1K27me2 was neither detected.

Altogether these results indicate that dH1K27me2 is largely restricted to pericentromeric and intercalary heterochromatic regions, but it is absent from other types of heterochromatic regions. Consistent with its preferential localization in heterochromatin, the presence of dH1K27me2 strongly correlates with other heterochromatic marks, namely H3K9me2 and, specially, H4K20me3. Interestingly, while H3K9me2 is mostly constrained to the chromocenter and chromosome 4, the pattern of H4K20me3 distribution in polytene chromosomes strongly resembles that of dH1K27me2, being detected at the chromocenter and in the DAPI-intense bands of intercalary heterochromatin, but largely absent from chromosome 4 (7). In addition, H4K20me3 distribution in human metaphase chromosomes is strikingly similar to the dH1K27me2 distribution observed here in fly cells (34). Finally, H4K20me3 localization in C127 cells has also been suggested to be cell cycle regulated (7).

Our results show that dH1K27me2 accumulation at pericentromeric heterochromatin is independent of H3K9 methylation by Su(var)3-9, suggesting that dH1K27 methylation is an early step in pericentromeric heterochromatin formation in *Drosophila*. In this regard, it has been reported that dH1 has a crucial contribution to heterochromatin organization and stability (20,24,35). In particular, it has been proposed that dH1 directly interacts with Su(var)3-9 and is responsible for its tethering to pericentromeric heterochromatin, enhancing its H3K9 methylation activity *in vitro* (35). How dH1, which is distributed all throughout the genome, recruits Su(var)3-9 only at pericentromeric heterochromatin is not well understood. It is tempting to speculate that dH1K27me2 regulates Su(var)3-9 recruitment to pericentromeric heterochromatin and initiates heterochromatin formation. Interestingly, while at the chromocenter Su(var)4-20 recruitment depends on the Su(var)3-9/HP1a pathway (6,7), the mechanisms involved in its recruitment at the intercalary heterochromatin of the DAPI-intense bands in the chromosome arms are unknown. In this regard, our results show that dH1K27me2 in intercalary heterochromatin acts upstream of H4K20 methylation by Su(var)4-20, suggesting that dH1K27me2 might also be involved in Su(var)4-20 recruitment in intercalary heterochromatin. Unfortunately, enzymes involved in dH1K27me2 remain unknown and we failed to identify any dH1K27me2 HTM and KDM in KD screens of possible candidates. Thus, further work is required to determine the mechanisms regulating dH1K27me2 and its actual contribution to heterochromatin formation.

We found that dH1K27me2 regions are statistically enriched in several TFs binding motifs, some of which have been reported to associate with pericentromeric heterochromatin, such as GAGA factor and the PAX family TF eyegone (eyg) (36,37). While the presence of TEs and repeated sequences was expected, the presence of TF binding motifs was somehow surprising. Indeed, TF binding motifs in *D. melanogaster* TEs have recently been analyzed (38) and none of them correspond to the ones found in our work. A similar enrichment of certain TF binding sequences in heterochromatin has already been reported in mice and a role in heterochromatin formation of certain transcription factors (Pax3 and Pax9) has already been suggested (39). In flies, a dynamic binding of some TFs to pericentromeric regions has also been reported (36,40). Notably, some of them (GAGA factor and Prod) appear to accumulate at the pericentromeric regions only in metaphase chromosomes and to redistribute in interphase chromatin. Remarkably, both are essentially absent from the chromocenter in polytene chromosomes (36). Since dH1K27me2 shows a cell cycle-dependent accumulation at pericentromeric regions in mitosis, it is possible that this PTM might be involved in such dynamic behavior.

In summary, in comparison to core histones, linker histones H1 PTMs have been poorly studied as potential epigenetic marks. Here, we have characterized dimethylation of lysine 27 on somatic *Drosophila* histone dH1 as a novel epigenetic modification that labels heterochromatin independently of H3K9 methylation and, thus, may be acting at the initial steps of heterochromatin formation. Interestingly, dH1K27 occurs within a short conserved sequence motif and it has been reported to be methylated in some mammalian H1 variants (12,13), opening the possibility that its heterochromatin localization is not constrained to *Drosophila*. These results suggest the possibility that other dH1 PTMs might also have restricted distributions as well and highlight the potential of linker histones H1 as epigenetic regulators of chromatin structure and function.

## ACKNOWLEDGEMENTS

We are thankful to Dr J. Kadonaga for generous gift of αdH1antibodies and to Dr. R. Duronio for 12xWT and 12XH3K9R fly stocks, and to all members of the IBMB Molecular Genomics department for constructive comments and support. We acknowledge the Advanced Digital Microscopy facility of IRB for help with confocal microscopy. OV acknowledges receipt of an IRB fellowship.

## SUPPLEMENTARY TABLES

**Supplementary Table I.**
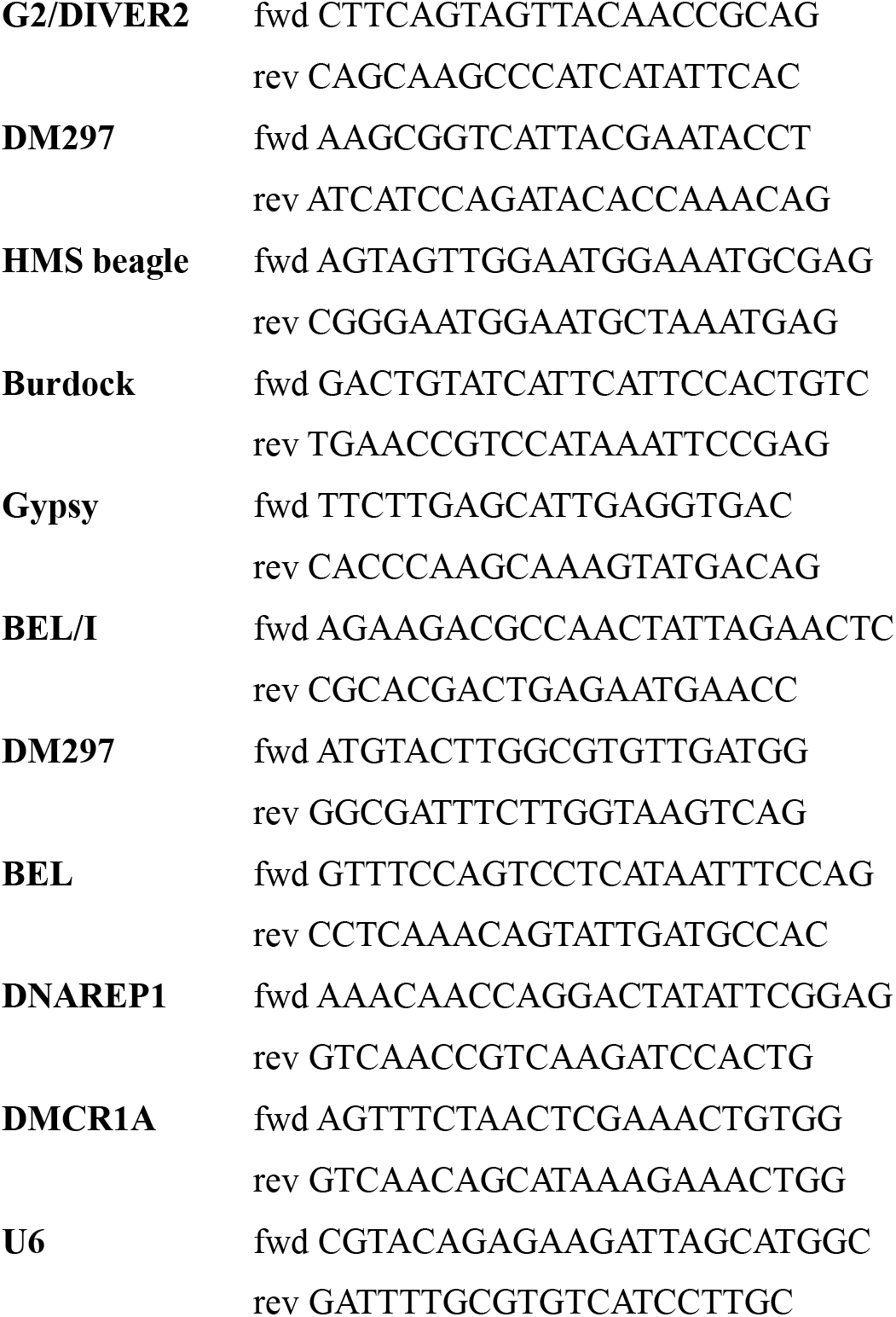
Primers used in ChIPqPCR experiments.

**Supplementary Table III.**
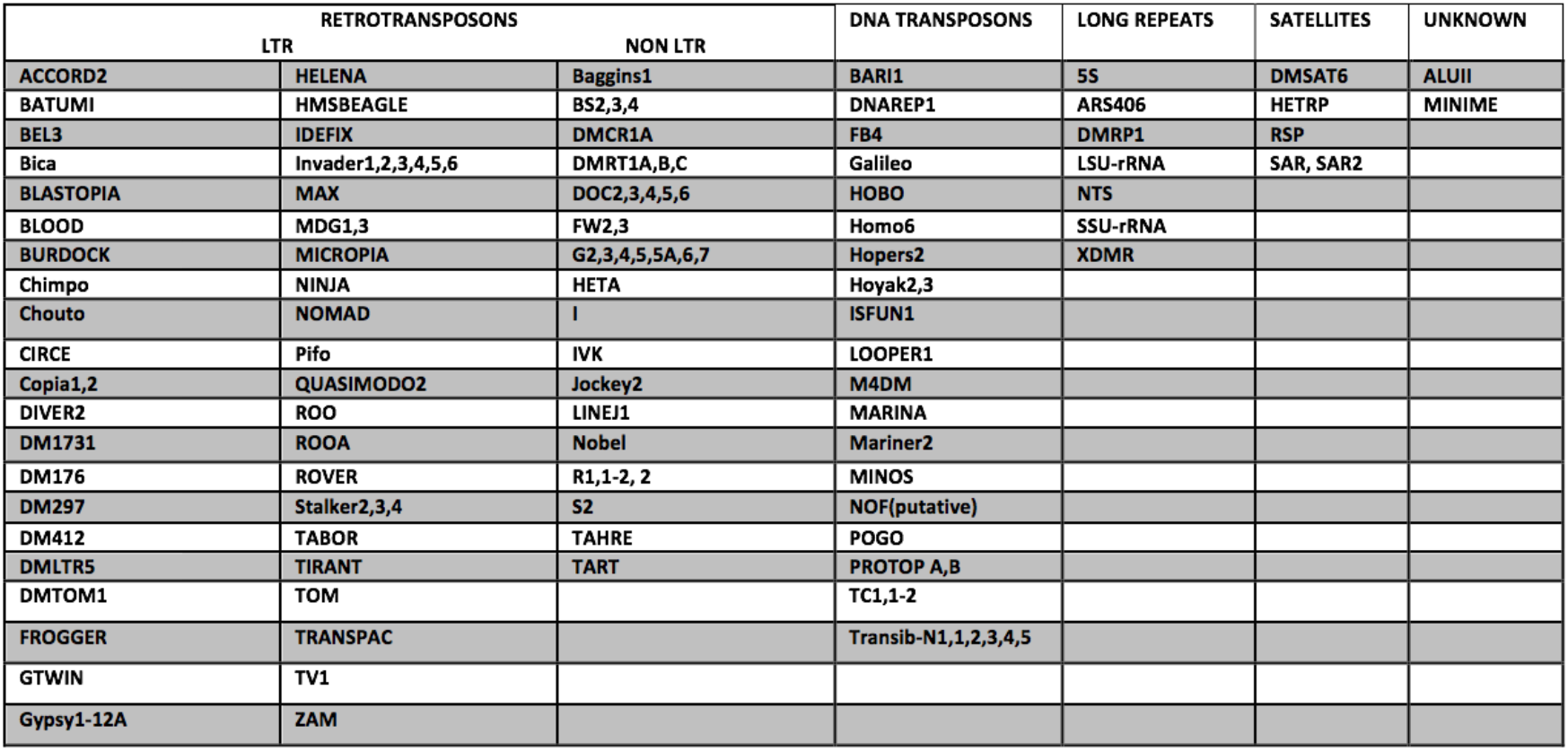
List of transposable elements (TE), long repeats and satellite DNAs identified at dH1K27me2 enriched regions.

**Supplementary Table IV.**
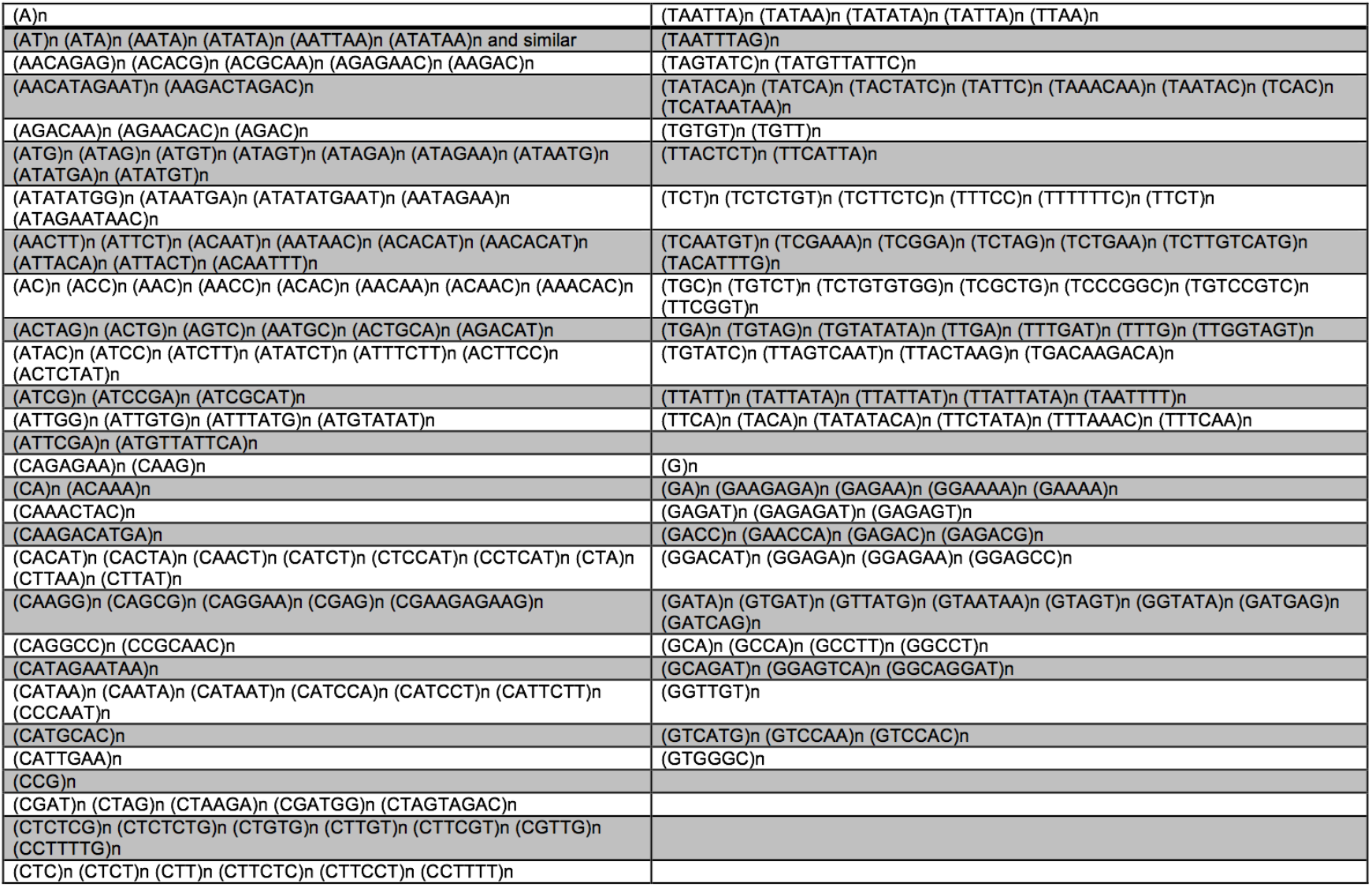
List of simple repeated DNA sequences identified at dH1K27me2 enriched regions.

**Supplementary Table V.**
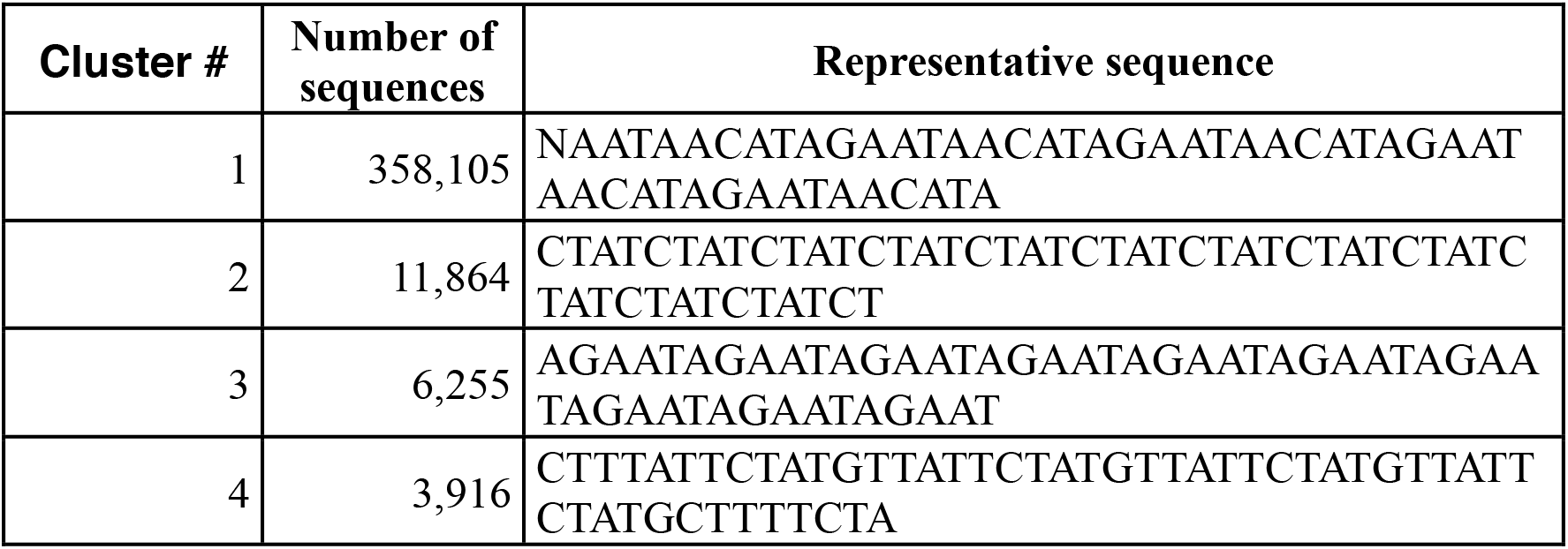
Clusters of simple repeated DNA sequences identified within the unaligned reads.

**Supplementary Table VI.**
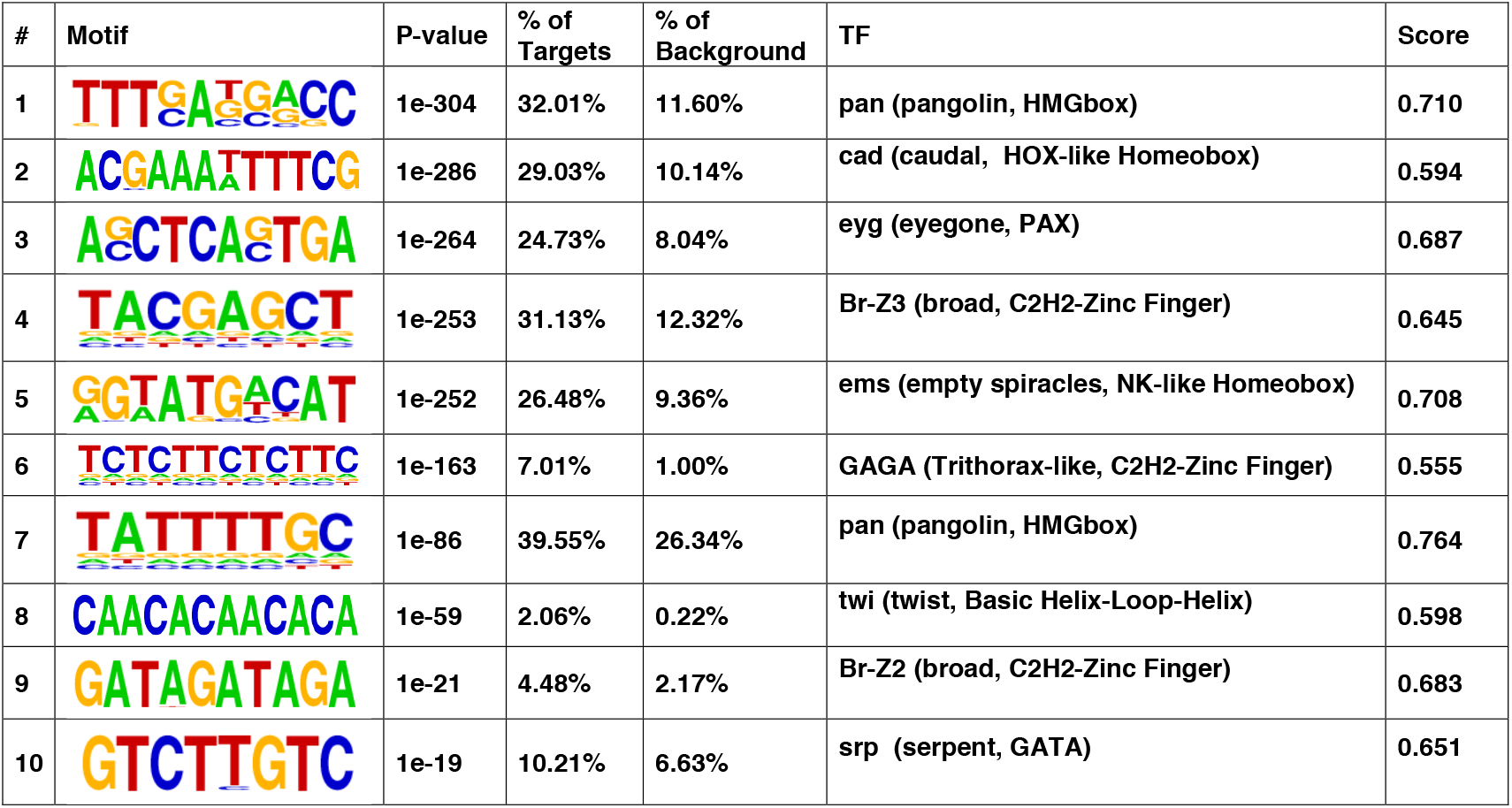
List of the top ten DNA sequence motifs identified at dH1K27me2 enriched regions.

| # | Motif | P-value | % of Targets | % of Background | TF | Score |
| --- | --- | --- | --- | --- | --- | --- |
| 1 |  | 1e-304 | 32.01% | 11.60% | pan (pangolin, HMGbox) | 0.710 |
| 2 |  | 1e-286 | 29.03% | 10.14% | cad (caudal, HOX-like Homeobox) | 0.594 |
| 3 |  | 1e-264 | 24.73% | 8.04% | eyg (eyegone, PAX) | 0.687 |
| 4 |  | 1e-253 | 31.13% | 12.32% | Br-Z3 (broad, C2H2-Zinc Finger) | 0.645 |
| 5 |  | 1e-252 | 26.48% | 9.36% | ems (empty spiracles, NK-like Homeobox) | 0.708 |
| 6 |  | 1e-163 | 7.01% | 1.00% | GAGA (Trithorax-like, C2H2-Zinc Finger) | 0.555 |
| 7 |  | 1e-86 | 39.55% | 26.34% | pan (pangolin, HMGbox) | 0.764 |
| 8 |  | 1e-59 | 2.06% | 0.22% | twi (twist, Basic Helix-Loop-Helix) | 0.598 |
| 9 |  | 1e-21 | 4.48% | 2.17% | Br-Z2 (broad, C2H2-Zinc Finger) | 0.683 |
| 10 |  | 1e-19 | 10.21% | 6.63% | srp (serpent, GATA) | 0.651 |

## LEGENDS TO SUPPLEMENTARY FIGURES

**Supplementary Figure S1.**
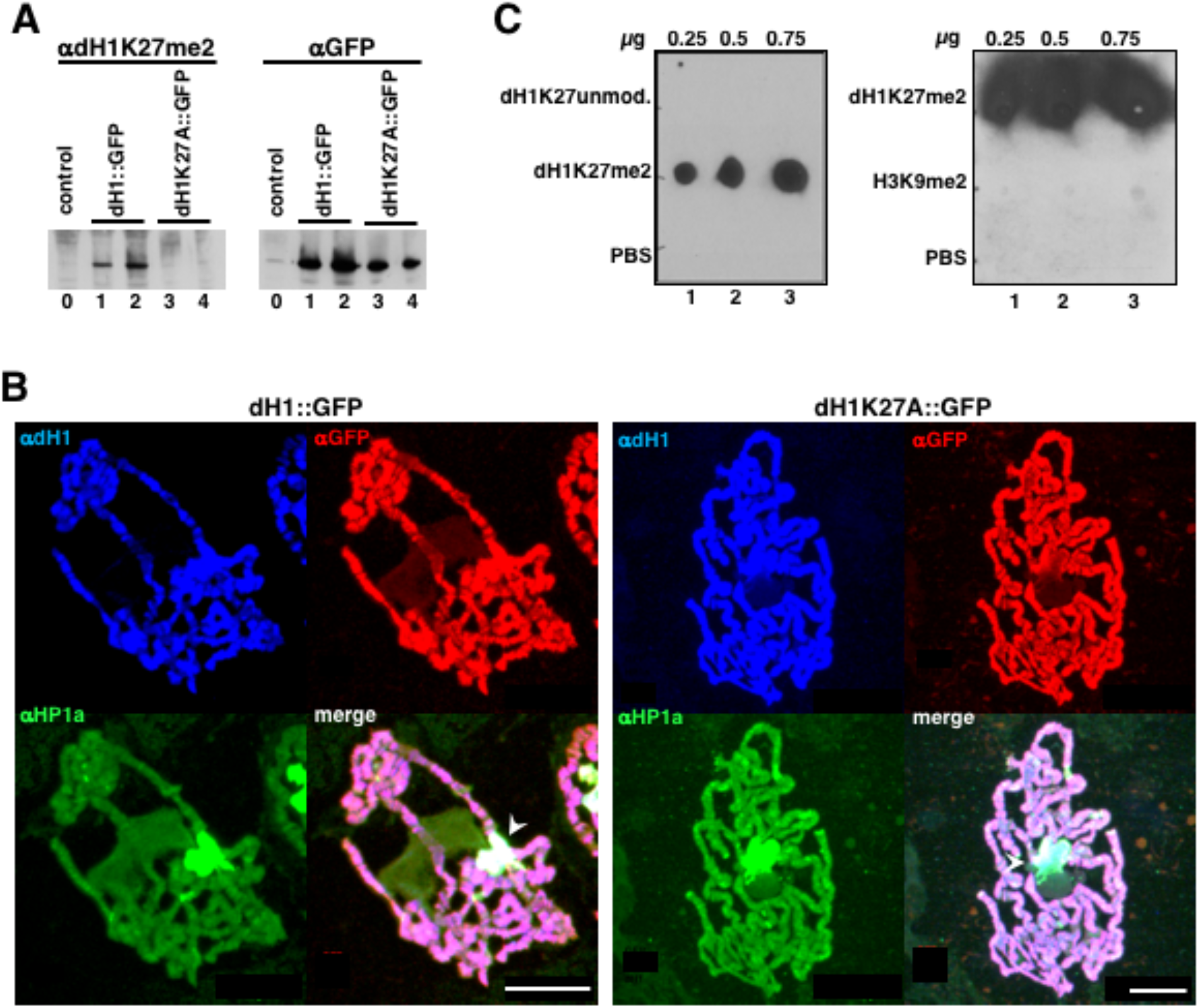
Specificity of αdH1K27me2 antibodies. **A**) WB analysis with αdH1K27me2 (left) and αGFP (right) of total extracts from 3^rd^ instar salivary glands from flies overexpressing dH1::GFP (lanes 1 and 2) and dH1K27A::GFP (lanes 3 and 4). Extracts from two independent experiments are shown. Lanes 0 correspond to extracts from control flies expressing no GFP-tagged construct. **B**) Immunostainings with αGFP (in red), αHP1a (in green) and αdH1 (in blue) of polytene chromosomes from flies expressing dH1::GFP (left) and dH1K27A::GFP (right). Arrowheads indicate the chromocenter. Scale bars correspond to 20μm. **C**) On the left, dot-blot assay using increasing amounts of a dH1-peptide carrying dimethylated K27 (dH1K27me2), unmethylated K27 (dH1K27unmod) or no peptide (PBS). Membranes were assayed for binding with αdH1K27me2 antibodies. On the right, a similar assay but with increasing amounts of a H3-peptide carrying trimethylated K9 (H3K9me3).

**Supplementary Figure S2.**
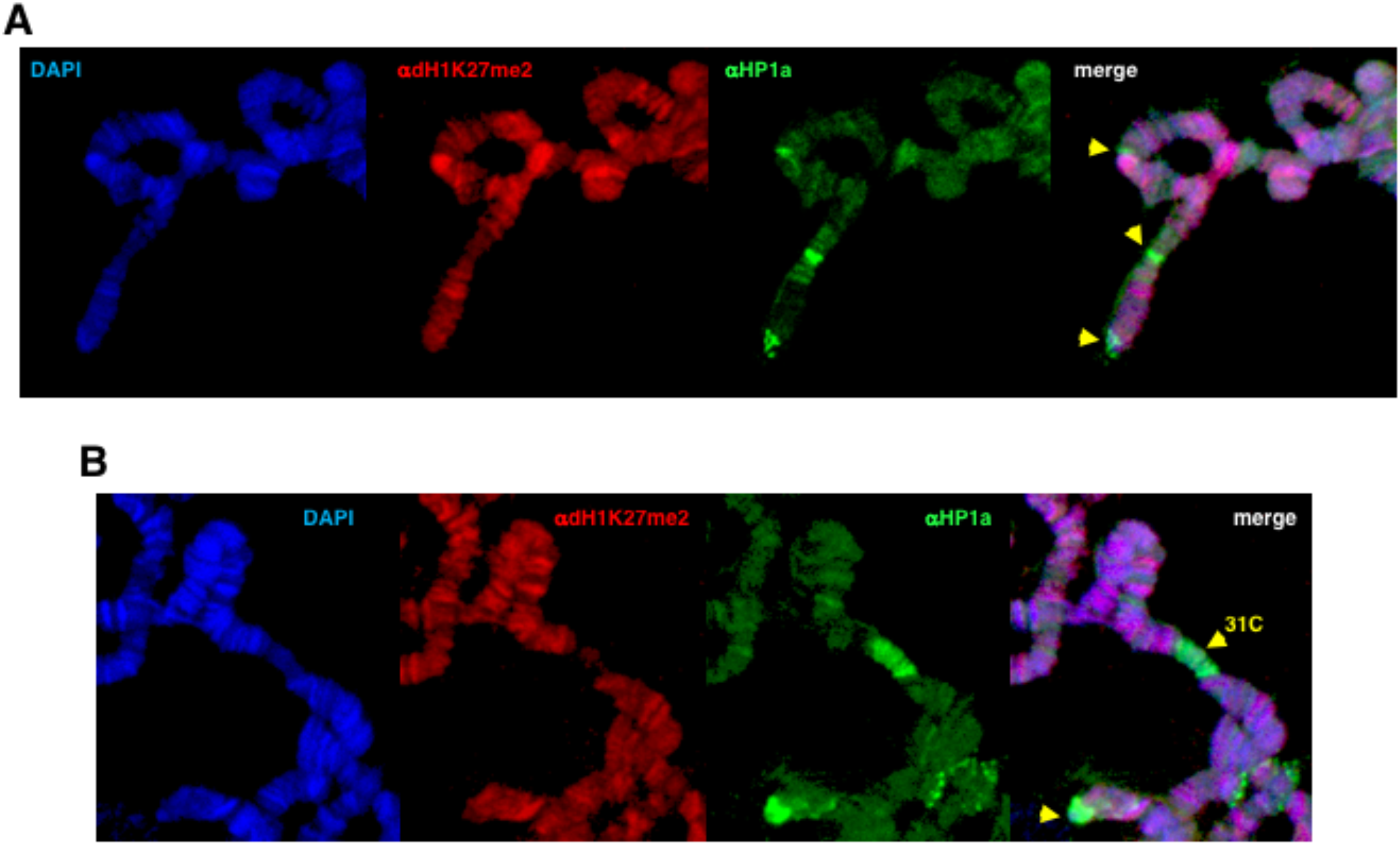
**A**) Enlarged image of the tip of a polytene chromosome immunostained with αdH1K27me2 (in red) and αHP1a (in green). DNA was stained with DAPI (in blue). **B**) As in A but for the HP1a positive euchromatic region 31C. In both panels, arrowheads indicate αHP1a bands not overlapping with αdH1K27me2 signals.

