## Supplementary material for "Linker histone dH1K27 dimethylation marks *Drosophila* heterochromatin independently of H3K9 methylation": Suplemental table II

### Supplementary Table II. dH1K27me2 enriched genomic regions

| Chr | start | end | width | fc | annot. Region |
|---|---|---|---|---|---|
| chr2L | 770143 | 772652 | 2510 | 2,86 | 0001 |
| chr2L | 2112422 | 2119126 | 6705 | 2,11 | 0002 |
| chr2L | 4662413 | 4668736 | 6324 | 2,46 | 0003 |
| chr2L | 5164681 | 5167897 | 3217 | 2,23 | 0004 |
| chr2L | 5169827 | 5171144 | 1318 | 2,14 | 0005 |
| chr2L | 5909460 | 5910855 | 1396 | 2,04 | 0006 |
| chr2L | 6991368 | 6997278 | 5911 | 2,89 | 0007 |
| chr2L | 8064034 | 8065557 | 1524 | 2,17 | 0008 |
| chr2L | 9899913 | 9902663 | 2751 | 3,33 | 0009 |
| chr2L | 9974993 | 9980542 | 5550 | 2,5 | 0010 |
| chr2L | 10593363 | 10599772 | 6410 | 2,2 | 0011 |
| chr2L | 10862963 | 10864564 | 1602 | 2,39 | 0012 |
| chr2L | 11541440 | 11547746 | 6307 | 2,68 | 0013 |
| chr2L | 12558726 | 12564198 | 5473 | 2,35 | 0014 |
| chr2L | 13030499 | 13031753 | 1255 | 2,2 | 0015 |
| chr2L | 13522122 | 13527669 | 5548 | 2,15 | 0016 |
| chr2L | 14985192 | 14990561 | 5370 | 2,48 | 0017 |
| chr2L | 15051509 | 15053117 | 1609 | 2,48 | 0018 |
| chr2L | 15723283 | 15724345 | 1063 | 2,3 | 0019 |
| chr2L | 15828332 | 15829458 | 1127 | 2,63 | 0020 |
| chr2L | 16243300 | 16245263 | 1964 | 2,16 | 0021 |
| chr2L | 16513269 | 16518754 | 5486 | 3,33 | 0022 |
| chr2L | 17144648 | 17151373 | 6726 | 2,41 | 0023 |
| chr2L | 17689231 | 17690855 | 1625 | 1,84 | 0024 |
| chr2L | 19657955 | 19664620 | 6666 | 2,5 | 0025 |
| chr2L | 20007207 | 20008938 | 1732 | 1,99 | 0026 |
| chr2L | 21358254 | 21360403 | 2150 | 2,52 | 0027 |
| chr2L | 21363040 | 21367392 | 4353 | 2,96 | 0028 |
| chr2L | 21367722 | 21370087 | 2366 | 2,01 | 0029 |
| chr2L | 21373866 | 21376380 | 2515 | 2,14 | 0030 |
| chr2L | 21385985 | 21387526 | 1542 | 1,96 | 0031 |
| chr2L | 21398600 | 21399660 | 1061 | 2,1 | 0032 |
| chr2L | 21405279 | 21406430 | 1152 | 4,71 | 0033 |
| chr2L | 21471058 | 21475994 | 4937 | 2,45 | 0034 |
| chr2L | 21476006 | 21481035 | 5030 | 2,58 | 0035 |
| chr2L | 21540570 | 21544100 | 3531 | 2,75 | 0036 |
| chr2L | 21544372 | 21549795 | 5424 | 2,47 | 0037 |
| chr2L | 21555446 | 21559861 | 4416 | 2,22 | 0038 |
| chr2L | 21583419 | 21587146 | 3728 | 1,84 | 0039 |
| chr2L | 21587376 | 21590433 | 3058 | 1,93 | 0040 |
| chr2L | 21590559 | 21596299 | 5741 | 2,27 | 0041 |
| chr2L | 21605939 | 21613256 | 7318 | 1,87 | 0042 |
| chr2L | 21665575 | 21671763 | 6189 | 1,98 | 0043 |
| chr2L | 21900695 | 21904049 | 3355 | 2,28 | 0044 |
| chr2L | 22087201 | 22104620 | 17420 | 3,19 | 0045 |

| | | | | | |
|---|---|---|---|---|---|
| chr2L | 22104710 | 22112879 | 8170 | 2,57 | 0046 |
| chr2L | 22115662 | 22117823 | 2162 | 2,06 | 0047 |
| chr2L | 22151217 | 22152950 | 1734 | 2,63 | 0048 |
| chr2L | 22152978 | 22159145 | 6168 | 2,3 | 0049 |
| chr2L | 22159170 | 22164882 | 5713 | 2,8 | 0050 |
| chr2L | 22187553 | 22191371 | 3819 | 2,47 | 0051 |
| chr2L | 22191531 | 22193191 | 1661 | 2,56 | 0052 |
| chr2L | 22193414 | 22195803 | 2390 | 3,18 | 0053 |
| chr2L | 22212561 | 22217038 | 4478 | 2,69 | 0054 |
| chr2L | 22217495 | 22218711 | 1217 | 2,36 | 0055 |
| chr2L | 22220570 | 22224717 | 4148 | 2,72 | 0056 |
| chr2L | 22224795 | 22226526 | 1732 | 2,88 | 0057 |
| chr2L | 22226774 | 22228836 | 2063 | 2,91 | 0058 |
| chr2L | 22231815 | 22232913 | 1099 | 2,04 | 0059 |
| chr2L | 22236881 | 22238034 | 1154 | 2,75 | 0060 |
| chr2L | 22261021 | 22263040 | 2020 | 2,36 | 0061 |
| chr2L | 22264063 | 22265631 | 1569 | 2,3 | 0062 |
| chr2L | 22271065 | 22273473 | 2409 | 2,78 | 0063 |
| chr2L | 22275978 | 22284529 | 8552 | 3,27 | 0064 |
| chr2L | 22289749 | 22292762 | 3014 | 2,79 | 0065 |
| chr2L | 22293036 | 22296120 | 3085 | 2,67 | 0066 |
| chr2L | 22296275 | 22297894 | 1620 | 2,62 | 0067 |
| chr2L | 22298377 | 22300548 | 2172 | 2,43 | 0068 |
| chr2L | 22308187 | 22310275 | 2089 | 2,73 | 0069 |
| chr2L | 22314913 | 22318630 | 3718 | 2,45 | 0070 |
| chr2L | 22318827 | 22322976 | 4150 | 3,07 | 0071 |
| chr2L | 22323048 | 22329744 | 6697 | 3,37 | 0072 |
| chr2L | 22333917 | 22336354 | 2438 | 3,43 | 0073 |
| chr2L | 22339296 | 22349240 | 9945 | 3,59 | 0074 |
| chr2L | 22349466 | 22356594 | 7129 | 3,01 | 0075 |
| chr2L | 22356669 | 22362189 | 5521 | 2,41 | 0076 |
| chr2L | 22369649 | 22373185 | 3537 | 2,75 | 0077 |
| chr2L | 22373228 | 22378079 | 4852 | 3,72 | 0078 |
| chr2L | 22381435 | 22382824 | 1390 | 2,14 | 0079 |
| chr2L | 22384632 | 22386238 | 1607 | 2,49 | 0080 |
| chr2L | 22389037 | 22396899 | 7863 | 2,98 | 0081 |
| chr2L | 22397641 | 22401458 | 3818 | 2,25 | 0082 |
| chr2L | 22402056 | 22406018 | 3963 | 2,82 | 0083 |
| chr2L | 22406157 | 22416103 | 9947 | 2,82 | 0084 |
| chr2L | 22420042 | 22426582 | 6541 | 2,91 | 0085 |
| chr2L | 22434008 | 22441294 | 7287 | 2,12 | 0086 |
| chr2L | 22444257 | 22448277 | 4021 | 3,49 | 0087 |
| chr2L | 22449477 | 22457521 | 8045 | 3,49 | 0088 |
| chr2L | 22463362 | 22465707 | 2346 | 2,18 | 0089 |
| chr2L | 22476583 | 22479986 | 3404 | 2,24 | 0090 |
| chr2L | 22480045 | 22486089 | 6045 | 2,74 | 0091 |
| chr2L | 22491050 | 22498414 | 7365 | 3,55 | 0092 |
| chr2L | 22509294 | 22515675 | 6382 | 2,9 | 0093 |

| | | | | | |
|---|---|---|---|---|---|
| chr2L | 22516947 | 22522985 | 6039 | 2,57 | 0094 |
| chr2L | 22523127 | 22531331 | 8205 | 3,24 | 0095 |
| chr2L | 22532472 | 22540620 | 8149 | 2,98 | 0096 |
| chr2L | 22541761 | 22545977 | 4217 | 2,78 | 0097 |
| chr2L | 22548134 | 22557790 | 9657 | 2,98 | 0098 |
| chr2L | 22558276 | 22562972 | 4697 | 2,97 | 0099 |
| chr2L | 22563015 | 22564275 | 1261 | 3,43 | 0100 |
| chr2L | 22564424 | 22572773 | 8350 | 3,32 | 0101 |
| chr2L | 22632220 | 22633866 | 1647 | 2,26 | 0102 |
| chr2L | 22634105 | 22644851 | 10747 | 2,77 | 0103 |
| chr2L | 22646367 | 22647520 | 1154 | 2,56 | 0104 |
| chr2L | 22650660 | 22658143 | 7484 | 3,07 | 0105 |
| chr2L | 22659761 | 22665910 | 6150 | 2,64 | 0106 |
| chr2L | 22666742 | 22669320 | 2579 | 3,37 | 0107 |
| chr2L | 22670889 | 22673605 | 2717 | 2,78 | 0108 |
| chr2L | 22673777 | 22676715 | 2939 | 2,59 | 0109 |
| chr2L | 22680601 | 22688478 | 7878 | 2,98 | 0110 |
| chr2L | 22688654 | 22689813 | 1160 | 2,32 | 0111 |
| chr2L | 22691802 | 22694167 | 2366 | 2,55 | 0112 |
| chr2L | 22696078 | 22698564 | 2487 | 2,59 | 0113 |
| chr2L | 22699562 | 22704966 | 5405 | 3,2 | 0114 |
| chr2L | 22705016 | 22718389 | 13374 | 3,37 | 0115 |
| chr2L | 22719685 | 22720881 | 1197 | 1,88 | 0116 |
| chr2L | 22721950 | 22723124 | 1175 | 2,36 | 0117 |
| chr2L | 22727703 | 22734231 | 6529 | 4,04 | 0118 |
| chr2L | 22748414 | 22754342 | 5929 | 3,3 | 0119 |
| chr2L | 22756255 | 22758595 | 2341 | 2,26 | 0120 |
| chr2L | 22759800 | 22766838 | 7039 | 3,69 | 0121 |
| chr2L | 22773843 | 22774946 | 1104 | 2,1 | 0122 |
| chr2L | 22807579 | 22808677 | 1099 | 2,07 | 0123 |
| chr2L | 22812700 | 22817111 | 4412 | 3,99 | 0124 |
| chr2L | 22822029 | 22828111 | 6083 | 3,01 | 0125 |
| chr2L | 22828296 | 22829514 | 1219 | 2,82 | 0126 |
| chr2L | 22829673 | 22832262 | 2590 | 2,88 | 0127 |
| chr2L | 22841641 | 22849223 | 7583 | 3,02 | 0128 |
| chr2L | 22849378 | 22850557 | 1180 | 2,17 | 0129 |
| chr2L | 22850811 | 22855590 | 4780 | 3,2 | 0130 |
| chr2L | 22855885 | 22858667 | 2783 | 2,69 | 0131 |
| chr2L | 22858891 | 22860117 | 1227 | 2,39 | 0132 |
| chr2L | 22860489 | 22869013 | 8525 | 2,83 | 0133 |
| chr2L | 22869320 | 22871296 | 1977 | 2,28 | 0134 |
| chr2L | 22879343 | 22885085 | 5743 | 2,09 | 0135 |
| chr2L | 22886974 | 22890572 | 3599 | 2,45 | 0136 |
| chr2L | 22917761 | 22922599 | 4839 | 3,34 | 0137 |
| chr2L | 22922935 | 22935320 | 12386 | 3,06 | 0138 |
| chr2L | 22937539 | 22942132 | 4594 | 3,72 | 0139 |
| chr2L | 22942244 | 22946768 | 4525 | 3,42 | 0140 |
| chr2L | 22947353 | 22948744 | 1392 | 2,39 | 0141 |

| | | | | | |
|---|---|---|---|---|---|
| chr2L | 22948961 | 22950455 | 1495 | 2,33 | 0142 |
| chr2L | 22956824 | 22959033 | 2210 | 2,59 | 0143 |
| chr2L | 22962576 | 22968183 | 5608 | 2,38 | 0144 |
| chr2L | 22968357 | 22979454 | 11098 | 3,26 | 0145 |
| chr2L | 22980076 | 22985388 | 5313 | 3,25 | 0146 |
| chr2LHet | 38876 | 42584 | 3709 | 2,87 | 0147 |
| chr2LHet | 45029 | 48425 | 3397 | 3,29 | 0148 |
| chr2LHet | 49422 | 53132 | 3711 | 3,79 | 0149 |
| chr2LHet | 133434 | 135788 | 2355 | 2,82 | 0150 |
| chr2LHet | 140251 | 144462 | 4212 | 3,11 | 0151 |
| chr2LHet | 150684 | 155688 | 5005 | 3,37 | 0152 |
| chr2LHet | 155846 | 159327 | 3482 | 2,98 | 0153 |
| chr2LHet | 159886 | 161820 | 1935 | 3,4 | 0154 |
| chr2LHet | 162712 | 168036 | 5325 | 2,85 | 0155 |
| chr2LHet | 169566 | 172489 | 2924 | 2,15 | 0156 |
| chr2LHet | 172810 | 174753 | 1944 | 2,94 | 0157 |
| chr2LHet | 175006 | 178703 | 3698 | 2,94 | 0158 |
| chr2LHet | 180398 | 184320 | 3923 | 3,32 | 0159 |
| chr2LHet | 184659 | 197924 | 13266 | 3,99 | 0160 |
| chr2LHet | 213250 | 214424 | 1175 | 2,3 | 0161 |
| chr2LHet | 236525 | 243055 | 6531 | 2,64 | 0162 |
| chr2LHet | 243125 | 247253 | 4129 | 3,19 | 0163 |
| chr2LHet | 247627 | 253299 | 5673 | 3,26 | 0164 |
| chr2LHet | 254457 | 261123 | 6667 | 3,43 | 0165 |
| chr2LHet | 272065 | 280548 | 8484 | 3,5 | 0166 |
| chr2LHet | 281612 | 294749 | 13138 | 3,03 | 0167 |
| chr2LHet | 295485 | 302854 | 7370 | 2,17 | 0168 |
| chr2LHet | 302943 | 306218 | 3276 | 2,63 | 0169 |
| chr2LHet | 306936 | 310593 | 3658 | 3,41 | 0170 |
| chr2LHet | 312131 | 316249 | 4119 | 2,62 | 0171 |
| chr2LHet | 317343 | 319083 | 1741 | 2,94 | 0172 |
| chr2LHet | 321161 | 322272 | 1112 | 2,62 | 0173 |
| chr2LHet | 327733 | 328984 | 1252 | 2,78 | 0174 |
| chr2LHet | 330297 | 331508 | 1212 | 2,23 | 0175 |
| chr2LHet | 332062 | 336762 | 4701 | 3,67 | 0176 |
| chr2LHet | 337030 | 342016 | 4987 | 2,75 | 0177 |
| chr2LHet | 342266 | 348195 | 5930 | 3,33 | 0178 |
| chr2LHet | 348459 | 350853 | 2395 | 2,99 | 0179 |
| chr2LHet | 354764 | 356219 | 1456 | 2,1 | 0180 |
| chr2LHet | 358890 | 361861 | 2972 | 2,94 | 0181 |
| chr2R | 12914 | 14533 | 1620 | 1,97 | 0182 |
| chr2R | 22185 | 24260 | 2076 | 2,56 | 0183 |
| chr2R | 26750 | 29413 | 2664 | 2,56 | 0184 |
| chr2R | 38303 | 40023 | 1721 | 2,36 | 0185 |
| chr2R | 40229 | 44177 | 3949 | 2,75 | 0186 |
| chr2R | 44613 | 47282 | 2670 | 2,26 | 0187 |
| chr2R | 50086 | 52548 | 2463 | 2,49 | 0188 |
| chr2R | 53693 | 55074 | 1382 | 2,17 | 0189 |

| | | | | | |
|---|---|---|---|---|---|
| chr2R | 55642 | 57177 | 1536 | 2,62 | 0190 |
| chr2R | 58606 | 60974 | 2369 | 3,12 | 0191 |
| chr2R | 67730 | 70043 | 2314 | 2,26 | 0192 |
| chr2R | 79254 | 81108 | 1855 | 2,29 | 0193 |
| chr2R | 82603 | 86555 | 3953 | 2,07 | 0194 |
| chr2R | 87679 | 88755 | 1077 | 2,49 | 0195 |
| chr2R | 88854 | 92430 | 3577 | 2,59 | 0196 |
| chr2R | 92601 | 94813 | 2213 | 2,52 | 0197 |
| chr2R | 120169 | 123080 | 2912 | 2,59 | 0198 |
| chr2R | 132829 | 135553 | 2725 | 3,1 | 0199 |
| chr2R | 137051 | 142976 | 5926 | 2,48 | 0200 |
| chr2R | 150835 | 152195 | 1361 | 2,46 | 0201 |
| chr2R | 157850 | 160454 | 2605 | 2,36 | 0202 |
| chr2R | 171969 | 173717 | 1749 | 2,49 | 0203 |
| chr2R | 174496 | 175794 | 1299 | 2,23 | 0204 |
| chr2R | 176426 | 177615 | 1190 | 2,59 | 0205 |
| chr2R | 177753 | 181859 | 4107 | 3,12 | 0206 |
| chr2R | 198160 | 201933 | 3774 | 2,43 | 0207 |
| chr2R | 204898 | 205997 | 1100 | 2,75 | 0208 |
| chr2R | 211313 | 217463 | 6151 | 2,49 | 0209 |
| chr2R | 222007 | 227130 | 5124 | 2,59 | 0210 |
| chr2R | 227302 | 228677 | 1376 | 2,49 | 0211 |
| chr2R | 229709 | 231289 | 1581 | 2,62 | 0212 |
| chr2R | 231779 | 242445 | 10667 | 2,88 | 0213 |
| chr2R | 248401 | 251224 | 2824 | 2,69 | 0214 |
| chr2R | 253296 | 255145 | 1850 | 2,14 | 0215 |
| chr2R | 255241 | 256408 | 1168 | 2,72 | 0216 |
| chr2R | 275852 | 278514 | 2663 | 2,12 | 0217 |
| chr2R | 278617 | 281333 | 2717 | 2,23 | 0218 |
| chr2R | 285162 | 286203 | 1042 | 2,2 | 0219 |
| chr2R | 290103 | 295234 | 5132 | 2,78 | 0220 |
| chr2R | 295405 | 298607 | 3203 | 2,36 | 0221 |
| chr2R | 299137 | 301846 | 2710 | 3,11 | 0222 |
| chr2R | 312474 | 314607 | 2134 | 2,88 | 0223 |
| chr2R | 314756 | 319718 | 4963 | 2,98 | 0224 |
| chr2R | 326326 | 329516 | 3191 | 2,94 | 0225 |
| chr2R | 337043 | 339703 | 2661 | 2,49 | 0226 |
| chr2R | 339910 | 341256 | 1347 | 2,43 | 0227 |
| chr2R | 359820 | 365951 | 6132 | 3,19 | 0228 |
| chr2R | 366341 | 368662 | 2322 | 2,65 | 0229 |
| chr2R | 376207 | 378065 | 1859 | 2,26 | 0230 |
| chr2R | 380515 | 382859 | 2345 | 2,69 | 0231 |
| chr2R | 385466 | 387489 | 2024 | 2,52 | 0232 |
| chr2R | 387671 | 395240 | 7570 | 3,04 | 0233 |
| chr2R | 406933 | 408330 | 1398 | 2,23 | 0234 |
| chr2R | 410827 | 412431 | 1605 | 2,84 | 0235 |
| chr2R | 413752 | 415712 | 1961 | 2,33 | 0236 |
| chr2R | 420342 | 423449 | 3108 | 2,65 | 0237 |

| | | | | | |
|---|---|---|---|---|---|
| chr2R | 423911 | 426970 | 3060 | 2,52 | 0238 |
| chr2R | 444989 | 446361 | 1373 | 2,33 | 0239 |
| chr2R | 453256 | 456275 | 3020 | 2,49 | 0240 |
| chr2R | 461866 | 463193 | 1328 | 2,04 | 0241 |
| chr2R | 465899 | 467825 | 1927 | 2,46 | 0242 |
| chr2R | 469455 | 471749 | 2295 | 2,26 | 0243 |
| chr2R | 477598 | 479215 | 1618 | 2,14 | 0244 |
| chr2R | 497160 | 502140 | 4981 | 2,52 | 0245 |
| chr2R | 517898 | 520159 | 2262 | 2,56 | 0246 |
| chr2R | 522528 | 523921 | 1394 | 2,46 | 0247 |
| chr2R | 524332 | 525741 | 1410 | 2,69 | 0248 |
| chr2R | 538810 | 540239 | 1430 | 2,56 | 0249 |
| chr2R | 542321 | 546629 | 4309 | 3,9 | 0250 |
| chr2R | 546724 | 547884 | 1161 | 2,07 | 0251 |
| chr2R | 561672 | 563713 | 2042 | 2,23 | 0252 |
| chr2R | 565418 | 567161 | 1744 | 2,78 | 0253 |
| chr2R | 575719 | 578255 | 2537 | 2,33 | 0254 |
| chr2R | 591147 | 594692 | 3546 | 2,85 | 0255 |
| chr2R | 595794 | 597706 | 1913 | 2,59 | 0256 |
| chr2R | 599213 | 601889 | 2677 | 2,23 | 0257 |
| chr2R | 604711 | 605772 | 1062 | 2,26 | 0258 |
| chr2R | 627445 | 630870 | 3426 | 2,61 | 0259 |
| chr2R | 653295 | 654338 | 1044 | 2,69 | 0260 |
| chr2R | 656890 | 663274 | 6385 | 2,78 | 0261 |
| chr2R | 663488 | 666721 | 3234 | 2,62 | 0262 |
| chr2R | 679568 | 683553 | 3986 | 3,04 | 0263 |
| chr2R | 683644 | 685517 | 1874 | 2,39 | 0264 |
| chr2R | 686765 | 688822 | 2058 | 2,17 | 0265 |
| chr2R | 691251 | 692305 | 1055 | 2,36 | 0266 |
| chr2R | 692779 | 697499 | 4721 | 2,46 | 0267 |
| chr2R | 721383 | 724405 | 3023 | 2,3 | 0268 |
| chr2R | 724672 | 726579 | 1908 | 2,1 | 0269 |
| chr2R | 727603 | 728811 | 1209 | 2,62 | 0270 |
| chr2R | 736014 | 737818 | 1805 | 2,2 | 0271 |
| chr2R | 738796 | 740280 | 1485 | 2,3 | 0272 |
| chr2R | 740321 | 747856 | 7536 | 3,88 | 0273 |
| chr2R | 756719 | 758315 | 1597 | 1,94 | 0274 |
| chr2R | 758665 | 760909 | 2245 | 2,43 | 0275 |
| chr2R | 761242 | 762795 | 1554 | 2,56 | 0276 |
| chr2R | 763092 | 768919 | 5828 | 2,49 | 0277 |
| chr2R | 773622 | 780482 | 6861 | 3,4 | 0278 |
| chr2R | 780539 | 782569 | 2031 | 2,56 | 0279 |
| chr2R | 783114 | 787969 | 4856 | 2,52 | 0280 |
| chr2R | 788309 | 789906 | 1598 | 2,49 | 0281 |
| chr2R | 790683 | 791875 | 1193 | 2,69 | 0282 |
| chr2R | 794341 | 797256 | 2916 | 2,39 | 0283 |
| chr2R | 799967 | 802817 | 2851 | 2,65 | 0284 |
| chr2R | 803162 | 804576 | 1415 | 2,33 | 0285 |

| | | | | | |
|---|---|---|---|---|---|
| chr2R | 867810 | 871815 | 4006 | 2,26 | 0286 |
| chr2R | 878240 | 879628 | 1389 | 2,17 | 0287 |
| chr2R | 885734 | 888262 | 2529 | 2,36 | 0288 |
| chr2R | 888368 | 890558 | 2191 | 2,52 | 0289 |
| chr2R | 891045 | 892790 | 1746 | 2,26 | 0290 |
| chr2R | 893035 | 894795 | 1761 | 2,3 | 0291 |
| chr2R | 895621 | 897637 | 2017 | 1,94 | 0292 |
| chr2R | 897804 | 899906 | 2103 | 2,36 | 0293 |
| chr2R | 904082 | 905397 | 1316 | 2,49 | 0294 |
| chr2R | 905430 | 906482 | 1053 | 2,3 | 0295 |
| chr2R | 914934 | 921855 | 6922 | 4,01 | 0296 |
| chr2R | 923114 | 926182 | 3069 | 2,33 | 0297 |
| chr2R | 928743 | 931180 | 2438 | 2,14 | 0298 |
| chr2R | 936864 | 944038 | 7175 | 1,88 | 0299 |
| chr2R | 949666 | 951519 | 1854 | 2,33 | 0300 |
| chr2R | 952547 | 953598 | 1052 | 2,78 | 0301 |
| chr2R | 953758 | 955214 | 1457 | 2,59 | 0302 |
| chr2R | 958208 | 959358 | 1151 | 2,07 | 0303 |
| chr2R | 960416 | 962057 | 1642 | 2,56 | 0304 |
| chr2R | 963227 | 966444 | 3218 | 2,3 | 0305 |
| chr2R | 966772 | 970986 | 4215 | 2,43 | 0306 |
| chr2R | 974316 | 975903 | 1588 | 2,17 | 0307 |
| chr2R | 982651 | 984397 | 1747 | 2,94 | 0308 |
| chr2R | 985713 | 988623 | 2911 | 2,14 | 0309 |
| chr2R | 988844 | 992023 | 3180 | 2,62 | 0310 |
| chr2R | 992539 | 993992 | 1454 | 2,1 | 0311 |
| chr2R | 995095 | 997822 | 2728 | 2,46 | 0312 |
| chr2R | 997916 | 1003598 | 5683 | 2,46 | 0313 |
| chr2R | 1018804 | 1022885 | 4082 | 2,6 | 0314 |
| chr2R | 1025870 | 1027724 | 1855 | 2,01 | 0315 |
| chr2R | 1041990 | 1044738 | 2749 | 2,75 | 0316 |
| chr2R | 1044870 | 1049069 | 4200 | 2,3 | 0317 |
| chr2R | 1057244 | 1059351 | 2108 | 2,72 | 0318 |
| chr2R | 1062152 | 1067476 | 5325 | 2,56 | 0319 |
| chr2R | 1074194 | 1075792 | 1599 | 2,07 | 0320 |
| chr2R | 1078396 | 1079641 | 1246 | 2,04 | 0321 |
| chr2R | 1092651 | 1094579 | 1929 | 2,01 | 0322 |
| chr2R | 1096009 | 1097450 | 1442 | 2,46 | 0323 |
| chr2R | 1097768 | 1100814 | 3047 | 2,56 | 0324 |
| chr2R | 1110396 | 1113942 | 3547 | 2,28 | 0325 |
| chr2R | 1114030 | 1115380 | 1351 | 2,46 | 0326 |
| chr2R | 1121329 | 1123451 | 2123 | 2,3 | 0327 |
| chr2R | 1123510 | 1127034 | 3525 | 2,88 | 0328 |
| chr2R | 1128408 | 1130061 | 1654 | 1,98 | 0329 |
| chr2R | 1140462 | 1142784 | 2323 | 2,2 | 0330 |
| chr2R | 1142893 | 1144981 | 2089 | 2,1 | 0331 |
| chr2R | 1185133 | 1186209 | 1077 | 2,25 | 0332 |
| chr2R | 1187107 | 1190168 | 3062 | 1,93 | 0333 |

| | | | | | |
|---|---|---|---|---|---|
| chr2R | 1190417 | 1192226 | 1810 | 2,08 | 0334 |
| chr2R | 1217041 | 1222476 | 5436 | 2,52 | 0335 |
| chr2R | 1252056 | 1253739 | 1684 | 2,46 | 0336 |
| chr2R | 1285693 | 1289711 | 4019 | 2,99 | 0337 |
| chr2R | 1289990 | 1295560 | 5571 | 2,51 | 0338 |
| chr2R | 1297394 | 1298544 | 1151 | 2,98 | 0339 |
| chr2R | 1300168 | 1302109 | 1942 | 2,22 | 0340 |
| chr2R | 1314175 | 1315339 | 1165 | 2,49 | 0341 |
| chr2R | 1318745 | 1320028 | 1284 | 2,43 | 0342 |
| chr2R | 1335780 | 1337343 | 1564 | 2,07 | 0343 |
| chr2R | 1338976 | 1340537 | 1562 | 2,14 | 0344 |
| chr2R | 1341775 | 1342881 | 1107 | 2,33 | 0345 |
| chr2R | 1344718 | 1348944 | 4227 | 2,3 | 0346 |
| chr2R | 1350235 | 1356770 | 6536 | 2,66 | 0347 |
| chr2R | 1356867 | 1359889 | 3023 | 2,72 | 0348 |
| chr2R | 1361138 | 1363006 | 1869 | 2,33 | 0349 |
| chr2R | 1363083 | 1365757 | 2675 | 2,36 | 0350 |
| chr2R | 1365983 | 1368507 | 2525 | 2,33 | 0351 |
| chr2R | 1374235 | 1377866 | 3632 | 2,52 | 0352 |
| chr2R | 1387631 | 1389310 | 1680 | 2,1 | 0353 |
| chr2R | 1391843 | 1393062 | 1220 | 1,94 | 0354 |
| chr2R | 1400982 | 1403445 | 2464 | 2,3 | 0355 |
| chr2R | 1419081 | 1420174 | 1094 | 2,56 | 0356 |
| chr2R | 1420805 | 1423203 | 2399 | 2,07 | 0357 |
| chr2R | 1426388 | 1429317 | 2930 | 2,46 | 0358 |
| chr2R | 1434934 | 1436667 | 1734 | 2,23 | 0359 |
| chr2R | 1438393 | 1440524 | 2132 | 2,14 | 0360 |
| chr2R | 1441803 | 1443998 | 2196 | 2,23 | 0361 |
| chr2R | 1464799 | 1467273 | 2475 | 2,46 | 0362 |
| chr2R | 1469754 | 1471044 | 1291 | 2,17 | 0363 |
| chr2R | 1473963 | 1475974 | 2012 | 2,36 | 0364 |
| chr2R | 1484950 | 1486168 | 1219 | 2,36 | 0365 |
| chr2R | 1499695 | 1503444 | 3750 | 2,33 | 0366 |
| chr2R | 1515245 | 1517898 | 2654 | 1,94 | 0367 |
| chr2R | 1520199 | 1521750 | 1552 | 2,1 | 0368 |
| chr2R | 1521903 | 1524524 | 2622 | 2,46 | 0369 |
| chr2R | 1524686 | 1529213 | 4528 | 2,64 | 0370 |
| chr2R | 1758113 | 1759138 | 1026 | 2,17 | 0371 |
| chr2R | 2175951 | 2177271 | 1321 | 2,39 | 0372 |
| chr2R | 2187730 | 2188785 | 1056 | 2,72 | 0373 |
| chr2R | 2209628 | 2212038 | 2411 | 1,97 | 0374 |
| chr2R | 2214551 | 2216356 | 1806 | 2,26 | 0375 |
| chr2R | 2234030 | 2239603 | 5574 | 1,86 | 0376 |
| chr2R | 2286196 | 2287277 | 1082 | 2,43 | 0377 |
| chr2R | 2287297 | 2290062 | 2766 | 2,55 | 0378 |
| chr2R | 2311865 | 2313992 | 2128 | 2,52 | 0379 |
| chr2R | 2320372 | 2324650 | 4279 | 2,62 | 0380 |
| chr2R | 2347931 | 2349818 | 1888 | 2,23 | 0381 |

| | | | | | |
|---|---|---|---|---|---|
| chr2R | 2352820 | 2355469 | 2650 | 2,33 | 0382 |
| chr2R | 2368916 | 2370094 | 1179 | 2,37 | 0383 |
| chr2R | 2370142 | 2371610 | 1469 | 2,14 | 0384 |
| chr2R | 2374082 | 2375714 | 1633 | 1,94 | 0385 |
| chr2R | 2378638 | 2381184 | 2547 | 2,1 | 0386 |
| chr2R | 2468465 | 2470612 | 2148 | 2,33 | 0387 |
| chr2R | 2840358 | 2845857 | 5500 | 2,04 | 0388 |
| chr2R | 3210885 | 3215754 | 4870 | 2,4 | 0389 |
| chr2R | 3218315 | 3221607 | 3293 | 2,17 | 0390 |
| chr2R | 3523628 | 3535006 | 11379 | 2,29 | 0391 |
| chr2R | 3683480 | 3687498 | 4019 | 1,66 | 0392 |
| chr2R | 3830417 | 3840601 | 10185 | 2,63 | 0393 |
| chr2R | 4098912 | 4100631 | 1720 | 1,97 | 0394 |
| chr2R | 4105002 | 4108813 | 3812 | 2,39 | 0395 |
| chr2R | 4378165 | 4379809 | 1645 | 2,01 | 0396 |
| chr2R | 5199581 | 5205761 | 6181 | 2,79 | 0397 |
| chr2R | 5207224 | 5209282 | 2059 | 2,19 | 0398 |
| chr2R | 8389471 | 8391244 | 1774 | 2,43 | 0399 |
| chr2R | 8750743 | 8753796 | 3054 | 2,46 | 0400 |
| chr2R | 9616845 | 9623014 | 6170 | 2,06 | 0401 |
| chr2R | 9772790 | 9779223 | 6434 | 2,82 | 0402 |
| chr2R | 9989151 | 9995111 | 5961 | 3,01 | 0403 |
| chr2R | 10777082 | 10781244 | 4163 | 2,91 | 0404 |
| chr2R | 10781492 | 10783620 | 2129 | 2,19 | 0405 |
| chr2R | 12641532 | 12643381 | 1850 | 2,33 | 0406 |
| chr2R | 12647084 | 12649408 | 2325 | 2,07 | 0407 |
| chr2R | 14745457 | 14747902 | 2446 | 3,04 | 0408 |
| chr2R | 15957211 | 15958756 | 1546 | 2,59 | 0409 |
| chr2R | 16667458 | 16670495 | 3038 | 3,94 | 0410 |
| chr2R | 18575797 | 18577671 | 1875 | 8,61 | 0411 |
| chr2R | 18847721 | 18849499 | 1779 | 2,06 | 0412 |
| chr2RHet | 4 | 1984 | 1981 | 2,91 | 0413 |
| chr2RHet | 6537 | 9537 | 3001 | 2,56 | 0414 |
| chr2RHet | 9555 | 17389 | 7835 | 2,78 | 0415 |
| chr2RHet | 17442 | 20110 | 2669 | 3,2 | 0416 |
| chr2RHet | 25958 | 31016 | 5059 | 2,35 | 0417 |
| chr2RHet | 44968 | 48314 | 3347 | 2,62 | 0418 |
| chr2RHet | 79363 | 81345 | 1983 | 3,39 | 0419 |
| chr2RHet | 86622 | 96044 | 9423 | 3,11 | 0420 |
| chr2RHet | 96338 | 104337 | 8000 | 2,94 | 0421 |
| chr2RHet | 104434 | 109466 | 5033 | 2,98 | 0422 |
| chr2RHet | 131263 | 146654 | 15392 | 2,91 | 0423 |
| chr2RHet | 147192 | 148705 | 1514 | 2,52 | 0424 |
| chr2RHet | 149931 | 156276 | 6346 | 3,04 | 0425 |
| chr2RHet | 156319 | 157958 | 1640 | 2,65 | 0426 |
| chr2RHet | 158285 | 164684 | 6400 | 2,91 | 0427 |
| chr2RHet | 164926 | 168086 | 3161 | 2,78 | 0428 |
| chr2RHet | 168401 | 174686 | 6286 | 2,78 | 0429 |

| | | | | | |
|---|---|---|---|---|---|
| chr2RHet | 174908 | 179390 | 4483 | 3,09 | 0430 |
| chr2RHet | 182018 | 185722 | 3705 | 2,6 | 0431 |
| chr2RHet | 188520 | 192341 | 3822 | 2,98 | 0432 |
| chr2RHet | 193327 | 196365 | 3039 | 2,36 | 0433 |
| chr2RHet | 215577 | 216866 | 1290 | 2,28 | 0434 |
| chr2RHet | 217014 | 221345 | 4332 | 2,39 | 0435 |
| chr2RHet | 222091 | 223947 | 1857 | 2,43 | 0436 |
| chr2RHet | 224492 | 226188 | 1697 | 1,91 | 0437 |
| chr2RHet | 228604 | 230351 | 1748 | 2,07 | 0438 |
| chr2RHet | 230926 | 232551 | 1626 | 2,1 | 0439 |
| chr2RHet | 233521 | 236257 | 2737 | 2,36 | 0440 |
| chr2RHet | 292311 | 293402 | 1092 | 2,56 | 0441 |
| chr2RHet | 294966 | 299131 | 4166 | 2,46 | 0442 |
| chr2RHet | 300019 | 302865 | 2847 | 2,3 | 0443 |
| chr2RHet | 302893 | 308711 | 5819 | 3,04 | 0444 |
| chr2RHet | 310661 | 312138 | 1478 | 2,88 | 0445 |
| chr2RHet | 312269 | 313362 | 1094 | 2,72 | 0446 |
| chr2RHet | 317616 | 319556 | 1941 | 2,78 | 0447 |
| chr2RHet | 324290 | 326570 | 2281 | 2,26 | 0448 |
| chr2RHet | 327549 | 328947 | 1399 | 2,62 | 0449 |
| chr2RHet | 345426 | 348340 | 2915 | 2,39 | 0450 |
| chr2RHet | 351170 | 354306 | 3137 | 2,75 | 0451 |
| chr2RHet | 371032 | 375327 | 4296 | 2,98 | 0452 |
| chr2RHet | 377542 | 380895 | 3354 | 2,33 | 0453 |
| chr2RHet | 381664 | 382817 | 1154 | 2,14 | 0454 |
| chr2RHet | 395722 | 397845 | 2124 | 2,85 | 0455 |
| chr2RHet | 398175 | 406919 | 8745 | 3,04 | 0456 |
| chr2RHet | 408263 | 412523 | 4261 | 2,69 | 0457 |
| chr2RHet | 412628 | 422898 | 10271 | 3,12 | 0458 |
| chr2RHet | 422909 | 429593 | 6685 | 2,73 | 0459 |
| chr2RHet | 429670 | 435172 | 5503 | 3,69 | 0460 |
| chr2RHet | 449244 | 452091 | 2848 | 2,65 | 0461 |
| chr2RHet | 463664 | 466082 | 2419 | 2,77 | 0462 |
| chr2RHet | 466092 | 468551 | 2460 | 2,27 | 0463 |
| chr2RHet | 473300 | 475631 | 2332 | 2,65 | 0464 |
| chr2RHet | 477681 | 479073 | 1393 | 2,1 | 0465 |
| chr2RHet | 493700 | 495646 | 1947 | 2,55 | 0466 |
| chr2RHet | 495682 | 500654 | 4973 | 3,17 | 0467 |
| chr2RHet | 501090 | 505638 | 4549 | 2,82 | 0468 |
| chr2RHet | 505733 | 510440 | 4708 | 2,39 | 0469 |
| chr2RHet | 512267 | 517992 | 5726 | 2,65 | 0470 |
| chr2RHet | 518396 | 519562 | 1167 | 2,49 | 0471 |
| chr2RHet | 519676 | 523311 | 3636 | 2,78 | 0472 |
| chr2RHet | 523676 | 535723 | 12048 | 2,98 | 0473 |
| chr2RHet | 536727 | 540721 | 3995 | 2,75 | 0474 |
| chr2RHet | 543859 | 548264 | 4406 | 2,91 | 0475 |
| chr2RHet | 549012 | 556104 | 7093 | 3,04 | 0476 |
| chr2RHet | 556576 | 559333 | 2758 | 2,78 | 0477 |

| | | | | | |
|---|---|---|---|---|---|
| chr2RHet | 559478 | 560628 | 1151 | 2,3 | 0478 |
| chr2RHet | 562660 | 579098 | 16439 | 3,4 | 0479 |
| chr2RHet | 585715 | 591551 | 5837 | 2,33 | 0480 |
| chr2RHet | 591942 | 594482 | 2541 | 2,21 | 0481 |
| chr2RHet | 596133 | 598193 | 2061 | 2,26 | 0482 |
| chr2RHet | 599023 | 600403 | 1381 | 2,33 | 0483 |
| chr2RHet | 600426 | 606287 | 5862 | 2,56 | 0484 |
| chr2RHet | 608949 | 610350 | 1402 | 2,26 | 0485 |
| chr2RHet | 610632 | 613534 | 2903 | 2,36 | 0486 |
| chr2RHet | 614485 | 616304 | 1820 | 3,63 | 0487 |
| chr2RHet | 616326 | 619643 | 3318 | 2,85 | 0488 |
| chr2RHet | 620758 | 625169 | 4412 | 2,52 | 0489 |
| chr2RHet | 625389 | 628042 | 2654 | 2,85 | 0490 |
| chr2RHet | 628432 | 636154 | 7723 | 3,01 | 0491 |
| chr2RHet | 638533 | 641181 | 2649 | 2,33 | 0492 |
| chr2RHet | 641468 | 643102 | 1635 | 2,62 | 0493 |
| chr2RHet | 643201 | 648362 | 5162 | 2,75 | 0494 |
| chr2RHet | 648503 | 650188 | 1686 | 2,62 | 0495 |
| chr2RHet | 657466 | 658562 | 1097 | 1,75 | 0496 |
| chr2RHet | 658838 | 675787 | 16950 | 3,17 | 0497 |
| chr2RHet | 725133 | 730386 | 5254 | 2,85 | 0498 |
| chr2RHet | 737478 | 743016 | 5539 | 2,88 | 0499 |
| chr2RHet | 763003 | 767049 | 4047 | 2,69 | 0500 |
| chr2RHet | 767542 | 769980 | 2439 | 2,91 | 0501 |
| chr2RHet | 771586 | 774363 | 2778 | 2,9 | 0502 |
| chr2RHet | 777465 | 782150 | 4686 | 2,65 | 0503 |
| chr2RHet | 782953 | 787278 | 4326 | 2,62 | 0504 |
| chr2RHet | 787574 | 797743 | 10170 | 3,17 | 0505 |
| chr2RHet | 798311 | 800956 | 2646 | 2,62 | 0506 |
| chr2RHet | 801001 | 805148 | 4148 | 2,52 | 0507 |
| chr2RHet | 817197 | 822630 | 5434 | 2,91 | 0508 |
| chr2RHet | 822686 | 826883 | 4198 | 2,82 | 0509 |
| chr2RHet | 827090 | 830394 | 3305 | 2,52 | 0510 |
| chr2RHet | 836044 | 839034 | 2991 | 2,65 | 0511 |
| chr2RHet | 841844 | 843467 | 1624 | 2,59 | 0512 |
| chr2RHet | 866478 | 868831 | 2354 | 2,62 | 0513 |
| chr2RHet | 868993 | 871909 | 2917 | 3,27 | 0514 |
| chr2RHet | 872233 | 873606 | 1374 | 2,62 | 0515 |
| chr2RHet | 880576 | 882445 | 1870 | 2,62 | 0516 |
| chr2RHet | 882977 | 884115 | 1139 | 2,36 | 0517 |
| chr2RHet | 884217 | 886475 | 2259 | 2,91 | 0518 |
| chr2RHet | 886516 | 890845 | 4330 | 2,52 | 0519 |
| chr2RHet | 899785 | 901432 | 1648 | 2,36 | 0520 |
| chr2RHet | 907109 | 912249 | 5141 | 2,36 | 0521 |
| chr2RHet | 913827 | 923268 | 9442 | 2,82 | 0522 |
| chr2RHet | 932636 | 937352 | 4717 | 2,94 | 0523 |
| chr2RHet | 938599 | 940615 | 2017 | 2,49 | 0524 |
| chr2RHet | 940994 | 942324 | 1331 | 2,01 | 0525 |

| | | | | | |
|---|---|---|---|---|---|
| chr2RHet | 942530 | 943824 | 1295 | 2,14 | 0526 |
| chr2RHet | 947918 | 954451 | 6534 | 2,85 | 0527 |
| chr2RHet | 954714 | 960692 | 5979 | 2,65 | 0528 |
| chr2RHet | 961322 | 966395 | 5074 | 2,82 | 0529 |
| chr2RHet | 966975 | 972146 | 5172 | 2,91 | 0530 |
| chr2RHet | 973739 | 977840 | 4102 | 2,98 | 0531 |
| chr2RHet | 999217 | 1000226 | 1010 | 3,01 | 0532 |
| chr2RHet | 1000465 | 1001975 | 1511 | 2,78 | 0533 |
| chr2RHet | 1003810 | 1005798 | 1989 | 3,37 | 0534 |
| chr2RHet | 1007201 | 1008633 | 1433 | 2,46 | 0535 |
| chr2RHet | 1009229 | 1012287 | 3059 | 2,49 | 0536 |
| chr2RHet | 1039268 | 1044272 | 5005 | 3,03 | 0537 |
| chr2RHet | 1076692 | 1078524 | 1833 | 2,94 | 0538 |
| chr2RHet | 1078691 | 1082787 | 4097 | 2,52 | 0539 |
| chr2RHet | 1083117 | 1091996 | 8880 | 2,94 | 0540 |
| chr2RHet | 1093930 | 1095629 | 1700 | 2,04 | 0541 |
| chr2RHet | 1111014 | 1113762 | 2749 | 3,04 | 0542 |
| chr2RHet | 1131178 | 1133766 | 2589 | 2,49 | 0543 |
| chr2RHet | 1134084 | 1136145 | 2062 | 2,49 | 0544 |
| chr2RHet | 1136991 | 1142762 | 5772 | 2,37 | 0545 |
| chr2RHet | 1268424 | 1269612 | 1189 | 2,52 | 0546 |
| chr2RHet | 1278861 | 1282489 | 3629 | 2,43 | 0547 |
| chr2RHet | 1285100 | 1296948 | 11849 | 3,11 | 0548 |
| chr2RHet | 1297908 | 1300726 | 2819 | 2,62 | 0549 |
| chr2RHet | 1300787 | 1304350 | 3564 | 2,2 | 0550 |
| chr2RHet | 1304421 | 1312041 | 7621 | 2,56 | 0551 |
| chr2RHet | 1313980 | 1315141 | 1162 | 2,1 | 0552 |
| chr2RHet | 1316355 | 1326100 | 9746 | 2,75 | 0553 |
| chr2RHet | 1337966 | 1339214 | 1249 | 2,39 | 0554 |
| chr2RHet | 1342462 | 1344335 | 1874 | 3,2 | 0555 |
| chr2RHet | 1359046 | 1362837 | 3792 | 3,1 | 0556 |
| chr2RHet | 1364554 | 1367166 | 2613 | 2,88 | 0557 |
| chr2RHet | 1367410 | 1373715 | 6306 | 2,73 | 0558 |
| chr2RHet | 1373780 | 1378044 | 4265 | 3 | 0559 |
| chr2RHet | 1384607 | 1386240 | 1634 | 2,07 | 0560 |
| chr2RHet | 1388269 | 1391561 | 3293 | 2,94 | 0561 |
| chr2RHet | 1396822 | 1400438 | 3617 | 2,91 | 0562 |
| chr2RHet | 1405289 | 1409095 | 3807 | 3,04 | 0563 |
| chr2RHet | 1427248 | 1436325 | 9078 | 2,97 | 0564 |
| chr2RHet | 1436610 | 1441329 | 4720 | 2,91 | 0565 |
| chr2RHet | 1441929 | 1451025 | 9097 | 3,04 | 0566 |
| chr2RHet | 1552606 | 1555965 | 3360 | 2,36 | 0567 |
| chr2RHet | 1556061 | 1557315 | 1255 | 2,3 | 0568 |
| chr2RHet | 1559022 | 1560763 | 1742 | 2,23 | 0569 |
| chr2RHet | 1563281 | 1564925 | 1645 | 2,39 | 0570 |
| chr2RHet | 1570164 | 1572942 | 2779 | 3,2 | 0571 |
| chr2RHet | 1573941 | 1576931 | 2991 | 2,72 | 0572 |
| chr2RHet | 1577041 | 1579646 | 2606 | 2,56 | 0573 |

| | | | | | |
|---|---|---|---|---|---|
| chr2RHet | 1582138 | 1583923 | 1786 | 2,39 | 0574 |
| chr2RHet | 1589382 | 1592499 | 3118 | 2,49 | 0575 |
| chr2RHet | 1598088 | 1599331 | 1244 | 2,1 | 0576 |
| chr2RHet | 1624365 | 1625445 | 1081 | 2,33 | 0577 |
| chr2RHet | 1633935 | 1637435 | 3501 | 2,79 | 0578 |
| chr2RHet | 1638110 | 1639491 | 1382 | 2,59 | 0579 |
| chr2RHet | 1641667 | 1643225 | 1559 | 2,43 | 0580 |
| chr2RHet | 1643461 | 1645478 | 2018 | 2,36 | 0581 |
| chr2RHet | 1647690 | 1648791 | 1102 | 2,2 | 0582 |
| chr2RHet | 1649306 | 1651337 | 2032 | 2,52 | 0583 |
| chr2RHet | 1651412 | 1655962 | 4551 | 2,49 | 0584 |
| chr2RHet | 1657055 | 1661110 | 4056 | 2,62 | 0585 |
| chr2RHet | 1677572 | 1679834 | 2263 | 2,43 | 0586 |
| chr2RHet | 1680847 | 1682010 | 1164 | 2,3 | 0587 |
| chr2RHet | 1682622 | 1683908 | 1287 | 2,2 | 0588 |
| chr2RHet | 1689558 | 1691263 | 1706 | 2,01 | 0589 |
| chr2RHet | 1694673 | 1695857 | 1185 | 2,01 | 0590 |
| chr2RHet | 1697294 | 1699100 | 1807 | 2,01 | 0591 |
| chr2RHet | 1703703 | 1706827 | 3125 | 2,39 | 0592 |
| chr2RHet | 1708826 | 1710122 | 1297 | 2,26 | 0593 |
| chr2RHet | 1713029 | 1715253 | 2225 | 2,33 | 0594 |
| chr2RHet | 1716118 | 1717832 | 1715 | 2,23 | 0595 |
| chr2RHet | 1718224 | 1720511 | 2288 | 2,2 | 0596 |
| chr2RHet | 1720728 | 1721764 | 1037 | 2,26 | 0597 |
| chr2RHet | 1721787 | 1723756 | 1970 | 2,23 | 0598 |
| chr2RHet | 1724347 | 1728118 | 3772 | 2,88 | 0599 |
| chr2RHet | 1744555 | 1749822 | 5268 | 2,59 | 0600 |
| chr2RHet | 1751644 | 1753357 | 1714 | 3,49 | 0601 |
| chr2RHet | 1755480 | 1756885 | 1406 | 2,72 | 0602 |
| chr2RHet | 1852837 | 1855878 | 3042 | 2,56 | 0603 |
| chr2RHet | 1857299 | 1859709 | 2411 | 2,39 | 0604 |
| chr2RHet | 1866011 | 1867598 | 1588 | 2,23 | 0605 |
| chr2RHet | 1873239 | 1874966 | 1728 | 2,17 | 0606 |
| chr2RHet | 1875062 | 1881049 | 5988 | 2,3 | 0607 |
| chr2RHet | 1884146 | 1885582 | 1437 | 2,23 | 0608 |
| chr2RHet | 1885629 | 1887621 | 1993 | 2,17 | 0609 |
| chr2RHet | 1890077 | 1892502 | 2426 | 2,2 | 0610 |
| chr2RHet | 1895404 | 1898003 | 2600 | 2,52 | 0611 |
| chr2RHet | 1900063 | 1901160 | 1098 | 2,07 | 0612 |
| chr2RHet | 1901314 | 1902818 | 1505 | 2,07 | 0613 |
| chr2RHet | 1903238 | 1906301 | 3064 | 3,26 | 0614 |
| chr2RHet | 1906886 | 1910549 | 3664 | 2,43 | 0615 |
| chr2RHet | 1912375 | 1913723 | 1349 | 2,1 | 0616 |
| chr2RHet | 1913735 | 1915123 | 1389 | 2,78 | 0617 |
| chr2RHet | 1915913 | 1919851 | 3939 | 2,23 | 0618 |
| chr2RHet | 1920946 | 1923747 | 2802 | 2,75 | 0619 |
| chr2RHet | 1923787 | 1925001 | 1215 | 2,07 | 0620 |
| chr2RHet | 1926896 | 1930691 | 3796 | 2,91 | 0621 |

| | | | | | |
|---|---|---|---|---|---|
| chr2RHet | 1943171 | 1944996 | 1826 | 2,14 | 0622 |
| chr2RHet | 1946194 | 1949935 | 3742 | 2,49 | 0623 |
| chr2RHet | 1955644 | 1958493 | 2850 | 2,23 | 0624 |
| chr2RHet | 1958516 | 1959947 | 1432 | 2,36 | 0625 |
| chr2RHet | 1960168 | 1966409 | 6242 | 2,49 | 0626 |
| chr2RHet | 1967266 | 1969470 | 2205 | 2,3 | 0627 |
| chr2RHet | 1970080 | 1974118 | 4039 | 2,36 | 0628 |
| chr2RHet | 1974844 | 1976030 | 1187 | 2,23 | 0629 |
| chr2RHet | 1979009 | 1986504 | 7496 | 3,01 | 0630 |
| chr2RHet | 1986661 | 1987751 | 1091 | 2,33 | 0631 |
| chr2RHet | 1988061 | 1989219 | 1159 | 2,36 | 0632 |
| chr2RHet | 1989396 | 1992281 | 2886 | 2,2 | 0633 |
| chr2RHet | 2057311 | 2058873 | 1563 | 2,98 | 0634 |
| chr2RHet | 2065897 | 2067856 | 1960 | 2,01 | 0635 |
| chr2RHet | 2068582 | 2071667 | 3086 | 2,65 | 0636 |
| chr2RHet | 2072006 | 2075868 | 3863 | 2,43 | 0637 |
| chr2RHet | 2076836 | 2081028 | 4193 | 3,07 | 0638 |
| chr2RHet | 2081325 | 2083896 | 2572 | 3,11 | 0639 |
| chr2RHet | 2084534 | 2094336 | 9803 | 1,96 | 0640 |
| chr2RHet | 2127599 | 2129016 | 1418 | 2,46 | 0641 |
| chr2RHet | 2129036 | 2130773 | 1738 | 2,2 | 0642 |
| chr2RHet | 2132616 | 2138060 | 5445 | 2,75 | 0643 |
| chr2RHet | 2138119 | 2141662 | 3544 | 2,33 | 0644 |
| chr2RHet | 2151548 | 2154456 | 2909 | 2,88 | 0645 |
| chr2RHet | 2154863 | 2161677 | 6815 | 3,43 | 0646 |
| chr2RHet | 2178546 | 2179901 | 1356 | 2,2 | 0647 |
| chr2RHet | 2188025 | 2189635 | 1611 | 1,84 | 0648 |
| chr2RHet | 2194368 | 2195597 | 1230 | 2,75 | 0649 |
| chr2RHet | 2197824 | 2199148 | 1325 | 2,3 | 0650 |
| chr2RHet | 2201235 | 2204244 | 3010 | 2,33 | 0651 |
| chr2RHet | 2205387 | 2210954 | 5568 | 2,07 | 0652 |
| chr2RHet | 2212291 | 2214193 | 1903 | 2,98 | 0653 |
| chr2RHet | 2215033 | 2216972 | 1940 | 2,17 | 0654 |
| chr2RHet | 2235599 | 2239717 | 4119 | 2,59 | 0655 |
| chr2RHet | 2240396 | 2243191 | 2796 | 2,75 | 0656 |
| chr2RHet | 2244230 | 2247409 | 3180 | 2,33 | 0657 |
| chr2RHet | 2252324 | 2254211 | 1888 | 2,36 | 0658 |
| chr2RHet | 2255271 | 2256507 | 1237 | 2,65 | 0659 |
| chr2RHet | 2257755 | 2259146 | 1392 | 2,75 | 0660 |
| chr2RHet | 2261325 | 2272068 | 10744 | 3,35 | 0661 |
| chr2RHet | 2272379 | 2276963 | 4585 | 3,83 | 0662 |
| chr2RHet | 2291286 | 2292769 | 1484 | 2,81 | 0663 |
| chr2RHet | 2295209 | 2296256 | 1048 | 2,94 | 0664 |
| chr2RHet | 2296395 | 2299000 | 2606 | 2,93 | 0665 |
| chr2RHet | 2301262 | 2304211 | 2950 | 2,65 | 0666 |
| chr2RHet | 2308903 | 2312842 | 3940 | 2,78 | 0667 |
| chr2RHet | 2313394 | 2315138 | 1745 | 2,51 | 0668 |
| chr2RHet | 2315559 | 2316749 | 1191 | 2,3 | 0669 |

| | | | | | |
|---|---|---|---|---|---|
| chr2RHet | 2322685 | 2324984 | 2300 | 2,69 | 0670 |
| chr2RHet | 2325156 | 2329295 | 4140 | 2,2 | 0671 |
| chr2RHet | 2339483 | 2341746 | 2264 | 2,07 | 0672 |
| chr2RHet | 2341911 | 2343106 | 1196 | 2,82 | 0673 |
| chr2RHet | 2343648 | 2347030 | 3383 | 2,36 | 0674 |
| chr2RHet | 2348426 | 2349951 | 1526 | 2,39 | 0675 |
| chr2RHet | 2353111 | 2355848 | 2738 | 2,39 | 0676 |
| chr2RHet | 2358402 | 2361738 | 3337 | 2,14 | 0677 |
| chr2RHet | 2366592 | 2368304 | 1713 | 2,1 | 0678 |
| chr2RHet | 2370642 | 2374258 | 3617 | 2,33 | 0679 |
| chr2RHet | 2374431 | 2384808 | 10378 | 3,11 | 0680 |
| chr2RHet | 2384815 | 2388493 | 3679 | 3,24 | 0681 |
| chr2RHet | 2388718 | 2391940 | 3223 | 2,91 | 0682 |
| chr2RHet | 2393246 | 2396075 | 2830 | 2,26 | 0683 |
| chr2RHet | 2396227 | 2397627 | 1401 | 2,62 | 0684 |
| chr2RHet | 2399043 | 2400939 | 1897 | 2,72 | 0685 |
| chr2RHet | 2401208 | 2402400 | 1193 | 2,23 | 0686 |
| chr2RHet | 2402806 | 2404023 | 1218 | 2,62 | 0687 |
| chr2RHet | 2404196 | 2406377 | 2182 | 2,1 | 0688 |
| chr2RHet | 2442924 | 2448515 | 5592 | 3,01 | 0689 |
| chr2RHet | 2448818 | 2453338 | 4521 | 2,75 | 0690 |
| chr2RHet | 2454765 | 2456235 | 1471 | 2,91 | 0691 |
| chr2RHet | 2559897 | 2561428 | 1532 | 2,88 | 0692 |
| chr2RHet | 2566940 | 2569545 | 2606 | 2,36 | 0693 |
| chr2RHet | 2572181 | 2576045 | 3865 | 2,65 | 0694 |
| chr2RHet | 2584572 | 2593704 | 9133 | 3,72 | 0695 |
| chr2RHet | 2597882 | 2599731 | 1850 | 3,73 | 0696 |
| chr2RHet | 2600053 | 2603126 | 3074 | 2,3 | 0697 |
| chr2RHet | 2618554 | 2620367 | 1814 | 2,39 | 0698 |
| chr2RHet | 2621807 | 2622994 | 1188 | 2,94 | 0699 |
| chr2RHet | 2623599 | 2625629 | 2031 | 3,2 | 0700 |
| chr2RHet | 2626359 | 2627967 | 1609 | 2,43 | 0701 |
| chr2RHet | 2628579 | 2635664 | 7086 | 2,36 | 0702 |
| chr2RHet | 2641134 | 2642292 | 1159 | 2,01 | 0703 |
| chr2RHet | 2644111 | 2646710 | 2600 | 2,43 | 0704 |
| chr2RHet | 2762174 | 2765984 | 3811 | 3,07 | 0705 |
| chr2RHet | 2766137 | 2769991 | 3855 | 2,94 | 0706 |
| chr2RHet | 2770617 | 2773163 | 2547 | 2,75 | 0707 |
| chr2RHet | 2785963 | 2788703 | 2741 | 2,43 | 0708 |
| chr2RHet | 2804066 | 2805081 | 1016 | 2,39 | 0709 |
| chr2RHet | 2818721 | 2821042 | 2322 | 2,23 | 0710 |
| chr2RHet | 2821145 | 2824644 | 3500 | 2,59 | 0711 |
| chr2RHet | 2825132 | 2827001 | 1870 | 2,3 | 0712 |
| chr2RHet | 2827308 | 2828495 | 1188 | 2,33 | 0713 |
| chr2RHet | 2829066 | 2831825 | 2760 | 2,78 | 0714 |
| chr2RHet | 2833756 | 2835978 | 2223 | 2,26 | 0715 |
| chr2RHet | 2840905 | 2846508 | 5604 | 2,94 | 0716 |
| chr2RHet | 2846600 | 2848717 | 2118 | 2,23 | 0717 |

| | | | | | |
|---|---|---|---|---|---|
| chr2RHet | 2850961 | 2854013 | 3053 | 2,56 | 0718 |
| chr2RHet | 2856210 | 2858359 | 2150 | 2,69 | 0719 |
| chr2RHet | 2861267 | 2863183 | 1917 | 2,26 | 0720 |
| chr2RHet | 2869874 | 2871277 | 1404 | 1,97 | 0721 |
| chr2RHet | 2885160 | 2886781 | 1622 | 2,26 | 0722 |
| chr2RHet | 2895463 | 2898743 | 3281 | 2,26 | 0723 |
| chr2RHet | 2917729 | 2921997 | 4269 | 2,39 | 0724 |
| chr2RHet | 2926140 | 2927659 | 1520 | 2,63 | 0725 |
| chr2RHet | 2979342 | 2981131 | 1790 | 2,69 | 0726 |
| chr2RHet | 2984470 | 2985534 | 1065 | 2,88 | 0727 |
| chr2RHet | 2986738 | 2989264 | 2527 | 2,91 | 0728 |
| chr2RHet | 2990657 | 2992399 | 1743 | 2,1 | 0729 |
| chr2RHet | 2997841 | 2999229 | 1389 | 2,2 | 0730 |
| chr2RHet | 2999254 | 3000446 | 1193 | 2,23 | 0731 |
| chr2RHet | 3001241 | 3004357 | 3117 | 3,33 | 0732 |
| chr2RHet | 3005840 | 3011979 | 6140 | 2,34 | 0733 |
| chr2RHet | 3012054 | 3016282 | 4229 | 2,15 | 0734 |
| chr2RHet | 3016850 | 3019169 | 2320 | 2,88 | 0735 |
| chr2RHet | 3020469 | 3021512 | 1044 | 2,49 | 0736 |
| chr2RHet | 3021564 | 3022719 | 1156 | 2,14 | 0737 |
| chr2RHet | 3026781 | 3029183 | 2403 | 2,39 | 0738 |
| chr2RHet | 3034778 | 3036250 | 1473 | 2,56 | 0739 |
| chr2RHet | 3042293 | 3045191 | 2899 | 2,26 | 0740 |
| chr2RHet | 3049334 | 3054347 | 5014 | 2,59 | 0741 |
| chr2RHet | 3058019 | 3059773 | 1755 | 2,2 | 0742 |
| chr2RHet | 3064931 | 3067400 | 2470 | 2,36 | 0743 |
| chr2RHet | 3067458 | 3072658 | 5201 | 2,62 | 0744 |
| chr2RHet | 3073305 | 3074576 | 1272 | 2,26 | 0745 |
| chr2RHet | 3075797 | 3078746 | 2950 | 2,3 | 0746 |
| chr2RHet | 3097043 | 3098518 | 1476 | 2,2 | 0747 |
| chr2RHet | 3098732 | 3100923 | 2192 | 2,26 | 0748 |
| chr2RHet | 3102095 | 3107271 | 5177 | 3,11 | 0749 |
| chr2RHet | 3107913 | 3109593 | 1681 | 1,94 | 0750 |
| chr2RHet | 3111375 | 3114099 | 2725 | 2,46 | 0751 |
| chr2RHet | 3115046 | 3116741 | 1696 | 2,52 | 0752 |
| chr2RHet | 3117349 | 3119752 | 2404 | 3,3 | 0753 |
| chr2RHet | 3121372 | 3126476 | 5105 | 2,45 | 0754 |
| chr2RHet | 3128143 | 3129924 | 1782 | 2,46 | 0755 |
| chr2RHet | 3132368 | 3134093 | 1726 | 2,14 | 0756 |
| chr2RHet | 3134255 | 3136296 | 2042 | 2,62 | 0757 |
| chr2RHet | 3136678 | 3137734 | 1057 | 2,72 | 0758 |
| chr2RHet | 3138729 | 3140231 | 1503 | 2,2 | 0759 |
| chr2RHet | 3143208 | 3144439 | 1232 | 2,1 | 0760 |
| chr2RHet | 3150309 | 3152728 | 2420 | 2,91 | 0761 |
| chr2RHet | 3172051 | 3173177 | 1127 | 3,34 | 0762 |
| chr2RHet | 3184726 | 3190193 | 5468 | 2,62 | 0763 |
| chr2RHet | 3190235 | 3191348 | 1114 | 2,52 | 0764 |
| chr2RHet | 3191422 | 3193974 | 2553 | 2,43 | 0765 |

| | | | | | |
|---|---|---|---|---|---|
| chr2RHet | 3194243 | 3195363 | 1121 | 2,1 | 0766 |
| chr2RHet | 3202396 | 3203623 | 1228 | 2,52 | 0767 |
| chr2RHet | 3206822 | 3208232 | 1411 | 2,14 | 0768 |
| chr2RHet | 3208319 | 3209956 | 1638 | 2,43 | 0769 |
| chr2RHet | 3210455 | 3213980 | 3526 | 2,46 | 0770 |
| chr2RHet | 3217621 | 3220039 | 2419 | 3,23 | 0771 |
| chr2RHet | 3237196 | 3245321 | 8126 | 2,84 | 0772 |
| chr2RHet | 3245696 | 3247389 | 1694 | 3,27 | 0773 |
| chr2RHet | 3247443 | 3249139 | 1697 | 2,59 | 0774 |
| chr2RHet | 3250967 | 3254074 | 3108 | 3,06 | 0775 |
| chr2RHet | 3254423 | 3256109 | 1687 | 2,69 | 0776 |
| chr2RHet | 3256215 | 3260169 | 3955 | 3,41 | 0777 |
| chr2RHet | 3260301 | 3268311 | 8011 | 2,7 | 0778 |
| chr2RHet | 3269146 | 3274915 | 5770 | 1,92 | 0779 |
| chr3L | 1524576 | 1526710 | 2135 | 2,39 | 0780 |
| chr3L | 1526790 | 1531900 | 5111 | 3,04 | 0781 |
| chr3L | 1957928 | 1961201 | 3274 | 2,75 | 0782 |
| chr3L | 2063203 | 2069919 | 6717 | 2,19 | 0783 |
| chr3L | 2751908 | 2753861 | 1954 | 2,49 | 0784 |
| chr3L | 3202157 | 3210162 | 8006 | 2,72 | 0785 |
| chr3L | 3352440 | 3358078 | 5639 | 1,67 | 0786 |
| chr3L | 3360238 | 3365688 | 5451 | 2,27 | 0787 |
| chr3L | 4201030 | 4202145 | 1116 | 1,91 | 0788 |
| chr3L | 5452861 | 5454886 | 2026 | 2,77 | 0789 |
| chr3L | 5454945 | 5459457 | 4513 | 2,42 | 0790 |
| chr3L | 5885983 | 5888158 | 2176 | 2,52 | 0791 |
| chr3L | 11605777 | 11607476 | 1700 | 3,19 | 0792 |
| chr3L | 11608577 | 11611835 | 3259 | 4,17 | 0793 |
| chr3L | 13571388 | 13572796 | 1409 | 1,94 | 0794 |
| chr3L | 14714624 | 14715717 | 1094 | 2,1 | 0795 |
| chr3L | 15417491 | 15419368 | 1878 | 2 | 0796 |
| chr3L | 16709200 | 16714907 | 5708 | 3,24 | 0797 |
| chr3L | 17911869 | 17913015 | 1147 | 2,69 | 0798 |
| chr3L | 18500818 | 18502016 | 1199 | 1,91 | 0799 |
| chr3L | 19531799 | 19536871 | 5073 | 2,46 | 0800 |
| chr3L | 20002976 | 20005266 | 2291 | 2,39 | 0801 |
| chr3L | 21367015 | 21372193 | 5179 | 2,92 | 0802 |
| chr3L | 21414445 | 21417188 | 2744 | 2,36 | 0803 |
| chr3L | 21417196 | 21419906 | 2711 | 2,3 | 0804 |
| chr3L | 22476593 | 22478497 | 1905 | 2,3 | 0805 |
| chr3L | 22481847 | 22483354 | 1508 | 2,14 | 0806 |
| chr3L | 22494524 | 22496088 | 1565 | 2,2 | 0807 |
| chr3L | 22499736 | 22501289 | 1554 | 2,46 | 0808 |
| chr3L | 22503145 | 22504510 | 1366 | 2,04 | 0809 |
| chr3L | 22506355 | 22508191 | 1837 | 2,04 | 0810 |
| chr3L | 22510292 | 22512287 | 1996 | 2,33 | 0811 |
| chr3L | 22536393 | 22537547 | 1155 | 2,2 | 0812 |
| chr3L | 22538021 | 22540093 | 2073 | 2,33 | 0813 |

| | | | | | |
|---|---|---|---|---|---|
| chr3L | 22564741 | 22566479 | 1739 | 2,39 | 0814 |
| chr3L | 22567019 | 22568426 | 1408 | 2,14 | 0815 |
| chr3L | 22576881 | 22578116 | 1236 | 1,97 | 0816 |
| chr3L | 22591305 | 22592954 | 1650 | 2,14 | 0817 |
| chr3L | 22596137 | 22597358 | 1222 | 2,33 | 0818 |
| chr3L | 22597913 | 22599981 | 2069 | 2,2 | 0819 |
| chr3L | 22628118 | 22629758 | 1641 | 2,23 | 0820 |
| chr3L | 22630057 | 22631855 | 1799 | 2,33 | 0821 |
| chr3L | 22636380 | 22644431 | 8052 | 2,56 | 0822 |
| chr3L | 22648495 | 22649789 | 1295 | 2,1 | 0823 |
| chr3L | 22649966 | 22651540 | 1575 | 2,04 | 0824 |
| chr3L | 22661761 | 22663122 | 1362 | 2,1 | 0825 |
| chr3L | 22667260 | 22668724 | 1465 | 2,26 | 0826 |
| chr3L | 22669446 | 22673392 | 3947 | 2,23 | 0827 |
| chr3L | 22677107 | 22678661 | 1555 | 2,43 | 0828 |
| chr3L | 22680124 | 22681602 | 1479 | 2,1 | 0829 |
| chr3L | 22681826 | 22683053 | 1228 | 1,84 | 0830 |
| chr3L | 22683218 | 22684419 | 1202 | 2,17 | 0831 |
| chr3L | 22692093 | 22693184 | 1092 | 2,2 | 0832 |
| chr3L | 22697296 | 22698333 | 1038 | 2,26 | 0833 |
| chr3L | 22698448 | 22699717 | 1270 | 2,14 | 0834 |
| chr3L | 22754340 | 22759950 | 5611 | 2,99 | 0835 |
| chr3L | 22797654 | 22804148 | 6495 | 2,09 | 0836 |
| chr3L | 22807740 | 22809538 | 1799 | 1,88 | 0837 |
| chr3L | 22975244 | 22979441 | 4198 | 2,17 | 0838 |
| chr3L | 23006778 | 23008875 | 2098 | 2,2 | 0839 |
| chr3L | 23010187 | 23011299 | 1113 | 2,1 | 0840 |
| chr3L | 23011376 | 23012742 | 1367 | 1,97 | 0841 |
| chr3L | 23018454 | 23019816 | 1363 | 1,91 | 0842 |
| chr3L | 23020541 | 23022147 | 1607 | 2,2 | 0843 |
| chr3L | 23022751 | 23026526 | 3776 | 2,36 | 0844 |
| chr3L | 23026542 | 23027697 | 1156 | 2,91 | 0845 |
| chr3L | 23031644 | 23034383 | 2740 | 2,36 | 0846 |
| chr3L | 23036377 | 23040119 | 3743 | 2,66 | 0847 |
| chr3L | 23040649 | 23041775 | 1127 | 2,07 | 0848 |
| chr3L | 23042032 | 23049378 | 7347 | 3,01 | 0849 |
| chr3L | 23049428 | 23054960 | 5533 | 2,36 | 0850 |
| chr3L | 23055428 | 23058694 | 3267 | 2,39 | 0851 |
| chr3L | 23059762 | 23061565 | 1804 | 2,43 | 0852 |
| chr3L | 23075597 | 23076755 | 1159 | 2,36 | 0853 |
| chr3L | 23077531 | 23079778 | 2248 | 2,46 | 0854 |
| chr3L | 23079973 | 23081414 | 1442 | 2,26 | 0855 |
| chr3L | 23097427 | 23099302 | 1876 | 2,36 | 0856 |
| chr3L | 23099327 | 23101311 | 1985 | 2,88 | 0857 |
| chr3L | 23101375 | 23104226 | 2852 | 2,39 | 0858 |
| chr3L | 23112161 | 23115853 | 3693 | 2,52 | 0859 |
| chr3L | 23135359 | 23138074 | 2716 | 2,43 | 0860 |
| chr3L | 23144943 | 23146800 | 1858 | 2,69 | 0861 |

| | | | | | |
|---|---|---|---|---|---|
| chr3L | 23169776 | 23172057 | 2282 | 2,43 | 0862 |
| chr3L | 23174875 | 23176199 | 1325 | 2,01 | 0863 |
| chr3L | 23180071 | 23182097 | 2027 | 2,23 | 0864 |
| chr3L | 23186057 | 23187331 | 1275 | 2,26 | 0865 |
| chr3L | 23197520 | 23199228 | 1709 | 2,01 | 0866 |
| chr3L | 23200573 | 23202568 | 1996 | 2,3 | 0867 |
| chr3L | 23205721 | 23208712 | 2992 | 2,56 | 0868 |
| chr3L | 23210796 | 23215698 | 4903 | 2,52 | 0869 |
| chr3L | 23217340 | 23218825 | 1486 | 2,07 | 0870 |
| chr3L | 23219138 | 23220822 | 1685 | 2,01 | 0871 |
| chr3L | 23220862 | 23223246 | 2385 | 2,43 | 0872 |
| chr3L | 23223415 | 23224632 | 1218 | 2,01 | 0873 |
| chr3L | 23226866 | 23228715 | 1850 | 2,23 | 0874 |
| chr3L | 23228832 | 23231701 | 2870 | 2,62 | 0875 |
| chr3L | 23235720 | 23237799 | 2080 | 2,46 | 0876 |
| chr3L | 23245519 | 23248546 | 3028 | 2,82 | 0877 |
| chr3L | 23249102 | 23250313 | 1212 | 2,17 | 0878 |
| chr3L | 23251080 | 23253926 | 2847 | 2,23 | 0879 |
| chr3L | 23257388 | 23259968 | 2581 | 2,56 | 0880 |
| chr3L | 23262496 | 23263761 | 1266 | 2,46 | 0881 |
| chr3L | 23271105 | 23272219 | 1115 | 2,23 | 0882 |
| chr3L | 23272380 | 23276450 | 4071 | 2,72 | 0883 |
| chr3L | 23280481 | 23283918 | 3438 | 2,49 | 0884 |
| chr3L | 23288994 | 23293900 | 4907 | 2,59 | 0885 |
| chr3L | 23297519 | 23311545 | 14027 | 3,56 | 0886 |
| chr3L | 23312840 | 23313940 | 1101 | 1,84 | 0887 |
| chr3L | 23315208 | 23316854 | 1647 | 2,14 | 0888 |
| chr3L | 23336340 | 23338725 | 2386 | 2,85 | 0889 |
| chr3L | 23338903 | 23341387 | 2485 | 3,3 | 0890 |
| chr3L | 23341814 | 23344082 | 2269 | 2,23 | 0891 |
| chr3L | 23344118 | 23347308 | 3191 | 2,59 | 0892 |
| chr3L | 23365871 | 23372567 | 6697 | 2,53 | 0893 |
| chr3L | 23373231 | 23378407 | 5177 | 2,62 | 0894 |
| chr3L | 23378762 | 23385239 | 6478 | 2,02 | 0895 |
| chr3L | 23387349 | 23389000 | 1652 | 2,43 | 0896 |
| chr3L | 23389160 | 23393729 | 4570 | 3,07 | 0897 |
| chr3L | 23393899 | 23394932 | 1034 | 2,78 | 0898 |
| chr3L | 23395240 | 23396316 | 1077 | 2,36 | 0899 |
| chr3L | 23396386 | 23397795 | 1410 | 2,39 | 0900 |
| chr3L | 23397986 | 23400086 | 2101 | 2,94 | 0901 |
| chr3L | 23409544 | 23415940 | 6397 | 3,43 | 0902 |
| chr3L | 23418751 | 23422173 | 3423 | 2,88 | 0903 |
| chr3L | 23423075 | 23424901 | 1827 | 2,2 | 0904 |
| chr3L | 23425483 | 23427205 | 1723 | 2,43 | 0905 |
| chr3L | 23427213 | 23428477 | 1265 | 2,43 | 0906 |
| chr3L | 23429219 | 23431047 | 1829 | 2,88 | 0907 |
| chr3L | 23438135 | 23441982 | 3848 | 3,01 | 0908 |
| chr3L | 23442001 | 23445100 | 3100 | 2,49 | 0909 |

| | | | | | |
|---|---|---|---|---|---|
| chr3L | 23445996 | 23449691 | 3696 | 2,49 | 0910 |
| chr3L | 23450068 | 23453486 | 3419 | 2,62 | 0911 |
| chr3L | 23458807 | 23464556 | 5750 | 2,89 | 0912 |
| chr3L | 23464874 | 23470123 | 5250 | 2,85 | 0913 |
| chr3L | 23470402 | 23480273 | 9872 | 2,62 | 0914 |
| chr3L | 23482184 | 23485162 | 2979 | 2,33 | 0915 |
| chr3L | 23491431 | 23492554 | 1124 | 2,07 | 0916 |
| chr3L | 23497078 | 23498392 | 1315 | 2,36 | 0917 |
| chr3L | 23498403 | 23505258 | 6856 | 2,69 | 0918 |
| chr3L | 23517750 | 23520019 | 2270 | 2,1 | 0919 |
| chr3L | 23521557 | 23525209 | 3653 | 2,3 | 0920 |
| chr3L | 23525515 | 23531409 | 5895 | 2,62 | 0921 |
| chr3L | 23531643 | 23534377 | 2735 | 2,43 | 0922 |
| chr3L | 23539948 | 23541406 | 1459 | 2,82 | 0923 |
| chr3L | 23541568 | 23543515 | 1948 | 2,56 | 0924 |
| chr3L | 23544278 | 23545470 | 1193 | 2,52 | 0925 |
| chr3L | 23549457 | 23550888 | 1432 | 2,59 | 0926 |
| chr3L | 23551886 | 23554516 | 2631 | 2,59 | 0927 |
| chr3L | 23554518 | 23556529 | 2012 | 2,36 | 0928 |
| chr3L | 23556657 | 23560156 | 3500 | 2,36 | 0929 |
| chr3L | 23563744 | 23565430 | 1687 | 2,43 | 0930 |
| chr3L | 23575737 | 23577231 | 1495 | 2,39 | 0931 |
| chr3L | 23580190 | 23583201 | 3012 | 2,49 | 0932 |
| chr3L | 23583441 | 23593088 | 9648 | 2,78 | 0933 |
| chr3L | 23593202 | 23595695 | 2494 | 2,62 | 0934 |
| chr3L | 23595899 | 23600352 | 4454 | 2,49 | 0935 |
| chr3L | 23600674 | 23605529 | 4856 | 3,14 | 0936 |
| chr3L | 23606005 | 23608279 | 2275 | 2,56 | 0937 |
| chr3L | 23608353 | 23610004 | 1652 | 2,07 | 0938 |
| chr3L | 23610100 | 23613927 | 3828 | 2,43 | 0939 |
| chr3L | 23618733 | 23623233 | 4501 | 2,98 | 0940 |
| chr3L | 23623522 | 23625137 | 1616 | 2,39 | 0941 |
| chr3L | 23625299 | 23627249 | 1951 | 2,56 | 0942 |
| chr3L | 23628236 | 23640032 | 11797 | 3,17 | 0943 |
| chr3L | 23641543 | 23643808 | 2266 | 2,49 | 0944 |
| chr3L | 23644203 | 23656434 | 12232 | 2,91 | 0945 |
| chr3L | 23691056 | 23692411 | 1356 | 2,39 | 0946 |
| chr3L | 23694010 | 23696013 | 2004 | 2,06 | 0947 |
| chr3L | 23696241 | 23697979 | 1739 | 2,3 | 0948 |
| chr3L | 23698541 | 23704306 | 5766 | 2,23 | 0949 |
| chr3L | 23712599 | 23719130 | 6532 | 3,18 | 0950 |
| chr3L | 23719482 | 23726415 | 6934 | 2,75 | 0951 |
| chr3L | 23727369 | 23728657 | 1289 | 2,62 | 0952 |
| chr3L | 23728696 | 23729894 | 1199 | 2,52 | 0953 |
| chr3L | 23746154 | 23747365 | 1212 | 2,04 | 0954 |
| chr3L | 23749243 | 23750407 | 1165 | 2,36 | 0955 |
| chr3L | 23758300 | 23759614 | 1315 | 2,1 | 0956 |
| chr3L | 23760457 | 23761848 | 1392 | 2,2 | 0957 |

| | | | | | |
|---|---|---|---|---|---|
| chr3L | 23765796 | 23767925 | 2130 | 2,01 | 0958 |
| chr3L | 23777668 | 23779189 | 1522 | 1,97 | 0959 |
| chr3L | 23780339 | 23783744 | 3406 | 3,04 | 0960 |
| chr3L | 23788933 | 23790697 | 1765 | 2,26 | 0961 |
| chr3L | 23791666 | 23793323 | 1658 | 2,97 | 0962 |
| chr3L | 23798565 | 23799915 | 1351 | 3,01 | 0963 |
| chr3L | 23801743 | 23803942 | 2200 | 2,69 | 0964 |
| chr3L | 23804191 | 23806208 | 2018 | 2,43 | 0965 |
| chr3L | 23806254 | 23808275 | 2022 | 3,37 | 0966 |
| chr3L | 23808481 | 23816872 | 8392 | 3,46 | 0967 |
| chr3L | 23835610 | 23836666 | 1057 | 2,72 | 0968 |
| chr3L | 23838300 | 23839397 | 1098 | 2,2 | 0969 |
| chr3L | 23839931 | 23842044 | 2114 | 2,3 | 0970 |
| chr3L | 23846912 | 23849007 | 2096 | 2,14 | 0971 |
| chr3L | 23849049 | 23850241 | 1193 | 2,23 | 0972 |
| chr3L | 23850467 | 23851574 | 1108 | 2,69 | 0973 |
| chr3L | 23860023 | 23861474 | 1452 | 2,33 | 0974 |
| chr3L | 23861504 | 23863072 | 1569 | 2,39 | 0975 |
| chr3L | 23863256 | 23864607 | 1352 | 2,01 | 0976 |
| chr3L | 23864794 | 23867395 | 2602 | 2,17 | 0977 |
| chr3L | 23867517 | 23873148 | 5632 | 2,65 | 0978 |
| chr3L | 23874821 | 23876065 | 1245 | 2,17 | 0979 |
| chr3L | 23876141 | 23877320 | 1180 | 2,75 | 0980 |
| chr3L | 23882035 | 23883759 | 1725 | 2,3 | 0981 |
| chr3L | 23884918 | 23886207 | 1290 | 2,14 | 0982 |
| chr3L | 23886463 | 23888757 | 2295 | 2,14 | 0983 |
| chr3L | 23889662 | 23891451 | 1790 | 2,04 | 0984 |
| chr3L | 23893886 | 23895344 | 1459 | 2,52 | 0985 |
| chr3L | 23899078 | 23901644 | 2567 | 2,3 | 0986 |
| chr3L | 23902099 | 23905556 | 3458 | 2,46 | 0987 |
| chr3L | 23907898 | 23911808 | 3911 | 3,11 | 0988 |
| chr3L | 23913022 | 23914763 | 1742 | 2,46 | 0989 |
| chr3L | 23923906 | 23927142 | 3237 | 3,51 | 0990 |
| chr3L | 23930307 | 23935359 | 5053 | 2,88 | 0991 |
| chr3L | 23935957 | 23940526 | 4570 | 2,09 | 0992 |
| chr3L | 23941427 | 23942903 | 1477 | 2,26 | 0993 |
| chr3L | 23946792 | 23948167 | 1376 | 2,14 | 0994 |
| chr3L | 23948252 | 23951813 | 3562 | 2,39 | 0995 |
| chr3L | 23953878 | 23958960 | 5083 | 2,69 | 0996 |
| chr3L | 23960740 | 23962593 | 1854 | 2,43 | 0997 |
| chr3L | 23963242 | 23968304 | 5063 | 2,69 | 0998 |
| chr3L | 23970030 | 23972968 | 2939 | 2,56 | 0999 |
| chr3L | 23973669 | 23974956 | 1288 | 3,14 | 1000 |
| chr3L | 23982256 | 23983259 | 1004 | 2,56 | 1001 |
| chr3L | 23989459 | 23991109 | 1651 | 2,23 | 1002 |
| chr3L | 23992462 | 23993697 | 1236 | 2,26 | 1003 |
| chr3L | 23994167 | 23999944 | 5778 | 3,14 | 1004 |
| chr3L | 24002202 | 24005120 | 2919 | 2,56 | 1005 |

| | | | | | |
|---|---|---|---|---|---|
| chr3L | 24018844 | 24020195 | 1352 | 2,14 | 1006 |
| chr3L | 24030604 | 24032455 | 1852 | 2,07 | 1007 |
| chr3L | 24032607 | 24034311 | 1705 | 2,33 | 1008 |
| chr3L | 24034754 | 24038272 | 3519 | 2,46 | 1009 |
| chr3L | 24039565 | 24048533 | 8969 | 2,62 | 1010 |
| chr3L | 24051951 | 24055520 | 3570 | 2,82 | 1011 |
| chr3L | 24056154 | 24063761 | 7608 | 3,01 | 1012 |
| chr3L | 24064143 | 24065238 | 1096 | 2,14 | 1013 |
| chr3L | 24068431 | 24078291 | 9861 | 2,72 | 1014 |
| chr3L | 24081784 | 24083671 | 1888 | 2,49 | 1015 |
| chr3L | 24102726 | 24107079 | 4354 | 2,62 | 1016 |
| chr3L | 24112227 | 24123759 | 11533 | 3,57 | 1017 |
| chr3L | 24138441 | 24140337 | 1897 | 1,81 | 1018 |
| chr3L | 24140416 | 24147999 | 7584 | 2,65 | 1019 |
| chr3L | 24148121 | 24151814 | 3694 | 2,62 | 1020 |
| chr3L | 24156180 | 24158251 | 2072 | 2,36 | 1021 |
| chr3L | 24160572 | 24162089 | 1518 | 2,46 | 1022 |
| chr3L | 24169061 | 24175463 | 6403 | 3,14 | 1023 |
| chr3L | 24178584 | 24180505 | 1922 | 2,2 | 1024 |
| chr3L | 24180603 | 24184656 | 4054 | 2,39 | 1025 |
| chr3L | 24184841 | 24187240 | 2400 | 2,3 | 1026 |
| chr3L | 24193856 | 24195289 | 1434 | 2,2 | 1027 |
| chr3L | 24197330 | 24200071 | 2742 | 2,82 | 1028 |
| chr3L | 24201285 | 24202472 | 1188 | 2,26 | 1029 |
| chr3L | 24204428 | 24206311 | 1884 | 2,59 | 1030 |
| chr3L | 24229056 | 24231106 | 2051 | 2,23 | 1031 |
| chr3L | 24244382 | 24245833 | 1452 | 2,39 | 1032 |
| chr3L | 24246067 | 24249289 | 3223 | 2,43 | 1033 |
| chr3L | 24249466 | 24250546 | 1081 | 2,23 | 1034 |
| chr3L | 24251543 | 24254384 | 2842 | 2,52 | 1035 |
| chr3L | 24254971 | 24259215 | 4245 | 2,69 | 1036 |
| chr3L | 24260053 | 24265423 | 5371 | 2,56 | 1037 |
| chr3L | 24265853 | 24271502 | 5650 | 2,33 | 1038 |
| chr3L | 24274939 | 24276152 | 1214 | 2,01 | 1039 |
| chr3L | 24281046 | 24284199 | 3154 | 2,36 | 1040 |
| chr3L | 24284258 | 24285357 | 1100 | 2,56 | 1041 |
| chr3L | 24286367 | 24291393 | 5027 | 2,33 | 1042 |
| chr3L | 24291558 | 24294045 | 2488 | 2,49 | 1043 |
| chr3L | 24295498 | 24297418 | 1921 | 2,75 | 1044 |
| chr3L | 24298550 | 24300838 | 2289 | 2,52 | 1045 |
| chr3L | 24301442 | 24302670 | 1229 | 2,04 | 1046 |
| chr3L | 24302694 | 24304719 | 2026 | 2,46 | 1047 |
| chr3L | 24306577 | 24309882 | 3306 | 2,36 | 1048 |
| chr3L | 24312536 | 24317649 | 5114 | 2,39 | 1049 |
| chr3L | 24320475 | 24321607 | 1133 | 2,26 | 1050 |
| chr3L | 24324331 | 24327208 | 2878 | 2,33 | 1051 |
| chr3L | 24328583 | 24331355 | 2773 | 2,49 | 1052 |
| chr3L | 24339849 | 24340922 | 1074 | 2,07 | 1053 |

| | | | | | |
|---|---|---|---|---|---|
| chr3L | 24342081 | 24346374 | 4294 | 2,62 | 1054 |
| chr3L | 24353140 | 24355195 | 2056 | 2,36 | 1055 |
| chr3L | 24355605 | 24357883 | 2279 | 2,14 | 1056 |
| chr3L | 24360128 | 24362282 | 2155 | 2,56 | 1057 |
| chr3L | 24366025 | 24369811 | 3787 | 2,36 | 1058 |
| chr3L | 24376644 | 24379267 | 2624 | 2,46 | 1059 |
| chr3L | 24379300 | 24380489 | 1190 | 2,24 | 1060 |
| chr3L | 24390095 | 24391669 | 1575 | 2,39 | 1061 |
| chr3L | 24394014 | 24396764 | 2751 | 2,17 | 1062 |
| chr3L | 24401655 | 24403565 | 1911 | 2,2 | 1063 |
| chr3L | 24409906 | 24411419 | 1514 | 2,49 | 1064 |
| chr3L | 24423264 | 24424925 | 1662 | 3,02 | 1065 |
| chr3L | 24425391 | 24427407 | 2017 | 2,59 | 1066 |
| chr3L | 24431963 | 24436932 | 4970 | 1,89 | 1067 |
| chr3L | 24439375 | 24440801 | 1427 | 2,1 | 1068 |
| chr3L | 24473041 | 24475705 | 2665 | 2,14 | 1069 |
| chr3L | 24482761 | 24486006 | 3246 | 2,94 | 1070 |
| chr3L | 24494883 | 24496249 | 1367 | 2,26 | 1071 |
| chr3L | 24498589 | 24502932 | 4344 | 3,25 | 1072 |
| chr3L | 24503580 | 24505279 | 1700 | 2,43 | 1073 |
| chr3L | 24507220 | 24513244 | 6025 | 2,62 | 1074 |
| chr3L | 24514501 | 24517358 | 2858 | 2,33 | 1075 |
| chr3LHet | 10629 | 11773 | 1145 | 2,3 | 1076 |
| chr3LHet | 12362 | 14601 | 2240 | 2,91 | 1077 |
| chr3LHet | 15571 | 17719 | 2149 | 2,14 | 1078 |
| chr3LHet | 17968 | 19496 | 1529 | 2,52 | 1079 |
| chr3LHet | 19926 | 21532 | 1607 | 2,36 | 1080 |
| chr3LHet | 87002 | 89050 | 2049 | 2,46 | 1081 |
| chr3LHet | 160686 | 162810 | 2125 | 1,84 | 1082 |
| chr3LHet | 163049 | 164919 | 1871 | 2,31 | 1083 |
| chr3LHet | 177172 | 183342 | 6171 | 3,04 | 1084 |
| chr3LHet | 183630 | 191103 | 7474 | 3,07 | 1085 |
| chr3LHet | 191536 | 201989 | 10454 | 3 | 1086 |
| chr3LHet | 211272 | 214083 | 2812 | 2,91 | 1087 |
| chr3LHet | 214307 | 216989 | 2683 | 2,43 | 1088 |
| chr3LHet | 217002 | 218855 | 1854 | 2,33 | 1089 |
| chr3LHet | 220337 | 223173 | 2837 | 2,37 | 1090 |
| chr3LHet | 241275 | 243444 | 2170 | 2,52 | 1091 |
| chr3LHet | 253434 | 256533 | 3100 | 2,59 | 1092 |
| chr3LHet | 257106 | 260389 | 3284 | 2,46 | 1093 |
| chr3LHet | 268399 | 270535 | 2137 | 2,1 | 1094 |
| chr3LHet | 271608 | 275348 | 3741 | 2,62 | 1095 |
| chr3LHet | 279948 | 286043 | 6096 | 2,8 | 1096 |
| chr3LHet | 291272 | 292362 | 1091 | 2,3 | 1097 |
| chr3LHet | 293004 | 297535 | 4532 | 3,39 | 1098 |
| chr3LHet | 297626 | 303779 | 6154 | 2,16 | 1099 |
| chr3LHet | 319099 | 327719 | 8621 | 2,85 | 1100 |
| chr3LHet | 328264 | 329886 | 1623 | 2,62 | 1101 |

| | | | | | |
|---|---|---|---|---|---|
| chr3LHet | 332860 | 334796 | 1937 | 3,33 | 1102 |
| chr3LHet | 336960 | 342080 | 5121 | 2,73 | 1103 |
| chr3LHet | 342589 | 346844 | 4256 | 2,72 | 1104 |
| chr3LHet | 349231 | 355517 | 6287 | 2,52 | 1105 |
| chr3LHet | 355760 | 357321 | 1562 | 2,65 | 1106 |
| chr3LHet | 358007 | 364798 | 6792 | 2,82 | 1107 |
| chr3LHet | 368953 | 370539 | 1587 | 2,2 | 1108 |
| chr3LHet | 392745 | 398819 | 6075 | 3,39 | 1109 |
| chr3LHet | 399001 | 400756 | 1756 | 3,42 | 1110 |
| chr3LHet | 404019 | 410633 | 6615 | 2,92 | 1111 |
| chr3LHet | 412398 | 425114 | 12717 | 3,4 | 1112 |
| chr3LHet | 426028 | 434079 | 8052 | 4,26 | 1113 |
| chr3LHet | 434999 | 438403 | 3405 | 3,01 | 1114 |
| chr3LHet | 455506 | 458474 | 2969 | 2,73 | 1115 |
| chr3LHet | 461416 | 464593 | 3178 | 2,39 | 1116 |
| chr3LHet | 464817 | 469460 | 4644 | 2,69 | 1117 |
| chr3LHet | 471048 | 472716 | 1669 | 2,39 | 1118 |
| chr3LHet | 474730 | 479615 | 4886 | 2,69 | 1119 |
| chr3LHet | 479789 | 497857 | 18069 | 3,85 | 1120 |
| chr3LHet | 501830 | 503734 | 1905 | 2,17 | 1121 |
| chr3LHet | 505149 | 508314 | 3166 | 3,13 | 1122 |
| chr3LHet | 511117 | 512844 | 1728 | 2,33 | 1123 |
| chr3LHet | 513446 | 514505 | 1060 | 2,69 | 1124 |
| chr3LHet | 532351 | 535403 | 3053 | 2,75 | 1125 |
| chr3LHet | 561587 | 562747 | 1161 | 2,36 | 1126 |
| chr3LHet | 573621 | 576297 | 2677 | 2,21 | 1127 |
| chr3LHet | 577894 | 586592 | 8699 | 2,85 | 1128 |
| chr3LHet | 587002 | 588130 | 1129 | 2,39 | 1129 |
| chr3LHet | 588822 | 592609 | 3788 | 2,59 | 1130 |
| chr3LHet | 601928 | 605450 | 3523 | 2,05 | 1131 |
| chr3LHet | 607821 | 615441 | 7621 | 3,27 | 1132 |
| chr3LHet | 628310 | 629726 | 1417 | 2,65 | 1133 |
| chr3LHet | 629894 | 641882 | 11989 | 3,4 | 1134 |
| chr3LHet | 645501 | 648689 | 3189 | 2,78 | 1135 |
| chr3LHet | 649982 | 652569 | 2588 | 2,39 | 1136 |
| chr3LHet | 676241 | 678096 | 1856 | 2,46 | 1137 |
| chr3LHet | 683894 | 686587 | 2694 | 2,45 | 1138 |
| chr3LHet | 704264 | 709802 | 5539 | 2,65 | 1139 |
| chr3LHet | 715088 | 716227 | 1140 | 2,75 | 1140 |
| chr3LHet | 716662 | 724750 | 8089 | 2,98 | 1141 |
| chr3LHet | 725044 | 731223 | 6180 | 3,04 | 1142 |
| chr3LHet | 740013 | 741138 | 1126 | 2,88 | 1143 |
| chr3LHet | 762508 | 772562 | 10055 | 2,98 | 1144 |
| chr3LHet | 772766 | 774015 | 1250 | 2,65 | 1145 |
| chr3LHet | 778744 | 780830 | 2087 | 2,2 | 1146 |
| chr3LHet | 787510 | 795661 | 8152 | 3,07 | 1147 |
| chr3LHet | 795668 | 798234 | 2567 | 3,11 | 1148 |
| chr3LHet | 799768 | 801362 | 1595 | 2,07 | 1149 |

| | | | | | |
|---|---|---|---|---|---|
| chr3LHet | 801414 | 807041 | 5628 | 3,17 | 1150 |
| chr3LHet | 807212 | 809446 | 2235 | 2,91 | 1151 |
| chr3LHet | 809582 | 812395 | 2814 | 2,42 | 1152 |
| chr3LHet | 812465 | 813621 | 1157 | 2,17 | 1153 |
| chr3LHet | 813765 | 823341 | 9577 | 3,4 | 1154 |
| chr3LHet | 823403 | 837156 | 13754 | 3,4 | 1155 |
| chr3LHet | 837183 | 844652 | 7470 | 3,13 | 1156 |
| chr3LHet | 866966 | 877445 | 10480 | 3,01 | 1157 |
| chr3LHet | 882364 | 895600 | 13237 | 3,07 | 1158 |
| chr3LHet | 905124 | 906832 | 1709 | 2,82 | 1159 |
| chr3LHet | 934035 | 935540 | 1506 | 2,59 | 1160 |
| chr3LHet | 935949 | 938686 | 2738 | 2,82 | 1161 |
| chr3LHet | 939208 | 946783 | 7576 | 2,88 | 1162 |
| chr3LHet | 946812 | 949877 | 3066 | 2,59 | 1163 |
| chr3LHet | 950248 | 951871 | 1624 | 2,62 | 1164 |
| chr3LHet | 954717 | 955882 | 1166 | 2,26 | 1165 |
| chr3LHet | 963155 | 964215 | 1061 | 2,23 | 1166 |
| chr3LHet | 972263 | 974899 | 2637 | 2,36 | 1167 |
| chr3LHet | 980310 | 984542 | 4233 | 3,88 | 1168 |
| chr3LHet | 984656 | 985893 | 1238 | 2,36 | 1169 |
| chr3LHet | 986112 | 988983 | 2872 | 3,37 | 1170 |
| chr3LHet | 989009 | 993052 | 4044 | 3,43 | 1171 |
| chr3LHet | 993275 | 997079 | 3805 | 2,94 | 1172 |
| chr3LHet | 999947 | 1011412 | 11466 | 3,75 | 1173 |
| chr3LHet | 1015769 | 1017114 | 1346 | 2,39 | 1174 |
| chr3LHet | 1017153 | 1019343 | 2191 | 3,17 | 1175 |
| chr3LHet | 1019366 | 1021487 | 2122 | 2,59 | 1176 |
| chr3LHet | 1029587 | 1030717 | 1131 | 2,26 | 1177 |
| chr3LHet | 1032380 | 1034125 | 1746 | 2,69 | 1178 |
| chr3LHet | 1034176 | 1038020 | 3845 | 3,26 | 1179 |
| chr3LHet | 1044569 | 1053061 | 8493 | 3,79 | 1180 |
| chr3LHet | 1117495 | 1118650 | 1156 | 2,46 | 1181 |
| chr3LHet | 1129965 | 1144183 | 14219 | 3,4 | 1182 |
| chr3LHet | 1144258 | 1146125 | 1868 | 2,9 | 1183 |
| chr3LHet | 1146357 | 1157365 | 11009 | 3,01 | 1184 |
| chr3LHet | 1165716 | 1167751 | 2036 | 2,39 | 1185 |
| chr3LHet | 1172570 | 1178951 | 6382 | 3,04 | 1186 |
| chr3LHet | 1179083 | 1184395 | 5313 | 2,72 | 1187 |
| chr3LHet | 1184524 | 1185809 | 1286 | 2,72 | 1188 |
| chr3LHet | 1188048 | 1192766 | 4719 | 2,82 | 1189 |
| chr3LHet | 1192881 | 1197781 | 4901 | 3,04 | 1190 |
| chr3LHet | 1197836 | 1212770 | 14935 | 2,91 | 1191 |
| chr3LHet | 1213237 | 1215098 | 1862 | 2,07 | 1192 |
| chr3LHet | 1219616 | 1223820 | 4205 | 3,07 | 1193 |
| chr3LHet | 1223863 | 1227365 | 3503 | 3,46 | 1194 |
| chr3LHet | 1227524 | 1231273 | 3750 | 2,85 | 1195 |
| chr3LHet | 1232086 | 1233892 | 1807 | 2,91 | 1196 |
| chr3LHet | 1233961 | 1237910 | 3950 | 3,2 | 1197 |

| | | | | | |
|---|---|---|---|---|---|
| chr3LHet | 1241126 | 1242167 | 1042 | 2,07 | 1198 |
| chr3LHet | 1249118 | 1250704 | 1587 | 2,07 | 1199 |
| chr3LHet | 1250970 | 1257321 | 6352 | 3,01 | 1200 |
| chr3LHet | 1258476 | 1262504 | 4029 | 3,43 | 1201 |
| chr3LHet | 1262694 | 1264506 | 1813 | 2,36 | 1202 |
| chr3LHet | 1266359 | 1267740 | 1382 | 2,52 | 1203 |
| chr3LHet | 1269855 | 1272692 | 2838 | 2,3 | 1204 |
| chr3LHet | 1272979 | 1275994 | 3016 | 2,59 | 1205 |
| chr3LHet | 1289986 | 1291115 | 1130 | 2,07 | 1206 |
| chr3LHet | 1291411 | 1297574 | 6164 | 2,72 | 1207 |
| chr3LHet | 1310199 | 1311390 | 1192 | 1,88 | 1208 |
| chr3LHet | 1314709 | 1315788 | 1080 | 3,11 | 1209 |
| chr3LHet | 1320211 | 1321279 | 1069 | 2,23 | 1210 |
| chr3LHet | 1323491 | 1324743 | 1253 | 2,2 | 1211 |
| chr3LHet | 1332775 | 1342286 | 9512 | 2,88 | 1212 |
| chr3LHet | 1342341 | 1343719 | 1379 | 2,04 | 1213 |
| chr3LHet | 1344722 | 1348201 | 3480 | 3,04 | 1214 |
| chr3LHet | 1348241 | 1350755 | 2515 | 2,33 | 1215 |
| chr3LHet | 1353014 | 1354482 | 1469 | 2,04 | 1216 |
| chr3LHet | 1354752 | 1360522 | 5771 | 2,94 | 1217 |
| chr3LHet | 1361372 | 1363713 | 2342 | 2,3 | 1218 |
| chr3LHet | 1364084 | 1365436 | 1353 | 2,43 | 1219 |
| chr3LHet | 1366041 | 1372506 | 6466 | 3,04 | 1220 |
| chr3LHet | 1378310 | 1381346 | 3037 | 2,46 | 1221 |
| chr3LHet | 1382552 | 1384151 | 1600 | 2,17 | 1222 |
| chr3LHet | 1385945 | 1387643 | 1699 | 2,14 | 1223 |
| chr3LHet | 1387857 | 1389333 | 1477 | 2,65 | 1224 |
| chr3LHet | 1412122 | 1416120 | 3999 | 2,75 | 1225 |
| chr3LHet | 1424735 | 1427755 | 3021 | 3,2 | 1226 |
| chr3LHet | 1438039 | 1444536 | 6498 | 3,11 | 1227 |
| chr3LHet | 1446328 | 1448384 | 2057 | 3,07 | 1228 |
| chr3LHet | 1449353 | 1456307 | 6955 | 2,91 | 1229 |
| chr3LHet | 1482221 | 1484663 | 2443 | 2,88 | 1230 |
| chr3LHet | 1485813 | 1499551 | 13739 | 3,33 | 1231 |
| chr3LHet | 1499997 | 1504909 | 4913 | 2,65 | 1232 |
| chr3LHet | 1518416 | 1520405 | 1990 | 2,78 | 1233 |
| chr3LHet | 1520911 | 1522969 | 2059 | 2,94 | 1234 |
| chr3LHet | 1526486 | 1529931 | 3446 | 2,82 | 1235 |
| chr3LHet | 1530444 | 1532018 | 1575 | 2,07 | 1236 |
| chr3LHet | 1533245 | 1535862 | 2618 | 2,75 | 1237 |
| chr3LHet | 1535900 | 1537978 | 2079 | 2,85 | 1238 |
| chr3LHet | 1545845 | 1551204 | 5360 | 2,89 | 1239 |
| chr3LHet | 1551441 | 1553907 | 2467 | 3,45 | 1240 |
| chr3LHet | 1554469 | 1556168 | 1700 | 2,6 | 1241 |
| chr3LHet | 1572702 | 1581798 | 9097 | 2,95 | 1242 |
| chr3LHet | 1590870 | 1592727 | 1858 | 2,62 | 1243 |
| chr3LHet | 1593210 | 1595563 | 2354 | 2,62 | 1244 |
| chr3LHet | 1605973 | 1608728 | 2756 | 2,46 | 1245 |

| | | | | | |
|---|---|---|---|---|---|
| chr3LHet | 1609847 | 1620863 | 11017 | 3,1 | 1246 |
| chr3LHet | 1622506 | 1623714 | 1209 | 2,72 | 1247 |
| chr3LHet | 1624121 | 1629727 | 5607 | 3,01 | 1248 |
| chr3LHet | 1629927 | 1634612 | 4686 | 3,3 | 1249 |
| chr3LHet | 1634852 | 1635968 | 1117 | 2,62 | 1250 |
| chr3LHet | 1636116 | 1639342 | 3227 | 2,88 | 1251 |
| chr3LHet | 1639403 | 1642820 | 3418 | 2,88 | 1252 |
| chr3LHet | 1642860 | 1645031 | 2172 | 2,52 | 1253 |
| chr3LHet | 1647066 | 1652504 | 5439 | 2,78 | 1254 |
| chr3LHet | 1680687 | 1681928 | 1242 | 2,23 | 1255 |
| chr3LHet | 1704296 | 1706276 | 1981 | 3,06 | 1256 |
| chr3LHet | 1708867 | 1711408 | 2542 | 2,52 | 1257 |
| chr3LHet | 1713437 | 1714592 | 1156 | 1,97 | 1258 |
| chr3LHet | 1735196 | 1736304 | 1109 | 2,96 | 1259 |
| chr3LHet | 1739584 | 1741804 | 2221 | 2,49 | 1260 |
| chr3LHet | 1741814 | 1743701 | 1888 | 3,14 | 1261 |
| chr3LHet | 1745902 | 1747640 | 1739 | 2,56 | 1262 |
| chr3LHet | 1750446 | 1755709 | 5264 | 3,01 | 1263 |
| chr3LHet | 1756879 | 1759241 | 2363 | 2,72 | 1264 |
| chr3LHet | 1760173 | 1762881 | 2709 | 2,23 | 1265 |
| chr3LHet | 1768482 | 1769760 | 1279 | 3,04 | 1266 |
| chr3LHet | 1772713 | 1774017 | 1305 | 2,46 | 1267 |
| chr3LHet | 1796059 | 1798740 | 2682 | 2,65 | 1268 |
| chr3LHet | 1801267 | 1802830 | 1564 | 2,33 | 1269 |
| chr3LHet | 1817546 | 1819167 | 1622 | 2,23 | 1270 |
| chr3LHet | 1819700 | 1822041 | 2342 | 2,72 | 1271 |
| chr3LHet | 1822076 | 1826534 | 4459 | 2,62 | 1272 |
| chr3LHet | 1826913 | 1828402 | 1490 | 2,69 | 1273 |
| chr3LHet | 1828853 | 1833048 | 4196 | 2,31 | 1274 |
| chr3LHet | 1841469 | 1844641 | 3173 | 2,43 | 1275 |
| chr3LHet | 1848061 | 1849318 | 1258 | 2,33 | 1276 |
| chr3LHet | 1850286 | 1852861 | 2576 | 3,07 | 1277 |
| chr3LHet | 1854495 | 1857691 | 3197 | 2,79 | 1278 |
| chr3LHet | 1859462 | 1862465 | 3004 | 2,74 | 1279 |
| chr3LHet | 1863512 | 1864572 | 1061 | 1,91 | 1280 |
| chr3LHet | 1866222 | 1868241 | 2020 | 2,17 | 1281 |
| chr3LHet | 1868300 | 1871283 | 2984 | 2,49 | 1282 |
| chr3LHet | 1871805 | 1872918 | 1114 | 2,43 | 1283 |
| chr3LHet | 1873076 | 1874255 | 1180 | 2,04 | 1284 |
| chr3LHet | 1874661 | 1877469 | 2809 | 2,33 | 1285 |
| chr3LHet | 1879509 | 1880890 | 1382 | 2,17 | 1286 |
| chr3LHet | 1882596 | 1883899 | 1304 | 2,36 | 1287 |
| chr3LHet | 1888858 | 1894273 | 5416 | 2,49 | 1288 |
| chr3LHet | 1895151 | 1899583 | 4433 | 3,24 | 1289 |
| chr3LHet | 1902465 | 1905800 | 3336 | 3,07 | 1290 |
| chr3LHet | 1924082 | 1925147 | 1066 | 2,2 | 1291 |
| chr3LHet | 1925732 | 1927567 | 1836 | 2,52 | 1292 |
| chr3LHet | 1927817 | 1930006 | 2190 | 1,88 | 1293 |

| | | | | | |
|---|---|---|---|---|---|
| chr3LHet | 1937742 | 1940745 | 3004 | 2,2 | 1294 |
| chr3LHet | 1941160 | 1942673 | 1514 | 2,52 | 1295 |
| chr3LHet | 1943039 | 1944338 | 1300 | 1,97 | 1296 |
| chr3LHet | 1944485 | 1947328 | 2844 | 2,62 | 1297 |
| chr3LHet | 1947366 | 1948871 | 1506 | 2,2 | 1298 |
| chr3LHet | 1949912 | 1960050 | 10139 | 3,07 | 1299 |
| chr3LHet | 1964410 | 1967124 | 2715 | 2,32 | 1300 |
| chr3LHet | 1967492 | 1977929 | 10438 | 3,71 | 1301 |
| chr3LHet | 1978742 | 1981803 | 3062 | 2,46 | 1302 |
| chr3LHet | 1981986 | 1984681 | 2696 | 2,3 | 1303 |
| chr3LHet | 1984694 | 1991837 | 7144 | 3,03 | 1304 |
| chr3LHet | 1992369 | 1994032 | 1664 | 2,1 | 1305 |
| chr3LHet | 1996660 | 2001606 | 4947 | 2,94 | 1306 |
| chr3LHet | 2002048 | 2004410 | 2363 | 2,94 | 1307 |
| chr3LHet | 2004557 | 2011481 | 6925 | 2,75 | 1308 |
| chr3LHet | 2014893 | 2016296 | 1404 | 2,59 | 1309 |
| chr3LHet | 2019509 | 2023524 | 4016 | 2,94 | 1310 |
| chr3LHet | 2024020 | 2027206 | 3187 | 2,3 | 1311 |
| chr3LHet | 2027743 | 2030653 | 2911 | 2,82 | 1312 |
| chr3LHet | 2033324 | 2037083 | 3760 | 2,62 | 1313 |
| chr3LHet | 2037115 | 2038188 | 1074 | 2,91 | 1314 |
| chr3LHet | 2038477 | 2045338 | 6862 | 3,29 | 1315 |
| chr3LHet | 2045648 | 2047625 | 1978 | 2,43 | 1316 |
| chr3LHet | 2054037 | 2055285 | 1249 | 2,1 | 1317 |
| chr3LHet | 2063748 | 2065946 | 2199 | 2,82 | 1318 |
| chr3LHet | 2073583 | 2075015 | 1433 | 2,85 | 1319 |
| chr3LHet | 2082246 | 2087776 | 5531 | 2,46 | 1320 |
| chr3LHet | 2088543 | 2090570 | 2028 | 2,26 | 1321 |
| chr3LHet | 2092093 | 2093262 | 1170 | 2,75 | 1322 |
| chr3LHet | 2099947 | 2102490 | 2544 | 2,52 | 1323 |
| chr3LHet | 2108725 | 2110948 | 2224 | 2,17 | 1324 |
| chr3LHet | 2111590 | 2118195 | 6606 | 2,52 | 1325 |
| chr3LHet | 2126353 | 2131284 | 4932 | 2,88 | 1326 |
| chr3LHet | 2131385 | 2134690 | 3306 | 2,46 | 1327 |
| chr3LHet | 2135860 | 2137078 | 1219 | 2,17 | 1328 |
| chr3LHet | 2146887 | 2153093 | 6207 | 2,39 | 1329 |
| chr3LHet | 2153179 | 2154561 | 1383 | 2,1 | 1330 |
| chr3LHet | 2154802 | 2157055 | 2254 | 2,23 | 1331 |
| chr3LHet | 2158773 | 2160358 | 1586 | 2,3 | 1332 |
| chr3LHet | 2166316 | 2167839 | 1524 | 1,79 | 1333 |
| chr3LHet | 2180329 | 2182426 | 2098 | 2,19 | 1334 |
| chr3LHet | 2182496 | 2185703 | 3208 | 2,36 | 1335 |
| chr3LHet | 2186713 | 2188141 | 1429 | 2,36 | 1336 |
| chr3LHet | 2194757 | 2197325 | 2569 | 2,46 | 1337 |
| chr3LHet | 2198653 | 2201978 | 3326 | 2,17 | 1338 |
| chr3LHet | 2203494 | 2212062 | 8569 | 2,75 | 1339 |
| chr3LHet | 2213248 | 2215096 | 1849 | 2,1 | 1340 |
| chr3LHet | 2218902 | 2224489 | 5588 | 2,75 | 1341 |

| | | | | | |
|---|---|---|---|---|---|
| chr3LHet | 2239666 | 2240960 | 1295 | 2,17 | 1342 |
| chr3LHet | 2265626 | 2273890 | 8265 | 2,78 | 1343 |
| chr3LHet | 2274300 | 2276918 | 2619 | 2,94 | 1344 |
| chr3LHet | 2279319 | 2281217 | 1899 | 2,6 | 1345 |
| chr3LHet | 2285570 | 2287776 | 2207 | 2,85 | 1346 |
| chr3LHet | 2383415 | 2385498 | 2084 | 2,82 | 1347 |
| chr3LHet | 2385652 | 2389296 | 3645 | 2,39 | 1348 |
| chr3LHet | 2389769 | 2391706 | 1938 | 2,26 | 1349 |
| chr3LHet | 2401152 | 2403668 | 2517 | 2,33 | 1350 |
| chr3LHet | 2405622 | 2407640 | 2019 | 2,22 | 1351 |
| chr3LHet | 2426241 | 2436041 | 9801 | 2,91 | 1352 |
| chr3LHet | 2436055 | 2443435 | 7381 | 3,33 | 1353 |
| chr3LHet | 2451506 | 2454108 | 2603 | 2,17 | 1354 |
| chr3LHet | 2458145 | 2462810 | 4666 | 3,11 | 1355 |
| chr3LHet | 2463045 | 2466238 | 3194 | 2,46 | 1356 |
| chr3LHet | 2469364 | 2473199 | 3836 | 2,85 | 1357 |
| chr3LHet | 2473362 | 2479253 | 5892 | 2,85 | 1358 |
| chr3LHet | 2480216 | 2486836 | 6621 | 2,59 | 1359 |
| chr3LHet | 2488099 | 2491927 | 3829 | 2,36 | 1360 |
| chr3LHet | 2496611 | 2497819 | 1209 | 2,04 | 1361 |
| chr3LHet | 2501360 | 2504035 | 2676 | 2,26 | 1362 |
| chr3LHet | 2504297 | 2506938 | 2642 | 2,85 | 1363 |
| chr3LHet | 2506982 | 2510550 | 3569 | 2,49 | 1364 |
| chr3LHet | 2512884 | 2515681 | 2798 | 2,49 | 1365 |
| chr3LHet | 2518634 | 2522860 | 4227 | 2,46 | 1366 |
| chr3LHet | 2542181 | 2546009 | 3829 | 2,72 | 1367 |
| chr3LHet | 2546275 | 2550812 | 4538 | 2,88 | 1368 |
| chr3R | 582024 | 583096 | 1073 | 2,07 | 1369 |
| chr3R | 752675 | 757412 | 4738 | 2,56 | 1370 |
| chr3R | 872893 | 878379 | 5487 | 2,66 | 1371 |
| chr3R | 1164357 | 1165802 | 1446 | 2,46 | 1372 |
| chr3R | 1352895 | 1354043 | 1149 | 2,43 | 1373 |
| chr3R | 3177609 | 3179063 | 1455 | 2,01 | 1374 |
| chr3R | 3924025 | 3925142 | 1118 | 2,69 | 1375 |
| chr3R | 4396110 | 4399571 | 3462 | 3,25 | 1376 |
| chr3R | 5839112 | 5840177 | 1066 | 2,23 | 1377 |
| chr3R | 5842914 | 5844316 | 1403 | 2,2 | 1378 |
| chr3R | 5845450 | 5847307 | 1858 | 2,2 | 1379 |
| chr3R | 6529034 | 6532245 | 3212 | 2,46 | 1380 |
| chr3R | 7720544 | 7722697 | 2154 | 2,01 | 1381 |
| chr3R | 9804903 | 9806200 | 1298 | 2,17 | 1382 |
| chr3R | 12151355 | 12153016 | 1662 | 2,01 | 1383 |
| chr3R | 13859236 | 13865894 | 6659 | 2,39 | 1384 |
| chr3R | 14962431 | 14964777 | 2347 | 2,66 | 1385 |
| chr3R | 16339980 | 16341155 | 1176 | 2,36 | 1386 |
| chr3R | 19383795 | 19386361 | 2567 | 2,46 | 1387 |
| chr3R | 19677121 | 19683522 | 6402 | 2,2 | 1388 |
| chr3R | 20034011 | 20036662 | 2652 | 2,28 | 1389 |

| | | | | | |
|---|---|---|---|---|---|
| chr3R | 23133792 | 23134809 | 1018 | 2,1 | 1390 |
| chr3R | 23136585 | 23139403 | 2819 | 2,26 | 1391 |
| chr3R | 23142968 | 23146008 | 3041 | 2,3 | 1392 |
| chr3R | 23149629 | 23152077 | 2449 | 2,07 | 1393 |
| chr3R | 23161111 | 23162819 | 1709 | 2,23 | 1394 |
| chr3R | 23186513 | 23187919 | 1407 | 2,01 | 1395 |
| chr3R | 23205531 | 23207550 | 2020 | 2,43 | 1396 |
| chr3R | 23211369 | 23212779 | 1411 | 2,2 | 1397 |
| chr3R | 23287040 | 23288146 | 1107 | 2,04 | 1398 |
| chr3R | 23288368 | 23289568 | 1201 | 1,91 | 1399 |
| chr3R | 23311470 | 23313476 | 2007 | 2,3 | 1400 |
| chr3R | 23313629 | 23315301 | 1673 | 1,97 | 1401 |
| chr3R | 23331807 | 23333828 | 2022 | 1,97 | 1402 |
| chr3R | 23335089 | 23336556 | 1468 | 2,49 | 1403 |
| chr3R | 23366921 | 23368805 | 1885 | 2,07 | 1404 |
| chr3R | 23608148 | 23609517 | 1370 | 2,23 | 1405 |
| chr3R | 23618718 | 23620331 | 1614 | 2,17 | 1406 |
| chr3R | 23636939 | 23638643 | 1705 | 2,1 | 1407 |
| chr3R | 23654732 | 23656629 | 1898 | 2,33 | 1408 |
| chr3R | 23678533 | 23679655 | 1123 | 2,17 | 1409 |
| chr3R | 23687648 | 23694176 | 6529 | 1,86 | 1410 |
| chr3R | 25050583 | 25054607 | 4025 | 2,18 | 1411 |
| chr3R | 27034239 | 27035342 | 1104 | 2,2 | 1412 |
| chr3RHet | 2 | 962 | 961 | 3,04 | 1413 |
| chr3RHet | 2767 | 5046 | 2280 | 2,14 | 1414 |
| chr3RHet | 6193 | 9149 | 2957 | 2,52 | 1415 |
| chr3RHet | 9332 | 11785 | 2454 | 2,43 | 1416 |
| chr3RHet | 11797 | 18764 | 6968 | 2,88 | 1417 |
| chr3RHet | 23379 | 24531 | 1153 | 2,56 | 1418 |
| chr3RHet | 24604 | 25795 | 1192 | 1,94 | 1419 |
| chr3RHet | 28689 | 30670 | 1982 | 2,52 | 1420 |
| chr3RHet | 31841 | 33552 | 1712 | 2,17 | 1421 |
| chr3RHet | 33828 | 38544 | 4717 | 2,17 | 1422 |
| chr3RHet | 38746 | 46530 | 7785 | 3,26 | 1423 |
| chr3RHet | 51014 | 52195 | 1182 | 3,24 | 1424 |
| chr3RHet | 52958 | 55603 | 2646 | 1,73 | 1425 |
| chr3RHet | 77925 | 80560 | 2636 | 2,72 | 1426 |
| chr3RHet | 80791 | 83943 | 3153 | 2,62 | 1427 |
| chr3RHet | 90776 | 92935 | 2160 | 2,39 | 1428 |
| chr3RHet | 97079 | 98261 | 1183 | 2,72 | 1429 |
| chr3RHet | 104441 | 109509 | 5069 | 2,39 | 1430 |
| chr3RHet | 111965 | 113210 | 1246 | 2,39 | 1431 |
| chr3RHet | 115651 | 119238 | 3588 | 3,39 | 1432 |
| chr3RHet | 119590 | 120737 | 1148 | 1,97 | 1433 |
| chr3RHet | 120904 | 122367 | 1464 | 2,14 | 1434 |
| chr3RHet | 122378 | 124289 | 1912 | 2,23 | 1435 |
| chr3RHet | 141808 | 143971 | 2164 | 2,23 | 1436 |
| chr3RHet | 145221 | 146923 | 1703 | 2,59 | 1437 |

| | | | | | |
|---|---|---|---|---|---|
| chr3RHet | 162709 | 165028 | 2320 | 2,09 | 1438 |
| chr3RHet | 178712 | 182162 | 3451 | 1,9 | 1439 |
| chr3RHet | 184978 | 187183 | 2206 | 2,26 | 1440 |
| chr3RHet | 187445 | 189771 | 2327 | 2,43 | 1441 |
| chr3RHet | 191955 | 203053 | 11099 | 3,75 | 1442 |
| chr3RHet | 205528 | 207931 | 2404 | 2,36 | 1443 |
| chr3RHet | 211090 | 213644 | 2555 | 2,56 | 1444 |
| chr3RHet | 219605 | 225010 | 5406 | 2,46 | 1445 |
| chr3RHet | 225815 | 227117 | 1303 | 2,36 | 1446 |
| chr3RHet | 227760 | 229957 | 2198 | 2,26 | 1447 |
| chr3RHet | 235499 | 237152 | 1654 | 2,36 | 1448 |
| chr3RHet | 237803 | 239188 | 1386 | 2,07 | 1449 |
| chr3RHet | 262442 | 264165 | 1724 | 2,1 | 1450 |
| chr3RHet | 266442 | 267784 | 1343 | 2,14 | 1451 |
| chr3RHet | 270081 | 271543 | 1463 | 2,04 | 1452 |
| chr3RHet | 272588 | 275892 | 3305 | 2,43 | 1453 |
| chr3RHet | 276088 | 282661 | 6574 | 2,56 | 1454 |
| chr3RHet | 286963 | 288736 | 1774 | 2,01 | 1455 |
| chr3RHet | 292207 | 295265 | 3059 | 2,36 | 1456 |
| chr3RHet | 296997 | 299274 | 2278 | 2,14 | 1457 |
| chr3RHet | 300617 | 303040 | 2424 | 2,33 | 1458 |
| chr3RHet | 312453 | 313911 | 1459 | 2,23 | 1459 |
| chr3RHet | 367596 | 369887 | 2292 | 2,65 | 1460 |
| chr3RHet | 372978 | 374799 | 1822 | 2,2 | 1461 |
| chr3RHet | 375261 | 384422 | 9162 | 2,82 | 1462 |
| chr3RHet | 393515 | 395426 | 1912 | 2,04 | 1463 |
| chr3RHet | 405775 | 406884 | 1110 | 2,52 | 1464 |
| chr3RHet | 409452 | 417157 | 7706 | 2,65 | 1465 |
| chr3RHet | 417367 | 423739 | 6373 | 2,72 | 1466 |
| chr3RHet | 424237 | 425453 | 1217 | 2,43 | 1467 |
| chr3RHet | 428450 | 429621 | 1172 | 2,52 | 1468 |
| chr3RHet | 429770 | 434825 | 5056 | 2,75 | 1469 |
| chr3RHet | 435194 | 438953 | 3760 | 2,12 | 1470 |
| chr3RHet | 522476 | 525508 | 3033 | 2,75 | 1471 |
| chr3RHet | 527706 | 532978 | 5273 | 3,07 | 1472 |
| chr3RHet | 543278 | 553439 | 10162 | 2,78 | 1473 |
| chr3RHet | 554390 | 560593 | 6204 | 3,3 | 1474 |
| chr3RHet | 560703 | 562879 | 2177 | 2,72 | 1475 |
| chr3RHet | 562892 | 569670 | 6779 | 2,39 | 1476 |
| chr3RHet | 569865 | 571190 | 1326 | 2,75 | 1477 |
| chr3RHet | 576467 | 581646 | 5180 | 3,17 | 1478 |
| chr3RHet | 582432 | 584737 | 2306 | 2,94 | 1479 |
| chr3RHet | 585301 | 587161 | 1861 | 2,39 | 1480 |
| chr3RHet | 591513 | 596183 | 4671 | 3,43 | 1481 |
| chr3RHet | 596793 | 600533 | 3741 | 2,56 | 1482 |
| chr3RHet | 601508 | 604600 | 3093 | 2,75 | 1483 |
| chr3RHet | 604721 | 616006 | 11286 | 3,2 | 1484 |
| chr3RHet | 634470 | 638786 | 4317 | 2,91 | 1485 |

| | | | | | |
|---|---|---|---|---|---|
| chr3RHet | 638970 | 641788 | 2819 | 2,75 | 1486 |
| chr3RHet | 643644 | 649003 | 5360 | 2,56 | 1487 |
| chr3RHet | 651641 | 659515 | 7875 | 3,43 | 1488 |
| chr3RHet | 663257 | 668314 | 5058 | 2,75 | 1489 |
| chr3RHet | 668543 | 670091 | 1549 | 1,97 | 1490 |
| chr3RHet | 670123 | 675768 | 5646 | 2,85 | 1491 |
| chr3RHet | 676708 | 681799 | 5092 | 2,65 | 1492 |
| chr3RHet | 682069 | 687365 | 5297 | 2,94 | 1493 |
| chr3RHet | 687367 | 688402 | 1036 | 2,82 | 1494 |
| chr3RHet | 701693 | 706510 | 4818 | 2,75 | 1495 |
| chr3RHet | 706588 | 710963 | 4376 | 2,69 | 1496 |
| chr3RHet | 710972 | 719696 | 8725 | 2,85 | 1497 |
| chr3RHet | 724702 | 730875 | 6174 | 2,65 | 1498 |
| chr3RHet | 731041 | 734495 | 3455 | 2,91 | 1499 |
| chr3RHet | 735573 | 739634 | 4062 | 2,69 | 1500 |
| chr3RHet | 740183 | 742144 | 1962 | 2,59 | 1501 |
| chr3RHet | 742344 | 745081 | 2738 | 2,33 | 1502 |
| chr3RHet | 745116 | 748242 | 3127 | 2,69 | 1503 |
| chr3RHet | 748253 | 750446 | 2194 | 2,23 | 1504 |
| chr3RHet | 750566 | 762340 | 11775 | 3,52 | 1505 |
| chr3RHet | 762822 | 764130 | 1309 | 2,62 | 1506 |
| chr3RHet | 764194 | 765546 | 1353 | 2,56 | 1507 |
| chr3RHet | 773490 | 775445 | 1956 | 2,39 | 1508 |
| chr3RHet | 776249 | 777410 | 1162 | 2,2 | 1509 |
| chr3RHet | 777520 | 784632 | 7113 | 3,82 | 1510 |
| chr3RHet | 795907 | 799315 | 3409 | 2,36 | 1511 |
| chr3RHet | 799807 | 802593 | 2787 | 2,56 | 1512 |
| chr3RHet | 803338 | 805045 | 1708 | 2,56 | 1513 |
| chr3RHet | 813124 | 818826 | 5703 | 4,3 | 1514 |
| chr3RHet | 819109 | 822132 | 3024 | 2,47 | 1515 |
| chr3RHet | 822388 | 825133 | 2746 | 2,78 | 1516 |
| chr3RHet | 826017 | 828461 | 2445 | 2,69 | 1517 |
| chr3RHet | 836918 | 838833 | 1916 | 2,26 | 1518 |
| chr3RHet | 839661 | 841294 | 1634 | 2,39 | 1519 |
| chr3RHet | 841464 | 843898 | 2435 | 2,1 | 1520 |
| chr3RHet | 858999 | 860316 | 1318 | 2,43 | 1521 |
| chr3RHet | 860635 | 864154 | 3520 | 2,43 | 1522 |
| chr3RHet | 864215 | 867110 | 2896 | 2,49 | 1523 |
| chr3RHet | 867531 | 869414 | 1884 | 2,75 | 1524 |
| chr3RHet | 869948 | 872409 | 2462 | 2,3 | 1525 |
| chr3RHet | 872555 | 873968 | 1414 | 2,52 | 1526 |
| chr3RHet | 876178 | 881111 | 4934 | 2,98 | 1527 |
| chr3RHet | 892856 | 895936 | 3081 | 2,36 | 1528 |
| chr3RHet | 902924 | 905491 | 2568 | 2,17 | 1529 |
| chr3RHet | 905614 | 906688 | 1075 | 2,33 | 1530 |
| chr3RHet | 907865 | 909421 | 1557 | 2,23 | 1531 |
| chr3RHet | 909493 | 910905 | 1413 | 2,46 | 1532 |
| chr3RHet | 914514 | 917897 | 3384 | 2,33 | 1533 |

| | | | | | |
|---|---|---|---|---|---|
| chr3RHet | 921653 | 923374 | 1722 | 2,04 | 1534 |
| chr3RHet | 937858 | 939123 | 1266 | 2,49 | 1535 |
| chr3RHet | 969449 | 970994 | 1546 | 2,85 | 1536 |
| chr3RHet | 972904 | 974761 | 1858 | 2,17 | 1537 |
| chr3RHet | 976513 | 980438 | 3926 | 2,69 | 1538 |
| chr3RHet | 982113 | 985418 | 3306 | 3,37 | 1539 |
| chr3RHet | 1016962 | 1018308 | 1347 | 2,33 | 1540 |
| chr3RHet | 1028049 | 1029481 | 1433 | 2,17 | 1541 |
| chr3RHet | 1029548 | 1031362 | 1815 | 2,23 | 1542 |
| chr3RHet | 1041530 | 1046287 | 4758 | 2,49 | 1543 |
| chr3RHet | 1046648 | 1048293 | 1646 | 2,36 | 1544 |
| chr3RHet | 1065603 | 1066717 | 1115 | 2,49 | 1545 |
| chr3RHet | 1090803 | 1092144 | 1342 | 2,14 | 1546 |
| chr3RHet | 1092177 | 1095717 | 3541 | 2,78 | 1547 |
| chr3RHet | 1096048 | 1097907 | 1860 | 3,49 | 1548 |
| chr3RHet | 1098669 | 1101822 | 3154 | 2,43 | 1549 |
| chr3RHet | 1110136 | 1111942 | 1807 | 2,14 | 1550 |
| chr3RHet | 1112828 | 1113908 | 1081 | 2,2 | 1551 |
| chr3RHet | 1117591 | 1119299 | 1709 | 2,69 | 1552 |
| chr3RHet | 1122381 | 1124018 | 1638 | 2,49 | 1553 |
| chr3RHet | 1124300 | 1126635 | 2336 | 2,91 | 1554 |
| chr3RHet | 1127250 | 1130205 | 2956 | 3,65 | 1555 |
| chr3RHet | 1132032 | 1134836 | 2805 | 3,43 | 1556 |
| chr3RHet | 1137247 | 1139924 | 2678 | 2,17 | 1557 |
| chr3RHet | 1156131 | 1157452 | 1322 | 2,14 | 1558 |
| chr3RHet | 1164330 | 1166119 | 1790 | 2,72 | 1559 |
| chr3RHet | 1167504 | 1169787 | 2284 | 2,3 | 1560 |
| chr3RHet | 1185366 | 1189544 | 4179 | 2,56 | 1561 |
| chr3RHet | 1191562 | 1193389 | 1828 | 2,17 | 1562 |
| chr3RHet | 1193450 | 1195419 | 1970 | 2,36 | 1563 |
| chr3RHet | 1202632 | 1204055 | 1424 | 2,04 | 1564 |
| chr3RHet | 1208881 | 1210582 | 1702 | 2,82 | 1565 |
| chr3RHet | 1212786 | 1214045 | 1260 | 1,97 | 1566 |
| chr3RHet | 1214058 | 1219179 | 5122 | 2,69 | 1567 |
| chr3RHet | 1219202 | 1220258 | 1057 | 2,01 | 1568 |
| chr3RHet | 1220860 | 1223821 | 2962 | 2,33 | 1569 |
| chr3RHet | 1227690 | 1228950 | 1261 | 2,39 | 1570 |
| chr3RHet | 1232143 | 1238973 | 6831 | 2,52 | 1571 |
| chr3RHet | 1245470 | 1248964 | 3495 | 4,17 | 1572 |
| chr3RHet | 1255116 | 1256457 | 1342 | 3,37 | 1573 |
| chr3RHet | 1258994 | 1260614 | 1621 | 2,49 | 1574 |
| chr3RHet | 1261074 | 1263432 | 2359 | 2,39 | 1575 |
| chr3RHet | 1264061 | 1266542 | 2482 | 2,26 | 1576 |
| chr3RHet | 1274337 | 1279749 | 5413 | 2,71 | 1577 |
| chr3RHet | 1289110 | 1291223 | 2114 | 2,62 | 1578 |
| chr3RHet | 1310682 | 1312240 | 1559 | 2,33 | 1579 |
| chr3RHet | 1312552 | 1313942 | 1391 | 2,26 | 1580 |
| chr3RHet | 1321712 | 1322859 | 1148 | 2,43 | 1581 |

| | | | | | |
|---|---|---|---|---|---|
| chr3RHet | 1336237 | 1337579 | 1343 | 2,07 | 1582 |
| chr3RHet | 1339003 | 1340398 | 1396 | 2,14 | 1583 |
| chr3RHet | 1341151 | 1342900 | 1750 | 2,26 | 1584 |
| chr3RHet | 1342995 | 1345168 | 2174 | 2,58 | 1585 |
| chr3RHet | 1347968 | 1350159 | 2192 | 2,49 | 1586 |
| chr3RHet | 1350391 | 1351681 | 1291 | 2,1 | 1587 |
| chr3RHet | 1363575 | 1365083 | 1509 | 2,65 | 1588 |
| chr3RHet | 1366157 | 1367340 | 1184 | 2,36 | 1589 |
| chr3RHet | 1367543 | 1368610 | 1068 | 2,1 | 1590 |
| chr3RHet | 1369267 | 1371242 | 1976 | 2,78 | 1591 |
| chr3RHet | 1371296 | 1372948 | 1653 | 2,26 | 1592 |
| chr3RHet | 1375162 | 1376793 | 1632 | 2,17 | 1593 |
| chr3RHet | 1376967 | 1380158 | 3192 | 2,85 | 1594 |
| chr3RHet | 1380238 | 1381615 | 1378 | 2,26 | 1595 |
| chr3RHet | 1381670 | 1383005 | 1336 | 2,43 | 1596 |
| chr3RHet | 1384640 | 1386095 | 1456 | 2,78 | 1597 |
| chr3RHet | 1386109 | 1387305 | 1197 | 2,07 | 1598 |
| chr3RHet | 1402974 | 1404639 | 1666 | 2,3 | 1599 |
| chr3RHet | 1405099 | 1406851 | 1753 | 2,17 | 1600 |
| chr3RHet | 1412153 | 1414810 | 2658 | 2,36 | 1601 |
| chr3RHet | 1414905 | 1416276 | 1372 | 2,1 | 1602 |
| chr3RHet | 1425809 | 1427902 | 2094 | 3,2 | 1603 |
| chr3RHet | 1428007 | 1429538 | 1532 | 2,33 | 1604 |
| chr3RHet | 1441786 | 1444458 | 2673 | 2,14 | 1605 |
| chr3RHet | 1444484 | 1447057 | 2574 | 2,49 | 1606 |
| chr3RHet | 1449241 | 1450356 | 1116 | 2,14 | 1607 |
| chr3RHet | 1451611 | 1453059 | 1449 | 2,39 | 1608 |
| chr3RHet | 1461742 | 1468501 | 6760 | 2,49 | 1609 |
| chr3RHet | 1468761 | 1471074 | 2314 | 3,88 | 1610 |
| chr3RHet | 1471720 | 1473090 | 1371 | 2,46 | 1611 |
| chr3RHet | 1480647 | 1482397 | 1751 | 2,14 | 1612 |
| chr3RHet | 1482531 | 1483859 | 1329 | 1,94 | 1613 |
| chr3RHet | 1490232 | 1491533 | 1302 | 1,91 | 1614 |
| chr3RHet | 1501241 | 1503221 | 1981 | 2,43 | 1615 |
| chr3RHet | 1503404 | 1504751 | 1348 | 2,1 | 1616 |
| chr3RHet | 1528261 | 1530231 | 1971 | 2,36 | 1617 |
| chr3RHet | 1536668 | 1537960 | 1293 | 2,07 | 1618 |
| chr3RHet | 1539931 | 1541981 | 2051 | 2,3 | 1619 |
| chr3RHet | 1542189 | 1543325 | 1137 | 2,52 | 1620 |
| chr3RHet | 1543450 | 1547413 | 3964 | 2,23 | 1621 |
| chr3RHet | 1547830 | 1549842 | 2013 | 2,14 | 1622 |
| chr3RHet | 1551519 | 1552700 | 1182 | 2,65 | 1623 |
| chr3RHet | 1561651 | 1563236 | 1586 | 2,07 | 1624 |
| chr3RHet | 1563555 | 1567710 | 4156 | 2,46 | 1625 |
| chr3RHet | 1567908 | 1570458 | 2551 | 2,17 | 1626 |
| chr3RHet | 1570523 | 1573507 | 2985 | 2,56 | 1627 |
| chr3RHet | 1574483 | 1576758 | 2276 | 2,77 | 1628 |
| chr3RHet | 1578262 | 1579527 | 1266 | 2,78 | 1629 |

| | | | | | |
|---|---|---|---|---|---|
| chr3RHet | 1581866 | 1585300 | 3435 | 3,14 | 1630 |
| chr3RHet | 1585831 | 1587847 | 2017 | 2,26 | 1631 |
| chr3RHet | 1588877 | 1591637 | 2761 | 2,78 | 1632 |
| chr3RHet | 1592150 | 1593277 | 1128 | 2,39 | 1633 |
| chr3RHet | 1593529 | 1594718 | 1190 | 2,33 | 1634 |
| chr3RHet | 1596104 | 1597384 | 1281 | 2,43 | 1635 |
| chr3RHet | 1598541 | 1600120 | 1580 | 4,71 | 1636 |
| chr3RHet | 1601533 | 1603033 | 1501 | 2,36 | 1637 |
| chr3RHet | 1603744 | 1605850 | 2107 | 2,62 | 1638 |
| chr3RHet | 1621607 | 1623163 | 1557 | 2,23 | 1639 |
| chr3RHet | 1623561 | 1626624 | 3064 | 2,2 | 1640 |
| chr3RHet | 1627023 | 1628144 | 1122 | 2,01 | 1641 |
| chr3RHet | 1628264 | 1629401 | 1138 | 1,94 | 1642 |
| chr3RHet | 1630246 | 1633594 | 3349 | 2,49 | 1643 |
| chr3RHet | 1633758 | 1636627 | 2870 | 2,69 | 1644 |
| chr3RHet | 1637403 | 1643453 | 6051 | 3,28 | 1645 |
| chr3RHet | 1672057 | 1673246 | 1190 | 2,33 | 1646 |
| chr3RHet | 1674536 | 1678015 | 3480 | 2,3 | 1647 |
| chr3RHet | 1678043 | 1680597 | 2555 | 2,72 | 1648 |
| chr3RHet | 1681949 | 1683432 | 1484 | 2,46 | 1649 |
| chr3RHet | 1691030 | 1701077 | 10048 | 2,48 | 1650 |
| chr3RHet | 1701386 | 1702950 | 1565 | 2,05 | 1651 |
| chr3RHet | 1707866 | 1710193 | 2328 | 2,31 | 1652 |
| chr3RHet | 1757867 | 1765067 | 7201 | 2,98 | 1653 |
| chr3RHet | 1772394 | 1773838 | 1445 | 2,07 | 1654 |
| chr3RHet | 1777328 | 1779781 | 2454 | 2,79 | 1655 |
| chr3RHet | 1780161 | 1781817 | 1657 | 2,49 | 1656 |
| chr3RHet | 1784441 | 1786800 | 2360 | 3,01 | 1657 |
| chr3RHet | 1787279 | 1788430 | 1152 | 1,88 | 1658 |
| chr3RHet | 1791962 | 1794410 | 2449 | 2,43 | 1659 |
| chr3RHet | 1799258 | 1801865 | 2608 | 2,56 | 1660 |
| chr3RHet | 1802566 | 1804200 | 1635 | 2,3 | 1661 |
| chr3RHet | 1804563 | 1805758 | 1196 | 2,56 | 1662 |
| chr3RHet | 1825332 | 1829019 | 3688 | 3,17 | 1663 |
| chr3RHet | 1829447 | 1833406 | 3960 | 2,72 | 1664 |
| chr3RHet | 1833934 | 1836500 | 2567 | 2,52 | 1665 |
| chr3RHet | 1837943 | 1839931 | 1989 | 2,36 | 1666 |
| chr3RHet | 1839941 | 1841858 | 1918 | 2,2 | 1667 |
| chr3RHet | 1841862 | 1845007 | 3146 | 2,52 | 1668 |
| chr3RHet | 1845291 | 1846874 | 1584 | 2,38 | 1669 |
| chr3RHet | 1851825 | 1856728 | 4904 | 2,59 | 1670 |
| chr3RHet | 1867124 | 1870649 | 3526 | 3,62 | 1671 |
| chr3RHet | 1872115 | 1877110 | 4996 | 3,46 | 1672 |
| chr3RHet | 1877231 | 1879412 | 2182 | 2,75 | 1673 |
| chr3RHet | 1879620 | 1881683 | 2064 | 2,17 | 1674 |
| chr3RHet | 1882592 | 1887440 | 4849 | 2,62 | 1675 |
| chr3RHet | 1887545 | 1890966 | 3422 | 2,23 | 1676 |
| chr3RHet | 1891049 | 1892991 | 1943 | 2,65 | 1677 |

| | | | | | |
|---|---|---|---|---|---|
| chr3RHet | 1902363 | 1910877 | 8515 | 2,75 | 1678 |
| chr3RHet | 1910881 | 1912213 | 1333 | 2,78 | 1679 |
| chr3RHet | 1913414 | 1915041 | 1628 | 2,17 | 1680 |
| chr3RHet | 1916459 | 1917892 | 1434 | 2,3 | 1681 |
| chr3RHet | 1918163 | 1930494 | 12332 | 2,78 | 1682 |
| chr3RHet | 1930518 | 1931702 | 1185 | 2,04 | 1683 |
| chr3RHet | 1942836 | 1945590 | 2755 | 2,23 | 1684 |
| chr3RHet | 1945653 | 1947958 | 2306 | 2,49 | 1685 |
| chr3RHet | 1992015 | 1993261 | 1247 | 2,43 | 1686 |
| chr3RHet | 1995156 | 2002372 | 7217 | 3,43 | 1687 |
| chr3RHet | 2017349 | 2019527 | 2179 | 2,62 | 1688 |
| chr3RHet | 2019886 | 2024554 | 4669 | 2,88 | 1689 |
| chr3RHet | 2024673 | 2027513 | 2841 | 2,36 | 1690 |
| chr3RHet | 2027771 | 2029748 | 1978 | 2,1 | 1691 |
| chr3RHet | 2031450 | 2034097 | 2648 | 2,36 | 1692 |
| chr3RHet | 2034619 | 2035607 | 989 | 2,46 | 1693 |
| chr3RHet | 2036020 | 2038863 | 2844 | 3,05 | 1694 |
| chr3RHet | 2040286 | 2045415 | 5130 | 2,85 | 1695 |
| chr3RHet | 2070964 | 2076279 | 5316 | 3,34 | 1696 |
| chr3RHet | 2081088 | 2082560 | 1473 | 2,26 | 1697 |
| chr3RHet | 2082747 | 2085358 | 2612 | 2,46 | 1698 |
| chr3RHet | 2091088 | 2097386 | 6299 | 2,46 | 1699 |
| chr3RHet | 2097429 | 2098531 | 1103 | 2,04 | 1700 |
| chr3RHet | 2102589 | 2106304 | 3716 | 2,52 | 1701 |
| chr3RHet | 2111835 | 2113105 | 1271 | 2,14 | 1702 |
| chr3RHet | 2117793 | 2121423 | 3631 | 2,82 | 1703 |
| chr3RHet | 2121485 | 2124609 | 3125 | 2,62 | 1704 |
| chr3RHet | 2125623 | 2129461 | 3839 | 2,36 | 1705 |
| chr3RHet | 2135800 | 2139969 | 4170 | 2,3 | 1706 |
| chr3RHet | 2140840 | 2142195 | 1356 | 2,91 | 1707 |
| chr3RHet | 2143008 | 2146413 | 3406 | 2,88 | 1708 |
| chr3RHet | 2147320 | 2149799 | 2480 | 2,33 | 1709 |
| chr3RHet | 2152162 | 2156771 | 4610 | 2,26 | 1710 |
| chr3RHet | 2156810 | 2158041 | 1232 | 2,49 | 1711 |
| chr3RHet | 2159853 | 2161299 | 1447 | 2,14 | 1712 |
| chr3RHet | 2162791 | 2167470 | 4680 | 3,64 | 1713 |
| chr3RHet | 2169707 | 2172665 | 2959 | 3,51 | 1714 |
| chr3RHet | 2174680 | 2177811 | 3132 | 3,52 | 1715 |
| chr3RHet | 2179128 | 2181491 | 2364 | 2,82 | 1716 |
| chr3RHet | 2181550 | 2182666 | 1117 | 1,97 | 1717 |
| chr3RHet | 2185261 | 2187800 | 2540 | 2,46 | 1718 |
| chr3RHet | 2187906 | 2192714 | 4809 | 2,39 | 1719 |
| chr3RHet | 2197818 | 2204475 | 6658 | 2,46 | 1720 |
| chr3RHet | 2207924 | 2210913 | 2990 | 2,52 | 1721 |
| chr3RHet | 2212535 | 2215703 | 3169 | 2,33 | 1722 |
| chr3RHet | 2221698 | 2223377 | 1680 | 2,17 | 1723 |
| chr3RHet | 2265004 | 2266356 | 1353 | 2,56 | 1724 |
| chr3RHet | 2267436 | 2270744 | 3309 | 2,17 | 1725 |

| | | | | | |
|---|---|---|---|---|---|
| chr3RHet | 2271423 | 2274385 | 2963 | 2,33 | 1726 |
| chr3RHet | 2274476 | 2276862 | 2387 | 2,75 | 1727 |
| chr3RHet | 2277162 | 2281677 | 4516 | 2,59 | 1728 |
| chr3RHet | 2282129 | 2296635 | 14507 | 2,85 | 1729 |
| chr3RHet | 2297303 | 2301210 | 3908 | 2,72 | 1730 |
| chr3RHet | 2301490 | 2306411 | 4922 | 2,82 | 1731 |
| chr3RHet | 2307738 | 2311609 | 3872 | 4,82 | 1732 |
| chr3RHet | 2312908 | 2315939 | 3032 | 2,14 | 1733 |
| chr3RHet | 2319234 | 2323565 | 4332 | 2,75 | 1734 |
| chr3RHet | 2323783 | 2325970 | 2188 | 2,49 | 1735 |
| chr3RHet | 2326355 | 2327694 | 1340 | 2,46 | 1736 |
| chr3RHet | 2329653 | 2332971 | 3319 | 2,46 | 1737 |
| chr3RHet | 2334438 | 2336575 | 2138 | 2,78 | 1738 |
| chr3RHet | 2338416 | 2340303 | 1888 | 3,01 | 1739 |
| chr3RHet | 2346295 | 2351693 | 5399 | 2,65 | 1740 |
| chr3RHet | 2351818 | 2353828 | 2011 | 2,82 | 1741 |
| chr3RHet | 2354318 | 2358816 | 4499 | 2,52 | 1742 |
| chr3RHet | 2359580 | 2360874 | 1295 | 2,23 | 1743 |
| chr3RHet | 2431764 | 2436400 | 4637 | 2,69 | 1744 |
| chr3RHet | 2436811 | 2440030 | 3220 | 2,39 | 1745 |
| chr3RHet | 2443417 | 2444793 | 1377 | 2,23 | 1746 |
| chr3RHet | 2456203 | 2458467 | 2265 | 2,17 | 1747 |
| chr3RHet | 2459105 | 2463890 | 4786 | 2,68 | 1748 |
| chr3RHet | 2465123 | 2470356 | 5234 | 2,85 | 1749 |
| chr3RHet | 2473771 | 2475168 | 1398 | 2,2 | 1750 |
| chr3RHet | 2483289 | 2485079 | 1791 | 2,43 | 1751 |
| chr3RHet | 2485303 | 2487452 | 2150 | 2,3 | 1752 |
| chr3RHet | 2488201 | 2492047 | 3847 | 3,07 | 1753 |
| chr4 | 28746 | 30677 | 1932 | 2,3 | 1754 |
| chr4 | 35490 | 40686 | 5197 | 2,88 | 1755 |
| chr4 | 42025 | 43455 | 1431 | 2,26 | 1756 |
| chr4 | 57802 | 58985 | 1184 | 2,62 | 1757 |
| chr4 | 306585 | 307825 | 1241 | 2,33 | 1758 |
| chr4 | 860235 | 862706 | 2472 | 2,53 | 1759 |
| chrU | 3453 | 8189 | 4737 | 2,66 | 1760 |
| chrU | 8719 | 14244 | 5526 | 3,3 | 1761 |
| chrU | 14509 | 18004 | 3496 | 2,72 | 1762 |
| chrU | 19044 | 24799 | 5756 | 3,37 | 1763 |
| chrU | 24833 | 29721 | 4889 | 4,32 | 1764 |
| chrU | 50327 | 58069 | 7743 | 2,69 | 1765 |
| chrU | 59062 | 63636 | 4575 | 2,43 | 1766 |
| chrU | 81743 | 90799 | 9057 | 3,4 | 1767 |
| chrU | 98886 | 106695 | 7810 | 3,11 | 1768 |
| chrU | 106805 | 111344 | 4540 | 2,6 | 1769 |
| chrU | 111366 | 113419 | 2054 | 2,52 | 1770 |
| chrU | 115209 | 118818 | 3610 | 2,65 | 1771 |
| chrU | 119608 | 130274 | 10667 | 2,94 | 1772 |
| chrU | 130741 | 139082 | 8342 | 3,2 | 1773 |

| | | | | | |
|---|---|---|---|---|---|
| chrU | 139296 | 142507 | 3212 | 2,65 | 1774 |
| chrU | 143126 | 147060 | 3935 | 2,69 | 1775 |
| chrU | 161486 | 163144 | 1659 | 4,11 | 1776 |
| chrU | 165225 | 177477 | 12253 | 2,47 | 1777 |
| chrU | 177782 | 179039 | 1258 | 2,56 | 1778 |
| chrU | 180243 | 182053 | 1811 | 2,36 | 1779 |
| chrU | 185040 | 186839 | 1800 | 1,91 | 1780 |
| chrU | 189134 | 190863 | 1730 | 2,17 | 1781 |
| chrU | 192058 | 193582 | 1525 | 2,56 | 1782 |
| chrU | 212481 | 221877 | 9397 | 3,07 | 1783 |
| chrU | 222060 | 224847 | 2788 | 2,1 | 1784 |
| chrU | 224917 | 226076 | 1160 | 2,17 | 1785 |
| chrU | 282954 | 286855 | 3902 | 2,13 | 1786 |
| chrU | 287467 | 288872 | 1406 | 2,07 | 1787 |
| chrU | 291995 | 293937 | 1943 | 2,3 | 1788 |
| chrU | 294637 | 296281 | 1645 | 2,01 | 1789 |
| chrU | 302334 | 305448 | 3115 | 2,26 | 1790 |
| chrU | 305769 | 318047 | 12279 | 2,69 | 1791 |
| chrU | 318412 | 321859 | 3448 | 2,39 | 1792 |
| chrU | 322386 | 323658 | 1273 | 2,26 | 1793 |
| chrU | 338413 | 339826 | 1414 | 2,2 | 1794 |
| chrU | 344682 | 350134 | 5453 | 3,2 | 1795 |
| chrU | 357615 | 359881 | 2267 | 2,2 | 1796 |
| chrU | 363121 | 366973 | 3853 | 3,06 | 1797 |
| chrU | 367345 | 370051 | 2707 | 2,82 | 1798 |
| chrU | 373124 | 374192 | 1069 | 2,46 | 1799 |
| chrU | 374392 | 375613 | 1222 | 2,07 | 1800 |
| chrU | 377163 | 379374 | 2212 | 2,52 | 1801 |
| chrU | 383815 | 389680 | 5866 | 2,14 | 1802 |
| chrU | 390196 | 392503 | 2308 | 2,39 | 1803 |
| chrU | 392523 | 393965 | 1443 | 2,23 | 1804 |
| chrU | 395498 | 397088 | 1591 | 2,01 | 1805 |
| chrU | 397501 | 401575 | 4075 | 2,3 | 1806 |
| chrU | 406185 | 407255 | 1071 | 2,23 | 1807 |
| chrU | 411961 | 415795 | 3835 | 2,43 | 1808 |
| chrU | 415997 | 418643 | 2647 | 3,3 | 1809 |
| chrU | 418647 | 429495 | 10849 | 3,24 | 1810 |
| chrU | 449185 | 451641 | 2457 | 2,69 | 1811 |
| chrU | 452575 | 454841 | 2267 | 2,91 | 1812 |
| chrU | 455037 | 459006 | 3970 | 3,85 | 1813 |
| chrU | 475656 | 477130 | 1475 | 2,23 | 1814 |
| chrU | 482729 | 484108 | 1380 | 2,23 | 1815 |
| chrU | 486972 | 489406 | 2435 | 2,26 | 1816 |
| chrU | 491427 | 495007 | 3581 | 2,23 | 1817 |
| chrU | 497653 | 499511 | 1859 | 2,33 | 1818 |
| chrU | 502312 | 505932 | 3621 | 3,01 | 1819 |
| chrU | 508423 | 512648 | 4226 | 2,59 | 1820 |
| chrU | 512774 | 515803 | 3030 | 2,82 | 1821 |

| | | | | | |
|---|---|---|---|---|---|
| chrU | 516053 | 518247 | 2195 | 2,36 | 1822 |
| chrU | 519346 | 520901 | 1556 | 2,3 | 1823 |
| chrU | 520981 | 523891 | 2911 | 2,14 | 1824 |
| chrU | 524852 | 530194 | 5343 | 4,37 | 1825 |
| chrU | 531097 | 532596 | 1500 | 2,46 | 1826 |
| chrU | 540300 | 541797 | 1498 | 2,36 | 1827 |
| chrU | 544165 | 548386 | 4222 | 2,62 | 1828 |
| chrU | 548861 | 549927 | 1067 | 2,3 | 1829 |
| chrU | 555812 | 557803 | 1992 | 2,39 | 1830 |
| chrU | 564469 | 566952 | 2484 | 2,23 | 1831 |
| chrU | 568183 | 570595 | 2413 | 2,39 | 1832 |
| chrU | 572555 | 575816 | 3262 | 2,52 | 1833 |
| chrU | 575923 | 577344 | 1422 | 1,94 | 1834 |
| chrU | 583527 | 586552 | 3026 | 2,2 | 1835 |
| chrU | 594799 | 596022 | 1224 | 2,07 | 1836 |
| chrU | 597293 | 599886 | 2594 | 2,52 | 1837 |
| chrU | 599922 | 601122 | 1201 | 2,23 | 1838 |
| chrU | 601243 | 602769 | 1527 | 2,2 | 1839 |
| chrU | 603105 | 604518 | 1414 | 2,07 | 1840 |
| chrU | 606924 | 608591 | 1668 | 2,23 | 1841 |
| chrU | 608900 | 610722 | 1823 | 2,72 | 1842 |
| chrU | 616216 | 620990 | 4775 | 3,66 | 1843 |
| chrU | 622041 | 623773 | 1733 | 2,65 | 1844 |
| chrU | 623809 | 624920 | 1112 | 2,26 | 1845 |
| chrU | 624952 | 628200 | 3249 | 2,39 | 1846 |
| chrU | 628423 | 629847 | 1425 | 2,43 | 1847 |
| chrU | 632895 | 634407 | 1513 | 2,2 | 1848 |
| chrU | 635817 | 637264 | 1448 | 1,94 | 1849 |
| chrU | 893751 | 894755 | 1005 | 4,14 | 1850 |
| chrU | 914697 | 917020 | 2324 | 2,78 | 1851 |
| chrU | 918217 | 920541 | 2325 | 2,26 | 1852 |
| chrU | 930700 | 932518 | 1819 | 2,2 | 1853 |
| chrU | 939937 | 941327 | 1391 | 2,43 | 1854 |
| chrU | 943509 | 944738 | 1230 | 1,97 | 1855 |
| chrU | 944808 | 945915 | 1108 | 2,26 | 1856 |
| chrU | 946533 | 948208 | 1676 | 2,62 | 1857 |
| chrU | 953716 | 956780 | 3065 | 2,88 | 1858 |
| chrU | 957291 | 959849 | 2559 | 2,75 | 1859 |
| chrU | 965243 | 969053 | 3811 | 2,72 | 1860 |
| chrU | 970633 | 972008 | 1376 | 3,17 | 1861 |
| chrU | 1032418 | 1033499 | 1082 | 2,36 | 1862 |
| chrU | 1033892 | 1035519 | 1628 | 2,49 | 1863 |
| chrU | 1036465 | 1037586 | 1122 | 2,39 | 1864 |
| chrU | 1041115 | 1044239 | 3125 | 3,42 | 1865 |
| chrU | 1044537 | 1048330 | 3794 | 2,72 | 1866 |
| chrU | 1053753 | 1055750 | 1998 | 2,62 | 1867 |
| chrU | 1057215 | 1058333 | 1119 | 2,33 | 1868 |
| chrU | 1061580 | 1063425 | 1846 | 2,3 | 1869 |

| | | | | | |
|---|---|---|---|---|---|
| chrU | 1063433 | 1065488 | 2056 | 2,56 | 1870 |
| chrU | 1075492 | 1082969 | 7478 | 2,24 | 1871 |
| chrU | 1084445 | 1086747 | 2303 | 2,37 | 1872 |
| chrU | 1104934 | 1106330 | 1397 | 2,9 | 1873 |
| chrU | 1154350 | 1157870 | 3521 | 2,75 | 1874 |
| chrU | 1159181 | 1160666 | 1486 | 1,84 | 1875 |
| chrU | 1165099 | 1169578 | 4480 | 3,01 | 1876 |
| chrU | 1187363 | 1190668 | 3306 | 2,65 | 1877 |
| chrU | 1190794 | 1195567 | 4774 | 2,82 | 1878 |
| chrU | 1198010 | 1200251 | 2242 | 2,49 | 1879 |
| chrU | 1200640 | 1206072 | 5433 | 2,93 | 1880 |
| chrU | 1206798 | 1210751 | 3954 | 2,82 | 1881 |
| chrU | 1213896 | 1215701 | 1806 | 2,65 | 1882 |
| chrU | 1225235 | 1226687 | 1453 | 2,04 | 1883 |
| chrU | 1227247 | 1233257 | 6011 | 2,86 | 1884 |
| chrU | 1235145 | 1243953 | 8809 | 2,72 | 1885 |
| chrU | 1244109 | 1245311 | 1203 | 3,43 | 1886 |
| chrU | 1245386 | 1247370 | 1985 | 3,37 | 1887 |
| chrU | 1274839 | 1285627 | 10789 | 2,72 | 1888 |
| chrU | 1315200 | 1320862 | 5663 | 2,72 | 1889 |
| chrU | 1321144 | 1328748 | 7605 | 2,78 | 1890 |
| chrU | 1328781 | 1332278 | 3498 | 2,56 | 1891 |
| chrU | 1332946 | 1336213 | 3268 | 2,72 | 1892 |
| chrU | 1365689 | 1372035 | 6347 | 2,72 | 1893 |
| chrU | 1372380 | 1377853 | 5474 | 2,72 | 1894 |
| chrU | 1392976 | 1398717 | 5742 | 4,22 | 1895 |
| chrU | 1399995 | 1401071 | 1077 | 2,65 | 1896 |
| chrU | 1407276 | 1409695 | 2420 | 2,26 | 1897 |
| chrU | 1435212 | 1438434 | 3223 | 2,82 | 1898 |
| chrU | 1444274 | 1449838 | 5565 | 2,52 | 1899 |
| chrU | 1450346 | 1452432 | 2087 | 2,33 | 1900 |
| chrU | 1453862 | 1455759 | 1898 | 2,56 | 1901 |
| chrU | 1458260 | 1460396 | 2137 | 2,17 | 1902 |
| chrU | 1472548 | 1475079 | 2532 | 2,89 | 1903 |
| chrU | 1550941 | 1552555 | 1615 | 2,35 | 1904 |
| chrU | 1554392 | 1556458 | 2067 | 2,46 | 1905 |
| chrU | 1557355 | 1559440 | 2086 | 2,41 | 1906 |
| chrU | 1560815 | 1563460 | 2646 | 2,43 | 1907 |
| chrU | 1563584 | 1564677 | 1094 | 2,65 | 1908 |
| chrU | 1564950 | 1572102 | 7153 | 3,07 | 1909 |
| chrU | 1585507 | 1587651 | 2145 | 2,46 | 1910 |
| chrU | 1587710 | 1594110 | 6401 | 2,96 | 1911 |
| chrU | 1594651 | 1598763 | 4113 | 2,47 | 1912 |
| chrU | 1598808 | 1600309 | 1502 | 2,36 | 1913 |
| chrU | 1613122 | 1614699 | 1578 | 1,97 | 1914 |
| chrU | 1621883 | 1623170 | 1288 | 2,01 | 1915 |
| chrU | 1626415 | 1627917 | 1503 | 1,97 | 1916 |
| chrU | 1637543 | 1639293 | 1751 | 2,88 | 1917 |

| | | | | | |
|---|---|---|---|---|---|
| chrU | 1650736 | 1653420 | 2685 | 2,33 | 1918 |
| chrU | 1655998 | 1657439 | 1442 | 2,46 | 1919 |
| chrU | 1660524 | 1662799 | 2276 | 2,3 | 1920 |
| chrU | 1664392 | 1665554 | 1163 | 2,82 | 1921 |
| chrU | 1665569 | 1668276 | 2708 | 2,62 | 1922 |
| chrU | 1669197 | 1673338 | 4142 | 2,56 | 1923 |
| chrU | 1675019 | 1677594 | 2576 | 2,82 | 1924 |
| chrU | 1696371 | 1697346 | 976 | 2,88 | 1925 |
| chrU | 1697688 | 1700324 | 2637 | 2,39 | 1926 |
| chrU | 1831192 | 1835432 | 4241 | 2,46 | 1927 |
| chrU | 1838457 | 1839890 | 1434 | 2,43 | 1928 |
| chrU | 1842854 | 1846266 | 3413 | 3,04 | 1929 |
| chrU | 1847065 | 1850690 | 3626 | 2,85 | 1930 |
| chrU | 1851542 | 1855597 | 4056 | 3,07 | 1931 |
| chrU | 1856405 | 1858629 | 2225 | 2,14 | 1932 |
| chrU | 1859319 | 1860488 | 1170 | 2,3 | 1933 |
| chrU | 1860692 | 1863442 | 2751 | 2,26 | 1934 |
| chrU | 1865105 | 1867514 | 2410 | 2,56 | 1935 |
| chrU | 1870541 | 1873519 | 2979 | 2,14 | 1936 |
| chrU | 1875156 | 1877568 | 2413 | 2,2 | 1937 |
| chrU | 1891915 | 1893419 | 1505 | 2,49 | 1938 |
| chrU | 1903316 | 1904660 | 1345 | 2,43 | 1939 |
| chrU | 1909837 | 1918754 | 8918 | 2,94 | 1940 |
| chrU | 1918777 | 1925252 | 6476 | 3,07 | 1941 |
| chrU | 1925343 | 1926434 | 1092 | 2,01 | 1942 |
| chrU | 1931172 | 1934320 | 3149 | 2,91 | 1943 |
| chrU | 1940733 | 1942496 | 1764 | 3,3 | 1944 |
| chrU | 1951627 | 1956065 | 4439 | 2,26 | 1945 |
| chrU | 1958876 | 1960170 | 1295 | 2,36 | 1946 |
| chrU | 1961231 | 1962463 | 1233 | 2,07 | 1947 |
| chrU | 1966275 | 1968590 | 2316 | 2,17 | 1948 |
| chrU | 1972232 | 1974170 | 1939 | 2,01 | 1949 |
| chrU | 1977046 | 1979151 | 2106 | 2,07 | 1950 |
| chrU | 1982272 | 1983411 | 1140 | 2,14 | 1951 |
| chrU | 2058094 | 2061059 | 2966 | 2,52 | 1952 |
| chrU | 2064032 | 2065724 | 1693 | 2,59 | 1953 |
| chrU | 2067902 | 2070239 | 2338 | 3,18 | 1954 |
| chrU | 2071216 | 2073461 | 2246 | 3,79 | 1955 |
| chrU | 2073513 | 2075113 | 1601 | 2,36 | 1956 |
| chrU | 2079816 | 2082949 | 3134 | 3,43 | 1957 |
| chrU | 2084311 | 2096143 | 11833 | 3,56 | 1958 |
| chrU | 2200834 | 2208602 | 7769 | 3,66 | 1959 |
| chrU | 2208680 | 2210840 | 2161 | 2,59 | 1960 |
| chrU | 2211534 | 2213756 | 2223 | 2,49 | 1961 |
| chrU | 2213873 | 2215892 | 2020 | 2,36 | 1962 |
| chrU | 2216208 | 2218037 | 1830 | 2,14 | 1963 |
| chrU | 2218484 | 2222760 | 4277 | 2,36 | 1964 |
| chrU | 2229333 | 2233456 | 4124 | 2,62 | 1965 |

| | | | | | |
|---|---|---|---|---|---|
| chrU | 2235095 | 2236280 | 1186 | 2,85 | 1966 |
| chrU | 2241996 | 2243334 | 1339 | 3,35 | 1967 |
| chrU | 2247873 | 2253863 | 5991 | 2,55 | 1968 |
| chrU | 2307430 | 2309022 | 1593 | 2,59 | 1969 |
| chrU | 2309081 | 2315079 | 5999 | 3,17 | 1970 |
| chrU | 2317458 | 2319223 | 1766 | 2,88 | 1971 |
| chrU | 2320123 | 2322663 | 2541 | 2,36 | 1972 |
| chrU | 2322894 | 2328694 | 5801 | 3,2 | 1973 |
| chrU | 2334020 | 2335847 | 1828 | 2,39 | 1974 |
| chrU | 2338772 | 2340223 | 1452 | 2,65 | 1975 |
| chrU | 2354295 | 2355693 | 1399 | 2,85 | 1976 |
| chrU | 2355947 | 2360292 | 4346 | 2,69 | 1977 |
| chrU | 2408095 | 2414528 | 6434 | 4,83 | 1978 |
| chrU | 2437670 | 2443494 | 5825 | 3,24 | 1979 |
| chrU | 2443526 | 2444764 | 1239 | 2,14 | 1980 |
| chrU | 2465017 | 2468457 | 3441 | 2,36 | 1981 |
| chrU | 2471752 | 2477814 | 6063 | 2,81 | 1982 |
| chrU | 2521496 | 2522815 | 1320 | 2,57 | 1983 |
| chrU | 2541642 | 2544140 | 2499 | 2,06 | 1984 |
| chrU | 2579744 | 2580709 | 966 | 3,25 | 1985 |
| chrU | 2606722 | 2616892 | 10171 | 3,3 | 1986 |
| chrU | 2616936 | 2620195 | 3260 | 2,98 | 1987 |
| chrU | 2620449 | 2624837 | 4389 | 2,59 | 1988 |
| chrU | 2663435 | 2664551 | 1117 | 2,17 | 1989 |
| chrU | 2674091 | 2676027 | 1937 | 2,65 | 1990 |
| chrU | 2696061 | 2698504 | 2444 | 2,59 | 1991 |
| chrU | 2698538 | 2700463 | 1926 | 2,07 | 1992 |
| chrU | 2700541 | 2704471 | 3931 | 2,56 | 1993 |
| chrU | 2712157 | 2713677 | 1521 | 2,07 | 1994 |
| chrU | 2717617 | 2719814 | 2198 | 2,07 | 1995 |
| chrU | 2768934 | 2769818 | 885 | 4,97 | 1996 |
| chrU | 2890085 | 2892983 | 2899 | 2,3 | 1997 |
| chrU | 2913452 | 2914931 | 1480 | 2,14 | 1998 |
| chrU | 2978569 | 2979680 | 1112 | 2,01 | 1999 |
| chrU | 2993298 | 2995982 | 2685 | 2,39 | 2000 |
| chrU | 3067012 | 3069769 | 2758 | 2,82 | 2001 |
| chrU | 3073041 | 3077318 | 4278 | 2,72 | 2002 |
| chrU | 3084228 | 3085917 | 1690 | 2,59 | 2003 |
| chrU | 3116850 | 3118197 | 1348 | 2,43 | 2004 |
| chrU | 3135925 | 3137379 | 1455 | 2,88 | 2005 |
| chrU | 3248192 | 3249445 | 1254 | 2,78 | 2006 |
| chrU | 3292444 | 3296179 | 3736 | 3,17 | 2007 |
| chrU | 3296364 | 3303670 | 7307 | 2,85 | 2008 |
| chrU | 3303679 | 3305075 | 1397 | 2,49 | 2009 |
| chrU | 3305108 | 3306434 | 1327 | 2,2 | 2010 |
| chrU | 3307283 | 3308377 | 1095 | 2,65 | 2011 |
| chrU | 3309917 | 3311551 | 1635 | 2,78 | 2012 |
| chrU | 3333782 | 3335997 | 2216 | 2,75 | 2013 |

| | | | | | |
|---|---|---|---|---|---|
| chrU | 3338489 | 3342846 | 4358 | 3,23 | 2014 |
| chrU | 3343227 | 3348812 | 5586 | 2,77 | 2015 |
| chrU | 3392703 | 3394384 | 1682 | 2,43 | 2016 |
| chrU | 3403438 | 3405363 | 1926 | 2,33 | 2017 |
| chrU | 3406090 | 3408817 | 2728 | 2,39 | 2018 |
| chrU | 3411297 | 3420892 | 9596 | 2,82 | 2019 |
| chrU | 3420943 | 3423659 | 2717 | 2,38 | 2020 |
| chrU | 3424322 | 3427630 | 3309 | 2,59 | 2021 |
| chrU | 3427850 | 3437576 | 9727 | 2,59 | 2022 |
| chrU | 3440274 | 3441648 | 1375 | 3,22 | 2023 |
| chrU | 3447511 | 3449913 | 2403 | 2,67 | 2024 |
| chrU | 3475064 | 3476189 | 1126 | 2,33 | 2025 |
| chrU | 3524271 | 3529296 | 5026 | 2,86 | 2026 |
| chrU | 3533485 | 3537775 | 4291 | 3,04 | 2027 |
| chrU | 3549368 | 3555962 | 6595 | 3,18 | 2028 |
| chrU | 3559174 | 3561517 | 2344 | 2,67 | 2029 |
| chrU | 3611798 | 3616955 | 5158 | 2,88 | 2030 |
| chrU | 3678044 | 3679294 | 1251 | 3,84 | 2031 |
| chrU | 3680360 | 3685334 | 4975 | 2,62 | 2032 |
| chrU | 3884144 | 3885501 | 1358 | 3,24 | 2033 |
| chrU | 3885663 | 3887418 | 1756 | 2,69 | 2034 |
| chrU | 3889502 | 3891186 | 1685 | 2,17 | 2035 |
| chrU | 3894908 | 3897833 | 2926 | 3,23 | 2036 |
| chrU | 3999408 | 4001642 | 2235 | 3,75 | 2037 |
| chrU | 4142390 | 4145647 | 3258 | 2,17 | 2038 |
| chrU | 4184693 | 4187392 | 2700 | 2,43 | 2039 |
| chrU | 4188710 | 4190270 | 1561 | 2,2 | 2040 |
| chrU | 4192629 | 4194206 | 1578 | 2,39 | 2041 |
| chrU | 4510191 | 4512370 | 2180 | 3,2 | 2042 |
| chrU | 4582960 | 4584405 | 1446 | 3,05 | 2043 |
| chrU | 4695059 | 4698767 | 3709 | 2,3 | 2044 |
| chrU | 4698784 | 4699932 | 1149 | 2,2 | 2045 |
| chrU | 4700840 | 4702615 | 1776 | 2,72 | 2046 |
| chrU | 4702660 | 4704398 | 1739 | 2,98 | 2047 |
| chrU | 4762955 | 4764381 | 1427 | 2,1 | 2048 |
| chrU | 4786796 | 4791744 | 4949 | 2,32 | 2049 |
| chrU | 4791878 | 4798258 | 6381 | 2,39 | 2050 |
| chrU | 4803899 | 4807180 | 3282 | 2,2 | 2051 |
| chrU | 4816282 | 4817432 | 1151 | 2,2 | 2052 |
| chrU | 5185344 | 5188872 | 3529 | 3,22 | 2053 |
| chrU | 5202689 | 5204089 | 1401 | 2,98 | 2054 |
| chrU | 5213053 | 5214223 | 1171 | 2,43 | 2055 |
| chrU | 5466807 | 5468175 | 1369 | 2,82 | 2056 |
| chrU | 5491656 | 5493304 | 1649 | 2,23 | 2057 |
| chrU | 5494665 | 5498703 | 4039 | 2,72 | 2058 |
| chrU | 5507986 | 5510519 | 2534 | 2,52 | 2059 |
| chrU | 5528127 | 5529448 | 1322 | 4,57 | 2060 |
| chrU | 5546256 | 5547510 | 1255 | 2,3 | 2061 |

| | | | | | |
|---|---|---|---|---|---|
| chrU | 5556829 | 5559535 | 2707 | 4,27 | 2062 |
| chrU | 5560114 | 5561837 | 1724 | 2,69 | 2063 |
| chrU | 5578722 | 5582323 | 3602 | 2,26 | 2064 |
| chrU | 5582518 | 5584295 | 1778 | 6,21 | 2065 |
| chrU | 5599053 | 5603332 | 4280 | 2,94 | 2066 |
| chrU | 5603832 | 5608076 | 4245 | 2,84 | 2067 |
| chrU | 5608978 | 5610235 | 1258 | 2,69 | 2068 |
| chrU | 5620372 | 5621952 | 1581 | 3,17 | 2069 |
| chrU | 5624544 | 5625636 | 1093 | 3,11 | 2070 |
| chrU | 5626596 | 5628860 | 2265 | 2,72 | 2071 |
| chrU | 5637672 | 5639966 | 2295 | 2,52 | 2072 |
| chrU | 5640039 | 5641840 | 1802 | 2,43 | 2073 |
| chrU | 5666706 | 5669534 | 2829 | 2,01 | 2074 |
| chrU | 5669729 | 5672574 | 2846 | 2,17 | 2075 |
| chrU | 5672794 | 5677128 | 4335 | 2,55 | 2076 |
| chrU | 5722003 | 5723501 | 1499 | 2,39 | 2077 |
| chrU | 5724276 | 5726471 | 2196 | 2,33 | 2078 |
| chrU | 5775819 | 5777003 | 1185 | 3,04 | 2079 |
| chrU | 5815861 | 5817643 | 1783 | 2,39 | 2080 |
| chrU | 5817691 | 5820997 | 3307 | 2,52 | 2081 |
| chrU | 5858084 | 5862943 | 4860 | 2,57 | 2082 |
| chrU | 5888343 | 5889545 | 1203 | 2,3 | 2083 |
| chrU | 5891201 | 5892423 | 1223 | 2,23 | 2084 |
| chrU | 5923835 | 5925735 | 1901 | 2,78 | 2085 |
| chrU | 5926629 | 5928016 | 1388 | 2,17 | 2086 |
| chrU | 5928774 | 5933094 | 4321 | 2,43 | 2087 |
| chrU | 5944176 | 5946271 | 2096 | 2,69 | 2088 |
| chrU | 5953539 | 5956806 | 3268 | 2,62 | 2089 |
| chrU | 5968109 | 5969472 | 1364 | 2,23 | 2090 |
| chrU | 5969798 | 5970976 | 1179 | 1,91 | 2091 |
| chrU | 5978018 | 5986488 | 8471 | 3,59 | 2092 |
| chrU | 5989130 | 5990589 | 1460 | 2,36 | 2093 |
| chrU | 6000847 | 6003523 | 2677 | 3,24 | 2094 |
| chrU | 6003772 | 6005082 | 1311 | 2,36 | 2095 |
| chrU | 6031268 | 6032906 | 1639 | 2,14 | 2096 |
| chrU | 6046582 | 6049493 | 2912 | 3,39 | 2097 |
| chrU | 6060084 | 6063781 | 3698 | 2,91 | 2098 |
| chrU | 6109083 | 6113709 | 4627 | 3,04 | 2099 |
| chrU | 6154483 | 6156870 | 2388 | 3,17 | 2100 |
| chrU | 6185642 | 6187076 | 1435 | 3,13 | 2101 |
| chrU | 6197426 | 6199399 | 1974 | 2,3 | 2102 |
| chrU | 6199778 | 6201050 | 1273 | 2,69 | 2103 |
| chrU | 6216314 | 6218629 | 2316 | 2,27 | 2104 |
| chrU | 6259579 | 6261331 | 1753 | 2,85 | 2105 |
| chrU | 6303080 | 6305534 | 2455 | 2,19 | 2106 |
| chrU | 6313982 | 6316806 | 2825 | 3,28 | 2107 |
| chrU | 6323202 | 6325743 | 2542 | 3,15 | 2108 |
| chrU | 6418331 | 6420910 | 2580 | 2,3 | 2109 |

| | | | | | |
|---|---|---|---|---|---|
| chrU | 6455810 | 6462815 | 7006 | 2,42 | 2110 |
| chrU | 6528429 | 6532270 | 3842 | 2,75 | 2111 |
| chrU | 6539607 | 6542491 | 2885 | 2,39 | 2112 |
| chrU | 6565161 | 6568675 | 3515 | 2,74 | 2113 |
| chrU | 6571590 | 6574663 | 3074 | 2,97 | 2114 |
| chrU | 6588769 | 6592337 | 3569 | 2,32 | 2115 |
| chrU | 6609688 | 6612663 | 2976 | 2,05 | 2116 |
| chrU | 6621157 | 6622364 | 1208 | 3,43 | 2117 |
| chrU | 6640477 | 6643842 | 3366 | 2,91 | 2118 |
| chrU | 6660793 | 6661882 | 1090 | 3,92 | 2119 |
| chrU | 6678512 | 6681017 | 2506 | 2,45 | 2120 |
| chrU | 6738078 | 6740735 | 2658 | 2,26 | 2121 |
| chrU | 6745439 | 6747101 | 1663 | 2,07 | 2122 |
| chrU | 6757103 | 6759492 | 2390 | 2,45 | 2123 |
| chrU | 6784068 | 6786066 | 1999 | 1,97 | 2124 |
| chrU | 6791133 | 6792861 | 1729 | 2,01 | 2125 |
| chrU | 6796885 | 6797966 | 1082 | 2,14 | 2126 |
| chrU | 6853575 | 6855944 | 2370 | 2,17 | 2127 |
| chrU | 6859469 | 6862720 | 3252 | 2,36 | 2128 |
| chrU | 6878805 | 6880328 | 1524 | 2,25 | 2129 |
| chrU | 6881273 | 6882367 | 1095 | 2,56 | 2130 |
| chrU | 6893800 | 6895618 | 1819 | 2,07 | 2131 |
| chrU | 6900014 | 6901975 | 1962 | 2,26 | 2132 |
| chrU | 6915173 | 6917868 | 2696 | 2,51 | 2133 |
| chrU | 6963404 | 6966106 | 2703 | 2,94 | 2134 |
| chrU | 6993155 | 6995987 | 2833 | 2,49 | 2135 |
| chrU | 7008704 | 7010322 | 1619 | 3,04 | 2136 |
| chrU | 7011149 | 7013739 | 2591 | 2,07 | 2137 |
| chrU | 7014344 | 7015545 | 1202 | 2,39 | 2138 |
| chrU | 7021069 | 7023124 | 2056 | 2,73 | 2139 |
| chrU | 7034910 | 7037614 | 2705 | 2,46 | 2140 |
| chrU | 7056149 | 7058211 | 2063 | 2,59 | 2141 |
| chrU | 7064171 | 7066093 | 1923 | 2,43 | 2142 |
| chrU | 7066922 | 7069528 | 2607 | 2,29 | 2143 |
| chrU | 7088768 | 7090367 | 1600 | 3,3 | 2144 |
| chrU | 7137052 | 7138909 | 1858 | 2,37 | 2145 |
| chrU | 7139280 | 7142230 | 2951 | 2,33 | 2146 |
| chrU | 7148090 | 7150475 | 2386 | 2,49 | 2147 |
| chrU | 7167805 | 7171003 | 3199 | 2,78 | 2148 |
| chrU | 7171309 | 7172533 | 1225 | 2,14 | 2149 |
| chrU | 7190340 | 7192985 | 2646 | 2,24 | 2150 |
| chrU | 7196318 | 7201226 | 4909 | 2,52 | 2151 |
| chrU | 7207035 | 7209957 | 2923 | 2,88 | 2152 |
| chrU | 7236196 | 7237473 | 1278 | 2,33 | 2153 |
| chrU | 7240277 | 7242997 | 2721 | 2,32 | 2154 |
| chrU | 7245646 | 7247859 | 2214 | 2,17 | 2155 |
| chrU | 7248525 | 7250926 | 2402 | 2,39 | 2156 |
| chrU | 7251149 | 7252376 | 1228 | 2,56 | 2157 |

| | | | | | |
|---|---|---|---|---|---|
| chrU | 7270211 | 7273093 | 2883 | 2,58 | 2158 |
| chrU | 7281130 | 7283870 | 2741 | 2,54 | 2159 |
| chrU | 7286464 | 7289221 | 2758 | 2,17 | 2160 |
| chrU | 7297263 | 7300042 | 2780 | 2,3 | 2161 |
| chrU | 7303595 | 7305388 | 1794 | 2,6 | 2162 |
| chrU | 7315131 | 7316829 | 1699 | 4,63 | 2163 |
| chrU | 7340285 | 7342410 | 2126 | 2,03 | 2164 |
| chrU | 7345613 | 7347550 | 1938 | 2,23 | 2165 |
| chrU | 7384996 | 7387042 | 2047 | 2,24 | 2166 |
| chrU | 7406001 | 7407129 | 1129 | 2,1 | 2167 |
| chrU | 7408501 | 7411276 | 2776 | 2,56 | 2168 |
| chrU | 7439958 | 7441991 | 2034 | 3,77 | 2169 |
| chrU | 7452230 | 7454918 | 2689 | 2,35 | 2170 |
| chrU | 7467450 | 7470142 | 2693 | 2,12 | 2171 |
| chrU | 7477814 | 7480135 | 2322 | 2,57 | 2172 |
| chrU | 7491891 | 7495406 | 3516 | 3,43 | 2173 |
| chrU | 7502793 | 7505061 | 2269 | 2,54 | 2174 |
| chrU | 7507808 | 7510406 | 2599 | 2,25 | 2175 |
| chrU | 7520263 | 7522322 | 2060 | 2,14 | 2176 |
| chrU | 7538712 | 7540080 | 1369 | 2,06 | 2177 |
| chrU | 7558059 | 7560112 | 2054 | 2,45 | 2178 |
| chrU | 7565257 | 7566872 | 1616 | 2,48 | 2179 |
| chrU | 7599187 | 7601895 | 2709 | 2,31 | 2180 |
| chrU | 7616392 | 7618751 | 2360 | 2,2 | 2181 |
| chrU | 7628264 | 7630647 | 2384 | 2,75 | 2182 |
| chrU | 7640299 | 7642634 | 2336 | 2,61 | 2183 |
| chrU | 7647577 | 7649592 | 2016 | 2,33 | 2184 |
| chrU | 7650255 | 7652328 | 2074 | 2,56 | 2185 |
| chrU | 7656038 | 7657030 | 993 | 2,14 | 2186 |
| chrU | 7659369 | 7660762 | 1394 | 2,43 | 2187 |
| chrU | 7667348 | 7668961 | 1614 | 2,1 | 2188 |
| chrU | 7671387 | 7673484 | 2098 | 2,36 | 2189 |
| chrU | 7673673 | 7674797 | 1125 | 2,36 | 2190 |
| chrU | 7674957 | 7675952 | 996 | 2,62 | 2191 |
| chrU | 7687906 | 7689899 | 1994 | 2,05 | 2192 |
| chrU | 7693347 | 7694730 | 1384 | 2,26 | 2193 |
| chrU | 7711382 | 7713456 | 2075 | 2,39 | 2194 |
| chrU | 7715585 | 7718660 | 3076 | 3,11 | 2195 |
| chrU | 7719161 | 7722687 | 3527 | 2,72 | 2196 |
| chrU | 7727442 | 7729119 | 1678 | 2,22 | 2197 |
| chrU | 7731727 | 7733859 | 2133 | 2,69 | 2198 |
| chrU | 7745534 | 7746865 | 1332 | 2,49 | 2199 |
| chrU | 7751337 | 7752415 | 1079 | 2,36 | 2200 |
| chrU | 7752695 | 7754558 | 1864 | 2,36 | 2201 |
| chrU | 7761544 | 7763939 | 2396 | 2,04 | 2202 |
| chrU | 7768369 | 7770540 | 2172 | 2,56 | 2203 |
| chrU | 7773683 | 7775300 | 1618 | 2,3 | 2204 |
| chrU | 7789994 | 7793187 | 3194 | 2,65 | 2205 |

| | | | | | |
|---|---|---|---|---|---|
| chrU | 7799941 | 7801528 | 1588 | 2,25 | 2206 |
| chrU | 7813077 | 7815502 | 2426 | 2,3 | 2207 |
| chrU | 7815596 | 7817546 | 1951 | 2,9 | 2208 |
| chrU | 7817757 | 7819549 | 1793 | 2,26 | 2209 |
| chrU | 7841769 | 7843939 | 2171 | 2,59 | 2210 |
| chrU | 7846315 | 7847728 | 1414 | 2,2 | 2211 |
| chrU | 7852860 | 7854507 | 1648 | 2,2 | 2212 |
| chrU | 7874809 | 7881005 | 6197 | 2,26 | 2213 |
| chrU | 7887430 | 7890190 | 2761 | 2,3 | 2214 |
| chrU | 7899249 | 7900712 | 1464 | 2,3 | 2215 |
| chrU | 7917915 | 7919147 | 1233 | 2,06 | 2216 |
| chrU | 7930930 | 7932881 | 1952 | 2,3 | 2217 |
| chrU | 7933606 | 7935328 | 1723 | 3,95 | 2218 |
| chrU | 7937325 | 7939167 | 1843 | 2,1 | 2219 |
| chrU | 7947790 | 7949363 | 1574 | 1,84 | 2220 |
| chrU | 7960302 | 7961934 | 1633 | 2,91 | 2221 |
| chrU | 7964885 | 7966167 | 1283 | 1,91 | 2222 |
| chrU | 7966463 | 7970807 | 4345 | 3,93 | 2223 |
| chrU | 7981184 | 7983170 | 1987 | 2,01 | 2224 |
| chrU | 7997638 | 7999973 | 2336 | 3,27 | 2225 |
| chrU | 8003935 | 8006086 | 2152 | 2,91 | 2226 |
| chrU | 8026913 | 8028728 | 1816 | 2,52 | 2227 |
| chrU | 8034803 | 8036787 | 1985 | 1,84 | 2228 |
| chrU | 8049137 | 8050811 | 1675 | 2,3 | 2229 |
| chrU | 8051005 | 8053181 | 2177 | 2,65 | 2230 |
| chrU | 8063293 | 8065320 | 2028 | 2,29 | 2231 |
| chrU | 8079533 | 8081315 | 1783 | 2,83 | 2232 |
| chrU | 8085619 | 8087398 | 1780 | 1,94 | 2233 |
| chrU | 8089351 | 8091305 | 1955 | 2,34 | 2234 |
| chrU | 8097399 | 8100108 | 2710 | 3,73 | 2235 |
| chrU | 8105578 | 8109117 | 3540 | 2,43 | 2236 |
| chrU | 8110137 | 8111236 | 1100 | 2,01 | 2237 |
| chrU | 8129392 | 8131307 | 1916 | 2,45 | 2238 |
| chrU | 8133255 | 8135327 | 2073 | 2,1 | 2239 |
| chrU | 8139097 | 8141123 | 2027 | 2,02 | 2240 |
| chrU | 8141212 | 8145237 | 4026 | 2,61 | 2241 |
| chrU | 8149226 | 8150399 | 1174 | 2,46 | 2242 |
| chrU | 8160636 | 8162752 | 2117 | 2,01 | 2243 |
| chrU | 8164709 | 8166515 | 1807 | 2,1 | 2244 |
| chrU | 8166543 | 8168642 | 2100 | 2,2 | 2245 |
| chrU | 8168972 | 8170218 | 1247 | 1,97 | 2246 |
| chrU | 8183985 | 8189673 | 5689 | 2,32 | 2247 |
| chrU | 8202938 | 8204905 | 1968 | 2,26 | 2248 |
| chrU | 8212827 | 8214486 | 1660 | 2,43 | 2249 |
| chrU | 8216206 | 8219949 | 3744 | 2,46 | 2250 |
| chrU | 8222203 | 8223830 | 1628 | 2,16 | 2251 |
| chrU | 8231250 | 8232477 | 1228 | 2,67 | 2252 |
| chrU | 8251791 | 8253521 | 1731 | 1,91 | 2253 |

| | | | | | |
|---|---|---|---|---|---|
| chrU | 8257300 | 8261161 | 3862 | 2,29 | 2254 |
| chrU | 8262979 | 8266692 | 3714 | 2,57 | 2255 |
| chrU | 8275885 | 8277630 | 1746 | 2,24 | 2256 |
| chrU | 8281081 | 8283003 | 1923 | 2,39 | 2257 |
| chrU | 8293589 | 8297402 | 3814 | 2,54 | 2258 |
| chrU | 8299207 | 8301105 | 1899 | 2,36 | 2259 |
| chrU | 8303174 | 8304656 | 1483 | 2,1 | 2260 |
| chrU | 8311916 | 8313416 | 1501 | 2,43 | 2261 |
| chrU | 8315373 | 8317330 | 1958 | 2,43 | 2262 |
| chrU | 8324256 | 8325712 | 1457 | 2,12 | 2263 |
| chrU | 8331852 | 8338484 | 6633 | 2,43 | 2264 |
| chrU | 8343526 | 8344661 | 1136 | 2,85 | 2265 |
| chrU | 8348715 | 8350578 | 1864 | 2,49 | 2266 |
| chrU | 8357585 | 8359199 | 1615 | 2,94 | 2267 |
| chrU | 8403168 | 8404904 | 1737 | 2,49 | 2268 |
| chrU | 8405046 | 8406621 | 1576 | 2,49 | 2269 |
| chrU | 8408231 | 8409693 | 1463 | 2,2 | 2270 |
| chrU | 8409913 | 8411435 | 1523 | 2,43 | 2271 |
| chrU | 8421470 | 8422946 | 1477 | 2,33 | 2272 |
| chrU | 8427871 | 8429747 | 1877 | 2,43 | 2273 |
| chrU | 8441019 | 8442480 | 1462 | 2,52 | 2274 |
| chrU | 8455680 | 8457145 | 1466 | 2,09 | 2275 |
| chrU | 8457637 | 8458912 | 1276 | 2,26 | 2276 |
| chrU | 8465359 | 8466908 | 1550 | 2,17 | 2277 |
| chrU | 8476859 | 8477955 | 1097 | 2,23 | 2278 |
| chrU | 8479847 | 8482948 | 3102 | 2,66 | 2279 |
| chrU | 8489344 | 8490918 | 1575 | 2,61 | 2280 |
| chrU | 8494276 | 8495460 | 1185 | 2,22 | 2281 |
| chrU | 8521127 | 8522309 | 1183 | 2,25 | 2282 |
| chrU | 8522322 | 8523950 | 1629 | 2,07 | 2283 |
| chrU | 8530421 | 8531759 | 1339 | 2,52 | 2284 |
| chrU | 8545565 | 8547140 | 1576 | 2,36 | 2285 |
| chrU | 8556830 | 8557910 | 1081 | 2,3 | 2286 |
| chrU | 8565832 | 8566932 | 1101 | 1,81 | 2287 |
| chrU | 8573043 | 8574772 | 1730 | 2,71 | 2288 |
| chrU | 8576573 | 8577645 | 1073 | 2,02 | 2289 |
| chrU | 8577731 | 8578808 | 1078 | 2,2 | 2290 |
| chrU | 8585260 | 8586639 | 1380 | 2,41 | 2291 |
| chrU | 8609278 | 8609856 | 579 | 4,86 | 2292 |
| chrU | 8624276 | 8625803 | 1528 | 2,14 | 2293 |
| chrU | 8637566 | 8639220 | 1655 | 2,12 | 2294 |
| chrU | 8668732 | 8670044 | 1313 | 2,1 | 2295 |
| chrU | 8683355 | 8684755 | 1401 | 2,2 | 2296 |
| chrU | 8686131 | 8687631 | 1501 | 2,24 | 2297 |
| chrU | 8707969 | 8709314 | 1346 | 2,14 | 2298 |
| chrU | 8717954 | 8719454 | 1501 | 2,04 | 2299 |
| chrU | 8742257 | 8743798 | 1542 | 2,49 | 2300 |
| chrU | 8788695 | 8790478 | 1784 | 3,17 | 2301 |

| | | | | | |
|---|---|---|---|---|---|
| chrU | 8812961 | 8814031 | 1071 | 2,26 | 2302 |
| chrU | 8830681 | 8832064 | 1384 | 2,75 | 2303 |
| chrU | 8851423 | 8852814 | 1392 | 2,07 | 2304 |
| chrU | 8869199 | 8870483 | 1285 | 1,91 | 2305 |
| chrU | 8882787 | 8883949 | 1163 | 2,65 | 2306 |
| chrU | 8895541 | 8896860 | 1320 | 3,68 | 2307 |
| chrU | 8900671 | 8901992 | 1322 | 2,01 | 2308 |
| chrU | 8919560 | 8922315 | 2756 | 2,14 | 2309 |
| chrU | 8931925 | 8932979 | 1055 | 3,69 | 2310 |
| chrU | 8934478 | 8935794 | 1317 | 2,1 | 2311 |
| chrU | 8940290 | 8942552 | 2263 | 2,49 | 2312 |
| chrU | 8947864 | 8949053 | 1190 | 2,05 | 2313 |
| chrU | 8951991 | 8953522 | 1532 | 2,59 | 2314 |
| chrU | 8959896 | 8961396 | 1501 | 2,1 | 2315 |
| chrU | 8989304 | 8990423 | 1120 | 2,17 | 2316 |
| chrU | 9001169 | 9002377 | 1209 | 2,11 | 2317 |
| chrU | 9006540 | 9008261 | 1722 | 1,99 | 2318 |
| chrU | 9046329 | 9047636 | 1308 | 2,33 | 2319 |
| chrU | 9089370 | 9092224 | 2855 | 3,11 | 2320 |
| chrU | 9099938 | 9101311 | 1374 | 2,01 | 2321 |
| chrU | 9116579 | 9118451 | 1873 | 2,29 | 2322 |
| chrU | 9123388 | 9126109 | 2722 | 3,07 | 2323 |
| chrU | 9142812 | 9143967 | 1156 | 8,09 | 2324 |
| chrU | 9150672 | 9151973 | 1302 | 9,52 | 2325 |
| chrU | 9153080 | 9154505 | 1426 | 2,39 | 2326 |
| chrU | 9169653 | 9170412 | 760 | 6,21 | 2327 |
| chrU | 9171180 | 9172436 | 1257 | 2,07 | 2328 |
| chrU | 9178907 | 9180147 | 1241 | 2,34 | 2329 |
| chrU | 9208396 | 9209736 | 1341 | 2,33 | 2330 |
| chrU | 9214691 | 9216078 | 1388 | 2,73 | 2331 |
| chrU | 9274168 | 9275624 | 1457 | 2,52 | 2332 |
| chrU | 9291950 | 9293184 | 1235 | 2,72 | 2333 |
| chrU | 9294645 | 9295769 | 1125 | 2,3 | 2334 |
| chrU | 9341873 | 9342916 | 1044 | 3,01 | 2335 |
| chrU | 9356861 | 9357779 | 919 | 2,26 | 2336 |
| chrU | 9357977 | 9359486 | 1510 | 3,2 | 2337 |
| chrU | 9365587 | 9366746 | 1160 | 2,17 | 2338 |
| chrU | 9415192 | 9416367 | 1176 | 2,26 | 2339 |
| chrU | 9452384 | 9453486 | 1103 | 3,33 | 2340 |
| chrU | 9456596 | 9458555 | 1960 | 2,41 | 2341 |
| chrU | 9488795 | 9489889 | 1095 | 2,49 | 2342 |
| chrU | 9551275 | 9553023 | 1749 | 2,46 | 2343 |
| chrU | 9554067 | 9555287 | 1221 | 2,13 | 2344 |
| chrU | 9582645 | 9584130 | 1486 | 3,4 | 2345 |
| chrU | 9586571 | 9588970 | 2400 | 2,39 | 2346 |
| chrU | 9591358 | 9593456 | 2099 | 3,61 | 2347 |
| chrU | 9597434 | 9598474 | 1041 | 2,01 | 2348 |
| chrU | 9598869 | 9599444 | 576 | 3,59 | 2349 |

| | | | | | |
|---|---|---|---|---|---|
| chrU | 9599816 | 9600931 | 1116 | 2,72 | 2350 |
| chrU | 9625503 | 9627401 | 1899 | 3,08 | 2351 |
| chrU | 9635300 | 9636343 | 1044 | 2,65 | 2352 |
| chrU | 9650679 | 9651933 | 1255 | 2,01 | 2353 |
| chrU | 9658787 | 9660398 | 1612 | 2,35 | 2354 |
| chrU | 9691686 | 9692873 | 1188 | 7,18 | 2355 |
| chrU | 9727056 | 9728319 | 1264 | 2,3 | 2356 |
| chrU | 9742131 | 9743446 | 1316 | 2,82 | 2357 |
| chrU | 9773434 | 9774700 | 1267 | 3,03 | 2358 |
| chrU | 9780367 | 9781593 | 1227 | 2,43 | 2359 |
| chrU | 9793014 | 9794317 | 1304 | 2,49 | 2360 |
| chrU | 9802346 | 9804563 | 2218 | 2,85 | 2361 |
| chrU | 9811409 | 9812795 | 1387 | 2,01 | 2362 |
| chrU | 9832265 | 9833380 | 1116 | 2,17 | 2363 |
| chrU | 9842365 | 9843268 | 904 | 4,08 | 2364 |
| chrU | 9876806 | 9879629 | 2824 | 2,82 | 2365 |
| chrU | 9932255 | 9933428 | 1174 | 3,11 | 2366 |
| chrU | 9934390 | 9935217 | 828 | 3,95 | 2367 |
| chrU | 9944481 | 9945593 | 1113 | 2,78 | 2368 |
| chrU | 9965880 | 9967007 | 1128 | 2,33 | 2369 |
| chrU | 9973668 | 9974830 | 1163 | 2,85 | 2370 |
| chrU | 9983871 | 9984941 | 1071 | 2,49 | 2371 |
| chrU | 10002584 | 10003744 | 1161 | 2,23 | 2372 |
| chrU | 10018129 | 10019374 | 1246 | 2,3 | 2373 |
| chrU | 10032883 | 10034797 | 1915 | 3,39 | 2374 |
| chrUextra | 1186 | 3156 | 1971 | 2,36 | 2375 |
| chrUextra | 5754 | 9861 | 4108 | 2,52 | 2376 |
| chrUextra | 23340 | 28032 | 4693 | 2,59 | 2377 |
| chrUextra | 28268 | 33344 | 5077 | 2,46 | 2378 |
| chrUextra | 45653 | 50859 | 5207 | 3,43 | 2379 |
| chrUextra | 52385 | 53531 | 1147 | 2,23 | 2380 |
| chrUextra | 53724 | 55434 | 1711 | 2,23 | 2381 |
| chrUextra | 56130 | 57545 | 1416 | 2,2 | 2382 |
| chrUextra | 57963 | 60373 | 2411 | 2,17 | 2383 |
| chrUextra | 66358 | 70825 | 4468 | 2,46 | 2384 |
| chrUextra | 71221 | 74091 | 2871 | 2,3 | 2385 |
| chrUextra | 75054 | 80350 | 5297 | 2,39 | 2386 |
| chrUextra | 80953 | 82490 | 1538 | 2,17 | 2387 |
| chrUextra | 83130 | 89025 | 5896 | 2,59 | 2388 |
| chrUextra | 89154 | 90280 | 1127 | 2,33 | 2389 |
| chrUextra | 91050 | 95360 | 4311 | 2,26 | 2390 |
| chrUextra | 95362 | 96853 | 1492 | 2,46 | 2391 |
| chrUextra | 98468 | 102012 | 3545 | 2,33 | 2392 |
| chrUextra | 114992 | 119627 | 4636 | 2,59 | 2393 |
| chrUextra | 122386 | 126204 | 3819 | 2,57 | 2394 |
| chrUextra | 127776 | 131163 | 3388 | 2,59 | 2395 |
| chrUextra | 150891 | 153397 | 2507 | 2,46 | 2396 |
| chrUextra | 157625 | 160237 | 2613 | 2,2 | 2397 |

| | | | | | |
|---|---|---|---|---|---|
| chrUextra | 162047 | 163679 | 1633 | 2,07 | 2398 |
| chrUextra | 164272 | 165956 | 1685 | 2,43 | 2399 |
| chrUextra | 173907 | 176447 | 2541 | 2,2 | 2400 |
| chrUextra | 186559 | 190703 | 4145 | 2,59 | 2401 |
| chrUextra | 193436 | 195094 | 1659 | 1,94 | 2402 |
| chrUextra | 195282 | 199958 | 4677 | 3,19 | 2403 |
| chrUextra | 204116 | 208879 | 4764 | 3,42 | 2404 |
| chrUextra | 225921 | 227912 | 1992 | 2,17 | 2405 |
| chrUextra | 231675 | 232831 | 1157 | 2,26 | 2406 |
| chrUextra | 237821 | 241255 | 3435 | 3,85 | 2407 |
| chrUextra | 241861 | 244704 | 2844 | 2,36 | 2408 |
| chrUextra | 253127 | 254162 | 1036 | 2,46 | 2409 |
| chrUextra | 256451 | 260321 | 3871 | 3,16 | 2410 |
| chrUextra | 260618 | 261927 | 1310 | 2,07 | 2411 |
| chrUextra | 271015 | 274705 | 3691 | 2,87 | 2412 |
| chrUextra | 278232 | 280955 | 2724 | 3,98 | 2413 |
| chrUextra | 295553 | 296804 | 1252 | 2,33 | 2414 |
| chrUextra | 297793 | 301092 | 3300 | 3,88 | 2415 |
| chrUextra | 302371 | 303629 | 1259 | 2,82 | 2416 |
| chrUextra | 304786 | 307310 | 2525 | 2,75 | 2417 |
| chrUextra | 308306 | 310563 | 2258 | 2,26 | 2418 |
| chrUextra | 313996 | 316514 | 2519 | 2,33 | 2419 |
| chrUextra | 316866 | 319749 | 2884 | 2,78 | 2420 |
| chrUextra | 329368 | 331740 | 2373 | 2,88 | 2421 |
| chrUextra | 332391 | 333602 | 1212 | 2,49 | 2422 |
| chrUextra | 333649 | 334781 | 1133 | 2,1 | 2423 |
| chrUextra | 335203 | 337102 | 1900 | 2,17 | 2424 |
| chrUextra | 341579 | 342892 | 1314 | 2,26 | 2425 |
| chrUextra | 359759 | 361998 | 2240 | 2,56 | 2426 |
| chrUextra | 386894 | 389437 | 2544 | 3,39 | 2427 |
| chrUextra | 395982 | 398375 | 2394 | 2,56 | 2428 |
| chrUextra | 398744 | 400221 | 1478 | 2,36 | 2429 |
| chrUextra | 411448 | 413397 | 1950 | 3,22 | 2430 |
| chrUextra | 415588 | 417973 | 2386 | 2,65 | 2431 |
| chrUextra | 422206 | 424233 | 2028 | 3,45 | 2432 |
| chrUextra | 426397 | 427903 | 1507 | 2,36 | 2433 |
| chrUextra | 448170 | 449802 | 1633 | 3,4 | 2434 |
| chrUextra | 450522 | 451608 | 1087 | 2,49 | 2435 |
| chrUextra | 455902 | 457695 | 1794 | 2,79 | 2436 |
| chrUextra | 470992 | 473809 | 2818 | 3,06 | 2437 |
| chrUextra | 476878 | 478682 | 1805 | 3,17 | 2438 |
| chrUextra | 479087 | 480178 | 1092 | 2,78 | 2439 |
| chrUextra | 486771 | 487797 | 1027 | 2,39 | 2440 |
| chrUextra | 491939 | 493353 | 1415 | 3,46 | 2441 |
| chrUextra | 493401 | 494855 | 1455 | 2,82 | 2442 |
| chrUextra | 502724 | 506090 | 3367 | 3,69 | 2443 |
| chrUextra | 508480 | 510168 | 1689 | 3,59 | 2444 |
| chrUextra | 511996 | 513608 | 1613 | 3,12 | 2445 |

| | | | | | |
|---|---|---|---|---|---|
| chrUextra | 515412 | 517230 | 1819 | 2,69 | 2446 |
| chrUextra | 524386 | 526359 | 1974 | 2,64 | 2447 |
| chrUextra | 526701 | 527780 | 1080 | 2,33 | 2448 |
| chrUextra | 540607 | 542302 | 1696 | 2,91 | 2449 |
| chrUextra | 542412 | 543595 | 1184 | 3,16 | 2450 |
| chrUextra | 544057 | 545256 | 1200 | 3,56 | 2451 |
| chrUextra | 545974 | 549259 | 3286 | 3,27 | 2452 |
| chrUextra | 559638 | 561243 | 1606 | 3,18 | 2453 |
| chrUextra | 566496 | 569640 | 3145 | 3,07 | 2454 |
| chrUextra | 604080 | 605219 | 1140 | 3,07 | 2455 |
| chrUextra | 605830 | 607098 | 1269 | 2,36 | 2456 |
| chrUextra | 613519 | 614685 | 1167 | 2,36 | 2457 |
| chrUextra | 641563 | 642614 | 1052 | 2,33 | 2458 |
| chrUextra | 644607 | 645881 | 1275 | 3,17 | 2459 |
| chrUextra | 651983 | 653214 | 1232 | 2,01 | 2460 |
| chrUextra | 668640 | 669929 | 1290 | 2,39 | 2461 |
| chrUextra | 677636 | 678671 | 1036 | 2,49 | 2462 |
| chrUextra | 678729 | 680577 | 1849 | 3,01 | 2463 |
| chrUextra | 683638 | 684808 | 1171 | 2,59 | 2464 |
| chrUextra | 699296 | 701750 | 2455 | 2,72 | 2465 |
| chrUextra | 702367 | 703999 | 1633 | 3,27 | 2466 |
| chrUextra | 706978 | 708074 | 1097 | 2,56 | 2467 |
| chrUextra | 722670 | 724279 | 1610 | 2,26 | 2468 |
| chrUextra | 724430 | 725687 | 1258 | 3,61 | 2469 |
| chrUextra | 740175 | 741574 | 1400 | 3,17 | 2470 |
| chrUextra | 741759 | 742940 | 1182 | 2,33 | 2471 |
| chrUextra | 744394 | 745701 | 1308 | 2,26 | 2472 |
| chrUextra | 745704 | 747577 | 1874 | 2,28 | 2473 |
| chrUextra | 772624 | 773654 | 1031 | 4,82 | 2474 |
| chrUextra | 784432 | 785513 | 1082 | 2,39 | 2475 |
| chrUextra | 788600 | 789765 | 1166 | 2,17 | 2476 |
| chrUextra | 795457 | 796605 | 1149 | 2,69 | 2477 |
| chrUextra | 801220 | 802265 | 1046 | 2,56 | 2478 |
| chrUextra | 807945 | 810069 | 2125 | 2,75 | 2479 |
| chrUextra | 814013 | 815138 | 1126 | 2,82 | 2480 |
| chrUextra | 817724 | 819172 | 1449 | 4,37 | 2481 |
| chrUextra | 828034 | 829116 | 1083 | 2,82 | 2482 |
| chrUextra | 834736 | 835840 | 1105 | 2,72 | 2483 |
| chrUextra | 835879 | 837179 | 1301 | 2,76 | 2484 |
| chrUextra | 847117 | 848299 | 1183 | 3,82 | 2485 |
| chrUextra | 862157 | 863374 | 1218 | 2,23 | 2486 |
| chrUextra | 888744 | 889756 | 1013 | 2,23 | 2487 |
| chrUextra | 907920 | 909853 | 1934 | 2,82 | 2488 |
| chrUextra | 919394 | 921044 | 1651 | 2,78 | 2489 |
| chrUextra | 922151 | 923451 | 1301 | 2,59 | 2490 |
| chrUextra | 930036 | 931978 | 1943 | 2,52 | 2491 |
| chrUextra | 937063 | 938546 | 1484 | 3,01 | 2492 |
| chrUextra | 953358 | 954639 | 1282 | 3,17 | 2493 |

| | | | | | |
|---|---|---|---|---|---|
| chrUextra | 982423 | 984498 | 2076 | 2,41 | 2494 |
| chrUextra | 988765 | 990709 | 1945 | 3,4 | 2495 |
| chrUextra | 1007967 | 1009534 | 1568 | 2,82 | 2496 |
| chrUextra | 1029366 | 1030499 | 1134 | 2,14 | 2497 |
| chrUextra | 1030964 | 1033380 | 2417 | 4,09 | 2498 |
| chrUextra | 1034830 | 1035874 | 1045 | 2,82 | 2499 |
| chrUextra | 1050258 | 1051785 | 1528 | 3,04 | 2500 |
| chrUextra | 1062480 | 1063569 | 1090 | 2,56 | 2501 |
| chrUextra | 1084339 | 1087327 | 2989 | 2,27 | 2502 |
| chrUextra | 1092259 | 1093739 | 1481 | 3,56 | 2503 |
| chrUextra | 1104331 | 1105712 | 1382 | 3,49 | 2504 |
| chrUextra | 1111885 | 1113200 | 1316 | 2,62 | 2505 |
| chrUextra | 1120758 | 1122113 | 1356 | 3,37 | 2506 |
| chrUextra | 1139047 | 1141942 | 2896 | 2,91 | 2507 |
| chrUextra | 1151977 | 1153185 | 1209 | 2,36 | 2508 |
| chrUextra | 1168953 | 1171114 | 2162 | 2,36 | 2509 |
| chrUextra | 1172570 | 1173717 | 1148 | 2,75 | 2510 |
| chrUextra | 1179523 | 1183374 | 3852 | 3,11 | 2511 |
| chrUextra | 1186991 | 1188307 | 1317 | 2,2 | 2512 |
| chrUextra | 1207772 | 1209013 | 1242 | 3,67 | 2513 |
| chrUextra | 1209936 | 1211034 | 1099 | 2,33 | 2514 |
| chrUextra | 1241971 | 1243152 | 1182 | 2,59 | 2515 |
| chrUextra | 1266065 | 1267585 | 1521 | 3,75 | 2516 |
| chrUextra | 1295402 | 1296675 | 1274 | 2,85 | 2517 |
| chrUextra | 1306732 | 1308007 | 1276 | 2,01 | 2518 |
| chrUextra | 1309112 | 1310614 | 1503 | 2,87 | 2519 |
| chrUextra | 1321762 | 1323206 | 1445 | 2,69 | 2520 |
| chrUextra | 1334395 | 1335571 | 1177 | 2,46 | 2521 |
| chrUextra | 1336832 | 1338348 | 1517 | 3,01 | 2522 |
| chrUextra | 1356046 | 1357163 | 1118 | 3,88 | 2523 |
| chrUextra | 1358389 | 1359527 | 1139 | 3,24 | 2524 |
| chrUextra | 1363416 | 1364591 | 1176 | 3,11 | 2525 |
| chrUextra | 1373451 | 1374730 | 1280 | 2,83 | 2526 |
| chrUextra | 1389617 | 1391079 | 1463 | 2,39 | 2527 |
| chrUextra | 1392017 | 1393260 | 1244 | 2,39 | 2528 |
| chrUextra | 1412025 | 1414663 | 2639 | 2,65 | 2529 |
| chrUextra | 1417115 | 1418323 | 1209 | 3,53 | 2530 |
| chrUextra | 1432527 | 1434620 | 2094 | 3,66 | 2531 |
| chrUextra | 1434871 | 1436147 | 1277 | 4,68 | 2532 |
| chrUextra | 1449193 | 1451033 | 1841 | 3,28 | 2533 |
| chrUextra | 1455657 | 1456829 | 1173 | 2,1 | 2534 |
| chrUextra | 1465469 | 1466933 | 1465 | 3,2 | 2535 |
| chrUextra | 1479051 | 1480248 | 1198 | 3,53 | 2536 |
| chrUextra | 1489013 | 1490658 | 1646 | 3,96 | 2537 |
| chrUextra | 1491704 | 1492897 | 1194 | 2,78 | 2538 |
| chrUextra | 1507525 | 1508769 | 1245 | 2,33 | 2539 |
| chrUextra | 1510020 | 1511216 | 1197 | 2,17 | 2540 |
| chrUextra | 1512343 | 1513701 | 1359 | 2,46 | 2541 |

| | | | | | |
|---|---|---|---|---|---|
| chrUextra | 1517258 | 1518304 | 1047 | 2,82 | 2542 |
| chrUextra | 1530852 | 1531920 | 1069 | 2,07 | 2543 |
| chrUextra | 1535765 | 1537209 | 1445 | 2,98 | 2544 |
| chrUextra | 1552222 | 1554156 | 1935 | 3,07 | 2545 |
| chrUextra | 1562729 | 1564139 | 1411 | 3,53 | 2546 |
| chrUextra | 1567629 | 1569026 | 1398 | 2,88 | 2547 |
| chrUextra | 1597846 | 1599535 | 1690 | 3,57 | 2548 |
| chrUextra | 1607742 | 1608928 | 1187 | 3,3 | 2549 |
| chrUextra | 1631185 | 1632391 | 1207 | 4,01 | 2550 |
| chrUextra | 1643165 | 1644376 | 1212 | 2,36 | 2551 |
| chrUextra | 1662367 | 1663715 | 1349 | 2,3 | 2552 |
| chrUextra | 1666094 | 1669532 | 3439 | 3,14 | 2553 |
| chrUextra | 1669729 | 1670961 | 1233 | 2,69 | 2554 |
| chrUextra | 1673809 | 1675848 | 2040 | 2,82 | 2555 |
| chrUextra | 1690147 | 1691424 | 1278 | 3,82 | 2556 |
| chrUextra | 1693834 | 1694920 | 1087 | 2,59 | 2557 |
| chrUextra | 1700895 | 1702001 | 1107 | 2,65 | 2558 |
| chrUextra | 1709008 | 1710863 | 1856 | 3,8 | 2559 |
| chrUextra | 1712982 | 1714149 | 1168 | 3,07 | 2560 |
| chrUextra | 1717675 | 1718954 | 1280 | 2,75 | 2561 |
| chrUextra | 1738019 | 1739268 | 1250 | 2,39 | 2562 |
| chrUextra | 1792582 | 1794333 | 1752 | 3,42 | 2563 |
| chrUextra | 1796607 | 1797624 | 1018 | 2,56 | 2564 |
| chrUextra | 1804583 | 1805897 | 1315 | 2,29 | 2565 |
| chrUextra | 1811661 | 1813134 | 1474 | 2,06 | 2566 |
| chrUextra | 1817650 | 1818837 | 1188 | 2,07 | 2567 |
| chrUextra | 1831872 | 1833102 | 1231 | 2,17 | 2568 |
| chrUextra | 1841174 | 1842627 | 1454 | 2,33 | 2569 |
| chrUextra | 1860156 | 1861292 | 1137 | 2,43 | 2570 |
| chrUextra | 1869685 | 1870817 | 1133 | 2,33 | 2571 |
| chrUextra | 1898920 | 1900308 | 1389 | 3,46 | 2572 |
| chrUextra | 1902302 | 1903839 | 1538 | 3,53 | 2573 |
| chrUextra | 1917661 | 1919191 | 1531 | 2,98 | 2574 |
| chrUextra | 1924646 | 1926244 | 1599 | 2,79 | 2575 |
| chrUextra | 1936618 | 1937673 | 1056 | 2,07 | 2576 |
| chrUextra | 1938779 | 1940157 | 1379 | 2,3 | 2577 |
| chrUextra | 1953108 | 1954096 | 989 | 3,07 | 2578 |
| chrUextra | 1966750 | 1968210 | 1461 | 2,64 | 2579 |
| chrUextra | 2013404 | 2014761 | 1358 | 3,03 | 2580 |
| chrUextra | 2028496 | 2029772 | 1277 | 2,39 | 2581 |
| chrUextra | 2055033 | 2056490 | 1458 | 3,4 | 2582 |
| chrUextra | 2057258 | 2058743 | 1486 | 2,87 | 2583 |
| chrUextra | 2062172 | 2063299 | 1128 | 3,01 | 2584 |
| chrUextra | 2065598 | 2066736 | 1139 | 2,1 | 2585 |
| chrUextra | 2069200 | 2070309 | 1110 | 3,1 | 2586 |
| chrUextra | 2077953 | 2079225 | 1273 | 3,86 | 2587 |
| chrUextra | 2087361 | 2088626 | 1266 | 2,32 | 2588 |
| chrUextra | 2089559 | 2091964 | 2406 | 2,54 | 2589 |

| | | | | | |
|---|---|---|---|---|---|
| chrUextra | 2095414 | 2096756 | 1343 | 2,37 | 2590 |
| chrUextra | 2109250 | 2110431 | 1182 | 2,43 | 2591 |
| chrUextra | 2121804 | 2123118 | 1315 | 2,36 | 2592 |
| chrUextra | 2124102 | 2125343 | 1242 | 2,72 | 2593 |
| chrUextra | 2149283 | 2151922 | 2640 | 2,88 | 2594 |
| chrUextra | 2163062 | 2165199 | 2138 | 2,46 | 2595 |
| chrUextra | 2166485 | 2168987 | 2503 | 3,62 | 2596 |
| chrUextra | 2172129 | 2173403 | 1275 | 3,72 | 2597 |
| chrUextra | 2173857 | 2175619 | 1763 | 2,39 | 2598 |
| chrUextra | 2179991 | 2182553 | 2563 | 4,14 | 2599 |
| chrUextra | 2205245 | 2206449 | 1205 | 2,3 | 2600 |
| chrUextra | 2214435 | 2215641 | 1207 | 2,85 | 2601 |
| chrUextra | 2217943 | 2219042 | 1100 | 3,04 | 2602 |
| chrUextra | 2230118 | 2231493 | 1376 | 3,1 | 2603 |
| chrUextra | 2238149 | 2239505 | 1357 | 3,37 | 2604 |
| chrUextra | 2245014 | 2246194 | 1181 | 2,56 | 2605 |
| chrUextra | 2262021 | 2263184 | 1164 | 2,04 | 2606 |
| chrUextra | 2267616 | 2270099 | 2484 | 3,17 | 2607 |
| chrUextra | 2285701 | 2287979 | 2279 | 2,69 | 2608 |
| chrUextra | 2302638 | 2304938 | 2301 | 2,98 | 2609 |
| chrUextra | 2312735 | 2313804 | 1070 | 3,37 | 2610 |
| chrUextra | 2344538 | 2346730 | 2193 | 2,48 | 2611 |
| chrUextra | 2362305 | 2363584 | 1280 | 2,26 | 2612 |
| chrUextra | 2375138 | 2377042 | 1905 | 2,27 | 2613 |
| chrUextra | 2393763 | 2394808 | 1046 | 2,46 | 2614 |
| chrUextra | 2400377 | 2401867 | 1491 | 3,59 | 2615 |
| chrUextra | 2403865 | 2405037 | 1173 | 2,01 | 2616 |
| chrUextra | 2409314 | 2410456 | 1143 | 2,04 | 2617 |
| chrUextra | 2449458 | 2450630 | 1173 | 2,52 | 2618 |
| chrUextra | 2468743 | 2471936 | 3194 | 2,56 | 2619 |
| chrUextra | 2473894 | 2475089 | 1196 | 2,2 | 2620 |
| chrUextra | 2487122 | 2488411 | 1290 | 3,46 | 2621 |
| chrUextra | 2492738 | 2493824 | 1087 | 3,24 | 2622 |
| chrUextra | 2494833 | 2496375 | 1543 | 3,29 | 2623 |
| chrUextra | 2512963 | 2513882 | 920 | 3,74 | 2624 |
| chrUextra | 2535887 | 2537249 | 1363 | 2,52 | 2625 |
| chrUextra | 2539282 | 2540456 | 1175 | 2,71 | 2626 |
| chrUextra | 2541487 | 2542625 | 1139 | 2,36 | 2627 |
| chrUextra | 2556945 | 2558180 | 1236 | 2,56 | 2628 |
| chrUextra | 2577906 | 2579052 | 1147 | 2,39 | 2629 |
| chrUextra | 2594341 | 2595776 | 1436 | 3,33 | 2630 |
| chrUextra | 2601080 | 2602182 | 1103 | 2,36 | 2631 |
| chrUextra | 2609815 | 2613060 | 3246 | 3,04 | 2632 |
| chrUextra | 2644898 | 2645553 | 656 | 4,66 | 2633 |
| chrUextra | 2645982 | 2648333 | 2352 | 2,46 | 2634 |
| chrUextra | 2652651 | 2654495 | 1845 | 2,17 | 2635 |
| chrUextra | 2672450 | 2673502 | 1053 | 2,59 | 2636 |
| chrUextra | 2675590 | 2676877 | 1288 | 2,46 | 2637 |

| | | | | | |
|---|---|---|---|---|---|
| chrUextra | 2684009 | 2685260 | 1252 | 9,61 | 2638 |
| chrUextra | 2696399 | 2697526 | 1128 | 2,39 | 2639 |
| chrUextra | 2714277 | 2715645 | 1369 | 2,36 | 2640 |
| chrUextra | 2720591 | 2721650 | 1060 | 2,2 | 2641 |
| chrUextra | 2724638 | 2726411 | 1774 | 3,66 | 2642 |
| chrUextra | 2732512 | 2733766 | 1255 | 2,12 | 2643 |
| chrUextra | 2736974 | 2738014 | 1041 | 2,56 | 2644 |
| chrUextra | 2746958 | 2749086 | 2129 | 3,04 | 2645 |
| chrUextra | 2771047 | 2771665 | 619 | 4,47 | 2646 |
| chrUextra | 2778382 | 2779669 | 1288 | 2,69 | 2647 |
| chrUextra | 2803289 | 2807552 | 4264 | 3,17 | 2648 |
| chrUextra | 2858975 | 2860292 | 1318 | 2,99 | 2649 |
| chrUextra | 2863222 | 2864240 | 1019 | 2,49 | 2650 |
| chrUextra | 2880657 | 2881931 | 1275 | 2,3 | 2651 |
| chrUextra | 2885078 | 2886169 | 1092 | 2,17 | 2652 |
| chrUextra | 2889317 | 2890496 | 1180 | 2,56 | 2653 |
| chrUextra | 2900608 | 2901286 | 679 | 4,53 | 2654 |
| chrUextra | 2904675 | 2907080 | 2406 | 4,07 | 2655 |
| chrUextra | 2907973 | 2909053 | 1081 | 2,65 | 2656 |
| chrUextra | 2911968 | 2913543 | 1576 | 3,36 | 2657 |
| chrUextra | 2921883 | 2924200 | 2318 | 2,96 | 2658 |
| chrUextra | 2932724 | 2934012 | 1289 | 2,59 | 2659 |
| chrUextra | 2964511 | 2965217 | 707 | 3,69 | 2660 |
| chrUextra | 2999438 | 3000523 | 1086 | 4,06 | 2661 |
| chrUextra | 3001431 | 3002921 | 1491 | 3,86 | 2662 |
| chrUextra | 3005336 | 3006448 | 1113 | 2,36 | 2663 |
| chrUextra | 3022543 | 3023531 | 989 | 5,36 | 2664 |
| chrUextra | 3044075 | 3045144 | 1070 | 2,46 | 2665 |
| chrUextra | 3047211 | 3048346 | 1136 | 3,2 | 2666 |
| chrUextra | 3073090 | 3074131 | 1042 | 2,75 | 2667 |
| chrUextra | 3074369 | 3076491 | 2123 | 3,14 | 2668 |
| chrUextra | 3079352 | 3081610 | 2259 | 5,91 | 2669 |
| chrUextra | 3089318 | 3090793 | 1476 | 3,46 | 2670 |
| chrUextra | 3094377 | 3095160 | 784 | 5,02 | 2671 |
| chrUextra | 3096704 | 3098134 | 1431 | 2,98 | 2672 |
| chrUextra | 3103349 | 3104493 | 1145 | 2,3 | 2673 |
| chrUextra | 3135859 | 3136992 | 1134 | 5,9 | 2674 |
| chrUextra | 3174078 | 3175159 | 1082 | 3,11 | 2675 |
| chrUextra | 3188079 | 3189216 | 1138 | 2,2 | 2676 |
| chrUextra | 3190332 | 3192040 | 1709 | 2,07 | 2677 |
| chrUextra | 3204572 | 3206511 | 1940 | 6,34 | 2678 |
| chrUextra | 3212028 | 3213214 | 1187 | 5,74 | 2679 |
| chrUextra | 3213840 | 3215002 | 1163 | 2,85 | 2680 |
| chrUextra | 3217128 | 3219333 | 2206 | 2,62 | 2681 |
| chrUextra | 3224468 | 3225748 | 1281 | 2,56 | 2682 |
| chrUextra | 3229889 | 3231060 | 1172 | 2,17 | 2683 |
| chrUextra | 3232148 | 3233265 | 1118 | 2,82 | 2684 |
| chrUextra | 3243910 | 3245927 | 2018 | 2,85 | 2685 |

| | | | | | |
|---|---|---|---|---|---|
| chrUextra | 3251234 | 3252446 | 1213 | 2,62 | 2686 |
| chrUextra | 3256622 | 3257835 | 1214 | 2,75 | 2687 |
| chrUextra | 3293981 | 3295188 | 1208 | 2,1 | 2688 |
| chrUextra | 3320615 | 3321859 | 1245 | 2,3 | 2689 |
| chrUextra | 3352538 | 3353826 | 1289 | 2,69 | 2690 |
| chrUextra | 3364409 | 3366413 | 2005 | 2,36 | 2691 |
| chrUextra | 3370959 | 3371885 | 927 | 4,59 | 2692 |
| chrUextra | 3387701 | 3388887 | 1187 | 2,2 | 2693 |
| chrUextra | 3397004 | 3398436 | 1433 | 2,62 | 2694 |
| chrUextra | 3399348 | 3400737 | 1390 | 3,73 | 2695 |
| chrUextra | 3414828 | 3415748 | 921 | 4,11 | 2696 |
| chrUextra | 3430154 | 3430923 | 770 | 4,94 | 2697 |
| chrUextra | 3431294 | 3432404 | 1111 | 2,26 | 2698 |
| chrUextra | 3437469 | 3438999 | 1531 | 3,32 | 2699 |
| chrUextra | 3446084 | 3447326 | 1243 | 2,3 | 2700 |
| chrUextra | 3452437 | 3453595 | 1159 | 2,82 | 2701 |
| chrUextra | 3456765 | 3458060 | 1296 | 3,37 | 2702 |
| chrUextra | 3460896 | 3462157 | 1262 | 2,72 | 2703 |
| chrUextra | 3478908 | 3479972 | 1065 | 2,36 | 2704 |
| chrUextra | 3481031 | 3482211 | 1181 | 2,17 | 2705 |
| chrUextra | 3482355 | 3484331 | 1977 | 2,39 | 2706 |
| chrUextra | 3508494 | 3510749 | 2256 | 2,56 | 2707 |
| chrUextra | 3533875 | 3534997 | 1123 | 3,04 | 2708 |
| chrUextra | 3550813 | 3551985 | 1173 | 2,75 | 2709 |
| chrUextra | 3577358 | 3578464 | 1107 | 2,69 | 2710 |
| chrUextra | 3579289 | 3580304 | 1016 | 2,78 | 2711 |
| chrUextra | 3597192 | 3598382 | 1191 | 2,52 | 2712 |
| chrUextra | 3603448 | 3604724 | 1277 | 2,3 | 2713 |
| chrUextra | 3609691 | 3610899 | 1209 | 2,43 | 2714 |
| chrUextra | 3616472 | 3618388 | 1917 | 2,46 | 2715 |
| chrUextra | 3623448 | 3624561 | 1114 | 2,43 | 2716 |
| chrUextra | 3633205 | 3634164 | 960 | 4,79 | 2717 |
| chrUextra | 3635034 | 3636151 | 1118 | 2,91 | 2718 |
| chrUextra | 3644476 | 3645723 | 1248 | 2,93 | 2719 |
| chrUextra | 3649790 | 3651016 | 1227 | 2,62 | 2720 |
| chrUextra | 3651877 | 3653025 | 1149 | 2,2 | 2721 |
| chrUextra | 3655048 | 3656236 | 1189 | 2,75 | 2722 |
| chrUextra | 3666598 | 3667838 | 1241 | 2,43 | 2723 |
| chrUextra | 3668080 | 3669583 | 1504 | 2,17 | 2724 |
| chrUextra | 3706450 | 3707706 | 1257 | 2,92 | 2725 |
| chrUextra | 3710577 | 3712405 | 1829 | 2,58 | 2726 |
| chrUextra | 3758704 | 3760025 | 1322 | 2,46 | 2727 |
| chrUextra | 3770234 | 3771333 | 1100 | 2,78 | 2728 |
| chrUextra | 3771391 | 3772571 | 1181 | 2,98 | 2729 |
| chrUextra | 3794461 | 3795547 | 1087 | 2,46 | 2730 |
| chrUextra | 3798627 | 3800346 | 1720 | 2,65 | 2731 |
| chrUextra | 3809201 | 3809997 | 797 | 8,83 | 2732 |
| chrUextra | 3812077 | 3813168 | 1092 | 2,3 | 2733 |

| | | | | | |
|---|---|---|---|---|---|
| chrUextra | 3849641 | 3850780 | 1140 | 2,52 | 2734 |
| chrUextra | 3852814 | 3854002 | 1189 | 2,59 | 2735 |
| chrUextra | 3858018 | 3859141 | 1124 | 2,33 | 2736 |
| chrUextra | 3864278 | 3867069 | 2792 | 3,82 | 2737 |
| chrUextra | 3892334 | 3893573 | 1240 | 2,23 | 2738 |
| chrUextra | 3900754 | 3903591 | 2838 | 3,6 | 2739 |
| chrUextra | 3913138 | 3914554 | 1417 | 2,82 | 2740 |
| chrUextra | 3932833 | 3934059 | 1227 | 3,49 | 2741 |
| chrUextra | 3934841 | 3936774 | 1934 | 3,33 | 2742 |
| chrUextra | 3940372 | 3941415 | 1044 | 2,36 | 2743 |
| chrUextra | 3949591 | 3950730 | 1140 | 2,36 | 2744 |
| chrUextra | 3956881 | 3957837 | 957 | 2,46 | 2745 |
| chrUextra | 3970260 | 3971351 | 1092 | 2,04 | 2746 |
| chrUextra | 3984004 | 3986617 | 2614 | 2,23 | 2747 |
| chrUextra | 3989865 | 3991234 | 1370 | 2,95 | 2748 |
| chrUextra | 4009594 | 4010734 | 1141 | 2,72 | 2749 |
| chrUextra | 4013719 | 4015044 | 1326 | 2,75 | 2750 |
| chrUextra | 4032367 | 4033592 | 1226 | 2,22 | 2751 |
| chrUextra | 4036555 | 4037691 | 1137 | 2,33 | 2752 |
| chrUextra | 4100533 | 4101581 | 1049 | 2,17 | 2753 |
| chrUextra | 4101692 | 4103809 | 2118 | 2,49 | 2754 |
| chrUextra | 4110822 | 4112041 | 1220 | 2,82 | 2755 |
| chrUextra | 4122243 | 4123327 | 1085 | 2,39 | 2756 |
| chrUextra | 4128361 | 4129397 | 1037 | 2,43 | 2757 |
| chrUextra | 4139933 | 4141682 | 1750 | 2,2 | 2758 |
| chrUextra | 4158337 | 4159359 | 1023 | 2,23 | 2759 |
| chrUextra | 4180941 | 4182028 | 1088 | 2,33 | 2760 |
| chrUextra | 4189075 | 4190219 | 1145 | 2,98 | 2761 |
| chrUextra | 4196189 | 4197389 | 1201 | 3,2 | 2762 |
| chrUextra | 4197589 | 4199384 | 1796 | 2,43 | 2763 |
| chrUextra | 4205591 | 4206665 | 1075 | 2,23 | 2764 |
| chrUextra | 4209899 | 4211623 | 1725 | 2,2 | 2765 |
| chrUextra | 4239374 | 4240615 | 1242 | 4,12 | 2766 |
| chrUextra | 4242443 | 4245631 | 3189 | 2,17 | 2767 |
| chrUextra | 4246526 | 4248018 | 1493 | 3,57 | 2768 |
| chrUextra | 4249820 | 4250833 | 1014 | 2,46 | 2769 |
| chrUextra | 4274156 | 4276509 | 2354 | 2,59 | 2770 |
| chrUextra | 4283596 | 4285328 | 1733 | 2,98 | 2771 |
| chrUextra | 4285530 | 4287711 | 2182 | 2,07 | 2772 |
| chrUextra | 4302978 | 4304793 | 1816 | 2,3 | 2773 |
| chrUextra | 4306059 | 4310166 | 4108 | 3,23 | 2774 |
| chrUextra | 4322447 | 4323665 | 1219 | 2,62 | 2775 |
| chrUextra | 4325534 | 4326677 | 1144 | 2,59 | 2776 |
| chrUextra | 4339774 | 4340810 | 1037 | 2,62 | 2777 |
| chrUextra | 4348849 | 4349933 | 1085 | 3,11 | 2778 |
| chrUextra | 4351059 | 4352099 | 1041 | 2,46 | 2779 |
| chrUextra | 4359263 | 4360452 | 1190 | 2,22 | 2780 |
| chrUextra | 4400927 | 4402084 | 1158 | 2,46 | 2781 |

| | | | | | |
|---|---|---|---|---|---|
| chrUextra | 4405247 | 4406387 | 1141 | 2,59 | 2782 |
| chrUextra | 4435646 | 4436822 | 1177 | 2,39 | 2783 |
| chrUextra | 4437801 | 4438939 | 1139 | 2,5 | 2784 |
| chrUextra | 4471320 | 4473520 | 2201 | 2,78 | 2785 |
| chrUextra | 4475505 | 4477518 | 2014 | 2,39 | 2786 |
| chrUextra | 4481459 | 4482723 | 1265 | 2,72 | 2787 |
| chrUextra | 4488760 | 4489806 | 1047 | 2,36 | 2788 |
| chrUextra | 4516067 | 4517113 | 1047 | 2,43 | 2789 |
| chrUextra | 4524101 | 4525264 | 1164 | 2,23 | 2790 |
| chrUextra | 4528995 | 4530332 | 1338 | 3,83 | 2791 |
| chrUextra | 4558611 | 4559640 | 1030 | 2,59 | 2792 |
| chrUextra | 4563604 | 4564720 | 1117 | 2,72 | 2793 |
| chrUextra | 4567648 | 4568841 | 1194 | 2,36 | 2794 |
| chrUextra | 4595857 | 4597263 | 1407 | 3,43 | 2795 |
| chrUextra | 4600951 | 4602241 | 1291 | 2,62 | 2796 |
| chrUextra | 4609171 | 4610329 | 1159 | 2,23 | 2797 |
| chrUextra | 4610459 | 4613778 | 3320 | 4,59 | 2798 |
| chrUextra | 4621297 | 4622401 | 1105 | 2,2 | 2799 |
| chrUextra | 4633367 | 4634458 | 1092 | 2,56 | 2800 |
| chrUextra | 4636393 | 4637597 | 1205 | 3,2 | 2801 |
| chrUextra | 4647690 | 4649228 | 1539 | 2,43 | 2802 |
| chrUextra | 4649537 | 4650750 | 1214 | 2,59 | 2803 |
| chrUextra | 4654537 | 4655937 | 1401 | 2,91 | 2804 |
| chrUextra | 4664713 | 4665788 | 1076 | 2,49 | 2805 |
| chrUextra | 4674732 | 4676010 | 1279 | 2,98 | 2806 |
| chrUextra | 4677857 | 4678936 | 1080 | 2,72 | 2807 |
| chrUextra | 4680786 | 4682576 | 1791 | 2,52 | 2808 |
| chrUextra | 4686958 | 4690800 | 3843 | 3,17 | 2809 |
| chrUextra | 4694987 | 4696095 | 1109 | 2,43 | 2810 |
| chrUextra | 4723063 | 4724111 | 1049 | 2,49 | 2811 |
| chrUextra | 4724307 | 4725341 | 1035 | 2,56 | 2812 |
| chrUextra | 4729425 | 4730526 | 1102 | 4,55 | 2813 |
| chrUextra | 4735134 | 4736350 | 1217 | 2,39 | 2814 |
| chrUextra | 4741162 | 4742224 | 1063 | 2,28 | 2815 |
| chrUextra | 4761279 | 4762457 | 1179 | 2,6 | 2816 |
| chrUextra | 4766281 | 4767493 | 1213 | 2,33 | 2817 |
| chrUextra | 4775304 | 4777444 | 2141 | 2,52 | 2818 |
| chrUextra | 4782341 | 4784305 | 1965 | 2,82 | 2819 |
| chrUextra | 4794370 | 4796584 | 2215 | 2,29 | 2820 |
| chrUextra | 4804501 | 4806565 | 2065 | 2,42 | 2821 |
| chrUextra | 4826502 | 4827654 | 1153 | 2,33 | 2822 |
| chrUextra | 4835532 | 4837559 | 2028 | 2,46 | 2823 |
| chrUextra | 4843533 | 4844685 | 1153 | 2,91 | 2824 |
| chrUextra | 4862503 | 4863555 | 1053 | 2,59 | 2825 |
| chrUextra | 4876653 | 4879410 | 2758 | 2,73 | 2826 |
| chrUextra | 4887505 | 4888663 | 1159 | 2,94 | 2827 |
| chrUextra | 4913606 | 4916612 | 3007 | 2,55 | 2828 |
| chrUextra | 4922529 | 4924494 | 1966 | 2,2 | 2829 |

| | | | | | |
|---|---|---|---|---|---|
| chrUextra | 4934471 | 4935648 | 1178 | 2,43 | 2830 |
| chrUextra | 4938453 | 4939696 | 1244 | 2,3 | 2831 |
| chrUextra | 4959504 | 4961584 | 2081 | 2,88 | 2832 |
| chrUextra | 4979441 | 4980861 | 1421 | 2,46 | 2833 |
| chrUextra | 4985349 | 4986538 | 1190 | 2,91 | 2834 |
| chrUextra | 4994492 | 4996122 | 1631 | 2,43 | 2835 |
| chrUextra | 5000495 | 5002151 | 1657 | 2,46 | 2836 |
| chrUextra | 5006496 | 5008362 | 1867 | 2,46 | 2837 |
| chrUextra | 5008391 | 5010442 | 2052 | 2,36 | 2838 |
| chrUextra | 5018155 | 5019398 | 1244 | 2,94 | 2839 |
| chrUextra | 5047059 | 5048246 | 1188 | 3,89 | 2840 |
| chrUextra | 5049174 | 5050262 | 1089 | 2,33 | 2841 |
| chrUextra | 5064145 | 5065883 | 1739 | 5,47 | 2842 |
| chrUextra | 5086846 | 5088321 | 1476 | 3,56 | 2843 |
| chrUextra | 5097720 | 5098743 | 1024 | 2,1 | 2844 |
| chrUextra | 5109372 | 5110372 | 1001 | 6,5 | 2845 |
| chrUextra | 5115516 | 5116692 | 1177 | 2,1 | 2846 |
| chrUextra | 5117007 | 5117744 | 738 | 4,98 | 2847 |
| chrUextra | 5118556 | 5119766 | 1211 | 2,22 | 2848 |
| chrUextra | 5122504 | 5123587 | 1084 | 2,17 | 2849 |
| chrUextra | 5142128 | 5143245 | 1118 | 5,84 | 2850 |
| chrUextra | 5164157 | 5165436 | 1280 | 3,56 | 2851 |
| chrUextra | 5174000 | 5175218 | 1219 | 2,39 | 2852 |
| chrUextra | 5193753 | 5194845 | 1093 | 2,88 | 2853 |
| chrUextra | 5208498 | 5209743 | 1246 | 2,2 | 2854 |
| chrUextra | 5216498 | 5217722 | 1225 | 2,49 | 2855 |
| chrUextra | 5223660 | 5224906 | 1247 | 11,71 | 2856 |
| chrUextra | 5254086 | 5255818 | 1733 | 2,9 | 2857 |
| chrUextra | 5263780 | 5265116 | 1337 | 2,75 | 2858 |
| chrUextra | 5267982 | 5269463 | 1482 | 2,23 | 2859 |
| chrUextra | 5288622 | 5289639 | 1018 | 2,33 | 2860 |
| chrUextra | 5297352 | 5299124 | 1773 | 3,83 | 2861 |
| chrUextra | 5306174 | 5307248 | 1075 | 2,49 | 2862 |
| chrUextra | 5315831 | 5317394 | 1564 | 3,58 | 2863 |
| chrUextra | 5324929 | 5326019 | 1091 | 2,46 | 2864 |
| chrUextra | 5326655 | 5327602 | 948 | 3,42 | 2865 |
| chrUextra | 5342562 | 5343718 | 1157 | 2,65 | 2866 |
| chrUextra | 5348298 | 5350564 | 2267 | 3,17 | 2867 |
| chrUextra | 5357389 | 5358460 | 1072 | 2,26 | 2868 |
| chrUextra | 5373899 | 5375075 | 1177 | 3,96 | 2869 |
| chrUextra | 5422222 | 5423216 | 995 | 2,75 | 2870 |
| chrUextra | 5424029 | 5424889 | 861 | 3,72 | 2871 |
| chrUextra | 5431870 | 5432895 | 1026 | 2,69 | 2872 |
| chrUextra | 5442619 | 5443795 | 1177 | 2,43 | 2873 |
| chrUextra | 5452494 | 5453555 | 1062 | 2,59 | 2874 |
| chrUextra | 5462224 | 5463350 | 1127 | 2,98 | 2875 |
| chrUextra | 5477973 | 5478989 | 1017 | 2,39 | 2876 |
| chrUextra | 5480728 | 5482180 | 1453 | 2,43 | 2877 |

| | | | | | |
|---|---|---|---|---|---|
| chrUextra | 5490736 | 5492772 | 2037 | 2,46 | 2878 |
| chrUextra | 5506277 | 5508131 | 1855 | 2,43 | 2879 |
| chrUextra | 5517039 | 5518206 | 1168 | 2,72 | 2880 |
| chrUextra | 5553196 | 5554234 | 1039 | 2,3 | 2881 |
| chrUextra | 5558110 | 5559187 | 1078 | 2,52 | 2882 |
| chrUextra | 5570748 | 5571793 | 1046 | 2,33 | 2883 |
| chrUextra | 5579522 | 5580317 | 796 | 6,73 | 2884 |
| chrUextra | 5588317 | 5589468 | 1152 | 2,56 | 2885 |
| chrUextra | 5643962 | 5645837 | 1876 | 2,26 | 2886 |
| chrUextra | 5647648 | 5649262 | 1615 | 2,2 | 2887 |
| chrUextra | 5653498 | 5655581 | 2084 | 2,43 | 2888 |
| chrUextra | 5662273 | 5664607 | 2335 | 3,24 | 2889 |
| chrUextra | 5674828 | 5676779 | 1952 | 2,78 | 2890 |
| chrUextra | 5677874 | 5678952 | 1079 | 2,3 | 2891 |
| chrUextra | 5684663 | 5685801 | 1139 | 2,75 | 2892 |
| chrUextra | 5686673 | 5687698 | 1026 | 2,04 | 2893 |
| chrUextra | 5707942 | 5709151 | 1210 | 2,77 | 2894 |
| chrUextra | 5735241 | 5736192 | 952 | 2,43 | 2895 |
| chrUextra | 5740060 | 5741115 | 1056 | 2,98 | 2896 |
| chrUextra | 5747537 | 5748595 | 1059 | 4,69 | 2897 |
| chrUextra | 5770902 | 5772103 | 1202 | 2,39 | 2898 |
| chrUextra | 5798092 | 5799156 | 1065 | 2,33 | 2899 |
| chrUextra | 5810638 | 5811772 | 1135 | 2,49 | 2900 |
| chrUextra | 5822252 | 5823293 | 1042 | 2,62 | 2901 |
| chrUextra | 5828610 | 5829399 | 790 | 5,34 | 2902 |
| chrUextra | 5831799 | 5832541 | 743 | 3,3 | 2903 |
| chrUextra | 5834897 | 5836933 | 2037 | 3,01 | 2904 |
| chrUextra | 5840697 | 5842757 | 2061 | 2,75 | 2905 |
| chrUextra | 5850231 | 5851334 | 1104 | 2,43 | 2906 |
| chrUextra | 5852151 | 5853372 | 1222 | 2,69 | 2907 |
| chrUextra | 5857983 | 5860143 | 2161 | 2,69 | 2908 |
| chrUextra | 5860735 | 5862114 | 1380 | 3,78 | 2909 |
| chrUextra | 5882018 | 5883224 | 1207 | 2,91 | 2910 |
| chrUextra | 5892665 | 5893937 | 1273 | 3,17 | 2911 |
| chrUextra | 5897270 | 5899270 | 2001 | 2,98 | 2912 |
| chrUextra | 5902306 | 5904271 | 1966 | 3,09 | 2913 |
| chrUextra | 5913100 | 5914606 | 1507 | 2,36 | 2914 |
| chrUextra | 5919721 | 5920873 | 1153 | 2,75 | 2915 |
| chrUextra | 5936961 | 5937967 | 1007 | 3,85 | 2916 |
| chrUextra | 5944660 | 5945778 | 1119 | 2,39 | 2917 |
| chrUextra | 5955286 | 5957470 | 2185 | 3,44 | 2918 |
| chrUextra | 5981336 | 5983210 | 1875 | 2,14 | 2919 |
| chrUextra | 5984151 | 5985216 | 1066 | 2,36 | 2920 |
| chrUextra | 5987905 | 5989077 | 1173 | 2,32 | 2921 |
| chrUextra | 6004242 | 6005321 | 1080 | 2,14 | 2922 |
| chrUextra | 6043597 | 6045561 | 1965 | 2,14 | 2923 |
| chrUextra | 6051280 | 6052318 | 1039 | 2,36 | 2924 |
| chrUextra | 6065687 | 6067292 | 1606 | 2,59 | 2925 |

| | | | | | |
|---|---|---|---|---|---|
| chrUextra | 6078928 | 6080075 | 1148 | 2,46 | 2926 |
| chrUextra | 6081869 | 6082924 | 1056 | 2,85 | 2927 |
| chrUextra | 6088567 | 6089611 | 1045 | 2,46 | 2928 |
| chrUextra | 6129688 | 6130216 | 529 | 4,84 | 2929 |
| chrUextra | 6146878 | 6147945 | 1068 | 2,14 | 2930 |
| chrUextra | 6151440 | 6152489 | 1050 | 2,89 | 2931 |
| chrUextra | 6157670 | 6158733 | 1064 | 4,54 | 2932 |
| chrUextra | 6165968 | 6166764 | 797 | 3,38 | 2933 |
| chrUextra | 6167921 | 6169106 | 1186 | 2,44 | 2934 |
| chrUextra | 6176584 | 6178479 | 1896 | 2,69 | 2935 |
| chrUextra | 6202264 | 6203384 | 1121 | 2,31 | 2936 |
| chrUextra | 6212675 | 6213767 | 1093 | 2,1 | 2937 |
| chrUextra | 6217519 | 6218626 | 1108 | 2,46 | 2938 |
| chrUextra | 6224103 | 6225054 | 952 | 2,46 | 2939 |
| chrUextra | 6235533 | 6236546 | 1014 | 2,33 | 2940 |
| chrUextra | 6238323 | 6239539 | 1217 | 2,91 | 2941 |
| chrUextra | 6246043 | 6247161 | 1119 | 2,14 | 2942 |
| chrUextra | 6248564 | 6250089 | 1526 | 3,71 | 2943 |
| chrUextra | 6281381 | 6283393 | 2013 | 3,62 | 2944 |
| chrUextra | 6294491 | 6295556 | 1066 | 2,36 | 2945 |
| chrUextra | 6345808 | 6347130 | 1323 | 9,31 | 2946 |
| chrUextra | 6361879 | 6362956 | 1078 | 2,39 | 2947 |
| chrUextra | 6365575 | 6366617 | 1043 | 3,04 | 2948 |
| chrUextra | 6394033 | 6395056 | 1024 | 2,33 | 2949 |
| chrUextra | 6408064 | 6409345 | 1282 | 2,69 | 2950 |
| chrUextra | 6418669 | 6419812 | 1144 | 3,01 | 2951 |
| chrUextra | 6420566 | 6422695 | 2130 | 6,36 | 2952 |
| chrUextra | 6466938 | 6467957 | 1020 | 2,52 | 2953 |
| chrUextra | 6493376 | 6494428 | 1053 | 2,36 | 2954 |
| chrUextra | 6500756 | 6502194 | 1439 | 3,1 | 2955 |
| chrUextra | 6519753 | 6520831 | 1079 | 2,62 | 2956 |
| chrUextra | 6541437 | 6543313 | 1877 | 2,26 | 2957 |
| chrUextra | 6549990 | 6550986 | 997 | 7,06 | 2958 |
| chrUextra | 6576430 | 6577993 | 1564 | 3,6 | 2959 |
| chrUextra | 6584561 | 6585825 | 1265 | 2,41 | 2960 |
| chrUextra | 6588145 | 6589254 | 1110 | 2,99 | 2961 |
| chrUextra | 6609112 | 6610149 | 1038 | 2,2 | 2962 |
| chrUextra | 6630679 | 6631769 | 1091 | 2,33 | 2963 |
| chrUextra | 6638961 | 6639884 | 924 | 3,3 | 2964 |
| chrUextra | 6647632 | 6648699 | 1068 | 2,46 | 2965 |
| chrUextra | 6655133 | 6656320 | 1188 | 2,49 | 2966 |
| chrUextra | 6662201 | 6663487 | 1287 | 4,34 | 2967 |
| chrUextra | 6684134 | 6686109 | 1976 | 2,65 | 2968 |
| chrUextra | 6702109 | 6704126 | 2018 | 2,91 | 2969 |
| chrUextra | 6706672 | 6707734 | 1063 | 2,88 | 2970 |
| chrUextra | 6709439 | 6710593 | 1155 | 2,78 | 2971 |
| chrUextra | 6713262 | 6715281 | 2020 | 2,43 | 2972 |
| chrUextra | 6723342 | 6724719 | 1378 | 3,57 | 2973 |

| | | | | | |
|---|---|---|---|---|---|
| chrUextra | 6745281 | 6746987 | 1707 | 2,33 | 2974 |
| chrUextra | 6750578 | 6752310 | 1733 | 2,39 | 2975 |
| chrUextra | 6779545 | 6780896 | 1352 | 3,31 | 2976 |
| chrUextra | 6788939 | 6790811 | 1873 | 2,78 | 2977 |
| chrUextra | 6797290 | 6799194 | 1905 | 2,3 | 2978 |
| chrUextra | 6801930 | 6803114 | 1185 | 2,3 | 2979 |
| chrUextra | 6804555 | 6805928 | 1374 | 3,22 | 2980 |
| chrUextra | 6824506 | 6826012 | 1507 | 2,26 | 2981 |
| chrUextra | 6826327 | 6828074 | 1748 | 3,17 | 2982 |
| chrUextra | 6834755 | 6835735 | 981 | 2,53 | 2983 |
| chrUextra | 6851394 | 6853323 | 1930 | 2,52 | 2984 |
| chrUextra | 6867231 | 6868313 | 1083 | 2,56 | 2985 |
| chrUextra | 6878558 | 6880613 | 2056 | 2,72 | 2986 |
| chrUextra | 6882220 | 6883227 | 1008 | 2,17 | 2987 |
| chrUextra | 6890398 | 6892335 | 1938 | 2,91 | 2988 |
| chrUextra | 6893342 | 6895132 | 1791 | 2,42 | 2989 |
| chrUextra | 6897018 | 6897900 | 883 | 2,97 | 2990 |
| chrUextra | 6900831 | 6901871 | 1041 | 2,49 | 2991 |
| chrUextra | 6923007 | 6924353 | 1347 | 2,88 | 2992 |
| chrUextra | 6926796 | 6927907 | 1112 | 2,2 | 2993 |
| chrUextra | 6937126 | 6939136 | 2011 | 2,59 | 2994 |
| chrUextra | 6940667 | 6941776 | 1110 | 2,65 | 2995 |
| chrUextra | 6949708 | 6951090 | 1383 | 3,82 | 2996 |
| chrUextra | 6952215 | 6953748 | 1534 | 2,36 | 2997 |
| chrUextra | 6958406 | 6959617 | 1212 | 3,13 | 2998 |
| chrUextra | 6963631 | 6964742 | 1112 | 4,32 | 2999 |
| chrUextra | 6973188 | 6974402 | 1215 | 2,56 | 3000 |
| chrUextra | 6988815 | 6990218 | 1404 | 3,14 | 3001 |
| chrUextra | 6997321 | 6999339 | 2019 | 3,2 | 3002 |
| chrUextra | 7016899 | 7018178 | 1280 | 3,82 | 3003 |
| chrUextra | 7018674 | 7019896 | 1223 | 2,73 | 3004 |
| chrUextra | 7020467 | 7021646 | 1180 | 2,86 | 3005 |
| chrUextra | 7024235 | 7025284 | 1050 | 2,49 | 3006 |
| chrUextra | 7026796 | 7028427 | 1632 | 2,68 | 3007 |
| chrUextra | 7031611 | 7032967 | 1357 | 2,68 | 3008 |
| chrUextra | 7038939 | 7041700 | 2762 | 9,58 | 3009 |
| chrUextra | 7072512 | 7074309 | 1798 | 2,49 | 3010 |
| chrUextra | 7080688 | 7081605 | 918 | 2,62 | 3011 |
| chrUextra | 7082592 | 7083695 | 1104 | 2,26 | 3012 |
| chrUextra | 7086659 | 7088284 | 1626 | 2,52 | 3013 |
| chrUextra | 7124161 | 7125334 | 1174 | 2,52 | 3014 |
| chrUextra | 7126005 | 7127840 | 1836 | 4,09 | 3015 |
| chrUextra | 7127870 | 7129100 | 1231 | 2,65 | 3016 |
| chrUextra | 7136220 | 7137243 | 1024 | 2,39 | 3017 |
| chrUextra | 7170289 | 7172231 | 1943 | 2,82 | 3018 |
| chrUextra | 7183309 | 7184364 | 1056 | 2,3 | 3019 |
| chrUextra | 7186938 | 7188053 | 1116 | 2,82 | 3020 |
| chrUextra | 7195233 | 7197193 | 1961 | 2,49 | 3021 |

| | | | | | |
|---|---|---|---|---|---|
| chrUextra | 7232926 | 7234003 | 1078 | 2,78 | 3022 |
| chrUextra | 7235797 | 7236813 | 1017 | 2,78 | 3023 |
| chrUextra | 7242242 | 7243275 | 1034 | 2,2 | 3024 |
| chrUextra | 7248682 | 7249796 | 1115 | 2,46 | 3025 |
| chrUextra | 7250490 | 7252132 | 1643 | 2,82 | 3026 |
| chrUextra | 7264318 | 7265351 | 1034 | 2,69 | 3027 |
| chrUextra | 7273690 | 7274851 | 1162 | 6,45 | 3028 |
| chrUextra | 7290156 | 7291202 | 1047 | 2,3 | 3029 |
| chrUextra | 7303846 | 7304977 | 1132 | 2,63 | 3030 |
| chrUextra | 7325182 | 7326977 | 1796 | 2,17 | 3031 |
| chrUextra | 7338694 | 7339946 | 1253 | 2,49 | 3032 |
| chrUextra | 7370032 | 7371200 | 1169 | 2,88 | 3033 |
| chrUextra | 7386613 | 7387671 | 1059 | 2,33 | 3034 |
| chrUextra | 7389353 | 7390399 | 1047 | 2,36 | 3035 |
| chrUextra | 7398486 | 7399548 | 1063 | 2,52 | 3036 |
| chrUextra | 7432463 | 7434085 | 1623 | 3,01 | 3037 |
| chrUextra | 7447982 | 7449829 | 1848 | 2,43 | 3038 |
| chrUextra | 7470161 | 7470861 | 701 | 4,98 | 3039 |
| chrUextra | 7482713 | 7483869 | 1157 | 2,43 | 3040 |
| chrUextra | 7486452 | 7487507 | 1056 | 3,32 | 3041 |
| chrUextra | 7496411 | 7497663 | 1253 | 3,85 | 3042 |
| chrUextra | 7500190 | 7501984 | 1795 | 2,3 | 3043 |
| chrUextra | 7516612 | 7517674 | 1063 | 2,69 | 3044 |
| chrUextra | 7538686 | 7540033 | 1348 | 2,46 | 3045 |
| chrUextra | 7598467 | 7599650 | 1184 | 4,5 | 3046 |
| chrUextra | 7616009 | 7617105 | 1097 | 3,01 | 3047 |
| chrUextra | 7629726 | 7630938 | 1213 | 2,59 | 3048 |
| chrUextra | 7633390 | 7634518 | 1129 | 2,01 | 3049 |
| chrUextra | 7656099 | 7657288 | 1190 | 2,39 | 3050 |
| chrUextra | 7681700 | 7682744 | 1045 | 3,17 | 3051 |
| chrUextra | 7703294 | 7704548 | 1255 | 3,8 | 3052 |
| chrUextra | 7713509 | 7716339 | 2831 | 3,27 | 3053 |
| chrUextra | 7729050 | 7730825 | 1776 | 2,65 | 3054 |
| chrUextra | 7733334 | 7734597 | 1264 | 3,86 | 3055 |
| chrUextra | 7753606 | 7755219 | 1614 | 3,27 | 3056 |
| chrUextra | 7768997 | 7770689 | 1693 | 2,88 | 3057 |
| chrUextra | 7785283 | 7786254 | 972 | 2,14 | 3058 |
| chrUextra | 7815847 | 7817244 | 1398 | 2,51 | 3059 |
| chrUextra | 7848128 | 7849176 | 1049 | 3,89 | 3060 |
| chrUextra | 7855127 | 7856772 | 1646 | 2,36 | 3061 |
| chrUextra | 7857815 | 7858936 | 1122 | 2,82 | 3062 |
| chrUextra | 7868814 | 7870669 | 1856 | 2,3 | 3063 |
| chrUextra | 7890392 | 7891556 | 1165 | 2,72 | 3064 |
| chrUextra | 7903232 | 7904160 | 929 | 2,69 | 3065 |
| chrUextra | 7905374 | 7906185 | 812 | 6,5 | 3066 |
| chrUextra | 7936605 | 7937646 | 1042 | 2,39 | 3067 |
| chrUextra | 7945149 | 7946664 | 1516 | 3,13 | 3068 |
| chrUextra | 7951885 | 7953271 | 1387 | 4 | 3069 |

| | | | | | |
|---|---|---|---|---|---|
| chrUextra | 7953767 | 7954711 | 945 | 2,88 | 3070 |
| chrUextra | 7958223 | 7960284 | 2062 | 4,13 | 3071 |
| chrUextra | 7979960 | 7981643 | 1684 | 2,91 | 3072 |
| chrUextra | 7983605 | 7985386 | 1782 | 3,33 | 3073 |
| chrUextra | 7989939 | 7990955 | 1017 | 2,59 | 3074 |
| chrUextra | 8001510 | 8002668 | 1159 | 3,27 | 3075 |
| chrUextra | 8038584 | 8040513 | 1930 | 2,94 | 3076 |
| chrUextra | 8043136 | 8044715 | 1580 | 2,69 | 3077 |
| chrUextra | 8045084 | 8046927 | 1844 | 5,28 | 3078 |
| chrUextra | 8066550 | 8068470 | 1921 | 2,39 | 3079 |
| chrUextra | 8072733 | 8074775 | 2043 | 3,14 | 3080 |
| chrUextra | 8075528 | 8077504 | 1977 | 2,62 | 3081 |
| chrUextra | 8078366 | 8080085 | 1720 | 2,88 | 3082 |
| chrUextra | 8089064 | 8090149 | 1086 | 2,88 | 3083 |
| chrUextra | 8103453 | 8105335 | 1883 | 2,14 | 3084 |
| chrUextra | 8126012 | 8127387 | 1376 | 3,88 | 3085 |
| chrUextra | 8128493 | 8129763 | 1271 | 2,62 | 3086 |
| chrUextra | 8133257 | 8133902 | 646 | 8,02 | 3087 |
| chrUextra | 8146817 | 8148223 | 1407 | 4,56 | 3088 |
| chrUextra | 8151191 | 8154025 | 2835 | 2,72 | 3089 |
| chrUextra | 8191545 | 8192584 | 1040 | 2,49 | 3090 |
| chrUextra | 8197675 | 8199479 | 1805 | 2,69 | 3091 |
| chrUextra | 8202234 | 8203432 | 1199 | 2,88 | 3092 |
| chrUextra | 8204791 | 8206188 | 1398 | 3,2 | 3093 |
| chrUextra | 8209489 | 8210545 | 1057 | 2,36 | 3094 |
| chrUextra | 8214035 | 8217972 | 3938 | 2,89 | 3095 |
| chrUextra | 8226502 | 8228337 | 1836 | 2,49 | 3096 |
| chrUextra | 8281146 | 8282437 | 1292 | 3,19 | 3097 |
| chrUextra | 8284858 | 8286798 | 1941 | 3,51 | 3098 |
| chrUextra | 8306467 | 8308251 | 1785 | 2,3 | 3099 |
| chrUextra | 8329586 | 8331344 | 1759 | 2,69 | 3100 |
| chrUextra | 8365454 | 8368078 | 2625 | 4,65 | 3101 |
| chrUextra | 8370779 | 8372730 | 1952 | 5,43 | 3102 |
| chrUextra | 8379616 | 8380922 | 1307 | 3,2 | 3103 |
| chrUextra | 8389268 | 8390903 | 1636 | 4,45 | 3104 |
| chrUextra | 8404730 | 8406721 | 1992 | 2,62 | 3105 |
| chrUextra | 8414924 | 8415829 | 906 | 4,34 | 3106 |
| chrUextra | 8435787 | 8438805 | 3019 | 4,52 | 3107 |
| chrUextra | 8440442 | 8441540 | 1099 | 2,3 | 3108 |
| chrUextra | 8455193 | 8456272 | 1080 | 4,08 | 3109 |
| chrUextra | 8461898 | 8463787 | 1890 | 2,46 | 3110 |
| chrUextra | 8466372 | 8468087 | 1716 | 3,58 | 3111 |
| chrUextra | 8483307 | 8484383 | 1077 | 2,3 | 3112 |
| chrUextra | 8493870 | 8496111 | 2242 | 3,34 | 3113 |
| chrUextra | 8503805 | 8505860 | 2056 | 2,7 | 3114 |
| chrUextra | 8510611 | 8511939 | 1329 | 6,62 | 3115 |
| chrUextra | 8519193 | 8521131 | 1939 | 4,53 | 3116 |
| chrUextra | 8528692 | 8531614 | 2923 | 3,6 | 3117 |

| | | | | | |
|---|---|---|---|---|---|
| chrUextra | 8559187 | 8560975 | 1789 | 2,52 | 3118 |
| chrUextra | 8573351 | 8574367 | 1017 | 3,17 | 3119 |
| chrUextra | 8575874 | 8577047 | 1174 | 2,78 | 3120 |
| chrUextra | 8596591 | 8598273 | 1683 | 2,52 | 3121 |
| chrUextra | 8607169 | 8608557 | 1389 | 2,49 | 3122 |
| chrUextra | 8610750 | 8612697 | 1948 | 2,78 | 3123 |
| chrUextra | 8614228 | 8615314 | 1087 | 3,27 | 3124 |
| chrUextra | 8664033 | 8665057 | 1025 | 2,26 | 3125 |
| chrUextra | 8670153 | 8671421 | 1269 | 3,84 | 3126 |
| chrUextra | 8682660 | 8684556 | 1897 | 2,39 | 3127 |
| chrUextra | 8746576 | 8747646 | 1071 | 2,56 | 3128 |
| chrUextra | 8749356 | 8750979 | 1624 | 2,56 | 3129 |
| chrUextra | 8752629 | 8753972 | 1344 | 2,82 | 3130 |
| chrUextra | 8755254 | 8756090 | 837 | 5,57 | 3131 |
| chrUextra | 8757151 | 8758004 | 854 | 3,22 | 3132 |
| chrUextra | 8765107 | 8766253 | 1147 | 2,58 | 3133 |
| chrUextra | 8775863 | 8777471 | 1609 | 4,33 | 3134 |
| chrUextra | 8782788 | 8784177 | 1390 | 3,35 | 3135 |
| chrUextra | 8790885 | 8791989 | 1105 | 2,56 | 3136 |
| chrUextra | 8793623 | 8796269 | 2647 | 2,65 | 3137 |
| chrUextra | 8813072 | 8814168 | 1097 | 3,17 | 3138 |
| chrUextra | 8830673 | 8832037 | 1365 | 3,35 | 3139 |
| chrUextra | 8835062 | 8836116 | 1055 | 3,03 | 3140 |
| chrUextra | 8854742 | 8855864 | 1123 | 2,85 | 3141 |
| chrUextra | 8861023 | 8863655 | 2633 | 2,78 | 3142 |
| chrUextra | 8875911 | 8877144 | 1234 | 3,49 | 3143 |
| chrUextra | 8926289 | 8927454 | 1166 | 3,07 | 3144 |
| chrUextra | 8933267 | 8934466 | 1200 | 3,52 | 3145 |
| chrUextra | 8935120 | 8936408 | 1289 | 3,79 | 3146 |
| chrUextra | 8942140 | 8943392 | 1253 | 3,13 | 3147 |
| chrUextra | 8955428 | 8956597 | 1170 | 2,75 | 3148 |
| chrUextra | 8972256 | 8974919 | 2664 | 2,43 | 3149 |
| chrUextra | 8980194 | 8982064 | 1871 | 2,62 | 3150 |
| chrUextra | 8992654 | 8994682 | 2029 | 3,48 | 3151 |
| chrUextra | 9006616 | 9007683 | 1068 | 2,56 | 3152 |
| chrUextra | 9015415 | 9016517 | 1103 | 2,88 | 3153 |
| chrUextra | 9029686 | 9031028 | 1343 | 5,99 | 3154 |
| chrUextra | 9036907 | 9037803 | 897 | 7,12 | 3155 |
| chrUextra | 9039489 | 9040308 | 820 | 3,98 | 3156 |
| chrUextra | 9061248 | 9062446 | 1199 | 2,88 | 3157 |
| chrUextra | 9077280 | 9079140 | 1861 | 2,78 | 3158 |
| chrUextra | 9083309 | 9084448 | 1140 | 4,44 | 3159 |
| chrUextra | 9093825 | 9094822 | 998 | 3,38 | 3160 |
| chrUextra | 9099012 | 9100224 | 1213 | 3,07 | 3161 |
| chrUextra | 9107210 | 9108141 | 932 | 2,26 | 3162 |
| chrUextra | 9115867 | 9117805 | 1939 | 3,14 | 3163 |
| chrUextra | 9122871 | 9124031 | 1161 | 2,52 | 3164 |
| chrUextra | 9126182 | 9127282 | 1101 | 4,08 | 3165 |

| | | | | | |
|---|---|---|---|---|---|
| chrUextra | 9132529 | 9133847 | 1319 | 3,62 | 3166 |
| chrUextra | 9140520 | 9141638 | 1119 | 2,46 | 3167 |
| chrUextra | 9142306 | 9144268 | 1963 | 2,43 | 3168 |
| chrUextra | 9147408 | 9148646 | 1239 | 3,43 | 3169 |
| chrUextra | 9169669 | 9170713 | 1045 | 3,14 | 3170 |
| chrUextra | 9173126 | 9174795 | 1670 | 2,36 | 3171 |
| chrUextra | 9176657 | 9177688 | 1032 | 2,52 | 3172 |
| chrUextra | 9186884 | 9188382 | 1499 | 2,66 | 3173 |
| chrUextra | 9203817 | 9204919 | 1103 | 2,36 | 3174 |
| chrUextra | 9207303 | 9208408 | 1106 | 2,98 | 3175 |
| chrUextra | 9224918 | 9226824 | 1907 | 2,65 | 3176 |
| chrUextra | 9233468 | 9234960 | 1493 | 3,96 | 3177 |
| chrUextra | 9243310 | 9244441 | 1132 | 3,14 | 3178 |
| chrUextra | 9258290 | 9259525 | 1236 | 2,69 | 3179 |
| chrUextra | 9265676 | 9267327 | 1652 | 4,05 | 3180 |
| chrUextra | 9272296 | 9276109 | 3814 | 3,04 | 3181 |
| chrUextra | 9292526 | 9294456 | 1931 | 2,3 | 3182 |
| chrUextra | 9301265 | 9302307 | 1043 | 2,49 | 3183 |
| chrUextra | 9311803 | 9312757 | 955 | 3,04 | 3184 |
| chrUextra | 9330170 | 9331382 | 1213 | 2,45 | 3185 |
| chrUextra | 9353903 | 9354981 | 1079 | 3,33 | 3186 |
| chrUextra | 9356504 | 9357384 | 881 | 6,34 | 3187 |
| chrUextra | 9384583 | 9385644 | 1062 | 2,82 | 3188 |
| chrUextra | 9389740 | 9390896 | 1157 | 2,56 | 3189 |
| chrUextra | 9402724 | 9404054 | 1331 | 4,18 | 3190 |
| chrUextra | 9407314 | 9408283 | 970 | 2,53 | 3191 |
| chrUextra | 9408368 | 9409809 | 1442 | 2,1 | 3192 |
| chrUextra | 9418684 | 9419750 | 1067 | 3,02 | 3193 |
| chrUextra | 9457265 | 9458207 | 943 | 2,98 | 3194 |
| chrUextra | 9465107 | 9466783 | 1677 | 3,04 | 3195 |
| chrUextra | 9472065 | 9473065 | 1001 | 2,56 | 3196 |
| chrUextra | 9493808 | 9495112 | 1305 | 3,11 | 3197 |
| chrUextra | 9509598 | 9510719 | 1122 | 2,75 | 3198 |
| chrUextra | 9524534 | 9525673 | 1140 | 3,24 | 3199 |
| chrUextra | 9538482 | 9539543 | 1062 | 2,49 | 3200 |
| chrUextra | 9540007 | 9541289 | 1283 | 2,67 | 3201 |
| chrUextra | 9544620 | 9546692 | 2073 | 4,37 | 3202 |
| chrUextra | 9566305 | 9569033 | 2729 | 3,19 | 3203 |
| chrUextra | 9572533 | 9573576 | 1044 | 2,56 | 3204 |
| chrUextra | 9592505 | 9594326 | 1822 | 3,11 | 3205 |
| chrUextra | 9596957 | 9598007 | 1051 | 2,52 | 3206 |
| chrUextra | 9598620 | 9599785 | 1166 | 2,36 | 3207 |
| chrUextra | 9603021 | 9604123 | 1103 | 2,65 | 3208 |
| chrUextra | 9616866 | 9618922 | 2057 | 2,51 | 3209 |
| chrUextra | 9619634 | 9621567 | 1934 | 2,62 | 3210 |
| chrUextra | 9637153 | 9638994 | 1842 | 2,39 | 3211 |
| chrUextra | 9644987 | 9646608 | 1622 | 2,33 | 3212 |
| chrUextra | 9648313 | 9649643 | 1331 | 3,78 | 3213 |

| | | | | | |
|---|---|---|---|---|---|
| chrUextra | 9653640 | 9655226 | 1587 | 3,69 | 3214 |
| chrUextra | 9664855 | 9666736 | 1882 | 3,49 | 3215 |
| chrUextra | 9674556 | 9675621 | 1066 | 2,94 | 3216 |
| chrUextra | 9682378 | 9683462 | 1085 | 2,49 | 3217 |
| chrUextra | 9686380 | 9687942 | 1563 | 3,88 | 3218 |
| chrUextra | 9733106 | 9734705 | 1600 | 2,17 | 3219 |
| chrUextra | 9737639 | 9739116 | 1478 | 3,04 | 3220 |
| chrUextra | 9757268 | 9759143 | 1876 | 3,26 | 3221 |
| chrUextra | 9769650 | 9770600 | 951 | 5,08 | 3222 |
| chrUextra | 9776625 | 9778246 | 1622 | 2,67 | 3223 |
| chrUextra | 9787715 | 9788742 | 1028 | 3,9 | 3224 |
| chrUextra | 9799878 | 9800944 | 1067 | 2,3 | 3225 |
| chrUextra | 9817264 | 9818939 | 1676 | 2,56 | 3226 |
| chrUextra | 9818970 | 9820890 | 1921 | 3,3 | 3227 |
| chrUextra | 9828665 | 9829649 | 985 | 2,26 | 3228 |
| chrUextra | 9834456 | 9835262 | 807 | 5,69 | 3229 |
| chrUextra | 9847478 | 9849366 | 1889 | 4,85 | 3230 |
| chrUextra | 9853127 | 9854750 | 1624 | 2,23 | 3231 |
| chrUextra | 9946604 | 9948410 | 1807 | 2,62 | 3232 |
| chrUextra | 9959618 | 9961231 | 1614 | 2,52 | 3233 |
| chrUextra | 9963893 | 9964903 | 1011 | 2,43 | 3234 |
| chrUextra | 10014281 | 10016144 | 1864 | 2,59 | 3235 |
| chrUextra | 10025478 | 10027533 | 2056 | 2,85 | 3236 |
| chrUextra | 10039365 | 10041231 | 1867 | 2,56 | 3237 |
| chrUextra | 10048729 | 10050003 | 1275 | 4,59 | 3238 |
| chrUextra | 10052315 | 10053350 | 1036 | 2,17 | 3239 |
| chrUextra | 10053561 | 10055200 | 1640 | 2,84 | 3240 |
| chrUextra | 10086689 | 10088863 | 2175 | 3,75 | 3241 |
| chrUextra | 10107703 | 10108759 | 1057 | 2,62 | 3242 |
| chrUextra | 10133652 | 10134779 | 1128 | 3,07 | 3243 |
| chrUextra | 10141483 | 10143346 | 1864 | 3,3 | 3244 |
| chrUextra | 10151175 | 10152290 | 1116 | 6,21 | 3245 |
| chrUextra | 10161945 | 10164037 | 2093 | 3,78 | 3246 |
| chrUextra | 10169067 | 10170398 | 1332 | 2,82 | 3247 |
| chrUextra | 10188996 | 10192232 | 3237 | 2,26 | 3248 |
| chrUextra | 10217609 | 10219328 | 1720 | 3,98 | 3249 |
| chrUextra | 10222679 | 10224310 | 1632 | 2,26 | 3250 |
| chrUextra | 10228670 | 10229724 | 1055 | 2,52 | 3251 |
| chrUextra | 10238159 | 10239370 | 1212 | 4,24 | 3252 |
| chrUextra | 10240600 | 10242085 | 1486 | 7,15 | 3253 |
| chrUextra | 10245030 | 10246616 | 1587 | 2,36 | 3254 |
| chrUextra | 10251137 | 10252145 | 1009 | 2,3 | 3255 |
| chrUextra | 10257974 | 10259085 | 1112 | 2,3 | 3256 |
| chrUextra | 10273615 | 10275014 | 1400 | 5,47 | 3257 |
| chrUextra | 10275989 | 10276650 | 662 | 5,89 | 3258 |
| chrUextra | 10276982 | 10278063 | 1082 | 2,49 | 3259 |
| chrUextra | 10286198 | 10287595 | 1398 | 3,16 | 3260 |
| chrUextra | 10295992 | 10297018 | 1027 | 2,46 | 3261 |

| | | | | | |
|---|---|---|---|---|---|
| chrUextra | 10298484 | 10299574 | 1091 | 2,61 | 3262 |
| chrUextra | 10307210 | 10308938 | 1729 | 2,43 | 3263 |
| chrUextra | 10342375 | 10343522 | 1148 | 3,62 | 3264 |
| chrUextra | 10402754 | 10403669 | 916 | 2,46 | 3265 |
| chrUextra | 10434613 | 10435648 | 1036 | 2,56 | 3266 |
| chrUextra | 10458898 | 10460459 | 1562 | 2,82 | 3267 |
| chrUextra | 10493954 | 10494953 | 1000 | 2,82 | 3268 |
| chrUextra | 10518859 | 10519856 | 998 | 2,91 | 3269 |
| chrUextra | 10534169 | 10535467 | 1299 | 4,12 | 3270 |
| chrUextra | 10545413 | 10546734 | 1322 | 3,27 | 3271 |
| chrUextra | 10556919 | 10558640 | 1722 | 3,08 | 3272 |
| chrUextra | 10559154 | 10561025 | 1872 | 2,91 | 3273 |
| chrUextra | 10572185 | 10573168 | 984 | 2,59 | 3274 |
| chrUextra | 10582466 | 10583468 | 1003 | 3,04 | 3275 |
| chrUextra | 10589091 | 10590485 | 1395 | 2,91 | 3276 |
| chrUextra | 10592741 | 10593826 | 1086 | 2,46 | 3277 |
| chrUextra | 10595242 | 10596307 | 1066 | 2,75 | 3278 |
| chrUextra | 10597859 | 10598799 | 941 | 2,26 | 3279 |
| chrUextra | 10609887 | 10612616 | 2730 | 2,62 | 3280 |
| chrUextra | 10618460 | 10619445 | 986 | 2,59 | 3281 |
| chrUextra | 10660352 | 10661458 | 1107 | 2,85 | 3282 |
| chrUextra | 10664601 | 10665959 | 1359 | 4,33 | 3283 |
| chrUextra | 10685329 | 10687257 | 1929 | 2,85 | 3284 |
| chrUextra | 10687972 | 10688955 | 984 | 2,56 | 3285 |
| chrUextra | 10711923 | 10713025 | 1103 | 2,72 | 3286 |
| chrUextra | 10725328 | 10725949 | 622 | 3,98 | 3287 |
| chrUextra | 10736143 | 10737791 | 1649 | 2,49 | 3288 |
| chrUextra | 10739160 | 10740587 | 1428 | 2,85 | 3289 |
| chrUextra | 10741886 | 10742929 | 1044 | 2,56 | 3290 |
| chrUextra | 10758066 | 10759155 | 1090 | 2,56 | 3291 |
| chrUextra | 10769992 | 10771316 | 1325 | 3,2 | 3292 |
| chrUextra | 10771919 | 10772866 | 948 | 2,33 | 3293 |
| chrUextra | 10795928 | 10797541 | 1614 | 2,43 | 3294 |
| chrUextra | 10798331 | 10799685 | 1355 | 2,94 | 3295 |
| chrUextra | 10800912 | 10802799 | 1888 | 2,97 | 3296 |
| chrUextra | 10805231 | 10806405 | 1175 | 3,85 | 3297 |
| chrUextra | 10806943 | 10808063 | 1121 | 2,62 | 3298 |
| chrUextra | 10828258 | 10829391 | 1134 | 3,2 | 3299 |
| chrUextra | 10829869 | 10831290 | 1422 | 2,98 | 3300 |
| chrUextra | 10834375 | 10835920 | 1546 | 2,17 | 3301 |
| chrUextra | 10871087 | 10872115 | 1029 | 2,78 | 3302 |
| chrUextra | 10884755 | 10886283 | 1529 | 2,26 | 3303 |
| chrUextra | 10887381 | 10888804 | 1424 | 6,53 | 3304 |
| chrUextra | 10902788 | 10904133 | 1346 | 4,28 | 3305 |
| chrUextra | 10904321 | 10906497 | 2177 | 2,89 | 3306 |
| chrUextra | 10917205 | 10918334 | 1130 | 3,01 | 3307 |
| chrUextra | 10922315 | 10923368 | 1054 | 2,75 | 3308 |
| chrUextra | 10926637 | 10927968 | 1332 | 4,76 | 3309 |

| | | | | | |
|---|---|---|---|---|---|
| chrUextra | 10932598 | 10933643 | 1046 | 2,49 | 3310 |
| chrUextra | 10936773 | 10938157 | 1385 | 3,47 | 3311 |
| chrUextra | 10942931 | 10943976 | 1046 | 2,62 | 3312 |
| chrUextra | 10945319 | 10946363 | 1045 | 3,49 | 3313 |
| chrUextra | 10958239 | 10959444 | 1206 | 3,21 | 3314 |
| chrUextra | 10984500 | 10986637 | 2138 | 2,74 | 3315 |
| chrUextra | 11005969 | 11007240 | 1272 | 3,45 | 3316 |
| chrUextra | 11017227 | 11018218 | 992 | 2,62 | 3317 |
| chrUextra | 11028330 | 11030335 | 2006 | 2,91 | 3318 |
| chrUextra | 11031643 | 11032672 | 1030 | 2,43 | 3319 |
| chrUextra | 11034202 | 11036295 | 2094 | 4,2 | 3320 |
| chrUextra | 11036877 | 11037853 | 977 | 2,65 | 3321 |
| chrUextra | 11042662 | 11043782 | 1121 | 2,23 | 3322 |
| chrUextra | 11049874 | 11053134 | 3261 | 4,83 | 3323 |
| chrUextra | 11058089 | 11059861 | 1773 | 2,75 | 3324 |
| chrUextra | 11069974 | 11071076 | 1103 | 2,17 | 3325 |
| chrUextra | 11081093 | 11082153 | 1061 | 2,82 | 3326 |
| chrUextra | 11104171 | 11105150 | 980 | 2,98 | 3327 |
| chrUextra | 11107526 | 11109287 | 1762 | 2,62 | 3328 |
| chrUextra | 11114338 | 11115383 | 1046 | 2,33 | 3329 |
| chrUextra | 11122138 | 11124599 | 2462 | 3,2 | 3330 |
| chrUextra | 11151011 | 11152764 | 1754 | 3,26 | 3331 |
| chrUextra | 11167153 | 11168198 | 1046 | 3,37 | 3332 |
| chrUextra | 11204504 | 11205701 | 1198 | 3,67 | 3333 |
| chrUextra | 11209472 | 11211040 | 1569 | 3,96 | 3334 |
| chrUextra | 11224942 | 11226031 | 1090 | 3,2 | 3335 |
| chrUextra | 11241293 | 11242840 | 1548 | 3,33 | 3336 |
| chrUextra | 11252267 | 11253292 | 1026 | 2,88 | 3337 |
| chrUextra | 11263250 | 11264282 | 1033 | 3,07 | 3338 |
| chrUextra | 11271787 | 11272850 | 1064 | 2,39 | 3339 |
| chrUextra | 11281235 | 11283556 | 2322 | 2,39 | 3340 |
| chrUextra | 11287107 | 11288182 | 1076 | 2,72 | 3341 |
| chrUextra | 11303242 | 11304326 | 1085 | 2,97 | 3342 |
| chrUextra | 11313450 | 11314501 | 1052 | 2,26 | 3343 |
| chrUextra | 11315007 | 11316417 | 1411 | 2,73 | 3344 |
| chrUextra | 11340731 | 11343242 | 2512 | 2,39 | 3345 |
| chrUextra | 11344154 | 11345393 | 1240 | 4,03 | 3346 |
| chrUextra | 11345759 | 11346831 | 1073 | 2,56 | 3347 |
| chrUextra | 11365217 | 11366407 | 1191 | 2,56 | 3348 |
| chrUextra | 11368633 | 11369709 | 1077 | 3,44 | 3349 |
| chrUextra | 11374569 | 11375693 | 1125 | 2,88 | 3350 |
| chrUextra | 11389059 | 11390253 | 1195 | 3,2 | 3351 |
| chrUextra | 11400087 | 11402703 | 2617 | 2,29 | 3352 |
| chrUextra | 11404456 | 11406268 | 1813 | 2,71 | 3353 |
| chrUextra | 11407061 | 11409615 | 2555 | 2,78 | 3354 |
| chrUextra | 11411986 | 11413005 | 1020 | 2,88 | 3355 |
| chrUextra | 11425869 | 11426796 | 928 | 4,34 | 3356 |
| chrUextra | 11435743 | 11436789 | 1047 | 3,4 | 3357 |

| | | | | | |
|---|---|---|---|---|---|
| chrUextra | 11437353 | 11438597 | 1245 | 3,53 | 3358 |
| chrUextra | 11439206 | 11441091 | 1886 | 2,65 | 3359 |
| chrUextra | 11447460 | 11448658 | 1199 | 3,96 | 3360 |
| chrUextra | 11456313 | 11458071 | 1759 | 3,3 | 3361 |
| chrUextra | 11461270 | 11462314 | 1045 | 2,59 | 3362 |
| chrUextra | 11465424 | 11466443 | 1020 | 2,43 | 3363 |
| chrUextra | 11471363 | 11472370 | 1008 | 2,72 | 3364 |
| chrUextra | 11483248 | 11485176 | 1929 | 2,23 | 3365 |
| chrUextra | 11507028 | 11508041 | 1014 | 2,39 | 3366 |
| chrUextra | 11518799 | 11520956 | 2158 | 3 | 3367 |
| chrUextra | 11539295 | 11540966 | 1672 | 2,26 | 3368 |
| chrUextra | 11541734 | 11542939 | 1206 | 2,82 | 3369 |
| chrUextra | 11544267 | 11546903 | 2637 | 3,76 | 3370 |
| chrUextra | 11561208 | 11562248 | 1041 | 2,91 | 3371 |
| chrUextra | 11579688 | 11580695 | 1008 | 3,72 | 3372 |
| chrUextra | 11584223 | 11585990 | 1768 | 2,52 | 3373 |
| chrUextra | 11596707 | 11598710 | 2004 | 2,72 | 3374 |
| chrUextra | 11602749 | 11604605 | 1857 | 2,2 | 3375 |
| chrUextra | 11649052 | 11650341 | 1290 | 3,37 | 3376 |
| chrUextra | 11678004 | 11679050 | 1047 | 2,78 | 3377 |
| chrUextra | 11685682 | 11687510 | 1829 | 2,65 | 3378 |
| chrUextra | 11706806 | 11708688 | 1883 | 2,62 | 3379 |
| chrUextra | 11719366 | 11720637 | 1272 | 3,25 | 3380 |
| chrUextra | 11736339 | 11739084 | 2746 | 2,72 | 3381 |
| chrUextra | 11749833 | 11750886 | 1054 | 2,82 | 3382 |
| chrUextra | 11780234 | 11781553 | 1320 | 4,2 | 3383 |
| chrUextra | 11804678 | 11806116 | 1439 | 2,56 | 3384 |
| chrUextra | 11821459 | 11823507 | 2049 | 3,49 | 3385 |
| chrUextra | 11832484 | 11833684 | 1201 | 3,04 | 3386 |
| chrUextra | 11838425 | 11840587 | 2163 | 4,04 | 3387 |
| chrUextra | 11847380 | 11848879 | 1500 | 2,52 | 3388 |
| chrUextra | 11854687 | 11855591 | 905 | 3,64 | 3389 |
| chrUextra | 11858758 | 11859784 | 1027 | 3,2 | 3390 |
| chrUextra | 11894341 | 11896048 | 1708 | 2,56 | 3391 |
| chrUextra | 11900899 | 11901785 | 887 | 3,9 | 3392 |
| chrUextra | 11911692 | 11912721 | 1030 | 2,69 | 3393 |
| chrUextra | 11937354 | 11939220 | 1867 | 3,72 | 3394 |
| chrUextra | 11940601 | 11941525 | 925 | 2,72 | 3395 |
| chrUextra | 11955844 | 11956949 | 1106 | 3,27 | 3396 |
| chrUextra | 11960788 | 11961793 | 1006 | 2,52 | 3397 |
| chrUextra | 11974364 | 11975437 | 1074 | 2,72 | 3398 |
| chrUextra | 11977669 | 11978830 | 1162 | 3,89 | 3399 |
| chrUextra | 11985431 | 11986683 | 1253 | 2,14 | 3400 |
| chrUextra | 12007929 | 12009876 | 1948 | 3,43 | 3401 |
| chrUextra | 12029921 | 12031499 | 1579 | 2,33 | 3402 |
| chrUextra | 12047581 | 12048803 | 1223 | 3,26 | 3403 |
| chrUextra | 12049344 | 12051901 | 2558 | 3,48 | 3404 |
| chrUextra | 12051910 | 12053331 | 1422 | 4,82 | 3405 |

| | | | | | |
|---|---|---|---|---|---|
| chrUextra | 12065360 | 12066202 | 843 | 2,56 | 3406 |
| chrUextra | 12072106 | 12073834 | 1729 | 2,72 | 3407 |
| chrUextra | 12080391 | 12082222 | 1832 | 2,36 | 3408 |
| chrUextra | 12088219 | 12089798 | 1580 | 2,2 | 3409 |
| chrUextra | 12093960 | 12095988 | 2029 | 3,25 | 3410 |
| chrUextra | 12101380 | 12102495 | 1116 | 2,46 | 3411 |
| chrUextra | 12104003 | 12105874 | 1872 | 2,75 | 3412 |
| chrUextra | 12122456 | 12123423 | 968 | 2,46 | 3413 |
| chrUextra | 12161865 | 12162983 | 1119 | 3,49 | 3414 |
| chrUextra | 12183929 | 12184876 | 948 | 2,56 | 3415 |
| chrUextra | 12187845 | 12189017 | 1173 | 4,05 | 3416 |
| chrUextra | 12209795 | 12211000 | 1206 | 3,17 | 3417 |
| chrUextra | 12219850 | 12221193 | 1344 | 2,44 | 3418 |
| chrUextra | 12225847 | 12227658 | 1812 | 3,11 | 3419 |
| chrUextra | 12240969 | 12242654 | 1686 | 2,33 | 3420 |
| chrUextra | 12260333 | 12262188 | 1856 | 3,01 | 3421 |
| chrUextra | 12266958 | 12267939 | 982 | 3,07 | 3422 |
| chrUextra | 12270952 | 12273087 | 2136 | 3,2 | 3423 |
| chrUextra | 12278029 | 12279746 | 1718 | 4,29 | 3424 |
| chrUextra | 12285696 | 12288170 | 2475 | 2,59 | 3425 |
| chrUextra | 12291294 | 12292343 | 1050 | 2,72 | 3426 |
| chrUextra | 12295707 | 12297471 | 1765 | 2,79 | 3427 |
| chrUextra | 12303721 | 12304644 | 924 | 5,47 | 3428 |
| chrUextra | 12322211 | 12323303 | 1093 | 2,62 | 3429 |
| chrUextra | 12325821 | 12328400 | 2580 | 2,43 | 3430 |
| chrUextra | 12329880 | 12331167 | 1288 | 3,17 | 3431 |
| chrUextra | 12354257 | 12355273 | 1017 | 2,36 | 3432 |
| chrUextra | 12361010 | 12362820 | 1811 | 2,52 | 3433 |
| chrUextra | 12363464 | 12364501 | 1038 | 2,43 | 3434 |
| chrUextra | 12381444 | 12382265 | 822 | 6,1 | 3435 |
| chrUextra | 12385225 | 12386307 | 1083 | 3,68 | 3436 |
| chrUextra | 12394590 | 12395520 | 931 | 2,49 | 3437 |
| chrUextra | 12417954 | 12419000 | 1047 | 2,85 | 3438 |
| chrUextra | 12441906 | 12443151 | 1246 | 8,61 | 3439 |
| chrUextra | 12448130 | 12449134 | 1005 | 2,56 | 3440 |
| chrUextra | 12451450 | 12453218 | 1769 | 2,43 | 3441 |
| chrUextra | 12455617 | 12456751 | 1135 | 2,82 | 3442 |
| chrUextra | 12458315 | 12459517 | 1203 | 3,85 | 3443 |
| chrUextra | 12460512 | 12461897 | 1386 | 3,3 | 3444 |
| chrUextra | 12498354 | 12499380 | 1027 | 3,49 | 3445 |
| chrUextra | 12500036 | 12501146 | 1111 | 3,24 | 3446 |
| chrUextra | 12503319 | 12504956 | 1638 | 3,52 | 3447 |
| chrUextra | 12510883 | 12512769 | 1887 | 3,03 | 3448 |
| chrUextra | 12546887 | 12547542 | 656 | 3,79 | 3449 |
| chrUextra | 12557806 | 12558967 | 1162 | 3,56 | 3450 |
| chrUextra | 12570336 | 12571405 | 1070 | 2,59 | 3451 |
| chrUextra | 12593830 | 12595602 | 1773 | 2,42 | 3452 |
| chrUextra | 12606214 | 12607363 | 1150 | 2,54 | 3453 |

| | | | | | |
|---|---|---|---|---|---|
| chrUextra | 12632176 | 12634090 | 1915 | 2,63 | 3454 |
| chrUextra | 12640344 | 12641580 | 1237 | 3,43 | 3455 |
| chrUextra | 12647201 | 12648294 | 1094 | 2,65 | 3456 |
| chrUextra | 12687332 | 12688498 | 1167 | 3,56 | 3457 |
| chrUextra | 12697389 | 12698314 | 926 | 2,23 | 3458 |
| chrUextra | 12698566 | 12699518 | 953 | 4,92 | 3459 |
| chrUextra | 12708305 | 12709788 | 1484 | 2,43 | 3460 |
| chrUextra | 12725741 | 12726876 | 1136 | 2,39 | 3461 |
| chrUextra | 12756634 | 12758503 | 1870 | 2,59 | 3462 |
| chrUextra | 12763251 | 12764285 | 1035 | 2,56 | 3463 |
| chrUextra | 12796060 | 12797716 | 1657 | 2,62 | 3464 |
| chrUextra | 12800766 | 12802090 | 1325 | 3,1 | 3465 |
| chrUextra | 12804316 | 12805893 | 1578 | 2,56 | 3466 |
| chrUextra | 12805902 | 12807809 | 1908 | 3,66 | 3467 |
| chrUextra | 12812702 | 12815230 | 2529 | 2,8 | 3468 |
| chrUextra | 12830026 | 12831141 | 1116 | 2,69 | 3469 |
| chrUextra | 12859204 | 12861046 | 1843 | 2,3 | 3470 |
| chrUextra | 12866740 | 12867793 | 1054 | 6,02 | 3471 |
| chrUextra | 12877606 | 12878638 | 1033 | 3,07 | 3472 |
| chrUextra | 12880935 | 12882018 | 1084 | 2,52 | 3473 |
| chrUextra | 12911772 | 12914547 | 2776 | 2,34 | 3474 |
| chrUextra | 12917622 | 12918694 | 1073 | 2,75 | 3475 |
| chrUextra | 12987683 | 12989475 | 1793 | 3,24 | 3476 |
| chrUextra | 12994032 | 12995400 | 1369 | 2,97 | 3477 |
| chrUextra | 12995885 | 12996896 | 1012 | 2,33 | 3478 |
| chrUextra | 13000072 | 13001214 | 1143 | 2,91 | 3479 |
| chrUextra | 13003369 | 13006319 | 2951 | 2,81 | 3480 |
| chrUextra | 13013478 | 13015073 | 1596 | 2,39 | 3481 |
| chrUextra | 13019248 | 13021940 | 2693 | 2,64 | 3482 |
| chrUextra | 13023418 | 13024360 | 943 | 2,56 | 3483 |
| chrUextra | 13077556 | 13079191 | 1636 | 2,1 | 3484 |
| chrUextra | 13087518 | 13088695 | 1178 | 2,65 | 3485 |
| chrUextra | 13104917 | 13105997 | 1081 | 2,78 | 3486 |
| chrUextra | 13111409 | 13112772 | 1364 | 2,94 | 3487 |
| chrUextra | 13117405 | 13118427 | 1023 | 2,98 | 3488 |
| chrUextra | 13122192 | 13123581 | 1390 | 4,31 | 3489 |
| chrUextra | 13126580 | 13127628 | 1049 | 3,01 | 3490 |
| chrUextra | 13130542 | 13131744 | 1203 | 4,38 | 3491 |
| chrUextra | 13136509 | 13137625 | 1117 | 2,85 | 3492 |
| chrUextra | 13146423 | 13147695 | 1273 | 4,18 | 3493 |
| chrUextra | 13149840 | 13150929 | 1090 | 3,19 | 3494 |
| chrUextra | 13158141 | 13159800 | 1660 | 3,37 | 3495 |
| chrUextra | 13163046 | 13164143 | 1098 | 2,85 | 3496 |
| chrUextra | 13197239 | 13198222 | 984 | 2,72 | 3497 |
| chrUextra | 13220511 | 13221492 | 982 | 2,82 | 3498 |
| chrUextra | 13250346 | 13252239 | 1894 | 3,01 | 3499 |
| chrUextra | 13256148 | 13257150 | 1003 | 2,65 | 3500 |
| chrUextra | 13258844 | 13260406 | 1563 | 2,23 | 3501 |

| | | | | | |
|---|---|---|---|---|---|
| chrUextra | 13264506 | 13265539 | 1034 | 2,52 | 3502 |
| chrUextra | 13266656 | 13267767 | 1112 | 5,94 | 3503 |
| chrUextra | 13271779 | 13272980 | 1202 | 3,32 | 3504 |
| chrUextra | 13274442 | 13275602 | 1161 | 2,75 | 3505 |
| chrUextra | 13283647 | 13285448 | 1802 | 2,23 | 3506 |
| chrUextra | 13290038 | 13291432 | 1395 | 2,74 | 3507 |
| chrUextra | 13307547 | 13308754 | 1208 | 3,72 | 3508 |
| chrUextra | 13323427 | 13324069 | 643 | 5,15 | 3509 |
| chrUextra | 13326419 | 13327110 | 692 | 4,95 | 3510 |
| chrUextra | 13344122 | 13346017 | 1896 | 2,88 | 3511 |
| chrUextra | 13364075 | 13367767 | 3693 | 3,7 | 3512 |
| chrUextra | 13377218 | 13378393 | 1176 | 2,59 | 3513 |
| chrUextra | 13391364 | 13393316 | 1953 | 2,42 | 3514 |
| chrUextra | 13403005 | 13404063 | 1059 | 2,62 | 3515 |
| chrUextra | 13412159 | 13414027 | 1869 | 3,4 | 3516 |
| chrUextra | 13422738 | 13424193 | 1456 | 3,62 | 3517 |
| chrUextra | 13432916 | 13433903 | 988 | 2,56 | 3518 |
| chrUextra | 13439457 | 13440541 | 1085 | 2,39 | 3519 |
| chrUextra | 13456619 | 13457790 | 1172 | 4,11 | 3520 |
| chrUextra | 13463504 | 13464567 | 1064 | 2,72 | 3521 |
| chrUextra | 13494677 | 13495442 | 766 | 4,43 | 3522 |
| chrUextra | 13495909 | 13497280 | 1372 | 2,23 | 3523 |
| chrUextra | 13499186 | 13500183 | 998 | 2,46 | 3524 |
| chrUextra | 13516521 | 13517719 | 1199 | 3,24 | 3525 |
| chrUextra | 13557847 | 13559701 | 1855 | 4 | 3526 |
| chrUextra | 13627619 | 13630776 | 3158 | 2,82 | 3527 |
| chrUextra | 13640609 | 13641789 | 1181 | 2,69 | 3528 |
| chrUextra | 13659678 | 13660647 | 970 | 2,43 | 3529 |
| chrUextra | 13672140 | 13673123 | 984 | 2,91 | 3530 |
| chrUextra | 13674018 | 13675107 | 1090 | 5,05 | 3531 |
| chrUextra | 13679466 | 13680566 | 1101 | 4,51 | 3532 |
| chrUextra | 13715097 | 13717846 | 2750 | 3,53 | 3533 |
| chrUextra | 13720760 | 13722004 | 1245 | 3,65 | 3534 |
| chrUextra | 13748137 | 13749193 | 1057 | 2,65 | 3535 |
| chrUextra | 13753046 | 13754810 | 1765 | 3,5 | 3536 |
| chrUextra | 13754863 | 13756586 | 1724 | 2,38 | 3537 |
| chrUextra | 13776132 | 13777295 | 1164 | 2,78 | 3538 |
| chrUextra | 13785292 | 13785724 | 433 | 3,86 | 3539 |
| chrUextra | 13789489 | 13790169 | 681 | 4,76 | 3540 |
| chrUextra | 13790303 | 13791529 | 1227 | 4,04 | 3541 |
| chrUextra | 13796540 | 13798558 | 2019 | 3,76 | 3542 |
| chrUextra | 13803473 | 13804425 | 953 | 2,94 | 3543 |
| chrUextra | 13808879 | 13809791 | 913 | 5,21 | 3544 |
| chrUextra | 13820767 | 13821956 | 1190 | 3,23 | 3545 |
| chrUextra | 13823313 | 13825250 | 1938 | 3,67 | 3546 |
| chrUextra | 13834864 | 13835874 | 1011 | 2,36 | 3547 |
| chrUextra | 13848712 | 13850069 | 1358 | 3,28 | 3548 |
| chrUextra | 13861986 | 13863244 | 1259 | 4,44 | 3549 |

| | | | | | |
|---|---|---|---|---|---|
| chrUextra | 13894868 | 13895738 | 871 | 3,88 | 3550 |
| chrUextra | 13896835 | 13898485 | 1651 | 2,14 | 3551 |
| chrUextra | 13917358 | 13918456 | 1099 | 3,52 | 3552 |
| chrUextra | 13934590 | 13935553 | 964 | 3,61 | 3553 |
| chrUextra | 13938676 | 13939900 | 1225 | 2,58 | 3554 |
| chrUextra | 13954502 | 13955557 | 1056 | 2,85 | 3555 |
| chrUextra | 13979140 | 13980216 | 1077 | 3,72 | 3556 |
| chrUextra | 13993194 | 13994206 | 1013 | 2,72 | 3557 |
| chrUextra | 14004567 | 14005777 | 1211 | 4,18 | 3558 |
| chrUextra | 14037689 | 14038880 | 1192 | 2,84 | 3559 |
| chrUextra | 14040329 | 14041890 | 1562 | 2,1 | 3560 |
| chrUextra | 14051786 | 14053405 | 1620 | 2,17 | 3561 |
| chrUextra | 14054193 | 14055188 | 996 | 2,46 | 3562 |
| chrUextra | 14055724 | 14057021 | 1298 | 3,34 | 3563 |
| chrUextra | 14072498 | 14074069 | 1572 | 2,62 | 3564 |
| chrUextra | 14077258 | 14078214 | 957 | 3,49 | 3565 |
| chrUextra | 14102582 | 14104548 | 1967 | 4,63 | 3566 |
| chrUextra | 14109286 | 14111163 | 1878 | 3,88 | 3567 |
| chrUextra | 14123391 | 14124415 | 1025 | 2,65 | 3568 |
| chrUextra | 14137444 | 14138486 | 1043 | 3,26 | 3569 |
| chrUextra | 14167692 | 14168925 | 1234 | 3,8 | 3570 |
| chrUextra | 14182697 | 14185272 | 2576 | 3,03 | 3571 |
| chrUextra | 14186042 | 14187773 | 1732 | 2,71 | 3572 |
| chrUextra | 14200730 | 14202414 | 1685 | 2,52 | 3573 |
| chrUextra | 14204918 | 14205839 | 922 | 4,04 | 3574 |
| chrUextra | 14206467 | 14207585 | 1119 | 2,96 | 3575 |
| chrUextra | 14223884 | 14225670 | 1787 | 2,65 | 3576 |
| chrUextra | 14236783 | 14237856 | 1074 | 3,37 | 3577 |
| chrUextra | 14266547 | 14267555 | 1009 | 3,3 | 3578 |
| chrUextra | 14293418 | 14295358 | 1941 | 3,33 | 3579 |
| chrUextra | 14298576 | 14299521 | 946 | 2,98 | 3580 |
| chrUextra | 14304220 | 14305310 | 1091 | 3,59 | 3581 |
| chrUextra | 14305941 | 14307145 | 1205 | 3,41 | 3582 |
| chrUextra | 14313829 | 14315174 | 1346 | 2,43 | 3583 |
| chrUextra | 14315810 | 14316699 | 890 | 2,72 | 3584 |
| chrUextra | 14319855 | 14321766 | 1912 | 3,2 | 3585 |
| chrUextra | 14325675 | 14327589 | 1915 | 2,64 | 3586 |
| chrUextra | 14328121 | 14329896 | 1776 | 2,41 | 3587 |
| chrUextra | 14330647 | 14331668 | 1022 | 3,4 | 3588 |
| chrUextra | 14338519 | 14339928 | 1410 | 3,15 | 3589 |
| chrUextra | 14344541 | 14345568 | 1028 | 2,36 | 3590 |
| chrUextra | 14360139 | 14362800 | 2662 | 2,82 | 3591 |
| chrUextra | 14363437 | 14364397 | 961 | 3,67 | 3592 |
| chrUextra | 14386991 | 14388121 | 1131 | 6,1 | 3593 |
| chrUextra | 14388142 | 14389881 | 1740 | 2,33 | 3594 |
| chrUextra | 14410236 | 14411228 | 993 | 2,69 | 3595 |
| chrUextra | 14411875 | 14413457 | 1583 | 3,3 | 3596 |
| chrUextra | 14415907 | 14417007 | 1101 | 3,37 | 3597 |

| | | | | | |
|---|---|---|---|---|---|
| chrUextra | 14419850 | 14420942 | 1093 | 3,16 | 3598 |
| chrUextra | 14440366 | 14441628 | 1263 | 3,87 | 3599 |
| chrUextra | 14444714 | 14445682 | 969 | 2,62 | 3600 |
| chrUextra | 14465186 | 14466289 | 1104 | 2,49 | 3601 |
| chrUextra | 14503864 | 14505693 | 1830 | 2,49 | 3602 |
| chrUextra | 14511105 | 14512541 | 1437 | 2,56 | 3603 |
| chrUextra | 14514453 | 14515636 | 1184 | 2,77 | 3604 |
| chrUextra | 14526622 | 14527707 | 1086 | 4,18 | 3605 |
| chrUextra | 14531663 | 14532697 | 1035 | 2,46 | 3606 |
| chrUextra | 14547097 | 14548616 | 1520 | 2,65 | 3607 |
| chrUextra | 14561209 | 14562958 | 1750 | 2,36 | 3608 |
| chrUextra | 14567745 | 14568857 | 1113 | 2,82 | 3609 |
| chrUextra | 14569347 | 14570439 | 1093 | 2,49 | 3610 |
| chrUextra | 14570933 | 14573422 | 2490 | 2,62 | 3611 |
| chrUextra | 14577591 | 14579363 | 1773 | 3,04 | 3612 |
| chrUextra | 14615919 | 14618868 | 2950 | 3,33 | 3613 |
| chrUextra | 14619233 | 14620338 | 1106 | 2,82 | 3614 |
| chrUextra | 14643913 | 14644971 | 1059 | 2,36 | 3615 |
| chrUextra | 14662723 | 14663770 | 1048 | 2,44 | 3616 |
| chrUextra | 14677260 | 14678326 | 1067 | 2,87 | 3617 |
| chrUextra | 14719220 | 14721664 | 2445 | 3,89 | 3618 |
| chrUextra | 14726425 | 14729287 | 2863 | 3,76 | 3619 |
| chrUextra | 14759402 | 14761159 | 1758 | 2,62 | 3620 |
| chrUextra | 14767489 | 14768516 | 1028 | 2,59 | 3621 |
| chrUextra | 14769962 | 14771013 | 1052 | 3,37 | 3622 |
| chrUextra | 14774018 | 14775062 | 1045 | 3,29 | 3623 |
| chrUextra | 14779029 | 14780778 | 1750 | 2,49 | 3624 |
| chrUextra | 14791997 | 14793085 | 1089 | 2,26 | 3625 |
| chrUextra | 14793664 | 14794736 | 1073 | 2,98 | 3626 |
| chrUextra | 14800216 | 14801350 | 1135 | 3,46 | 3627 |
| chrUextra | 14815729 | 14817476 | 1748 | 3,42 | 3628 |
| chrUextra | 14823150 | 14824843 | 1694 | 2,56 | 3629 |
| chrUextra | 14828795 | 14829844 | 1050 | 2,72 | 3630 |
| chrUextra | 14833762 | 14834634 | 873 | 4,21 | 3631 |
| chrUextra | 14845358 | 14846993 | 1636 | 2,17 | 3632 |
| chrUextra | 14870302 | 14871543 | 1242 | 4,05 | 3633 |
| chrUextra | 14897440 | 14898491 | 1052 | 2,56 | 3634 |
| chrUextra | 14943105 | 14944349 | 1245 | 3,05 | 3635 |
| chrUextra | 14956348 | 14957983 | 1636 | 2,2 | 3636 |
| chrUextra | 14962849 | 14964724 | 1876 | 2,78 | 3637 |
| chrUextra | 14974240 | 14975277 | 1038 | 2,52 | 3638 |
| chrUextra | 15002679 | 15004024 | 1346 | 3,92 | 3639 |
| chrUextra | 15029655 | 15030974 | 1320 | 4,15 | 3640 |
| chrUextra | 15035372 | 15036674 | 1303 | 3,55 | 3641 |
| chrUextra | 15037718 | 15038924 | 1207 | 3,12 | 3642 |
| chrUextra | 15071386 | 15073240 | 1855 | 2,46 | 3643 |
| chrUextra | 15079331 | 15080699 | 1369 | 3,94 | 3644 |
| chrUextra | 15087768 | 15089611 | 1844 | 2,79 | 3645 |

| | | | | | |
|---|---|---|---|---|---|
| chrUextra | 15094196 | 15095196 | 1001 | 2,94 | 3646 |
| chrUextra | 15098308 | 15099191 | 884 | 2,49 | 3647 |
| chrUextra | 15125030 | 15128194 | 3165 | 2,91 | 3648 |
| chrUextra | 15132531 | 15135142 | 2612 | 3,4 | 3649 |
| chrUextra | 15150450 | 15151440 | 991 | 2,85 | 3650 |
| chrUextra | 15161742 | 15162878 | 1137 | 3,41 | 3651 |
| chrUextra | 15177431 | 15179211 | 1781 | 2,75 | 3652 |
| chrUextra | 15189621 | 15191404 | 1784 | 2,26 | 3653 |
| chrUextra | 15193678 | 15195162 | 1485 | 2,62 | 3654 |
| chrUextra | 15201608 | 15202850 | 1243 | 3,6 | 3655 |
| chrUextra | 15204239 | 15205231 | 993 | 3,17 | 3656 |
| chrUextra | 15216358 | 15217668 | 1311 | 3,52 | 3657 |
| chrUextra | 15219729 | 15220826 | 1098 | 2,78 | 3658 |
| chrUextra | 15243289 | 15244294 | 1006 | 2,88 | 3659 |
| chrUextra | 15257061 | 15258329 | 1269 | 2,82 | 3660 |
| chrUextra | 15273591 | 15275494 | 1904 | 2,79 | 3661 |
| chrUextra | 15290660 | 15291723 | 1064 | 3,08 | 3662 |
| chrUextra | 15297838 | 15298969 | 1132 | 2,85 | 3663 |
| chrUextra | 15306665 | 15308086 | 1422 | 2,73 | 3664 |
| chrUextra | 15315792 | 15317347 | 1556 | 2,69 | 3665 |
| chrUextra | 15324740 | 15326602 | 1863 | 3,56 | 3666 |
| chrUextra | 15391501 | 15394116 | 2616 | 2,49 | 3667 |
| chrUextra | 15421629 | 15422664 | 1036 | 2,65 | 3668 |
| chrUextra | 15447550 | 15448550 | 1001 | 3,3 | 3669 |
| chrUextra | 15458010 | 15459339 | 1330 | 4,54 | 3670 |
| chrUextra | 15459628 | 15462213 | 2586 | 3,02 | 3671 |
| chrUextra | 15474462 | 15476197 | 1736 | 3,5 | 3672 |
| chrUextra | 15481597 | 15482906 | 1310 | 4,01 | 3673 |
| chrUextra | 15485630 | 15486837 | 1208 | 3,29 | 3674 |
| chrUextra | 15488194 | 15489197 | 1004 | 2,65 | 3675 |
| chrUextra | 15493014 | 15494134 | 1121 | 3,27 | 3676 |
| chrUextra | 15502808 | 15503824 | 1017 | 2,91 | 3677 |
| chrUextra | 15504434 | 15505448 | 1015 | 2,56 | 3678 |
| chrUextra | 15508537 | 15509562 | 1026 | 3,04 | 3679 |
| chrUextra | 15519002 | 15520079 | 1078 | 3,3 | 3680 |
| chrUextra | 15523879 | 15524933 | 1055 | 2,36 | 3681 |
| chrUextra | 15534348 | 15535589 | 1242 | 3,75 | 3682 |
| chrUextra | 15535962 | 15539440 | 3479 | 3,25 | 3683 |
| chrUextra | 15544238 | 15546313 | 2076 | 2,39 | 3684 |
| chrUextra | 15553301 | 15555703 | 2403 | 2,33 | 3685 |
| chrUextra | 15561184 | 15562497 | 1314 | 3,85 | 3686 |
| chrUextra | 15572407 | 15573053 | 647 | 3,75 | 3687 |
| chrUextra | 15577615 | 15578640 | 1026 | 3,04 | 3688 |
| chrUextra | 15584032 | 15585754 | 1723 | 2,46 | 3689 |
| chrUextra | 15598557 | 15599712 | 1156 | 2,59 | 3690 |
| chrUextra | 15600254 | 15601299 | 1046 | 2,59 | 3691 |
| chrUextra | 15601673 | 15602889 | 1217 | 3,1 | 3692 |
| chrUextra | 15606699 | 15607781 | 1083 | 3,43 | 3693 |

| | | | | | |
|---|---|---|---|---|---|
| chrUextra | 15615702 | 15616687 | 986 | 2,44 | 3694 |
| chrUextra | 15632067 | 15633721 | 1655 | 2,69 | 3695 |
| chrUextra | 15651487 | 15653993 | 2507 | 2,75 | 3696 |
| chrUextra | 15679780 | 15680811 | 1032 | 2,65 | 3697 |
| chrUextra | 15696024 | 15697009 | 986 | 2,85 | 3698 |
| chrUextra | 15706544 | 15707627 | 1084 | 2,69 | 3699 |
| chrUextra | 15726599 | 15728427 | 1829 | 3,84 | 3700 |
| chrUextra | 15747916 | 15750045 | 2130 | 2,62 | 3701 |
| chrUextra | 15763281 | 15764335 | 1055 | 2,46 | 3702 |
| chrUextra | 15768191 | 15769848 | 1658 | 2,59 | 3703 |
| chrUextra | 15773951 | 15775676 | 1726 | 2,56 | 3704 |
| chrUextra | 15806246 | 15808075 | 1830 | 2,56 | 3705 |
| chrUextra | 15840294 | 15841416 | 1123 | 2,52 | 3706 |
| chrUextra | 15841801 | 15843485 | 1685 | 2,99 | 3707 |
| chrUextra | 15867753 | 15869008 | 1256 | 3,2 | 3708 |
| chrUextra | 15874315 | 15875373 | 1059 | 2,94 | 3709 |
| chrUextra | 15886361 | 15887599 | 1239 | 3 | 3710 |
| chrUextra | 15899391 | 15900402 | 1012 | 2,49 | 3711 |
| chrUextra | 15906776 | 15907965 | 1190 | 3,49 | 3712 |
| chrUextra | 15908948 | 15910318 | 1371 | 3,07 | 3713 |
| chrUextra | 15913296 | 15915206 | 1911 | 3,19 | 3714 |
| chrUextra | 15946403 | 15947439 | 1037 | 3,04 | 3715 |
| chrUextra | 15956049 | 15957133 | 1085 | 2,62 | 3716 |
| chrUextra | 15958526 | 15959500 | 975 | 2,52 | 3717 |
| chrUextra | 15960895 | 15962053 | 1159 | 2,73 | 3718 |
| chrUextra | 15975352 | 15977208 | 1857 | 2,88 | 3719 |
| chrUextra | 15995775 | 15996940 | 1166 | 6,99 | 3720 |
| chrUextra | 16002306 | 16004438 | 2133 | 2,36 | 3721 |
| chrUextra | 16010111 | 16011316 | 1206 | 3,51 | 3722 |
| chrUextra | 16011880 | 16013779 | 1900 | 3,1 | 3723 |
| chrUextra | 16015919 | 16016919 | 1001 | 2,36 | 3724 |
| chrUextra | 16023203 | 16024930 | 1728 | 3,46 | 3725 |
| chrUextra | 16041029 | 16043812 | 2784 | 3,54 | 3726 |
| chrUextra | 16063453 | 16064464 | 1012 | 3,79 | 3727 |
| chrUextra | 16074176 | 16075188 | 1013 | 2,3 | 3728 |
| chrUextra | 16081511 | 16083239 | 1729 | 2,36 | 3729 |
| chrUextra | 16140393 | 16141538 | 1146 | 2,31 | 3730 |
| chrUextra | 16155039 | 16156528 | 1490 | 2,43 | 3731 |
| chrUextra | 16163927 | 16166390 | 2464 | 3,51 | 3732 |
| chrUextra | 16171825 | 16172954 | 1130 | 3,16 | 3733 |
| chrUextra | 16188905 | 16189879 | 975 | 2,36 | 3734 |
| chrUextra | 16198621 | 16199593 | 973 | 2,56 | 3735 |
| chrUextra | 16209042 | 16210084 | 1043 | 2,52 | 3736 |
| chrUextra | 16230923 | 16231860 | 938 | 2,46 | 3737 |
| chrUextra | 16232414 | 16234312 | 1899 | 2,85 | 3738 |
| chrUextra | 16241366 | 16242567 | 1202 | 2,93 | 3739 |
| chrUextra | 16242939 | 16244003 | 1065 | 2,91 | 3740 |
| chrUextra | 16254181 | 16255360 | 1180 | 3,45 | 3741 |

| | | | | | |
|---|---|---|---|---|---|
| chrUextra | 16272943 | 16275340 | 2398 | 2,26 | 3742 |
| chrUextra | 16279464 | 16281169 | 1706 | 4,14 | 3743 |
| chrUextra | 16338282 | 16339211 | 930 | 2,17 | 3744 |
| chrUextra | 16351909 | 16352913 | 1005 | 2,39 | 3745 |
| chrUextra | 16381515 | 16382805 | 1291 | 2,49 | 3746 |
| chrUextra | 16409697 | 16411225 | 1529 | 3,52 | 3747 |
| chrUextra | 16417816 | 16419122 | 1307 | 2,83 | 3748 |
| chrUextra | 16431483 | 16432733 | 1251 | 3,6 | 3749 |
| chrUextra | 16434952 | 16435937 | 986 | 2,23 | 3750 |
| chrUextra | 16437352 | 16438296 | 945 | 3,59 | 3751 |
| chrUextra | 16447093 | 16448780 | 1688 | 2,49 | 3752 |
| chrUextra | 16471846 | 16473005 | 1160 | 3,45 | 3753 |
| chrUextra | 16481672 | 16482716 | 1045 | 2,78 | 3754 |
| chrUextra | 16483239 | 16484222 | 984 | 2,88 | 3755 |
| chrUextra | 16517911 | 16518925 | 1015 | 2,3 | 3756 |
| chrUextra | 16519391 | 16520522 | 1132 | 4,01 | 3757 |
| chrUextra | 16547600 | 16549393 | 1794 | 2,91 | 3758 |
| chrUextra | 16552336 | 16553647 | 1312 | 3,49 | 3759 |
| chrUextra | 16567702 | 16568823 | 1122 | 3,56 | 3760 |
| chrUextra | 16588899 | 16590478 | 1580 | 2,14 | 3761 |
| chrUextra | 16624902 | 16626096 | 1195 | 2,99 | 3762 |
| chrUextra | 16633148 | 16634851 | 1704 | 2,75 | 3763 |
| chrUextra | 16645883 | 16646934 | 1052 | 2,3 | 3764 |
| chrUextra | 16651595 | 16653391 | 1797 | 2,98 | 3765 |
| chrUextra | 16678634 | 16679297 | 664 | 3,53 | 3766 |
| chrUextra | 16706251 | 16707614 | 1364 | 2,75 | 3767 |
| chrUextra | 16717502 | 16719232 | 1731 | 2,49 | 3768 |
| chrUextra | 16739819 | 16741793 | 1975 | 3,19 | 3769 |
| chrUextra | 16744827 | 16746378 | 1552 | 2,43 | 3770 |
| chrUextra | 16760772 | 16762715 | 1944 | 3,24 | 3771 |
| chrUextra | 16765624 | 16766845 | 1222 | 3,64 | 3772 |
| chrUextra | 16780125 | 16781235 | 1111 | 3,43 | 3773 |
| chrUextra | 16786598 | 16788423 | 1826 | 4,24 | 3774 |
| chrUextra | 16791398 | 16792437 | 1040 | 2,2 | 3775 |
| chrUextra | 16797022 | 16799752 | 2731 | 3,51 | 3776 |
| chrUextra | 16800055 | 16801761 | 1707 | 3,7 | 3777 |
| chrUextra | 16803414 | 16804650 | 1237 | 3,52 | 3778 |
| chrUextra | 16809847 | 16811731 | 1885 | 2,47 | 3779 |
| chrUextra | 16823589 | 16825185 | 1597 | 2,43 | 3780 |
| chrUextra | 16861226 | 16862434 | 1209 | 2,62 | 3781 |
| chrUextra | 16866676 | 16868020 | 1345 | 3,23 | 3782 |
| chrUextra | 16891148 | 16892729 | 1582 | 2,65 | 3783 |
| chrUextra | 16899633 | 16900771 | 1139 | 3,16 | 3784 |
| chrUextra | 16913204 | 16914603 | 1400 | 4,43 | 3785 |
| chrUextra | 16942333 | 16943292 | 960 | 2,75 | 3786 |
| chrUextra | 16973441 | 16975365 | 1925 | 2,37 | 3787 |
| chrUextra | 16980005 | 16981062 | 1058 | 3,24 | 3788 |
| chrUextra | 16982206 | 16985097 | 2892 | 2,69 | 3789 |

| | | | | | |
|---|---|---|---|---|---|
| chrUextra | 17004087 | 17005784 | 1698 | 2,33 | 3790 |
| chrUextra | 17024109 | 17025105 | 997 | 2,82 | 3791 |
| chrUextra | 17025650 | 17027333 | 1684 | 5,21 | 3792 |
| chrUextra | 17029736 | 17030709 | 974 | 2,14 | 3793 |
| chrUextra | 17035268 | 17036243 | 976 | 3,33 | 3794 |
| chrUextra | 17051532 | 17053324 | 1793 | 3,74 | 3795 |
| chrUextra | 17071388 | 17073109 | 1722 | 2,59 | 3796 |
| chrUextra | 17078670 | 17080273 | 1604 | 2,39 | 3797 |
| chrUextra | 17084038 | 17085346 | 1309 | 3,29 | 3798 |
| chrUextra | 17100030 | 17101526 | 1497 | 3,04 | 3799 |
| chrUextra | 17103214 | 17104623 | 1410 | 2,95 | 3800 |
| chrUextra | 17105844 | 17107582 | 1739 | 2,88 | 3801 |
| chrUextra | 17111460 | 17112437 | 978 | 2,75 | 3802 |
| chrUextra | 17141081 | 17142046 | 966 | 3,05 | 3803 |
| chrUextra | 17144347 | 17146084 | 1738 | 2,91 | 3804 |
| chrUextra | 17156317 | 17157287 | 971 | 2,46 | 3805 |
| chrUextra | 17160110 | 17162065 | 1956 | 3,47 | 3806 |
| chrUextra | 17223510 | 17224483 | 974 | 2,39 | 3807 |
| chrUextra | 17237939 | 17239340 | 1402 | 2,56 | 3808 |
| chrUextra | 17266777 | 17268637 | 1861 | 4,24 | 3809 |
| chrUextra | 17289931 | 17290958 | 1028 | 3,04 | 3810 |
| chrUextra | 17297166 | 17299168 | 2003 | 5,12 | 3811 |
| chrUextra | 17301956 | 17303676 | 1721 | 2,33 | 3812 |
| chrUextra | 17325332 | 17326800 | 1469 | 2,2 | 3813 |
| chrUextra | 17373290 | 17374425 | 1136 | 5,79 | 3814 |
| chrUextra | 17384294 | 17385958 | 1665 | 2,43 | 3815 |
| chrUextra | 17398017 | 17399619 | 1603 | 2,26 | 3816 |
| chrUextra | 17400143 | 17400852 | 710 | 3,43 | 3817 |
| chrUextra | 17402698 | 17405311 | 2614 | 2,38 | 3818 |
| chrUextra | 17412105 | 17413492 | 1388 | 3,55 | 3819 |
| chrUextra | 17420994 | 17422065 | 1072 | 2,46 | 3820 |
| chrUextra | 17422544 | 17425229 | 2686 | 2,61 | 3821 |
| chrUextra | 17429006 | 17430021 | 1016 | 2,36 | 3822 |
| chrUextra | 17458425 | 17460161 | 1737 | 2,77 | 3823 |
| chrUextra | 17469810 | 17471405 | 1596 | 3,14 | 3824 |
| chrUextra | 17485664 | 17486666 | 1003 | 2,39 | 3825 |
| chrUextra | 17494675 | 17496304 | 1630 | 2,36 | 3826 |
| chrUextra | 17501854 | 17502943 | 1090 | 2,9 | 3827 |
| chrUextra | 17516108 | 17517885 | 1778 | 2,55 | 3828 |
| chrUextra | 17518074 | 17518840 | 767 | 5,21 | 3829 |
| chrUextra | 17519129 | 17520994 | 1866 | 3,95 | 3830 |
| chrUextra | 17526471 | 17527986 | 1516 | 3,23 | 3831 |
| chrUextra | 17538427 | 17539480 | 1054 | 2,39 | 3832 |
| chrUextra | 17542483 | 17544231 | 1749 | 2,82 | 3833 |
| chrUextra | 17546551 | 17548101 | 1551 | 2,62 | 3834 |
| chrUextra | 17564774 | 17566471 | 1698 | 2,2 | 3835 |
| chrUextra | 17574266 | 17575400 | 1135 | 3,58 | 3836 |
| chrUextra | 17585536 | 17586512 | 977 | 2,69 | 3837 |

| | | | | | |
|---|---|---|---|---|---|
| chrUextra | 17588742 | 17589690 | 949 | 2,85 | 3838 |
| chrUextra | 17591014 | 17592815 | 1802 | 3,11 | 3839 |
| chrUextra | 17608643 | 17609834 | 1192 | 3,51 | 3840 |
| chrUextra | 17615097 | 17616700 | 1604 | 2,36 | 3841 |
| chrUextra | 17621426 | 17622701 | 1276 | 2,67 | 3842 |
| chrUextra | 17651597 | 17652698 | 1102 | 2,62 | 3843 |
| chrUextra | 17670154 | 17671980 | 1827 | 2,88 | 3844 |
| chrUextra | 17679637 | 17680728 | 1092 | 2,82 | 3845 |
| chrUextra | 17684516 | 17687047 | 2532 | 2,59 | 3846 |
| chrUextra | 17688342 | 17690188 | 1847 | 3,41 | 3847 |
| chrUextra | 17714051 | 17715660 | 1610 | 2,46 | 3848 |
| chrUextra | 17722596 | 17723655 | 1060 | 2,46 | 3849 |
| chrUextra | 17726662 | 17728477 | 1816 | 2,36 | 3850 |
| chrUextra | 17733731 | 17734989 | 1259 | 3,46 | 3851 |
| chrUextra | 17756784 | 17758096 | 1313 | 3,33 | 3852 |
| chrUextra | 17785547 | 17786613 | 1067 | 3,11 | 3853 |
| chrUextra | 17806420 | 17808131 | 1712 | 2,82 | 3854 |
| chrUextra | 17835745 | 17838255 | 2511 | 3,82 | 3855 |
| chrUextra | 17838887 | 17840660 | 1774 | 3,11 | 3856 |
| chrUextra | 17841355 | 17843146 | 1792 | 2,36 | 3857 |
| chrUextra | 17866935 | 17868407 | 1473 | 2,39 | 3858 |
| chrUextra | 17876381 | 17877936 | 1556 | 2,52 | 3859 |
| chrUextra | 17882016 | 17883105 | 1090 | 3,4 | 3860 |
| chrUextra | 17899351 | 17900315 | 965 | 3,27 | 3861 |
| chrUextra | 17901057 | 17903628 | 2572 | 2,56 | 3862 |
| chrUextra | 17916189 | 17917165 | 977 | 2,52 | 3863 |
| chrUextra | 17929571 | 17930256 | 686 | 3,69 | 3864 |
| chrUextra | 17933734 | 17934660 | 927 | 2,43 | 3865 |
| chrUextra | 17939101 | 17940400 | 1300 | 3,37 | 3866 |
| chrUextra | 17944606 | 17945875 | 1270 | 3,67 | 3867 |
| chrUextra | 17960688 | 17961734 | 1047 | 2,75 | 3868 |
| chrUextra | 17971908 | 17974477 | 2570 | 3,63 | 3869 |
| chrUextra | 17985172 | 17985916 | 745 | 5,18 | 3870 |
| chrUextra | 18012422 | 18014219 | 1798 | 2,59 | 3871 |
| chrUextra | 18018034 | 18019690 | 1657 | 7,34 | 3872 |
| chrUextra | 18026756 | 18028461 | 1706 | 6,54 | 3873 |
| chrUextra | 18039407 | 18040459 | 1053 | 2,39 | 3874 |
| chrUextra | 18041861 | 18042783 | 923 | 2,39 | 3875 |
| chrUextra | 18055412 | 18057903 | 2492 | 2,23 | 3876 |
| chrUextra | 18067901 | 18068660 | 760 | 4,92 | 3877 |
| chrUextra | 18083063 | 18084831 | 1769 | 3,34 | 3878 |
| chrUextra | 18093440 | 18094500 | 1061 | 2,36 | 3879 |
| chrUextra | 18111824 | 18113446 | 1623 | 2,36 | 3880 |
| chrUextra | 18118961 | 18120095 | 1135 | 7,64 | 3881 |
| chrUextra | 18122012 | 18123024 | 1013 | 3,64 | 3882 |
| chrUextra | 18126843 | 18127749 | 907 | 2,72 | 3883 |
| chrUextra | 18149146 | 18150567 | 1422 | 2,46 | 3884 |
| chrUextra | 18162728 | 18164286 | 1559 | 3,29 | 3885 |

| | | | | | |
|---|---|---|---|---|---|
| chrUextra | 18186446 | 18188114 | 1669 | 2,33 | 3886 |
| chrUextra | 18188703 | 18190511 | 1809 | 2,36 | 3887 |
| chrUextra | 18194161 | 18194789 | 629 | 4,01 | 3888 |
| chrUextra | 18208762 | 18211058 | 2297 | 2,8 | 3889 |
| chrUextra | 18221117 | 18222305 | 1189 | 3,45 | 3890 |
| chrUextra | 18227575 | 18228638 | 1064 | 2,72 | 3891 |
| chrUextra | 18232389 | 18233320 | 932 | 2,49 | 3892 |
| chrUextra | 18250386 | 18251768 | 1383 | 3,16 | 3893 |
| chrUextra | 18279068 | 18280251 | 1184 | 3,56 | 3894 |
| chrUextra | 18285596 | 18286655 | 1060 | 3,24 | 3895 |
| chrUextra | 18295798 | 18296847 | 1050 | 2,56 | 3896 |
| chrUextra | 18300293 | 18301546 | 1254 | 2,91 | 3897 |
| chrUextra | 18308377 | 18310069 | 1693 | 2,69 | 3898 |
| chrUextra | 18319697 | 18321340 | 1644 | 2,9 | 3899 |
| chrUextra | 18326686 | 18328346 | 1661 | 2,36 | 3900 |
| chrUextra | 18337790 | 18339683 | 1894 | 2,62 | 3901 |
| chrUextra | 18362361 | 18363395 | 1035 | 2,82 | 3902 |
| chrUextra | 18398029 | 18399065 | 1037 | 2,59 | 3903 |
| chrUextra | 18418787 | 18420361 | 1575 | 3,01 | 3904 |
| chrUextra | 18421714 | 18422736 | 1023 | 3,3 | 3905 |
| chrUextra | 18424242 | 18426567 | 2326 | 2,46 | 3906 |
| chrUextra | 18430566 | 18432254 | 1689 | 4,38 | 3907 |
| chrUextra | 18436075 | 18437644 | 1570 | 2,39 | 3908 |
| chrUextra | 18516830 | 18518441 | 1612 | 2,33 | 3909 |
| chrUextra | 18534054 | 18535299 | 1246 | 4,19 | 3910 |
| chrUextra | 18539470 | 18540716 | 1247 | 4,66 | 3911 |
| chrUextra | 18541970 | 18543252 | 1283 | 4,02 | 3912 |
| chrUextra | 18553195 | 18554135 | 941 | 2,98 | 3913 |
| chrUextra | 18561324 | 18562857 | 1534 | 2,82 | 3914 |
| chrUextra | 18564328 | 18566020 | 1693 | 2,77 | 3915 |
| chrUextra | 18568027 | 18569161 | 1135 | 5,66 | 3916 |
| chrUextra | 18613550 | 18614455 | 906 | 3,92 | 3917 |
| chrUextra | 18625091 | 18626165 | 1075 | 2,65 | 3918 |
| chrUextra | 18629709 | 18631029 | 1321 | 3,54 | 3919 |
| chrUextra | 18646456 | 18647448 | 993 | 2,52 | 3920 |
| chrUextra | 18650243 | 18651744 | 1502 | 3,14 | 3921 |
| chrUextra | 18651969 | 18652972 | 1004 | 2,56 | 3922 |
| chrUextra | 18653713 | 18655295 | 1583 | 2,69 | 3923 |
| chrUextra | 18658441 | 18660087 | 1647 | 2,56 | 3924 |
| chrUextra | 18688226 | 18690674 | 2449 | 3,46 | 3925 |
| chrUextra | 18696956 | 18698049 | 1094 | 2,39 | 3926 |
| chrUextra | 18700009 | 18701108 | 1100 | 3,46 | 3927 |
| chrUextra | 18730096 | 18731311 | 1216 | 3,65 | 3928 |
| chrUextra | 18736545 | 18737534 | 990 | 2,43 | 3929 |
| chrUextra | 18799366 | 18800496 | 1131 | 2,39 | 3930 |
| chrUextra | 18811445 | 18814028 | 2584 | 3,62 | 3931 |
| chrUextra | 18819400 | 18820951 | 1552 | 3,03 | 3932 |
| chrUextra | 18821910 | 18823658 | 1749 | 2,78 | 3933 |

| | | | | | |
|---|---|---|---|---|---|
| chrUextra | 18828902 | 18830630 | 1729 | 2,91 | 3934 |
| chrUextra | 18840038 | 18841662 | 1625 | 3,27 | 3935 |
| chrUextra | 18854103 | 18855087 | 985 | 2,85 | 3936 |
| chrUextra | 18868200 | 18869627 | 1428 | 2,82 | 3937 |
| chrUextra | 18895147 | 18896086 | 940 | 2,39 | 3938 |
| chrUextra | 18903716 | 18904715 | 1000 | 4,59 | 3939 |
| chrUextra | 18921935 | 18923695 | 1761 | 2,82 | 3940 |
| chrUextra | 18930537 | 18931583 | 1047 | 2,78 | 3941 |
| chrUextra | 18935989 | 18938654 | 2666 | 3,69 | 3942 |
| chrUextra | 18942407 | 18944176 | 1770 | 2,33 | 3943 |
| chrUextra | 18968099 | 18969458 | 1360 | 3,34 | 3944 |
| chrUextra | 18976156 | 18977327 | 1172 | 2,91 | 3945 |
| chrUextra | 18978709 | 18980654 | 1946 | 3,05 | 3946 |
| chrUextra | 19012573 | 19013767 | 1195 | 2,83 | 3947 |
| chrUextra | 19029814 | 19030853 | 1040 | 2,78 | 3948 |
| chrUextra | 19033051 | 19034812 | 1762 | 2,46 | 3949 |
| chrUextra | 19038569 | 19040170 | 1602 | 2,11 | 3950 |
| chrUextra | 19046348 | 19048048 | 1701 | 3,41 | 3951 |
| chrUextra | 19064625 | 19067071 | 2447 | 3,04 | 3952 |
| chrUextra | 19073241 | 19074864 | 1624 | 2,73 | 3953 |
| chrUextra | 19109516 | 19111141 | 1626 | 2,3 | 3954 |
| chrUextra | 19146339 | 19147998 | 1660 | 2,23 | 3955 |
| chrUextra | 19166142 | 19167263 | 1122 | 4,63 | 3956 |
| chrUextra | 19173949 | 19175717 | 1769 | 2,49 | 3957 |
| chrUextra | 19181806 | 19183534 | 1729 | 2,69 | 3958 |
| chrUextra | 19185731 | 19187535 | 1805 | 2,12 | 3959 |
| chrUextra | 19215642 | 19217361 | 1720 | 2,88 | 3960 |
| chrUextra | 19221027 | 19222363 | 1337 | 3,72 | 3961 |
| chrUextra | 19241425 | 19242587 | 1163 | 2,46 | 3962 |
| chrUextra | 19270727 | 19272357 | 1631 | 2,39 | 3963 |
| chrUextra | 19289556 | 19292057 | 2502 | 2,75 | 3964 |
| chrUextra | 19292441 | 19293578 | 1138 | 3,35 | 3965 |
| chrUextra | 19339568 | 19340867 | 1300 | 3,41 | 3966 |
| chrUextra | 19463598 | 19464930 | 1333 | 3,47 | 3967 |
| chrUextra | 19475099 | 19475667 | 569 | 4,14 | 3968 |
| chrUextra | 19476667 | 19477471 | 805 | 3,33 | 3969 |
| chrUextra | 19486455 | 19491334 | 4880 | 2,75 | 3970 |
| chrUextra | 19499023 | 19500829 | 1807 | 2,26 | 3971 |
| chrUextra | 19501363 | 19502312 | 950 | 2,62 | 3972 |
| chrUextra | 19519351 | 19520968 | 1618 | 2,55 | 3973 |
| chrUextra | 19533513 | 19536112 | 2600 | 3,11 | 3974 |
| chrUextra | 19536629 | 19539563 | 2935 | 2,36 | 3975 |
| chrUextra | 19540252 | 19541550 | 1299 | 4,46 | 3976 |
| chrUextra | 19545256 | 19546976 | 1721 | 2,17 | 3977 |
| chrUextra | 19564267 | 19565835 | 1569 | 2,69 | 3978 |
| chrUextra | 19578945 | 19580030 | 1086 | 2,75 | 3979 |
| chrUextra | 19582861 | 19583875 | 1015 | 2,98 | 3980 |
| chrUextra | 19587565 | 19589233 | 1669 | 3,24 | 3981 |

| | | | | | |
|---|---|---|---|---|---|
| chrUextra | 19597723 | 19599483 | 1761 | 2,52 | 3982 |
| chrUextra | 19600975 | 19602635 | 1661 | 3,17 | 3983 |
| chrUextra | 19603746 | 19604530 | 785 | 4,92 | 3984 |
| chrUextra | 19621012 | 19621791 | 780 | 4,01 | 3985 |
| chrUextra | 19624918 | 19626286 | 1369 | 2,78 | 3986 |
| chrUextra | 19637677 | 19638685 | 1009 | 2,62 | 3987 |
| chrUextra | 19646237 | 19648908 | 2672 | 3,58 | 3988 |
| chrUextra | 19654904 | 19657211 | 2308 | 3,15 | 3989 |
| chrUextra | 19683062 | 19684835 | 1774 | 2,59 | 3990 |
| chrUextra | 19690252 | 19691815 | 1564 | 2,2 | 3991 |
| chrUextra | 19698693 | 19699726 | 1034 | 2,9 | 3992 |
| chrUextra | 19740255 | 19741851 | 1597 | 2,43 | 3993 |
| chrUextra | 19758187 | 19759939 | 1753 | 2,26 | 3994 |
| chrUextra | 19776164 | 19777115 | 952 | 2,59 | 3995 |
| chrUextra | 19821532 | 19822527 | 996 | 4,76 | 3996 |
| chrUextra | 19854306 | 19856065 | 1760 | 2,49 | 3997 |
| chrUextra | 19871490 | 19874146 | 2657 | 3,46 | 3998 |
| chrUextra | 19926935 | 19928706 | 1772 | 2,59 | 3999 |
| chrUextra | 19932433 | 19934223 | 1791 | 2,72 | 4000 |
| chrUextra | 19940192 | 19941215 | 1024 | 2,56 | 4001 |
| chrUextra | 19941799 | 19943466 | 1668 | 2,23 | 4002 |
| chrUextra | 19949757 | 19951776 | 2020 | 2,56 | 4003 |
| chrUextra | 19952062 | 19953697 | 1636 | 2,88 | 4004 |
| chrUextra | 19961965 | 19963280 | 1316 | 4,04 | 4005 |
| chrUextra | 19972239 | 19974014 | 1776 | 2,94 | 4006 |
| chrUextra | 19994695 | 19996585 | 1891 | 4,16 | 4007 |
| chrUextra | 20009688 | 20010662 | 975 | 2,26 | 4008 |
| chrUextra | 20013684 | 20014690 | 1007 | 4,17 | 4009 |
| chrUextra | 20017442 | 20018495 | 1054 | 3,27 | 4010 |
| chrUextra | 20027645 | 20028753 | 1109 | 2,72 | 4011 |
| chrUextra | 20045684 | 20047148 | 1465 | 3,66 | 4012 |
| chrUextra | 20067346 | 20068381 | 1036 | 2,46 | 4013 |
| chrUextra | 20071149 | 20072291 | 1143 | 3,17 | 4014 |
| chrUextra | 20072931 | 20074630 | 1700 | 2,78 | 4015 |
| chrUextra | 20095483 | 20097184 | 1702 | 2,52 | 4016 |
| chrUextra | 20100930 | 20101941 | 1012 | 3,43 | 4017 |
| chrUextra | 20122923 | 20124535 | 1613 | 3,14 | 4018 |
| chrUextra | 20126802 | 20128410 | 1609 | 2,62 | 4019 |
| chrUextra | 20133831 | 20135361 | 1531 | 2,78 | 4020 |
| chrUextra | 20206123 | 20207505 | 1383 | 2,06 | 4021 |
| chrUextra | 20209772 | 20210904 | 1133 | 6,01 | 4022 |
| chrUextra | 20234663 | 20235893 | 1231 | 3,28 | 4023 |
| chrUextra | 20269980 | 20271570 | 1591 | 2,2 | 4024 |
| chrUextra | 20286076 | 20287396 | 1321 | 3,53 | 4025 |
| chrUextra | 20292562 | 20293420 | 859 | 2,69 | 4026 |
| chrUextra | 20305809 | 20307484 | 1676 | 2,26 | 4027 |
| chrUextra | 20320407 | 20321519 | 1113 | 2,94 | 4028 |
| chrUextra | 20337578 | 20338623 | 1046 | 2,75 | 4029 |

| | | | | | |
|---|---|---|---|---|---|
| chrUextra | 20415821 | 20416403 | 583 | 4,24 | 4030 |
| chrUextra | 20423825 | 20426397 | 2573 | 2,75 | 4031 |
| chrUextra | 20432932 | 20433698 | 767 | 6,05 | 4032 |
| chrUextra | 20450236 | 20451209 | 974 | 2,62 | 4033 |
| chrUextra | 20458086 | 20458970 | 885 | 2,94 | 4034 |
| chrUextra | 20460271 | 20461378 | 1108 | 3,45 | 4035 |
| chrUextra | 20466542 | 20468270 | 1729 | 2,33 | 4036 |
| chrUextra | 20470525 | 20472207 | 1683 | 2,69 | 4037 |
| chrUextra | 20482856 | 20484525 | 1670 | 2,6 | 4038 |
| chrUextra | 20486079 | 20487548 | 1470 | 2,17 | 4039 |
| chrUextra | 20502964 | 20503990 | 1027 | 2,62 | 4040 |
| chrUextra | 20522422 | 20523416 | 995 | 2,94 | 4041 |
| chrUextra | 20548684 | 20550457 | 1774 | 3,2 | 4042 |
| chrUextra | 20553922 | 20555156 | 1235 | 5,04 | 4043 |
| chrUextra | 20568206 | 20569933 | 1728 | 2,33 | 4044 |
| chrUextra | 20586770 | 20587711 | 942 | 2,46 | 4045 |
| chrUextra | 20590748 | 20591810 | 1063 | 3,74 | 4046 |
| chrUextra | 20637940 | 20638948 | 1009 | 2,91 | 4047 |
| chrUextra | 20658136 | 20660055 | 1920 | 4,05 | 4048 |
| chrUextra | 20679716 | 20681580 | 1865 | 2,72 | 4049 |
| chrUextra | 20683709 | 20684731 | 1023 | 3,08 | 4050 |
| chrUextra | 20712248 | 20713343 | 1096 | 3,01 | 4051 |
| chrUextra | 20720167 | 20721787 | 1621 | 2,49 | 4052 |
| chrUextra | 20724666 | 20725740 | 1075 | 4,04 | 4053 |
| chrUextra | 20774978 | 20776303 | 1326 | 3,91 | 4054 |
| chrUextra | 20802758 | 20804047 | 1290 | 3,93 | 4055 |
| chrUextra | 20809156 | 20810705 | 1550 | 2,69 | 4056 |
| chrUextra | 20872122 | 20873080 | 959 | 4,79 | 4057 |
| chrUextra | 20873371 | 20874387 | 1017 | 2,46 | 4058 |
| chrUextra | 20880495 | 20882070 | 1576 | 2,56 | 4059 |
| chrUextra | 20883408 | 20885920 | 2513 | 2,43 | 4060 |
| chrUextra | 20892479 | 20893888 | 1410 | 3,29 | 4061 |
| chrUextra | 20915890 | 20917402 | 1513 | 3,07 | 4062 |
| chrUextra | 20933577 | 20935323 | 1747 | 3,49 | 4063 |
| chrUextra | 20937535 | 20938503 | 969 | 2,75 | 4064 |
| chrUextra | 20938905 | 20940461 | 1557 | 3,95 | 4065 |
| chrUextra | 20942966 | 20944392 | 1427 | 2,56 | 4066 |
| chrUextra | 21008410 | 21009566 | 1157 | 3,98 | 4067 |
| chrUextra | 21016934 | 21018190 | 1257 | 4,57 | 4068 |
| chrUextra | 21049511 | 21051192 | 1682 | 2,39 | 4069 |
| chrUextra | 21068845 | 21071187 | 2343 | 2,78 | 4070 |
| chrUextra | 21080427 | 21081963 | 1537 | 2,72 | 4071 |
| chrUextra | 21108100 | 21109234 | 1135 | 3,33 | 4072 |
| chrUextra | 21109731 | 21110693 | 963 | 2,43 | 4073 |
| chrUextra | 21113645 | 21115245 | 1601 | 2,49 | 4074 |
| chrUextra | 21125864 | 21126812 | 949 | 2,78 | 4075 |
| chrUextra | 21134500 | 21136265 | 1766 | 2,9 | 4076 |
| chrUextra | 21143713 | 21145557 | 1845 | 3,17 | 4077 |

| | | | | | |
|---|---|---|---|---|---|
| chrUextra | 21150570 | 21152335 | 1766 | 2,62 | 4078 |
| chrUextra | 21155210 | 21156191 | 982 | 2,49 | 4079 |
| chrUextra | 21185167 | 21186251 | 1085 | 2,98 | 4080 |
| chrUextra | 21223627 | 21224854 | 1228 | 3,34 | 4081 |
| chrUextra | 21268626 | 21270127 | 1502 | 2,49 | 4082 |
| chrUextra | 21277769 | 21278830 | 1062 | 3,3 | 4083 |
| chrUextra | 21294510 | 21295641 | 1132 | 4,46 | 4084 |
| chrUextra | 21312478 | 21313955 | 1478 | 2,23 | 4085 |
| chrUextra | 21355505 | 21356572 | 1068 | 2,56 | 4086 |
| chrUextra | 21377900 | 21379630 | 1731 | 2,33 | 4087 |
| chrUextra | 21384880 | 21386389 | 1510 | 3,27 | 4088 |
| chrUextra | 21387689 | 21390277 | 2589 | 3,2 | 4089 |
| chrUextra | 21402481 | 21403518 | 1038 | 2,46 | 4090 |
| chrUextra | 21411849 | 21413379 | 1531 | 2,36 | 4091 |
| chrUextra | 21415543 | 21419385 | 3843 | 3,11 | 4092 |
| chrUextra | 21443141 | 21444215 | 1075 | 3,37 | 4093 |
| chrUextra | 21446274 | 21447265 | 992 | 3,08 | 4094 |
| chrUextra | 21463953 | 21464899 | 947 | 3,04 | 4095 |
| chrUextra | 21472450 | 21473460 | 1011 | 2,91 | 4096 |
| chrUextra | 21483789 | 21484947 | 1159 | 2,46 | 4097 |
| chrUextra | 21487002 | 21488025 | 1024 | 3,56 | 4098 |
| chrUextra | 21503199 | 21504856 | 1658 | 2,69 | 4099 |
| chrUextra | 21505588 | 21507312 | 1725 | 2,91 | 4100 |
| chrUextra | 21539869 | 21540938 | 1070 | 4,78 | 4101 |
| chrUextra | 21543997 | 21545635 | 1639 | 3,88 | 4102 |
| chrUextra | 21546237 | 21547721 | 1485 | 3,43 | 4103 |
| chrUextra | 21560026 | 21561103 | 1078 | 2,56 | 4104 |
| chrUextra | 21567925 | 21570209 | 2285 | 4,18 | 4105 |
| chrUextra | 21615321 | 21617059 | 1739 | 2,59 | 4106 |
| chrUextra | 21625207 | 21626353 | 1147 | 3,11 | 4107 |
| chrUextra | 21632960 | 21633955 | 996 | 2,46 | 4108 |
| chrUextra | 21636793 | 21639926 | 3134 | 2,69 | 4109 |
| chrUextra | 21681282 | 21682284 | 1003 | 2,36 | 4110 |
| chrUextra | 21682874 | 21684256 | 1383 | 2,49 | 4111 |
| chrUextra | 21686681 | 21688363 | 1683 | 2,36 | 4112 |
| chrUextra | 21700193 | 21701288 | 1096 | 6,28 | 4113 |
| chrUextra | 21728197 | 21730586 | 2390 | 3,1 | 4114 |
| chrUextra | 21732555 | 21733626 | 1072 | 4,21 | 4115 |
| chrUextra | 21734294 | 21735962 | 1669 | 1,97 | 4116 |
| chrUextra | 21773400 | 21775067 | 1668 | 2,62 | 4117 |
| chrUextra | 21780182 | 21781862 | 1681 | 2,69 | 4118 |
| chrUextra | 21800044 | 21801155 | 1112 | 2,49 | 4119 |
| chrUextra | 21806228 | 21807215 | 988 | 2,62 | 4120 |
| chrUextra | 21814546 | 21815575 | 1030 | 4,25 | 4121 |
| chrUextra | 21816289 | 21818084 | 1796 | 3,47 | 4122 |
| chrUextra | 21849841 | 21850890 | 1050 | 2,75 | 4123 |
| chrUextra | 21861276 | 21862385 | 1110 | 3,08 | 4124 |
| chrUextra | 21869922 | 21871522 | 1601 | 2,52 | 4125 |

| | | | | | |
|---|---|---|---|---|---|
| chrUextra | 21874304 | 21875360 | 1057 | 3,04 | 4126 |
| chrUextra | 21891559 | 21892433 | 875 | 4,08 | 4127 |
| chrUextra | 21907889 | 21908912 | 1024 | 6,31 | 4128 |
| chrUextra | 21968484 | 21969523 | 1040 | 2,95 | 4129 |
| chrUextra | 21974655 | 21976314 | 1660 | 2,33 | 4130 |
| chrUextra | 21990569 | 21991800 | 1232 | 3,21 | 4131 |
| chrUextra | 21993630 | 21994824 | 1195 | 4,49 | 4132 |
| chrUextra | 21998451 | 21999987 | 1537 | 2,26 | 4133 |
| chrUextra | 22020399 | 22021409 | 1011 | 3,69 | 4134 |
| chrUextra | 22027347 | 22028335 | 989 | 3,04 | 4135 |
| chrUextra | 22034808 | 22035848 | 1041 | 6,12 | 4136 |
| chrUextra | 22042437 | 22043493 | 1057 | 4,5 | 4137 |
| chrUextra | 22056384 | 22057725 | 1342 | 4,33 | 4138 |
| chrUextra | 22069039 | 22070159 | 1121 | 4,14 | 4139 |
| chrUextra | 22079773 | 22080771 | 999 | 4,72 | 4140 |
| chrUextra | 22080837 | 22081789 | 953 | 3,72 | 4141 |
| chrUextra | 22084486 | 22085662 | 1177 | 3,98 | 4142 |
| chrUextra | 22152716 | 22154147 | 1432 | 5,69 | 4143 |
| chrUextra | 22164199 | 22165329 | 1131 | 5,54 | 4144 |
| chrUextra | 22174564 | 22175375 | 812 | 5,27 | 4145 |
| chrUextra | 22178302 | 22179650 | 1349 | 5,19 | 4146 |
| chrUextra | 22261685 | 22262645 | 961 | 2,56 | 4147 |
| chrUextra | 22263264 | 22264875 | 1612 | 2,59 | 4148 |
| chrUextra | 22267790 | 22268780 | 991 | 2,69 | 4149 |
| chrUextra | 22278129 | 22278979 | 851 | 3,4 | 4150 |
| chrUextra | 22283842 | 22284804 | 963 | 3,11 | 4151 |
| chrUextra | 22286145 | 22287096 | 952 | 2,59 | 4152 |
| chrUextra | 22296812 | 22298578 | 1767 | 2,43 | 4153 |
| chrUextra | 22304540 | 22306803 | 2264 | 3,58 | 4154 |
| chrUextra | 22317946 | 22319215 | 1270 | 5,47 | 4155 |
| chrUextra | 22336357 | 22338150 | 1794 | 2,43 | 4156 |
| chrUextra | 22357731 | 22359366 | 1636 | 2,62 | 4157 |
| chrUextra | 22386156 | 22389123 | 2968 | 2,33 | 4158 |
| chrUextra | 22389759 | 22390751 | 993 | 2,36 | 4159 |
| chrUextra | 22413364 | 22414915 | 1552 | 2,07 | 4160 |
| chrUextra | 22440096 | 22440944 | 849 | 3,45 | 4161 |
| chrUextra | 22459329 | 22459971 | 643 | 3,17 | 4162 |
| chrUextra | 22490216 | 22491931 | 1716 | 2,46 | 4163 |
| chrUextra | 22541153 | 22542766 | 1614 | 2,56 | 4164 |
| chrUextra | 22558637 | 22560324 | 1688 | 2,36 | 4165 |
| chrUextra | 22582956 | 22583963 | 1008 | 2,33 | 4166 |
| chrUextra | 22594404 | 22595400 | 997 | 3,04 | 4167 |
| chrUextra | 22615637 | 22616588 | 952 | 2,52 | 4168 |
| chrUextra | 22646721 | 22647704 | 984 | 2,59 | 4169 |
| chrUextra | 22709745 | 22710719 | 975 | 2,56 | 4170 |
| chrUextra | 22719031 | 22720038 | 1008 | 5,02 | 4171 |
| chrUextra | 22720281 | 22721384 | 1104 | 2,56 | 4172 |
| chrUextra | 22732506 | 22733464 | 959 | 2,56 | 4173 |

| | | | | | |
|---|---|---|---|---|---|
| chrUextra | 22733826 | 22735709 | 1884 | 2,83 | 4174 |
| chrUextra | 22776481 | 22778107 | 1627 | 2,23 | 4175 |
| chrUextra | 22787056 | 22788076 | 1021 | 2,62 | 4176 |
| chrUextra | 22819681 | 22821305 | 1625 | 2,56 | 4177 |
| chrUextra | 22830296 | 22831909 | 1614 | 2,23 | 4178 |
| chrUextra | 22839379 | 22840746 | 1368 | 3,43 | 4179 |
| chrUextra | 22846802 | 22847897 | 1096 | 2,98 | 4180 |
| chrUextra | 22865961 | 22867437 | 1477 | 2,98 | 4181 |
| chrUextra | 22876564 | 22879646 | 3083 | 2,49 | 4182 |
| chrUextra | 22910613 | 22912176 | 1564 | 2,26 | 4183 |
| chrUextra | 22940770 | 22942472 | 1703 | 2,72 | 4184 |
| chrUextra | 22952137 | 22953761 | 1625 | 2,2 | 4185 |
| chrUextra | 22961855 | 22962916 | 1062 | 2,85 | 4186 |
| chrUextra | 22965016 | 22966938 | 1923 | 2,87 | 4187 |
| chrUextra | 23003568 | 23005212 | 1645 | 2,2 | 4188 |
| chrUextra | 23010330 | 23012000 | 1671 | 2,46 | 4189 |
| chrUextra | 23014107 | 23015391 | 1285 | 3,85 | 4190 |
| chrUextra | 23025399 | 23026643 | 1245 | 3,74 | 4191 |
| chrUextra | 23050480 | 23052103 | 1624 | 2,33 | 4192 |
| chrUextra | 23052661 | 23054383 | 1723 | 2,33 | 4193 |
| chrUextra | 23054828 | 23055951 | 1124 | 2,91 | 4194 |
| chrUextra | 23061550 | 23062749 | 1200 | 3,88 | 4195 |
| chrUextra | 23067144 | 23067956 | 813 | 5,44 | 4196 |
| chrUextra | 23079049 | 23081345 | 2297 | 2,49 | 4197 |
| chrUextra | 23121415 | 23122954 | 1540 | 2,33 | 4198 |
| chrUextra | 23133399 | 23135126 | 1728 | 3,19 | 4199 |
| chrUextra | 23137214 | 23138855 | 1642 | 2,26 | 4200 |
| chrUextra | 23168900 | 23171376 | 2477 | 2,85 | 4201 |
| chrUextra | 23175610 | 23176595 | 986 | 3,92 | 4202 |
| chrUextra | 23179542 | 23181035 | 1494 | 5,31 | 4203 |
| chrUextra | 23201871 | 23202553 | 683 | 4,79 | 4204 |
| chrUextra | 23205668 | 23206467 | 800 | 5,57 | 4205 |
| chrUextra | 23213370 | 23214362 | 993 | 2,59 | 4206 |
| chrUextra | 23227026 | 23228666 | 1641 | 2,39 | 4207 |
| chrUextra | 23232963 | 23234660 | 1698 | 2,56 | 4208 |
| chrUextra | 23269808 | 23270774 | 967 | 2,46 | 4209 |
| chrUextra | 23276564 | 23278285 | 1722 | 2,33 | 4210 |
| chrUextra | 23293788 | 23294628 | 841 | 5,66 | 4211 |
| chrUextra | 23299185 | 23300300 | 1116 | 3,01 | 4212 |
| chrUextra | 23303880 | 23306796 | 2917 | 2,52 | 4213 |
| chrUextra | 23311232 | 23312372 | 1141 | 3,64 | 4214 |
| chrUextra | 23317254 | 23319628 | 2375 | 2,88 | 4215 |
| chrUextra | 23324088 | 23325737 | 1650 | 2,33 | 4216 |
| chrUextra | 23336286 | 23338064 | 1779 | 5,21 | 4217 |
| chrUextra | 23352701 | 23353384 | 684 | 4,01 | 4218 |
| chrUextra | 23365351 | 23366478 | 1128 | 3,4 | 4219 |
| chrUextra | 23382009 | 23383657 | 1649 | 2,2 | 4220 |
| chrUextra | 23395578 | 23397176 | 1599 | 2,1 | 4221 |

| | | | | | |
|---|---|---|---|---|---|
| chrUextra | 23406735 | 23408216 | 1482 | 5,24 | 4222 |
| chrUextra | 23408501 | 23410014 | 1514 | 2,65 | 4223 |
| chrUextra | 23450492 | 23452063 | 1572 | 2,43 | 4224 |
| chrUextra | 23466824 | 23467892 | 1069 | 3,95 | 4225 |
| chrUextra | 23499143 | 23501566 | 2424 | 2,44 | 4226 |
| chrUextra | 23511103 | 23512277 | 1175 | 2,83 | 4227 |
| chrUextra | 23514932 | 23517237 | 2306 | 3,49 | 4228 |
| chrUextra | 23533744 | 23535450 | 1707 | 2,62 | 4229 |
| chrUextra | 23597606 | 23599275 | 1670 | 2,43 | 4230 |
| chrUextra | 23617004 | 23618625 | 1622 | 2,98 | 4231 |
| chrUextra | 23657658 | 23659105 | 1448 | 2,17 | 4232 |
| chrUextra | 23678657 | 23680609 | 1953 | 2,3 | 4233 |
| chrUextra | 23681534 | 23683952 | 2419 | 2,82 | 4234 |
| chrUextra | 23698022 | 23699687 | 1666 | 2,65 | 4235 |
| chrUextra | 23700231 | 23701901 | 1671 | 2,78 | 4236 |
| chrUextra | 23702462 | 23703595 | 1134 | 3,4 | 4237 |
| chrUextra | 23703691 | 23704957 | 1267 | 5,96 | 4238 |
| chrUextra | 23739873 | 23740917 | 1045 | 3,15 | 4239 |
| chrUextra | 23791567 | 23793073 | 1507 | 2,1 | 4240 |
| chrUextra | 23854291 | 23855992 | 1702 | 2,69 | 4241 |
| chrUextra | 23865376 | 23866707 | 1332 | 2,74 | 4242 |
| chrUextra | 23894572 | 23896256 | 1685 | 5,74 | 4243 |
| chrUextra | 23940962 | 23942029 | 1068 | 3,66 | 4244 |
| chrUextra | 23953702 | 23955326 | 1625 | 2,65 | 4245 |
| chrUextra | 23959622 | 23960594 | 973 | 3,75 | 4246 |
| chrUextra | 24002713 | 24004581 | 1869 | 6,05 | 4247 |
| chrUextra | 24017886 | 24018858 | 973 | 2,59 | 4248 |
| chrUextra | 24026046 | 24027388 | 1343 | 2,95 | 4249 |
| chrUextra | 24034999 | 24036531 | 1533 | 2,26 | 4250 |
| chrUextra | 24041704 | 24045634 | 3931 | 3,56 | 4251 |
| chrUextra | 24055095 | 24056317 | 1223 | 4,23 | 4252 |
| chrUextra | 24101190 | 24101927 | 738 | 4,85 | 4253 |
| chrUextra | 24141643 | 24143122 | 1480 | 2,49 | 4254 |
| chrUextra | 24147459 | 24149127 | 1669 | 2,39 | 4255 |
| chrUextra | 24175831 | 24177384 | 1554 | 2,59 | 4256 |
| chrUextra | 24208518 | 24210914 | 2397 | 2,65 | 4257 |
| chrUextra | 24227006 | 24228720 | 1715 | 2,36 | 4258 |
| chrUextra | 24228802 | 24229786 | 985 | 4,34 | 4259 |
| chrUextra | 24233026 | 24234848 | 1823 | 2,83 | 4260 |
| chrUextra | 24250157 | 24251638 | 1482 | 2,33 | 4261 |
| chrUextra | 24263432 | 24264392 | 961 | 2,65 | 4262 |
| chrUextra | 24267806 | 24269874 | 2069 | 2,96 | 4263 |
| chrUextra | 24271622 | 24272574 | 953 | 3,51 | 4264 |
| chrUextra | 24272891 | 24274089 | 1199 | 3,07 | 4265 |
| chrUextra | 24277413 | 24278039 | 627 | 4,04 | 4266 |
| chrUextra | 24281953 | 24283406 | 1454 | 2,74 | 4267 |
| chrUextra | 24287103 | 24288312 | 1210 | 4,51 | 4268 |
| chrUextra | 24298312 | 24299288 | 977 | 2,43 | 4269 |

| | | | | | |
|---|---|---|---|---|---|
| chrUextra | 24312573 | 24314816 | 2244 | 2,55 | 4270 |
| chrUextra | 24345858 | 24346869 | 1012 | 3,14 | 4271 |
| chrUextra | 24351035 | 24352685 | 1651 | 2,82 | 4272 |
| chrUextra | 24365246 | 24366408 | 1163 | 6,7 | 4273 |
| chrUextra | 24368494 | 24369789 | 1296 | 3,7 | 4274 |
| chrUextra | 24379694 | 24380905 | 1212 | 3,66 | 4275 |
| chrUextra | 24388569 | 24389646 | 1078 | 4,69 | 4276 |
| chrUextra | 24392573 | 24394204 | 1632 | 2,39 | 4277 |
| chrUextra | 24411759 | 24412835 | 1077 | 2,94 | 4278 |
| chrUextra | 24413287 | 24414853 | 1567 | 2,46 | 4279 |
| chrUextra | 24425810 | 24427437 | 1628 | 2,46 | 4280 |
| chrUextra | 24447334 | 24448648 | 1315 | 3,34 | 4281 |
| chrUextra | 24482946 | 24483834 | 889 | 3,65 | 4282 |
| chrUextra | 24513224 | 24514201 | 978 | 3,04 | 4283 |
| chrUextra | 24518385 | 24520835 | 2451 | 2,46 | 4284 |
| chrUextra | 24522807 | 24523849 | 1043 | 2,49 | 4285 |
| chrUextra | 24529543 | 24531164 | 1622 | 2,52 | 4286 |
| chrUextra | 24540628 | 24542309 | 1682 | 2,82 | 4287 |
| chrUextra | 24547279 | 24548898 | 1620 | 2,59 | 4288 |
| chrUextra | 24559526 | 24560620 | 1095 | 4,04 | 4289 |
| chrUextra | 24586358 | 24587692 | 1335 | 2,46 | 4290 |
| chrUextra | 24601324 | 24603650 | 2327 | 2,65 | 4291 |
| chrUextra | 24640243 | 24641332 | 1090 | 2,59 | 4292 |
| chrUextra | 24643351 | 24647201 | 3851 | 3,08 | 4293 |
| chrUextra | 24648500 | 24649406 | 907 | 3,56 | 4294 |
| chrUextra | 24663861 | 24664935 | 1075 | 2,98 | 4295 |
| chrUextra | 24675787 | 24678180 | 2394 | 2,85 | 4296 |
| chrUextra | 24687603 | 24689319 | 1717 | 2,52 | 4297 |
| chrUextra | 24694241 | 24694678 | 438 | 3,07 | 4298 |
| chrUextra | 24711209 | 24712815 | 1607 | 2,36 | 4299 |
| chrUextra | 24727409 | 24728419 | 1011 | 2,97 | 4300 |
| chrUextra | 24750998 | 24752305 | 1308 | 5,02 | 4301 |
| chrUextra | 24753227 | 24754882 | 1656 | 2,43 | 4302 |
| chrUextra | 24759038 | 24760010 | 973 | 2,65 | 4303 |
| chrUextra | 24792078 | 24793014 | 937 | 3,22 | 4304 |
| chrUextra | 24806138 | 24807273 | 1136 | 4,4 | 4305 |
| chrUextra | 24835956 | 24836706 | 751 | 5,33 | 4306 |
| chrUextra | 24848894 | 24849824 | 931 | 2,62 | 4307 |
| chrUextra | 24851846 | 24853234 | 1389 | 2,49 | 4308 |
| chrUextra | 24876346 | 24877144 | 799 | 3,14 | 4309 |
| chrUextra | 24900043 | 24900814 | 772 | 4,22 | 4310 |
| chrUextra | 24904212 | 24905743 | 1532 | 2,91 | 4311 |
| chrUextra | 24909700 | 24910364 | 665 | 3,92 | 4312 |
| chrUextra | 24933326 | 24934498 | 1173 | 4,01 | 4313 |
| chrUextra | 24984703 | 24985899 | 1197 | 6,33 | 4314 |
| chrUextra | 24988442 | 24990024 | 1583 | 2,33 | 4315 |
| chrUextra | 25000314 | 25001473 | 1160 | 6,06 | 4316 |
| chrUextra | 25027246 | 25028852 | 1607 | 2,72 | 4317 |

| | | | | | |
|---|---|---|---|---|---|
| chrUextra | 25085243 | 25086745 | 1503 | 2,04 | 4318 |
| chrUextra | 25091822 | 25093392 | 1571 | 2,69 | 4319 |
| chrUextra | 25093756 | 25094685 | 930 | 5,55 | 4320 |
| chrUextra | 25127607 | 25129495 | 1889 | 4,15 | 4321 |
| chrUextra | 25144520 | 25146024 | 1505 | 2,36 | 4322 |
| chrUextra | 25150864 | 25151696 | 833 | 3,84 | 4323 |
| chrUextra | 25175359 | 25176891 | 1533 | 2,72 | 4324 |
| chrUextra | 25184256 | 25184957 | 702 | 4,3 | 4325 |
| chrUextra | 25190374 | 25192155 | 1782 | 3 | 4326 |
| chrUextra | 25194375 | 25195283 | 909 | 4,3 | 4327 |
| chrUextra | 25217486 | 25218458 | 973 | 3,49 | 4328 |
| chrUextra | 25235856 | 25237438 | 1583 | 2,56 | 4329 |
| chrUextra | 25257090 | 25258657 | 1568 | 3,78 | 4330 |
| chrUextra | 25262947 | 25264414 | 1468 | 2,3 | 4331 |
| chrUextra | 25285391 | 25286325 | 935 | 2,66 | 4332 |
| chrUextra | 25314688 | 25316243 | 1556 | 6,37 | 4333 |
| chrUextra | 25316751 | 25318286 | 1536 | 2,26 | 4334 |
| chrUextra | 25379867 | 25381558 | 1692 | 2,2 | 4335 |
| chrUextra | 25407501 | 25408635 | 1135 | 5,04 | 4336 |
| chrUextra | 25440423 | 25442070 | 1648 | 2,36 | 4337 |
| chrUextra | 25447683 | 25450052 | 2370 | 3,53 | 4338 |
| chrUextra | 25478329 | 25479439 | 1111 | 5,97 | 4339 |
| chrUextra | 25504393 | 25506270 | 1878 | 4,76 | 4340 |
| chrUextra | 25508567 | 25509628 | 1062 | 3,53 | 4341 |
| chrUextra | 25522237 | 25523156 | 920 | 5,53 | 4342 |
| chrUextra | 25626745 | 25627847 | 1103 | 3,82 | 4343 |
| chrUextra | 25642698 | 25644687 | 1990 | 3,33 | 4344 |
| chrUextra | 25656053 | 25656854 | 802 | 4,42 | 4345 |
| chrUextra | 25667260 | 25668681 | 1422 | 3,79 | 4346 |
| chrUextra | 25685332 | 25686316 | 985 | 3,56 | 4347 |
| chrUextra | 25688400 | 25689876 | 1477 | 2,33 | 4348 |
| chrUextra | 25691404 | 25693371 | 1968 | 2,17 | 4349 |
| chrUextra | 25715685 | 25718043 | 2359 | 2,23 | 4350 |
| chrUextra | 25758956 | 25759865 | 910 | 2,85 | 4351 |
| chrUextra | 25767103 | 25769297 | 2195 | 2,72 | 4352 |
| chrUextra | 25787124 | 25788013 | 890 | 2,36 | 4353 |
| chrUextra | 25800832 | 25802467 | 1636 | 2,26 | 4354 |
| chrUextra | 25836131 | 25837765 | 1635 | 2,39 | 4355 |
| chrUextra | 25842007 | 25843426 | 1420 | 2,14 | 4356 |
| chrUextra | 25871527 | 25872524 | 998 | 6,08 | 4357 |
| chrUextra | 25879374 | 25880969 | 1596 | 2,88 | 4358 |
| chrUextra | 25891710 | 25893196 | 1487 | 2,82 | 4359 |
| chrUextra | 25896656 | 25898239 | 1584 | 2,39 | 4360 |
| chrUextra | 25904602 | 25907623 | 3022 | 2,65 | 4361 |
| chrUextra | 25908839 | 25909358 | 520 | 4,72 | 4362 |
| chrUextra | 25911023 | 25911908 | 886 | 2,39 | 4363 |
| chrUextra | 25928201 | 25929323 | 1123 | 3,04 | 4364 |
| chrUextra | 25944158 | 25946408 | 2251 | 2,52 | 4365 |

| | | | | | |
|---|---|---|---|---|---|
| chrUextra | 25946900 | 25948518 | 1619 | 2,56 | 4366 |
| chrUextra | 25949519 | 25950663 | 1145 | 2,36 | 4367 |
| chrUextra | 25964898 | 25966504 | 1607 | 2,43 | 4368 |
| chrUextra | 25973564 | 25975860 | 2297 | 2,43 | 4369 |
| chrUextra | 25992870 | 25993808 | 939 | 2,52 | 4370 |
| chrUextra | 26034332 | 26035432 | 1101 | 6,23 | 4371 |
| chrUextra | 26040065 | 26041093 | 1029 | 5,14 | 4372 |
| chrUextra | 26041685 | 26042654 | 970 | 2,65 | 4373 |
| chrUextra | 26068934 | 26070485 | 1552 | 2,98 | 4374 |
| chrUextra | 26093263 | 26094734 | 1472 | 2,36 | 4375 |
| chrUextra | 26106100 | 26106972 | 873 | 2,65 | 4376 |
| chrUextra | 26115384 | 26117009 | 1626 | 3,2 | 4377 |
| chrUextra | 26122560 | 26124582 | 2023 | 2,23 | 4378 |
| chrUextra | 26131923 | 26133495 | 1573 | 2,56 | 4379 |
| chrUextra | 26138258 | 26138672 | 415 | 3,69 | 4380 |
| chrUextra | 26159498 | 26160733 | 1236 | 5,59 | 4381 |
| chrUextra | 26185300 | 26186246 | 947 | 4,87 | 4382 |
| chrUextra | 26195921 | 26196856 | 936 | 4,92 | 4383 |
| chrUextra | 26206172 | 26208246 | 2075 | 5,48 | 4384 |
| chrUextra | 26225339 | 26226827 | 1489 | 2,3 | 4385 |
| chrUextra | 26243004 | 26244343 | 1340 | 3,87 | 4386 |
| chrUextra | 26280831 | 26282477 | 1647 | 3,26 | 4387 |
| chrUextra | 26282969 | 26284118 | 1150 | 3,97 | 4388 |
| chrUextra | 26303839 | 26305381 | 1543 | 2,52 | 4389 |
| chrUextra | 26310779 | 26311928 | 1150 | 3,97 | 4390 |
| chrUextra | 26317279 | 26319351 | 2073 | 2,23 | 4391 |
| chrUextra | 26328429 | 26329322 | 894 | 4,89 | 4392 |
| chrUextra | 26341468 | 26342595 | 1128 | 4,92 | 4393 |
| chrUextra | 26346068 | 26346953 | 886 | 5,31 | 4394 |
| chrUextra | 26383993 | 26386187 | 2195 | 5,94 | 4395 |
| chrUextra | 26395465 | 26396549 | 1085 | 4,53 | 4396 |
| chrUextra | 26401272 | 26403353 | 2082 | 2,39 | 4397 |
| chrUextra | 26418047 | 26419751 | 1705 | 4,24 | 4398 |
| chrUextra | 26470463 | 26472786 | 2324 | 2,75 | 4399 |
| chrUextra | 26495937 | 26497855 | 1919 | 6,01 | 4400 |
| chrUextra | 26505192 | 26506266 | 1075 | 3,7 | 4401 |
| chrUextra | 26512257 | 26513806 | 1550 | 2,59 | 4402 |
| chrUextra | 26525341 | 26526305 | 965 | 5,14 | 4403 |
| chrUextra | 26527767 | 26529258 | 1492 | 2,36 | 4404 |
| chrUextra | 26553895 | 26554800 | 906 | 2,72 | 4405 |
| chrUextra | 26558915 | 26561756 | 2842 | 5,32 | 4406 |
| chrUextra | 26564263 | 26565313 | 1051 | 4,53 | 4407 |
| chrUextra | 26565912 | 26566992 | 1081 | 4,57 | 4408 |
| chrUextra | 26567850 | 26568621 | 772 | 6,68 | 4409 |
| chrUextra | 26577201 | 26578767 | 1567 | 2,26 | 4410 |
| chrUextra | 26581917 | 26582833 | 917 | 5,14 | 4411 |
| chrUextra | 26642022 | 26642876 | 855 | 2,88 | 4412 |
| chrUextra | 26648334 | 26650629 | 2296 | 3,01 | 4413 |

| | | | | | |
|---|---|---|---|---|---|
| chrUextra | 26664614 | 26665683 | 1070 | 3,07 | 4414 |
| chrUextra | 26706712 | 26708769 | 2058 | 2,93 | 4415 |
| chrUextra | 26710172 | 26710970 | 799 | 5,02 | 4416 |
| chrUextra | 26716542 | 26717336 | 795 | 2,82 | 4417 |
| chrUextra | 26728459 | 26730030 | 1572 | 2,62 | 4418 |
| chrUextra | 26743645 | 26744767 | 1123 | 3,53 | 4419 |
| chrUextra | 26750244 | 26751781 | 1538 | 2,39 | 4420 |
| chrUextra | 26756301 | 26757088 | 788 | 5,27 | 4421 |
| chrUextra | 26790382 | 26791234 | 853 | 3,34 | 4422 |
| chrUextra | 26802033 | 26803567 | 1535 | 2,46 | 4423 |
| chrUextra | 26844149 | 26844995 | 847 | 5,82 | 4424 |
| chrUextra | 26849484 | 26851056 | 1573 | 2,39 | 4425 |
| chrUextra | 26851475 | 26852734 | 1260 | 4,86 | 4426 |
| chrUextra | 26906200 | 26907061 | 862 | 4,39 | 4427 |
| chrUextra | 26936543 | 26937978 | 1436 | 2,2 | 4428 |
| chrUextra | 26941151 | 26942131 | 981 | 4,96 | 4429 |
| chrUextra | 26947748 | 26949056 | 1309 | 2,2 | 4430 |
| chrUextra | 26953315 | 26955477 | 2163 | 2,01 | 4431 |
| chrUextra | 26969931 | 26972129 | 2199 | 5,82 | 4432 |
| chrUextra | 26976725 | 26977639 | 915 | 2,72 | 4433 |
| chrUextra | 26995531 | 26996537 | 1007 | 6,56 | 4434 |
| chrUextra | 27019756 | 27020660 | 905 | 2,83 | 4435 |
| chrUextra | 27056524 | 27057838 | 1315 | 6,37 | 4436 |
| chrUextra | 27069313 | 27070286 | 974 | 5,27 | 4437 |
| chrUextra | 27070736 | 27072016 | 1281 | 4,65 | 4438 |
| chrUextra | 27092313 | 27093066 | 754 | 6,08 | 4439 |
| chrUextra | 27099170 | 27100213 | 1044 | 3,04 | 4440 |
| chrUextra | 27110195 | 27111771 | 1577 | 2,69 | 4441 |
| chrUextra | 27145311 | 27146251 | 941 | 2,72 | 4442 |
| chrUextra | 27214044 | 27214588 | 545 | 4,76 | 4443 |
| chrUextra | 27225068 | 27225962 | 895 | 4,25 | 4444 |
| chrUextra | 27238536 | 27239592 | 1057 | 5,69 | 4445 |
| chrUextra | 27292749 | 27294204 | 1456 | 2,52 | 4446 |
| chrUextra | 27319321 | 27319911 | 591 | 3,98 | 4447 |
| chrUextra | 27346321 | 27347785 | 1465 | 11,42 | 4448 |
| chrUextra | 27366505 | 27367015 | 511 | 3,66 | 4449 |
| chrUextra | 27367838 | 27369188 | 1351 | 5,79 | 4450 |
| chrUextra | 27370176 | 27372151 | 1976 | 4,47 | 4451 |
| chrUextra | 27390448 | 27391582 | 1135 | 3,56 | 4452 |
| chrUextra | 27431645 | 27432396 | 752 | 3,29 | 4453 |
| chrUextra | 27449820 | 27451228 | 1409 | 2,39 | 4454 |
| chrUextra | 27476758 | 27477907 | 1150 | 2,52 | 4455 |
| chrUextra | 27480664 | 27481304 | 641 | 4,56 | 4456 |
| chrUextra | 27481447 | 27482541 | 1095 | 3,11 | 4457 |
| chrUextra | 27484885 | 27485926 | 1042 | 6,31 | 4458 |
| chrUextra | 27511131 | 27513281 | 2151 | 3,57 | 4459 |
| chrUextra | 27519036 | 27519867 | 832 | 5,63 | 4460 |
| chrUextra | 27526169 | 27526737 | 569 | 3,46 | 4461 |

| | | | | | |
|---|---|---|---|---|---|
| chrUextra | 27534578 | 27536267 | 1690 | 2,66 | 4462 |
| chrUextra | 27550026 | 27550898 | 873 | 3,23 | 4463 |
| chrUextra | 27556747 | 27557625 | 879 | 3,14 | 4464 |
| chrUextra | 27590688 | 27591782 | 1095 | 6,94 | 4465 |
| chrUextra | 27599403 | 27601308 | 1906 | 5,29 | 4466 |
| chrUextra | 27663297 | 27664448 | 1152 | 4,89 | 4467 |
| chrUextra | 27666872 | 27668414 | 1543 | 2,3 | 4468 |
| chrUextra | 27669983 | 27671175 | 1193 | 3,93 | 4469 |
| chrUextra | 27685320 | 27686688 | 1369 | 5,76 | 4470 |
| chrUextra | 27700569 | 27701601 | 1033 | 5,56 | 4471 |
| chrUextra | 27709322 | 27710187 | 866 | 4,43 | 4472 |
| chrUextra | 27711785 | 27712947 | 1163 | 4,19 | 4473 |
| chrUextra | 27744232 | 27745785 | 1554 | 3,24 | 4474 |
| chrUextra | 27790147 | 27791377 | 1231 | 3,86 | 4475 |
| chrUextra | 27809176 | 27810251 | 1076 | 5,7 | 4476 |
| chrUextra | 27819578 | 27820968 | 1391 | 2,43 | 4477 |
| chrUextra | 27825366 | 27826661 | 1296 | 5,12 | 4478 |
| chrUextra | 27828106 | 27829483 | 1378 | 2,49 | 4479 |
| chrUextra | 27837498 | 27838309 | 812 | 4,09 | 4480 |
| chrUextra | 27850430 | 27851208 | 779 | 5,95 | 4481 |
| chrUextra | 27862580 | 27863592 | 1013 | 3,72 | 4482 |
| chrUextra | 27871930 | 27873161 | 1232 | 4,32 | 4483 |
| chrUextra | 27890346 | 27891665 | 1320 | 5,31 | 4484 |
| chrUextra | 27894871 | 27895724 | 854 | 2,62 | 4485 |
| chrUextra | 27903862 | 27904947 | 1086 | 4,72 | 4486 |
| chrUextra | 27905099 | 27906744 | 1646 | 4,07 | 4487 |
| chrUextra | 27908891 | 27909927 | 1037 | 5,4 | 4488 |
| chrUextra | 27922611 | 27923964 | 1354 | 4,05 | 4489 |
| chrUextra | 27932170 | 27933740 | 1571 | 4,5 | 4490 |
| chrUextra | 27973569 | 27974762 | 1194 | 3,88 | 4491 |
| chrUextra | 27976224 | 27977087 | 864 | 5,47 | 4492 |
| chrUextra | 27979609 | 27980902 | 1294 | 5,41 | 4493 |
| chrUextra | 27982634 | 27983703 | 1070 | 4,68 | 4494 |
| chrUextra | 27985180 | 27986298 | 1119 | 5,94 | 4495 |
| chrUextra | 27997383 | 27998206 | 824 | 4,04 | 4496 |
| chrUextra | 27998838 | 27999642 | 805 | 3,07 | 4497 |
| chrUextra | 28014418 | 28015540 | 1123 | 6,63 | 4498 |
| chrUextra | 28029209 | 28029987 | 779 | 3,2 | 4499 |
| chrUextra | 28031409 | 28032256 | 848 | 4,05 | 4500 |
| chrUextra | 28073185 | 28074263 | 1079 | 5,66 | 4501 |
| chrUextra | 28090112 | 28091136 | 1025 | 5,02 | 4502 |
| chrUextra | 28091655 | 28092140 | 486 | 3,4 | 4503 |
| chrUextra | 28093322 | 28094997 | 1676 | 6,37 | 4504 |
| chrUextra | 28106589 | 28108135 | 1547 | 5,05 | 4505 |
| chrUextra | 28130374 | 28130850 | 477 | 3,14 | 4506 |
| chrUextra | 28136793 | 28137409 | 617 | 3,2 | 4507 |
| chrUextra | 28137524 | 28138601 | 1078 | 4,68 | 4508 |
| chrUextra | 28142386 | 28143220 | 835 | 5,08 | 4509 |

| | | | | | |
|---|---|---|---|---|---|
| chrUextra | 28164219 | 28165438 | 1220 | 5,64 | 4510 |
| chrUextra | 28172030 | 28172847 | 818 | 3,89 | 4511 |
| chrUextra | 28193711 | 28194546 | 836 | 5,24 | 4512 |
| chrUextra | 28213334 | 28214557 | 1224 | 3,77 | 4513 |
| chrUextra | 28219429 | 28220669 | 1241 | 3,51 | 4514 |
| chrUextra | 28222544 | 28223639 | 1096 | 6,15 | 4515 |
| chrUextra | 28245598 | 28246399 | 802 | 3,14 | 4516 |
| chrUextra | 28247065 | 28247565 | 501 | 4,56 | 4517 |
| chrUextra | 28248885 | 28249774 | 890 | 4,64 | 4518 |
| chrUextra | 28250397 | 28251132 | 736 | 5,05 | 4519 |
| chrUextra | 28253017 | 28254860 | 1844 | 5,02 | 4520 |
| chrUextra | 28270062 | 28270844 | 783 | 3,53 | 4521 |
| chrUextra | 28271271 | 28271975 | 705 | 4,39 | 4522 |
| chrUextra | 28280190 | 28281114 | 925 | 4,14 | 4523 |
| chrUextra | 28281437 | 28282105 | 669 | 5,93 | 4524 |
| chrUextra | 28290552 | 28291324 | 773 | 3,01 | 4525 |
| chrUextra | 28299674 | 28300280 | 607 | 2,86 | 4526 |
| chrUextra | 28308477 | 28309645 | 1169 | 4,78 | 4527 |
| chrUextra | 28323799 | 28325002 | 1204 | 5,68 | 4528 |
| chrUextra | 28325060 | 28326218 | 1159 | 6,04 | 4529 |
| chrUextra | 28334049 | 28334623 | 575 | 4,81 | 4530 |
| chrUextra | 28339283 | 28340170 | 888 | 4,83 | 4531 |
| chrUextra | 28354679 | 28355499 | 821 | 4,06 | 4532 |
| chrUextra | 28359943 | 28360784 | 842 | 5,37 | 4533 |
| chrUextra | 28367034 | 28367995 | 962 | 6,79 | 4534 |
| chrUextra | 28368205 | 28368923 | 719 | 4,04 | 4535 |
| chrUextra | 28388524 | 28389588 | 1065 | 6,44 | 4536 |
| chrUextra | 28391359 | 28392506 | 1148 | 4,89 | 4537 |
| chrUextra | 28393861 | 28394498 | 638 | 4,4 | 4538 |
| chrUextra | 28407584 | 28408628 | 1045 | 5,5 | 4539 |
| chrUextra | 28422465 | 28423752 | 1288 | 7,47 | 4540 |
| chrUextra | 28427302 | 28427976 | 675 | 2,62 | 4541 |
| chrUextra | 28432849 | 28434003 | 1155 | 4,51 | 4542 |
| chrUextra | 28436223 | 28437403 | 1181 | 5,92 | 4543 |
| chrUextra | 28440793 | 28441726 | 934 | 6,76 | 4544 |
| chrUextra | 28446150 | 28446888 | 739 | 2,85 | 4545 |
| chrUextra | 28447184 | 28448267 | 1084 | 6,08 | 4546 |
| chrUextra | 28448808 | 28450195 | 1388 | 4 | 4547 |
| chrUextra | 28455611 | 28456290 | 680 | 3,72 | 4548 |
| chrUextra | 28456391 | 28457427 | 1037 | 5,99 | 4549 |
| chrUextra | 28459785 | 28460259 | 475 | 2,77 | 4550 |
| chrUextra | 28461314 | 28462309 | 996 | 6,21 | 4551 |
| chrUextra | 28462417 | 28464023 | 1607 | 5,47 | 4552 |
| chrUextra | 28477051 | 28478108 | 1058 | 4,39 | 4553 |
| chrUextra | 28480101 | 28481114 | 1014 | 5,69 | 4554 |
| chrUextra | 28487084 | 28488724 | 1641 | 4,58 | 4555 |
| chrUextra | 28495198 | 28495946 | 749 | 3,01 | 4556 |
| chrUextra | 28501671 | 28503041 | 1371 | 4,02 | 4557 |

| | | | | | |
|---|---|---|---|---|---|
| chrUextra | 28508273 | 28509548 | 1276 | 6,6 | 4558 |
| chrUextra | 28518451 | 28519497 | 1047 | 6 | 4559 |
| chrUextra | 28522719 | 28523929 | 1211 | 4,89 | 4560 |
| chrUextra | 28524039 | 28525093 | 1055 | 5,64 | 4561 |
| chrUextra | 28530800 | 28531673 | 874 | 5,47 | 4562 |
| chrUextra | 28535683 | 28536427 | 745 | 3,12 | 4563 |
| chrUextra | 28539324 | 28540210 | 887 | 5,28 | 4564 |
| chrUextra | 28541773 | 28542821 | 1049 | 6,13 | 4565 |
| chrUextra | 28543724 | 28544630 | 907 | 4,17 | 4566 |
| chrUextra | 28549170 | 28550161 | 992 | 4,85 | 4567 |
| chrUextra | 28551368 | 28552496 | 1129 | 4,03 | 4568 |
| chrUextra | 28555184 | 28557860 | 2677 | 3,78 | 4569 |
| chrUextra | 28561018 | 28562533 | 1516 | 4,45 | 4570 |
| chrUextra | 28567567 | 28569215 | 1649 | 6,36 | 4571 |
| chrUextra | 28570447 | 28571300 | 854 | 5,47 | 4572 |
| chrUextra | 28572057 | 28573072 | 1016 | 5,87 | 4573 |
| chrUextra | 28573661 | 28574605 | 945 | 6,22 | 4574 |
| chrUextra | 28577940 | 28578424 | 485 | 4,43 | 4575 |
| chrUextra | 28579536 | 28580272 | 737 | 5,92 | 4576 |
| chrUextra | 28580602 | 28581329 | 728 | 4,04 | 4577 |
| chrUextra | 28581668 | 28583935 | 2268 | 4,03 | 4578 |
| chrUextra | 28584853 | 28585567 | 715 | 5,55 | 4579 |
| chrUextra | 28586701 | 28587721 | 1021 | 5,69 | 4580 |
| chrUextra | 28587815 | 28589056 | 1242 | 4,69 | 4581 |
| chrUextra | 28592381 | 28593422 | 1042 | 3,13 | 4582 |
| chrUextra | 28594191 | 28595119 | 929 | 4,14 | 4583 |
| chrUextra | 28595232 | 28596365 | 1134 | 3,67 | 4584 |
| chrUextra | 28596845 | 28598087 | 1243 | 5,23 | 4585 |
| chrUextra | 28601772 | 28602643 | 872 | 5,02 | 4586 |
| chrUextra | 28604688 | 28606243 | 1556 | 5,68 | 4587 |
| chrUextra | 28608464 | 28609505 | 1042 | 6,97 | 4588 |
| chrUextra | 28609647 | 28610421 | 775 | 6,37 | 4589 |
| chrUextra | 28611143 | 28612099 | 957 | 3,92 | 4590 |
| chrUextra | 28612792 | 28613492 | 701 | 4,76 | 4591 |
| chrUextra | 28615127 | 28615871 | 745 | 5,45 | 4592 |
| chrUextra | 28617510 | 28618529 | 1020 | 5,43 | 4593 |
| chrUextra | 28618537 | 28619259 | 723 | 4,75 | 4594 |
| chrUextra | 28620626 | 28622066 | 1441 | 6,15 | 4595 |
| chrUextra | 28623745 | 28624451 | 707 | 5,27 | 4596 |
| chrUextra | 28624599 | 28625509 | 911 | 6,08 | 4597 |
| chrUextra | 28625830 | 28626505 | 676 | 4,78 | 4598 |
| chrUextra | 28630310 | 28631165 | 856 | 4,44 | 4599 |
| chrUextra | 28631898 | 28632697 | 800 | 2,82 | 4600 |
| chrUextra | 28637659 | 28639574 | 1916 | 4,3 | 4601 |
| chrUextra | 28641099 | 28642564 | 1466 | 4,66 | 4602 |
| chrUextra | 28646437 | 28647656 | 1220 | 5,7 | 4603 |
| chrUextra | 28648963 | 28650019 | 1057 | 5,78 | 4604 |
| chrUextra | 28652078 | 28655854 | 3777 | 5,87 | 4605 |

| | | | | | |
|---|---|---|---|---|---|
| chrUextra | 28656466 | 28657834 | 1369 | 4,27 | 4606 |
| chrUextra | 28658507 | 28659628 | 1122 | 6,49 | 4607 |
| chrUextra | 28668244 | 28669346 | 1103 | 6,5 | 4608 |
| chrUextra | 28669753 | 28670264 | 512 | 3,37 | 4609 |
| chrUextra | 28671261 | 28671819 | 559 | 4,59 | 4610 |
| chrUextra | 28675598 | 28676446 | 849 | 6,04 | 4611 |
| chrUextra | 28676775 | 28677658 | 884 | 3,89 | 4612 |
| chrUextra | 28681753 | 28682538 | 786 | 3,37 | 4613 |
| chrUextra | 28686158 | 28687167 | 1010 | 5,86 | 4614 |
| chrUextra | 28687705 | 28688906 | 1202 | 3,07 | 4615 |
| chrUextra | 28695058 | 28696064 | 1007 | 4,4 | 4616 |
| chrUextra | 28701264 | 28702431 | 1168 | 4,34 | 4617 |
| chrUextra | 28702805 | 28703969 | 1165 | 4 | 4618 |
| chrUextra | 28705184 | 28706326 | 1143 | 6,41 | 4619 |
| chrUextra | 28708254 | 28709077 | 824 | 5,88 | 4620 |
| chrUextra | 28711320 | 28712264 | 945 | 3,2 | 4621 |
| chrUextra | 28719926 | 28722250 | 2325 | 3,45 | 4622 |
| chrUextra | 28722594 | 28723801 | 1208 | 4,2 | 4623 |
| chrUextra | 28724366 | 28725283 | 918 | 4,08 | 4624 |
| chrUextra | 28726554 | 28727670 | 1117 | 4,36 | 4625 |
| chrUextra | 28730658 | 28731676 | 1019 | 6,62 | 4626 |
| chrUextra | 28735834 | 28737711 | 1878 | 5,6 | 4627 |
| chrUextra | 28738247 | 28739058 | 812 | 4,19 | 4628 |
| chrUextra | 28739794 | 28742465 | 2672 | 6,48 | 4629 |
| chrUextra | 28743100 | 28743754 | 655 | 3,4 | 4630 |
| chrUextra | 28747755 | 28749017 | 1263 | 5,95 | 4631 |
| chrUextra | 28749188 | 28749832 | 645 | 4,14 | 4632 |
| chrUextra | 28751894 | 28754518 | 2625 | 4,42 | 4633 |
| chrUextra | 28754627 | 28756131 | 1505 | 5,73 | 4634 |
| chrUextra | 28756446 | 28758184 | 1739 | 5,99 | 4635 |
| chrUextra | 28759401 | 28760335 | 935 | 6,36 | 4636 |
| chrUextra | 28760798 | 28761800 | 1003 | 4,51 | 4637 |
| chrUextra | 28763713 | 28764911 | 1199 | 5,66 | 4638 |
| chrUextra | 28766870 | 28767713 | 844 | 6,18 | 4639 |
| chrUextra | 28768218 | 28769526 | 1309 | 5,45 | 4640 |
| chrUextra | 28769557 | 28771823 | 2267 | 5,09 | 4641 |
| chrUextra | 28775518 | 28776655 | 1138 | 5,17 | 4642 |
| chrUextra | 28780600 | 28781607 | 1008 | 5,89 | 4643 |
| chrUextra | 28786725 | 28787698 | 974 | 4,98 | 4644 |
| chrUextra | 28791904 | 28792814 | 911 | 6,19 | 4645 |
| chrUextra | 28795464 | 28796112 | 649 | 4,81 | 4646 |
| chrUextra | 28797038 | 28798626 | 1589 | 6,47 | 4647 |
| chrUextra | 28800392 | 28801205 | 814 | 6,42 | 4648 |
| chrUextra | 28803502 | 28805062 | 1561 | 4,01 | 4649 |
| chrUextra | 28805726 | 28806588 | 863 | 5,14 | 4650 |
| chrUextra | 28807771 | 28808612 | 842 | 6,13 | 4651 |
| chrUextra | 28809635 | 28810652 | 1018 | 5,37 | 4652 |
| chrUextra | 28810959 | 28812464 | 1506 | 3,79 | 4653 |

| | | | | | |
|---|---|---|---|---|---|
| chrUextra | 28815184 | 28818771 | 3588 | 4 | 4654 |
| chrUextra | 28821261 | 28822954 | 1694 | 4,95 | 4655 |
| chrUextra | 28823377 | 28824224 | 848 | 3,84 | 4656 |
| chrUextra | 28824311 | 28825143 | 833 | 5,34 | 4657 |
| chrUextra | 28826542 | 28827438 | 897 | 5,19 | 4658 |
| chrUextra | 28828488 | 28829848 | 1361 | 4,13 | 4659 |
| chrUextra | 28830739 | 28832679 | 1941 | 6,78 | 4660 |
| chrUextra | 28832900 | 28833950 | 1051 | 4,11 | 4661 |
| chrUextra | 28835632 | 28836553 | 922 | 4,21 | 4662 |
| chrUextra | 28838053 | 28839350 | 1298 | 5,27 | 4663 |
| chrUextra | 28845721 | 28846587 | 867 | 4,08 | 4664 |
| chrUextra | 28849369 | 28850025 | 657 | 4,72 | 4665 |
| chrUextra | 28850379 | 28851269 | 891 | 4,24 | 4666 |
| chrUextra | 28853119 | 28853940 | 822 | 6,18 | 4667 |
| chrUextra | 28854169 | 28855186 | 1018 | 5,66 | 4668 |
| chrUextra | 28856536 | 28857567 | 1032 | 5,05 | 4669 |
| chrUextra | 28858796 | 28859413 | 618 | 4,13 | 4670 |
| chrUextra | 28860123 | 28861583 | 1461 | 5,24 | 4671 |
| chrUextra | 28865467 | 28866266 | 800 | 4,79 | 4672 |
| chrUextra | 28868017 | 28869649 | 1633 | 4,25 | 4673 |
| chrUextra | 28874469 | 28876251 | 1783 | 3,95 | 4674 |
| chrUextra | 28877038 | 28877949 | 912 | 6,37 | 4675 |
| chrUextra | 28878259 | 28879238 | 980 | 5,06 | 4676 |
| chrUextra | 28879591 | 28880277 | 687 | 4,53 | 4677 |
| chrUextra | 28881534 | 28882148 | 615 | 5,6 | 4678 |
| chrUextra | 28883795 | 28885155 | 1361 | 4,68 | 4679 |
| chrUextra | 28885615 | 28886151 | 537 | 3,56 | 4680 |
| chrUextra | 28886371 | 28888203 | 1833 | 4,82 | 4681 |
| chrUextra | 28894807 | 28898891 | 4085 | 6,25 | 4682 |
| chrUextra | 28899301 | 28902831 | 3531 | 4,2 | 4683 |
| chrUextra | 28906983 | 28908135 | 1153 | 4,24 | 4684 |
| chrUextra | 28909009 | 28912533 | 3525 | 5,03 | 4685 |
| chrUextra | 28913483 | 28914029 | 547 | 3,72 | 4686 |
| chrUextra | 28914818 | 28916513 | 1696 | 5,61 | 4687 |
| chrUextra | 28916585 | 28918218 | 1634 | 3,33 | 4688 |
| chrUextra | 28918298 | 28920078 | 1781 | 6,28 | 4689 |
| chrUextra | 28920862 | 28921400 | 539 | 5,01 | 4690 |
| chrUextra | 28928492 | 28928986 | 495 | 4,11 | 4691 |
| chrUextra | 28930811 | 28933928 | 3118 | 5,6 | 4692 |
| chrUextra | 28934406 | 28934970 | 565 | 2,85 | 4693 |
| chrUextra | 28935387 | 28936327 | 941 | 4,01 | 4694 |
| chrUextra | 28936641 | 28937945 | 1305 | 6,28 | 4695 |
| chrUextra | 28939968 | 28944158 | 4191 | 5,01 | 4696 |
| chrUextra | 28945027 | 28945630 | 604 | 5,05 | 4697 |
| chrUextra | 28949800 | 28950363 | 564 | 4,24 | 4698 |
| chrUextra | 28951969 | 28952671 | 703 | 3,98 | 4699 |
| chrUextra | 28957967 | 28959694 | 1728 | 5,05 | 4700 |
| chrUextra | 28960041 | 28962294 | 2254 | 4,85 | 4701 |

| | | | | | |
|---|---|---|---|---|---|
| chrUextra | 28964209 | 28966029 | 1821 | 4,79 | 4702 |
| chrUextra | 28966402 | 28967773 | 1372 | 5,08 | 4703 |
| chrUextra | 28968635 | 28969445 | 811 | 5,08 | 4704 |
| chrUextra | 28969766 | 28971628 | 1863 | 3,49 | 4705 |
| chrUextra | 28972323 | 28974740 | 2418 | 5,51 | 4706 |
| chrUextra | 28977047 | 28979072 | 2026 | 4,37 | 4707 |
| chrUextra | 28979196 | 28983536 | 4341 | 4,79 | 4708 |
| chrUextra | 28983647 | 28984464 | 818 | 3,62 | 4709 |
| chrUextra | 28988331 | 28990002 | 1672 | 4,17 | 4710 |
| chrUextra | 28990190 | 28991896 | 1707 | 4,01 | 4711 |
| chrUextra | 28992035 | 28992802 | 768 | 3,43 | 4712 |
| chrUextra | 28992903 | 28993672 | 770 | 2,88 | 4713 |
| chrUextra | 28993823 | 28996935 | 3113 | 4,43 | 4714 |
| chrUextra | 28997092 | 28998748 | 1657 | 4,4 | 4715 |
| chrUextra | 28999850 | 29002421 | 2572 | 3,52 | 4716 |
| chrUextra | 29002500 | 29003780 | 1281 | 3,33 | 4717 |
| chrX | 213862 | 215853 | 1992 | 2,01 | 4718 |
| chrX | 1251885 | 1253529 | 1645 | 2,97 | 4719 |
| chrX | 1258062 | 1262114 | 4053 | 2,72 | 4720 |
| chrX | 1824846 | 1831369 | 6524 | 2,6 | 4721 |
| chrX | 2504383 | 2510279 | 5897 | 2,66 | 4722 |
| chrX | 2683834 | 2687721 | 3888 | 11,39 | 4723 |
| chrX | 2689560 | 2691274 | 1715 | 11,16 | 4724 |
| chrX | 2969622 | 2974756 | 5135 | 1,74 | 4725 |
| chrX | 3073224 | 3078790 | 5567 | 2,06 | 4726 |
| chrX | 4990050 | 4998248 | 8199 | 2 | 4727 |
| chrX | 6068913 | 6070668 | 1756 | 2,1 | 4728 |
| chrX | 6324099 | 6325929 | 1831 | 1,97 | 4729 |
| chrX | 6917943 | 6924248 | 6306 | 2,79 | 4730 |
| chrX | 10991315 | 10996770 | 5456 | 2,78 | 4731 |
| chrX | 11080543 | 11081996 | 1454 | 2,08 | 4732 |
| chrX | 12276854 | 12278571 | 1718 | 2,13 | 4733 |
| chrX | 12824543 | 12830499 | 5957 | 3,38 | 4734 |
| chrX | 13940380 | 13955882 | 15503 | 2,57 | 4735 |
| chrX | 14446149 | 14452759 | 6611 | 2,21 | 4736 |
| chrX | 20151018 | 20154024 | 3007 | 2,18 | 4737 |
| chrX | 21329712 | 21335296 | 5585 | 2,77 | 4738 |
| chrX | 21342291 | 21343763 | 1473 | 2,07 | 4739 |
| chrX | 21393252 | 21394371 | 1120 | 2,43 | 4740 |
| chrX | 21673983 | 21677444 | 3462 | 2,65 | 4741 |
| chrX | 21678505 | 21680170 | 1666 | 2,07 | 4742 |
| chrX | 21681326 | 21683889 | 2564 | 3,95 | 4743 |
| chrX | 21764614 | 21765736 | 1123 | 2,69 | 4744 |
| chrX | 21834360 | 21835496 | 1137 | 6,21 | 4745 |
| chrXHet | 1 | 1211 | 1211 | 3,2 | 4746 |
| chrXHet | 1394 | 3056 | 1663 | 2,1 | 4747 |
| chrYHet | 4435 | 4901 | 467 | 3,69 | 4748 |
| chrYHet | 71378 | 76698 | 5321 | 1,88 | 4749 |

| | | | | | |
|---|---|---|---|---|---|
| chrYHet | 114028 | 115285 | 1258 | 1,84 | 4750 |
| chrYHet | 136720 | 137633 | 914 | 5,37 | 4751 |

**Chromatin State**
9
2
9
None
None
2
9
2
None
2
9
2
9
6
2
4
9
2
None
9
2
3
9
9
2
9
None
None
None
None
None
None
None
None
None
None
None
7
7
7
7
7
2,7
7
7

7
1
2,7
7
7
7
7
7
7
7
7
7
7
7
7
7
7
7
7
7
7
7
7
7
7
7
7
7
7
7,8
1,7
7
7
7
7
7
7
7
7
7
7
7
2,7
7
7,9
7
7
7
7

7
7
7
7
7
7
7
7
7
7
7
7
7
7
7
7
7
None
7
7
7
7
7
7
7
7
7
7
7
7
7
7
7
7
7
7
7
7
7
7
7
2,7
7
7
7
7
7
7

7
7
1,7
1,7
7
7
7
7
7
7
7
7
7
1,2,7
7
7
7
7
7
7
7
7
7
7
7
7
7
7
7
7
7
7
7
7
7
7
7
7
7
7
None
7
7
7
7
7
7
1,7

1
7
7
7
7
7
7
7
7
None
7
None
7
7
7
7
7
7
7
7
7
7
7
7
7
7
7
7
7
7
7
7
7
7
7
7
7
7
7
7
7
7
7
7
7
7
7
7
7
7

7
7
7
7
7
7
7
7
7
7
7
7
1,7
1
7,8
7
7
7
7
7
7
7
7
7
None
7
7
7
7
7
7
7
7
7
7
7
7
7
7
7
7
7
7
7
7
7
7
7

7
7
7
7
7
7
7
7
8
7
7
7
7
7
7
7
7
7
7
7
7
7
7
7
7
7
7
7
7
7
7
7
7
7
7
9
7
7
7
7
7
7
7
7
7
7
7
7

7
7
7
7
7
2
2
7
7
7
7
7
7
7
7
7
7
7
7
7
7
7
7
7
7
7
7
7
7
7
7
7
7
7
7
7
7
7
7
7
7
7
7
7
7
7
7
7

7
7
7
7
7
7
None
2,7
2,7
2,7
2
2
2
2
9
7,8,9
7,8,9
2
3
9
2
4
4
4
2
2,7
1,9
9
None
3,4
9
None
None
None
None
None
None
None
None
None
None
None
None
None
None
None
None
None

None
None
None
None
7
7
7
7
7
7
7
7
7
7
7
7
7
7
7
7
7
7
7
7
7
7
7
7
7
7
7
7
7
7
7
7
7
7
7
7
7
7
7
7
7
7
7
7

7
7
7
7
7
7
7
7
7
7
7
7
7
7
7
7
7
7
7
7
7
7
7
7
7
7
7
7
7
7
7
7
7
7
7
7
7
7
7
7
7
7
7
7
7
7
7
7

7
7
7
7
7
7
7
7
7
7
7
7
7
7
7
None
7
7
7
7
7
7
7
7
7
7
7
7
7
7
7
7
7
7
7
7
7
7
7
7
7
7
7
7
7
7
7
7

7
7
7
7
None
1,7
7
7
7
7
7
7
7
7
7
7
7
7
7
7
7
7
7
7
3,7
7
7
7
7
7
7
7
7
7
7
7
7
7
7
7
7
7
7
7
7
7
7
7

7
7
7
7
7
7
7
7
7
7
7
7
7
7
7
7
7
7
7
7
7
7
7
7
7
7
7
7
7
7
7
7
7
7
7
7
7
7
7
7
7
7
7
7
7
7
7
7

7
7
7
7
7
7
7
7
7
7
7
7
7,8
7
7
7
7
7
7
7
7
7
7
7
7
7
7
7
7
7
7
7
7
7
7
7
7
7
7
7
7
7
7
7
7
7
7
8

7
7
7
7
7
7
7
7
7
7
7
7
7
7
7
7
7
7
7
7
7
7
7
7
7
7
7
7
7
7
7
7
7
7
7
7
7
7
7
7
7
7
7
7
7
7
7
7

7
7
7
7
7
7
7
7
7
7
7
7
7
7
7
2,7
2,7
None
4
2
2
2
2
9
9
2,7,8
None
None
None
9
9
2
3
9
2,8,9
2
4
2
2
7,8
7
7,8
7
7,8
7,8
7
7
7

7,8
7
7
7,8
7
7
7
7
7,8
8
8
7,8
7,8
7,8
7,8
7,8
7
7
7,8
7
7,8
7
2
2
7
7
7
7
7
7
7
7
7
7
7
7,8
7
7
7
7
7
7
7
7
7
7
7
7

7
7
7
7
7
7
7
7
7
7
7
7
7
7
7
7
7
1,7
7
7
7,9
7
7
7
7
7
7
7
7
7
7
7
7
7
7
7
7
7
7
7
7
7
7
7
7
7
7
7

7
7
7
7
7
7
7
7
7
7
7
7
7
7
7
7
7
7
7
7
7
7
7
7
7
7
7
7
7
7
7
7
7
7
7
7
7
7
7
7
7
7
7
7
7
7
7
7,8

7
7
7
7
7
7
7
7
7
7
7
7
7
7
7
7
7
7
7
7
7
7
7
7
7
7
7
7
7
7
7
7
7
7
7
7
7
7
7
7
7
7
7
7
7
7
7
7

7
7
7
7
7
7
7
7
7
7
7
7
7
7
7
7
7
7
7
7
7
7
7
7
7
7
7
7
7
7
7
7
7
7
7
7
7
7
7
7
7
7
7
7
7
7
7
7

7
7
7
7
7
7
7
7
7
7
7
7
7
7
7
7
7
7
7
7
7
7
7
7
7
7
7
9
7
7
7
7
7
7
7
7
7
7
7
7
7
7
7
7
7
7
7
7

7
7
7
7
7
7
7
7
7
1,7
7
7
7
7
7
7
7
7
7
7
7
7
7
7
7
7
7
7
7
7
7
7
7
7
7
7
7
7
7
7
7
7
7
7
7
7
7
7
7

7
7
7
7
7,8
7
7
7
7
7
7
7
7
7
7
7
7
7
7
7
7
7
7
7
7
7
7
7
7
7
7
7
7
7
7
7
7
7
7
7
7
7
7
7
7
7
7
7

7
7
7
7
7
7
7
7
7
7
7
7
7
7
7
7
7
7
7
7
7
7
7
7
7
7
7
7
7
7
7
7
7
7
7
7
7
7
7
7
7
7
7
7
7
7
7
7

7
7
7
7
7
7
7
7
7
7
7
7
7
7
7
7
7
7
7
7
7
7
7
7
7
7
7
7
7
7
7
7
7
7
7
7
7
7
7
7
7
7
7
7
7
7
7
7

7
7
7
7
7
7
7
7
7
7
7
7
7
7
7
7
7
7
7
7
7
7
7
7
7
7
7
7
7
7
7
7
7
7
7
7
7
7
7
None
None
None
None
7
7
7
7
7

7
7
7
7
7
7
7
7
7
7
7
7
7
7
7
7
7
7
7
7
7
7
7
7
7
7
7
9
2,7
4
9
2
None
4
4
2
2
2
2,7
2,7
None
4
9
2
9
9
4
2

9
6,9
9
7,8
9
9
9
9
9
9
9
9
9
9
9
9
9
9
9
9
9
1,2,7
2
None
7
7
7
7
7
7
7
7
7
7
7
7
7
7
7
7
7
7
7
7
7
7
7
7

7
7
7
7
7
7
7
7
7
7
7
7
7
7
7
7
7
7
7
7
7
7
7
7
7
7
7
7
7,8
7
7
7
7
7
7
7
7
7
7
7
7,8
7
7
7
7
7
7,9
7

7
7
7
7
7
7
7
7
7
7
7
7
7
7
7
7
7
7
7
7
7
7
7
7
7
7
7
7
7
7
7
7
7
7
7
7
7
7
7
7
7
7
7
7
7
7
7
7

7
7
7
7
7
7
7
7
7
7
7
7
7
7
7
7
7
7
7
7
7
7
7
7
7
7
7
7
7
7
7
7
7
7
7
7
7
7
7
7
7
7
7
7
7
7
7
7

7
7
7
7
7
7
7
7
7
7
7
7
7
7
7
7
7
7
7
7
7
7
7
7
7
7
7
7
7
7
7
7
7
7
7
7
7
7
7
7
7
7
7
7
7
7
7
7

7
7
7
7
7
7
7
7
7
7
7
7
7
7
7
7
7
7,8
7
7
7
7
7
7
7
7
7
7
7
7
7
7
7
7
7
7
7
7
7
7
7
7
7
7
7
7
7
7

7
7
7
7
7
7
7
7
7
7
7
7
7
9
7
7
7
7
7
7
7
7,8
8
7
7
7
7
7
7
7
7
7
7
7
7
7
7
7
7
7
7
7
7
7
7
7
7
7

7
7
7
7
7
7
7
7
7
7
7
7
7
7
7
7
7
7
7,8
7,8
7
7
7
7
7
7
7
7
7
7
7
7
7
7
None
None
None
None
None
None
None
None
None
None
None
None
None
None

None
None
None
None
None
None
None
None
None
None
None
None
None
None
None
None
None
None
None
None
None
None
None
None
None
None
None
None
None
None
None
None
None
None
None
None
None
None
None
None
None
None
None
None
None
None
None
None

None
None
None
None
None
None
None
None
None
None
None
None
None
None
None
None
None
None
None
None
None
None
None
None
None
None
None
None
None
None
None
None
None
None
None
None
None
None
None
None
None
None
None
None
None
None
None
None

None
None
None
None
None
None
None
None
None
None
None
None
None
None
None
None
None
None
None
None
None
None
None
None
None
None
None
None
None
None
None
None
None
None
None
None
None
None
None
None
None
None
None
None
None
None
None
None

None
None
None
None
None
None
None
None
None
None
None
None
None
None
None
None
None
None
None
None
None
None
None
None
None
None
None
None
None
None
None
None
None
None
None
None
None
None
None
None
None
None
None
None
None
None
None
None

None
None
None
None
None
None
None
None
None
None
None
None
None
None
None
None
None
None
None
None
None
None
None
None
None
None
None
None
None
None
None
None
None
None
None
None
None
None
None
None
None
None
None
None
None
None
None
None

None
None
None
None
None
None
None
None
None
None
None
None
None
None
None
None
None
None
None
None
None
None
None
None
None
None
None
None
None
None
None
None
None
None
None
None
None
None
None
None
None
None
None
None
None
None
None
None

None
None
None
None
None
None
None
None
None
None
None
None
None
None
None
None
None
None
None
None
None
None
None
None
None
None
None
None
None
None
None
None
None
None
None
None
None
None
None
None
None
None
None
None
None
None
None
None

None
None
None
None
None
None
None
None
None
None
None
None
None
None
None
None
None
None
None
None
None
None
None
None
None
None
None
None
None
None
None
None
None
None
None
None
None
None
None
None
None
None
None
None
None
None
None
None

None
None
None
None
None
None
None
None
None
None
None
None
None
None
None
None
None
None
None
None
None
None
None
None
None
None
None
None
None
None
None
None
None
None
None
None
None
None
None
None
None
None
None
None
None
None
None
None

None
None
None
None
None
None
None
None
None
None
None
None
None
None
None
None
None
None
None
None
None
None
None
None
None
None
None
None
None
None
None
None
None
None
None
None
None
None
None
None
None
None
None
None
None
None
None
None

None
None
None
None
None
None
None
None
None
None
None
None
None
None
None
None
None
None
None
None
None
None
None
None
None
None
None
None
None
None
None
None
None
None
None
None
None
None
None
None
None
None
None
None
None
None
None
None

None
None
None
None
None
None
None
None
None
None
None
None
None
None
None
None
None
None
None
None
None
None
None
None
None
None
None
None
None
None
None
None
None
None
None
None
None
None
None
None
None
None
None
None
None
None
None
None

None
None
None
None
None
None
None
None
None
None
None
None
None
None
None
None
None
None
None
None
None
None
None
None
None
None
None
None
None
None
None
None
None
None
None
None
None
None
None
None
None
None
None
None
None
None
None
None

None
None
None
None
None
None
None
None
None
None
None
None
None
None
None
None
None
None
None
None
None
None
None
None
None
None
None
None
None
None
None
None
None
None
None
None
None
None
None
None
None
None
None
None
None
None
None
None

None
None
None
None
None
None
None
None
None
None
None
None
None
None
None
None
None
None
None
None
None
None
None
None
None
None
None
None
None
None
None
None
None
None
None
None
None
None
None
None
None
None
None
None
None
None
None
None

None
None
None
None
None
None
None
None
None
None
None
None
None
None
None
None
None
None
None
None
None
None
None
None
None
None
None
None
None
None
None
None
None
None
None
None
None
None
None
None
None
None
None
None
None
None
None
None

None
None
None
None
None
None
None
None
None
None
None
None
None
None
None
None
None
None
None
None
None
None
None
None
None
None
None
None
None
None
None
None
None
None
None
None
None
None
None
None
None
None
None
None
None
None
None
None

None
None
None
None
None
None
None
None
None
None
None
None
None
None
None
None
None
None
None
None
None
None
None
None
None
None
None
None
None
None
None
None
None
None
None
None
None
None
None
None
None
None
None
None
None
None
None
None

None
None
None
None
None
None
None
None
None
None
None
None
None
None
None
None
None
None
None
None
None
None
None
None
None
None
None
None
None
None
None
None
None
None
None
None
None
None
None
None
None
None
None
None
None
None
None
None

None
None
None
None
None
None
None
None
None
None
None
None
None
None
None
None
None
None
None
None
None
None
None
None
None
None
None
None
None
None
None
None
None
None
None
None
None
None
None
None
None
None
None
None
None
None
None
None

None
None
None
None
None
None
None
None
None
None
None
None
None
None
None
None
None
None
None
None
None
None
None
None
None
None
None
None
None
None
None
None
None
None
None
None
None
None
None
None
None
None
None
None
None
None
None
None

None
None
None
None
None
None
None
None
None
None
None
None
None
None
None
None
None
None
None
None
None
None
None
None
None
None
None
None
None
None
None
None
None
None
None
None
None
None
None
None
None
None
None
None
None
None
None
None

None
None
None
None
None
None
None
None
None
None
None
None
None
None
None
None
None
None
None
None
None
None
None
None
None
None
None
None
None
None
None
None
None
None
None
None
None
None
None
None
None
None
None
None
None
None
None
None

None
None
None
None
None
None
None
None
None
None
None
None
None
None
None
None
None
None
None
None
None
None
None
None
None
None
None
None
None
None
None
None
None
None
None
None
None
None
None
None
None
None
None
None
None
None
None
None

None
None
None
None
None
None
None
None
None
None
None
None
None
None
None
None
None
None
None
None
None
None
None
None
None
None
None
None
None
None
None
None
None
None
None
None
None
None
None
None
None
None
None
None
None
None
None
None

None
None
None
None
None
None
None
None
None
None
None
None
None
None
None
None
None
None
None
None
None
None
None
None
None
None
None
None
None
None
None
None
None
None
None
None
None
None
None
None
None
None
None
None
None
None
None
None

None
None
None
None
None
None
None
None
None
None
None
None
None
None
None
None
None
None
None
None
None
None
None
None
None
None
None
None
None
None
None
None
None
None
None
None
None
None
None
None
None
None
None
None
None
None
None
None

None
None
None
None
None
None
None
None
None
None
None
None
None
None
None
None
None
None
None
None
None
None
None
None
None
None
None
None
None
None
None
None
None
None
None
None
None
None
None
None
None
None
None
None
None
None
None
None

None
None
None
None
None
None
None
None
None
None
None
None
None
None
None
None
None
None
None
None
None
None
None
None
None
None
None
None
None
None
None
None
None
None
None
None
None
None
None
None
None
None
None
None
None
None
None
None

None
None
None
None
None
None
None
None
None
None
None
None
None
None
None
None
None
None
None
None
None
None
None
None
None
None
None
None
None
None
None
None
None
None
None
None
None
None
None
None
None
None
None
None
None
None
None
None

None
None
None
None
None
None
None
None
None
None
None
None
None
None
None
None
None
None
None
None
None
None
None
None
None
None
None
None
None
None
None
None
None
None
None
None
None
None
None
None
None
None
None
None
None
None
None
None

None
None
None
None
None
None
None
None
None
None
None
None
None
None
None
None
None
None
None
None
None
None
None
None
None
None
None
None
None
None
None
None
None
None
None
None
None
None
None
None
None
None
None
None
None
None
None
None

None
None
None
None
None
None
None
None
None
None
None
None
None
None
None
None
None
None
None
None
None
None
None
None
None
None
None
None
None
None
None
None
None
None
None
None
None
None
None
None
None
None
None
None
None
None
None
None

None
None
None
None
None
None
None
None
None
None
None
None
None
None
None
None
None
None
None
None
None
None
None
None
None
None
None
None
None
None
None
None
None
None
None
None
None
None
None
None
None
None
None
None
None
None
None
None

None
None
None
None
None
None
None
None
None
None
None
None
None
None
None
None
None
None
None
None
None
None
None
None
None
None
None
None
None
None
None
None
None
None
None
None
None
None
None
None
None
None
None
None
None
None
None
None

None
None
None
None
None
None
None
None
None
None
None
None
None
None
None
None
None
None
None
None
None
None
None
None
None
None
None
None
None
None
None
None
None
None
None
None
None
None
None
None
None
None
None
None
None
None
None
None

None
None
None
None
None
None
None
None
None
None
None
None
None
None
None
None
None
None
None
None
None
None
None
None
None
None
None
None
None
None
None
None
None
None
None
None
None
None
None
None
None
None
None
None
None
None
None
None

None
None
None
None
None
None
None
None
None
None
None
None
None
None
None
None
None
None
None
None
None
None
None
None
None
None
None
None
None
None
None
None
None
None
None
None
None
None
None
None
None
None
None
None
None
None
None
None

None
None
None
None
None
None
None
None
None
None
None
None
None
None
None
None
None
None
None
None
None
None
None
None
None
None
None
None
None
None
None
None
None
None
None
None
None
None
None
None
None
None
None
None
None
None
None
None

None
None
None
None
None
None
None
None
None
None
None
None
None
None
None
None
None
None
None
None
None
None
None
None
None
None
None
None
None
None
None
None
None
None
None
None
None
None
None
None
None
None
None
None
None
None
None
None

None
None
None
None
None
None
None
None
None
None
None
None
None
None
None
None
None
None
None
None
None
None
None
None
None
None
None
None
None
None
None
None
None
None
None
None
None
None
None
None
None
None
None
None
None
None
None
None

None
None
None
None
None
None
None
None
None
None
None
None
None
None
None
None
None
None
None
None
None
None
None
None
None
None
None
None
None
None
None
None
None
None
None
None
None
None
None
None
None
None
None
None
None
None
None
None

None
None
None
None
None
None
None
None
None
None
None
None
None
None
None
None
None
None
None
None
None
None
None
None
None
None
None
None
None
None
None
None
None
None
None
None
None
None
None
None
None
None
None
None
None
None
None
None

None
None
None
None
None
None
None
None
None
None
None
None
None
None
None
None
None
None
None
None
None
None
None
None
None
None
None
None
None
None
None
None
None
None
None
None
None
None
None
None
None
None
None
None
None
None
None
None

None
None
None
None
None
None
None
None
None
None
None
None
None
None
None
None
None
None
None
None
None
None
None
None
None
None
None
None
None
None
None
None
None
None
None
None
None
None
None
None
None
None
None
None
None
None
None
None

None
None
None
None
None
None
None
None
None
None
None
None
None
None
None
None
None
None
None
None
None
None
None
None
None
None
None
None
None
None
None
None
None
None
None
None
None
None
None
None
None
None
None
None
None
None
None
None

None
None
None
None
None
None
None
None
None
None
None
None
None
None
None
None
None
None
None
None
None
None
None
None
None
None
None
None
None
None
None
None
None
None
None
None
None
None
None
None
None
None
None
None
None
None
None
None

None
None
None
None
None
None
None
None
None
None
None
None
None
None
None
None
None
None
None
None
None
None
None
None
None
None
None
None
None
None
None
None
None
None
None
None
None
None
None
None
None
None
None
None
None
None
None
None

None
None
None
None
None
None
None
None
None
None
None
None
None
None
None
None
None
None
None
None
None
None
None
None
None
None
None
None
None
None
None
None
None
None
None
None
None
None
None
None
None
None
None
None
None
None
None
None

None
None
None
None
None
None
None
None
None
None
None
None
None
None
None
None
None
None
None
None
None
None
None
None
None
None
None
None
None
None
None
None
None
None
None
None
None
None
None
None
None
None
None
None
None
None
None
None

None
None
None
None
None
None
None
None
None
None
None
None
None
None
None
None
None
None
None
None
None
None
None
None
None
None
None
None
None
None
None
None
None
None
None
None
None
None
None
None
None
None
None
None
None
None
None
None

None
None
None
None
None
None
None
None
None
None
None
None
None
None
None
None
None
None
None
None
None
None
None
None
None
None
None
None
None
None
None
None
None
None
None
None
None
None
None
None
None
None
None
None
None
None
None
None

None
None
None
None
None
None
None
None
None
None
None
None
None
None
None
None
None
None
None
None
None
None
None
None
None
None
None
None
None
None
None
None
None
None
None
None
None
None
None
None
None
None
None
None
None
None
None
None

None
None
None
None
None
None
None
None
None
None
None
None
None
None
None
None
None
None
None
None
None
None
None
None
None
None
None
None
None
None
None
None
None
None
None
None
None
None
None
None
None
None
None
None
None
None
None
None

None
None
None
None
None
None
None
None
None
None
None
None
None
None
None
None
None
None
None
None
None
None
None
None
None
None
None
None
None
None
None
None
None
None
None
None
None
None
None
None
None
None
None
None
None
None
None
None

None
None
None
None
None
None
None
None
None
None
None
None
None
None
None
None
None
None
None
None
None
None
None
None
None
None
None
None
None
None
None
None
None
None
None
None
None
None
None
None
None
None
None
None
None
None
None
None

None
None
None
None
None
None
None
None
None
None
None
None
None
None
None
None
None
None
None
None
None
None
None
None
None
None
None
None
None
None
None
None
None
None
None
None
None
None
None
None
None
None
None
None
None
None
None
None

None
None
None
None
None
None
None
None
None
None
None
None
None
None
None
None
None
None
None
None
None
None
None
None
None
None
None
None
None
None
None
None
None
None
None
None
None
None
None
None
None
None
None
None
None
None
None
None

None
None
None
None
None
None
None
None
None
None
None
None
None
None
None
None
None
None
None
None
None
None
None
None
None
None
None
None
None
None
None
None
None
None
None
None
None
None
None
None
None
None
None
None
None
None
None
None

None
None
None
None
None
None
None
None
None
None
None
None
None
None
None
None
None
None
None
None
None
None
None
None
None
None
None
None
None
None
None
None
None
None
None
None
None
None
None
None
None
None
None
None
None
None
None
None

None
None
None
None
None
None
None
None
None
None
None
None
None
None
None
None
None
None
None
None
None
None
None
None
None
None
None
None
None
None
None
None
None
None
None
None
None
None
None
None
None
None
None
None
None
None
None
None

None
None
None
None
None
None
None
None
None
None
None
None
None
None
None
None
4
1
1
5
1
3,5,6
5
9
9
9
9
None
9
5
9
9
8
9
9
8
8
8
7
7
7
7
7
8
7
7
7
9
